# Scale space calibrates present and subsequent spatial learning in Barnes maze in mice

**DOI:** 10.1101/2022.12.14.520510

**Authors:** Yuto Tachiki, Yusuke Suzuki, Mutsumi Kurahashi, Keisuke Oki, Özgün Mavuk, Takuma Nakagawa, Shogo Ishihara, Yuichiro Gyoten, Akira Yamamoto, Itaru Imayoshi

**Author notes:** Correspondence Correspondence should be addressed to Yusuke Suzuki II at and Itaru Imayoshi at. These authors equally contributed to this work (Shogo Ishihara, Yuichiro Gyoten, Akira Yamamoto). Author Contributions Y.T.,Y.S.II. and I.I. Designed Research; Y.T.,Y.S.II., M.K., K.O., Ö.M., T.N., S.I., Y.G., A.Y. Performed Research; Y.T., Y.S.II. Analyzed data; Y.T.,Y.S.II., I.I. Wrote the paper.

## Abstract

Animals including humans are capable of representing different scale spaces from smaller to larger ones. However, most laboratory animals live their life in a narrow range of scale spaces like home-cages and experimental setups, making it hard to extrapolate the spatial representation and learning process in large scale spaces from those in conventional scale spaces. Here, we developed a 3-meter diameter Barnes maze (BM3), then explored whether spatial learning in Barnes maze (BM) is calibrated by scale spaces. In the BM3, mice exhibited lower learning rate compared to a conventional 1-meter diameter Barnes maze (BM1), suggesting that the BM3 requires more trial-and-error and larger computational resources to solve the task than the BM1. Analyzing network structures of moving trajectories, betweenness centrality would contrast spatial learning in a larger scale space with that in a smaller one, as it diverges between the BM1 and the BM3 along with the learning progression. We then explored whether prior learning in either BM scale calibrates subsequent spatial learning in the other BM scale, and found asymmetric facilitation such that the prior learning in the BM3 facilitated the subsequent learning in the BM1, but not *vice versa*. Network structures of trajectories in the subsequent BM scale were changed by both prior and subsequent BM scale. These results suggest that scale space calibrates both the present and subsequent BM learning. This is the first study to explore and demonstrate scale-dependent spatial learning in Barnes maze in mice.

**Significance Statement:** Animals are capable of representing different scale spaces. However, whether scale space calibrates goal-directed spatial learning remains unclear. The Barnes maze is a well-established experimental paradigm to evaluate spatial learning in rodents. Here, we developed a larger scale 3-meter diameter Barnes maze (BM3) then compared various navigation features in mice between the BM3 and a conventional 1-meter diameter Barnes maze (BM1). We demonstrated that learning on the BM3 required more computational resources than in the BM1, prompting mice to exploit unique navigation patterns. Such learning experiences in the BM3 facilitated subsequent spatial learning in the BM1, but not *vice versa*. These results suggest that scale space calibrates immediate and subsequent spatial learning.

## Introduction

Animals including humans are capable of representing different scale spaces from smaller to larger ones. For example, bats and wild rodents can navigate from the order of centimeters of detail in the vicinity of burrows and feeding grounds, to the order of kilometers between their burrow and feeding grounds (Geva-Sagiv et al., 2015). Understanding how animals acquire spatial representations over different scale spaces and how to incorporate them into a *cognitive map* are close to the nature of spatial representation and learning. However, most laboratory animals live their life in a narrow range of scale space, between home-cages and experimental setups. Therefore, it is hard to extrapolate the spatial representation and learning process in large scale spaces from those in conventional scale spaces.

Several types of neurons act as a unit interactively accounting for a cognitive map (Knierim et al., 1998). For example, hippocampal place cells encode one or more places termed place fields where they fired with the spatial resolution of dozens of centimeters order when rodents were located in the places (O’Keefe & Conway, 1978). Grid cells in the medial entorhinal cortex are active when the animal passes the apex of a hexagonal grid over its environment (Hafting et al., 2005). Compared to place cells, spatial resolution of grid cells widely ranges from about dozens of centimeters to several meters (Brun et al., 2008).

However, it is unlikely that any scale spaces are represented at a single, common spatial resolution; if one were to try to perform navigation on the order of kilometers with place cells of centimeter resolution, the number of required spatially selective neurons would be larger than the total number of neurons in a brain. Rather, spatial resolution would be different per scale space to be represented, or each place cell would have multiple or enlarged place fields (Geva-Sagiv et al., 2015). Several studies demonstrated that spatial scale in the environment controls the number and the size of place fields of place cells (Muller & Kubie, 1987; O’Keefe & Burgess, 1996; Fenton et al., 2008; Kjelstrup et al., 2008; Park et al., 2011; Rich et al., 2014; Harland et al., 2021; Sarel et al., 2022). Though these studies suggested the existence of scale-dependent spatial representation, experimental systems or paradigms to test them are yet to be developed.

The Barnes circular maze test (BM) was originally developed by Carol A. Barnes in 1979 (Barnes, 1979), and nowadays is one of well-established experimental paradigms to test spatial learning and memory in rodents. Briefly, a mouse was released in a bright, circular open field with 12 holes equally spaced along with the edge of the field. One escape box is attached under any one of the holes. The mouse can escape from the field by entering the escape box, which is the “goal” in this maze. Because the location of the goal is fixed per mouse, it can optimize navigation from start to goal, by updating spatial representation of the maze across repeated training. It is well documented that the BM learning depends on the hippocampus, as a number of studies reported that hippocampal lesioned rodents exhibited impaired spatial memory and navigation in the BM task (Bach et al., 1995; Mayford et al., 1996; Raber et al., 2004). Conventional Barnes maze test is performed on a field varying from 70 to 130 cm in diameter (Bach et al., 1995; Raber et al., 2004; Rosenfeld & Ferguson, 2014; Pitts, 2018), and no studies compare spatial learning between Barnes mazes of different scale spaces.

Here, we developed a 3-meter diameter Barnes maze (BM3) which is 3 times larger than a conventional 1-meter diameter Barnes maze (BM1). Comparing a variety of behavioral features in mice in the BM3 with those in the BM1, we examined whether spatial learning in Barnes maze (BM) is calibrated by scale spaces. We also explore whether prior learning in either BM scale facilitates subsequent spatial learning in the other BM scale. It has been widely accepted that animals learn not only solutions of an immediate task per se, but also how to find the solutions (learning to learn; Harlow, 1949). This meta-learning process leads to few-shot learning in future tasks that are variants of the previously learned task (for review, see Wang, 2021). To explore whether the meta-learning process is calibrated by scale spaces, facilitation effects from prior learning in either BM scale (e.g. BM3) to subsequent spatial learning in the other BM scale (e.g. BM1) were evaluated.

## Material and Methods

### Animal

111 naive male C57BL/6J mice (Japan SLC, Shizuoka, Japan) were used in this study (**Table 1**). All mice were 2∼3-months old (m.o) on the first day of the initial task. The mice were group-housed in a standard laboratory environment, maintained on a 12-h light/12-h dark cycle at 23–24°C temperature and 40–50% relative humidity. Food (pellets; Japan SLC) and water were provided *ad libitum*. The present behavioral testing was done during the dark phase. After the behavioral experiments, all mice were sacrificed by cervical dislocation. All animal procedures were performed in accordance with the [Author University] animal care committee’s regulations (permit numbers: Lif-K22008).

**Table 1.** Cohort and task instance n = number of mice in each cohort at instance 1. CFC = contextual fear conditioning. ^a^Scopolamine hydrobromide was injected 20 min before the beginning of the probe test. ^b^One mouse was dead in a blank between instance 1 and 2.

### Cohort

Each mouse was assigned to one of 7 cohorts, depending on the history of tasks that they were engaged in **Table 1**. Cohort 1 was engaged only in the 1-meter diameter maze (BM1). Cohorts 2 and 3 were engaged only in the 3-meter diameter maze (BM3), while Cohort 3 was injected with scopolamine hydrobromide (Santa Cruz Biotechnology, TX, USA) intraperitoneally 20 min before starting the probe test, where the scopolamine was prepared as a 0.3 mg/ml stock solution in 0.9% saline, so as to be 3 mg of scopolamine hydrobromide/kg of body weight. Cohorts 4 to 6 were engaged in the 2 tasks with the blank of 14 days between the tasks. Cohort 4 were engaged in the BM3 at the first task, then the BM1 at the second task. Cohort 5 was engaged in the modified BM1 (BM1’) at the first task, then the BM1 task at the second task; the BM1’ task was the same as the BM1 task (see above), except that the spatial cues were replaced by new independent ones, and that the training lasted for 12 days as much as that in the BM3. Cohort 6 were engaged in the BM3 then the BM1. Cohort S1 was engaged in the contextual fear conditioning at the first task, then the BM1 at the second task, with 30 days blank between the tasks.

### The 1-merter diameter Barnes maze (BM1)

#### Apparatus

The BM1 task was conducted on a custom-made Barnes maze system (Bio-Medica, Osaka, Japan) (**Fig. 1A, C, E**). Through the task, the mice learned to elaborate cognitive maps of the maze, and to take efficient navigation from the start to the goal on the maze. The maze was composed of a circular open arena with 98 cm diameter and 72 cm height from the floor. A start-lift was set at the center of the arena surface. The scaffold of the lift (10 cm diameter) was made of the same material as the arena and was held 20 cm below the surface of the arena before the initiation of each trial. At the start of each trial, the lift transported a mouse to the center of the arena surface. The vertical movement of the lift was programmed so that the experimenter could control it externally at any time. Twelve holes were equally spaced around the perimeter at a distance of 40 cm from the centroid where the diameter of each hole was 4 cm. A black iron escape box (17 × 14 × 7 cm), which had paper cage bedding on its bottom, was located under one of the holes. This hole is the goal of this navigation task, and the remaining holes were considered as dummies of the goal. The location of the goal was consistent for a given mouse, but was randomized across mice. The entire apparatus was set within a cube-shaped outer enclosure (130 × 130 × 180 cm) with black curtains in order to obscure the outside scene of the arena and absorb background noise. One white color projection LED light was mounted on the center of the ceiling of the enclosure to ensure uniform and intense illumination of the arena (600 lx on the arena surface). Unique 3D object as a spatial cue was set at each of 4 corners of the enclosure at a height of 86 cm from the floor.

**Figure 1.**
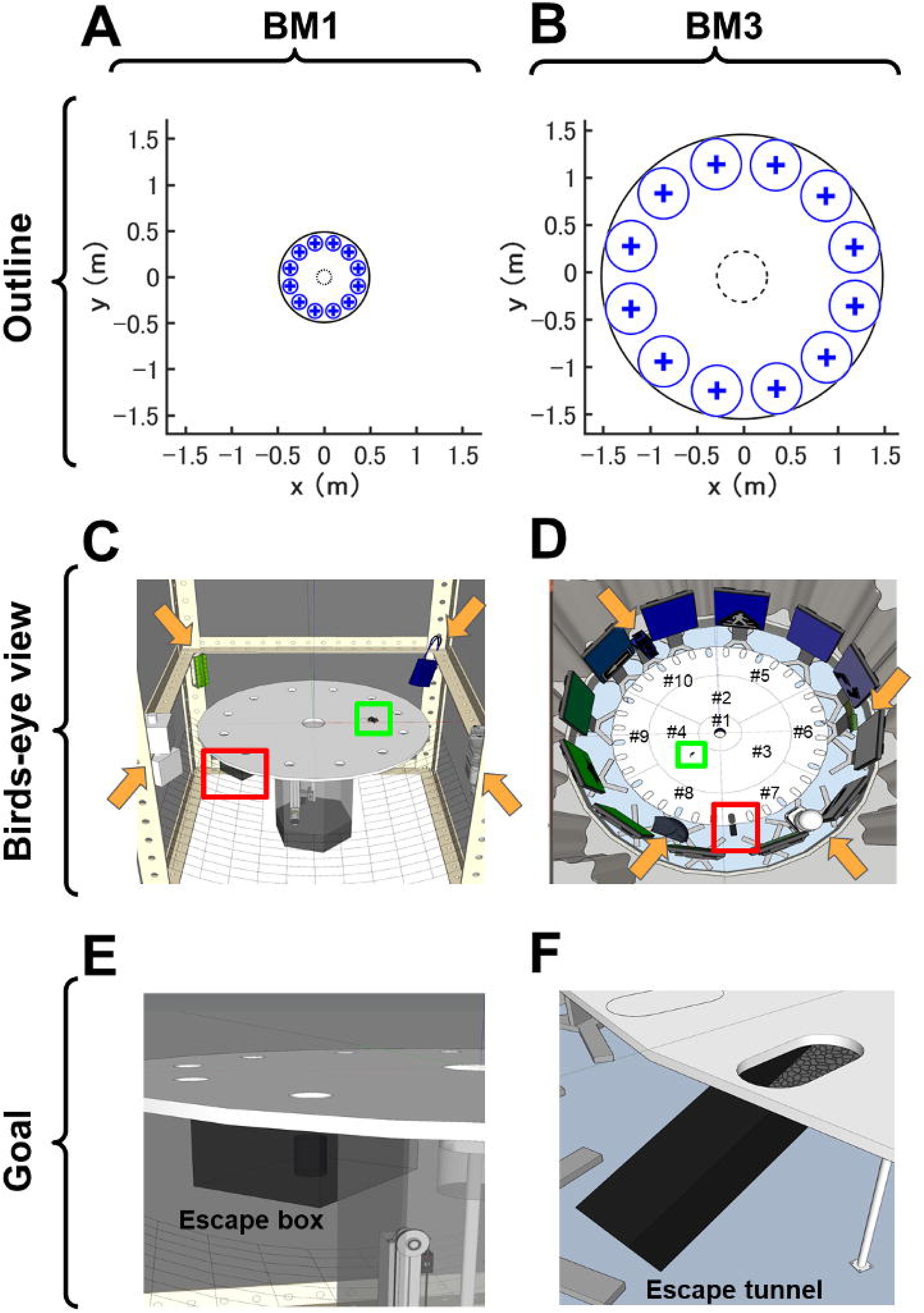
Architecture of the conventional 1-meter diameter Barnes maze (BM1) and the 3-meter diameter Barnes maze (BM3). ***A–F,*** The architecture of BM1 and BM3. Top, middle, and bottom rows indicate outlines (***A,B***), birds-eye views (***C,D***), and enlarged images of goals (***E,F***), respectively. In the outline, the display unit of vertical (*y*) and horizontal (*x*) axes in meters. The largest circle drawn by black solid line represents the edge of each maze. The “+” markers and blue-line circles around them represent the goal or dummy holes and the areas where hole visiting was scored, respectively. Namely, the number of errors was determined by the number that mice entered the blue-line circled areas until they goaled. Likewise, in the analysis of the probe test of spatial memory, time spent around each hole was the duration that mice stayed in the blue-line circled area. The black dashed-line circles represent the start areas with a lift transporting mice to the field. In strategy analysis, trajectory data while mice stayed in the start area since they entered the field were discarded, so that initial wobbly trajectory around the lift did not affect the classification of navigation strategy (see details in Material and Methods). ***C, D,*** In the birds-eye views, the orange arrows point to the locations of distal spatial cues. The red and green squares represent escape box/tunnel and mice, respectively. Each number (#) in ***D*** corresponds to each component of the BM3 arena. Escape box and tunnel were zoomed in ***E, F***, respectively.

Behavior during the trials was recorded using a GigE Vision camera (UI-5240SE-NIR; IDS Imaging Development Systems GmbH, Obersulm, Germany). The camera was mounted on the ceiling of the enclosure (93.5 cm above the maze center), which could achieve a spatial resolution of 1.96 mm/pixel. Each image frame (500 × 500 pixels, with ×2 binning) was acquired at the rising edge on a 20 Hz pulse counter and displayed on a monitor outside of the enclosure. All programs used for data acquisition, processing, saving, and synchronized device controls, were written in LabVIEW 2013 (National Instruments, TX, US).

#### Experimental procedure

The protocol of the BM1 task consists of three phases: habituation on Day 0, training from Days 1 to 6, and the probe test on Day 7. In all phases, mice were moved from the breeding rack to the experimental room 30 min before the experiment, and stayed in their home cages on a standby-rack in the experimental room while they were waiting for each trial. Immediately before the trial, the mouse was moved from the cage to the bottom of the lift via an opaque acrylic cylinder. At the start of the trial, the mouse was lifted up to the arena surface simultaneously with the start of recording. After every trial, the maze was thoroughly cleaned with 70% ethanol solution and dried. In the habituation phase, mice were allowed to freely explore the arena for 5 min. Afterwards they were moved to the escape box for 5 min, then they were returned to the home cage. This procedure was executed once per mouse. In the training phase, mice explored the arena until the mouse had successfully entered the escape box within 10 min. When the experimenter confirmed it, the recording was turned off, then brought the mouse back to the homecage. Otherwise, the experimenter picked up the mouse manually and returned it to the homecage by the cylinder. Trial was incremented by 1 if all mice completed the current trial, and 3 trials were performed in each training day. Twenty-four hours after the last day of training, mice were subjected to the probe test. In the probe test, mice explored the arena without the escape box for 5 min. This trial was done once per mouse.

### The 3-meter diameter Barnes maze (BM3)

#### Apparatus

The dedicated system for the BM3 was custom-built (Bio-Medica, Osaka, Japan) (**Fig. 1B, D, F**). The material of the circular arena was the same as the BM1, whereas the diameter was extended to 300 cm. The arena was constituted from 10 parts (#1-10, **Fig. 1D**), and these parts were seamlessly connected to generate a 3-meter diameter circular arena. A start-lift was set at the center of the arena surface. The lift was made of the same material as the arena, and the diameter was 11.6 cm. This lift vertically transported each mouse to the arena surface (79 cm height from the floor) from lower end (22 cm below the arena surface) at each trial start. Thirty-six slotted holes were equally spaced along with the parts #5∼10 which are the circumference parts of the arana. The width, length, radius of each hole was 8, 16, 4 cm, respectively. In cohorts of this study, twenty-four holes were closed by lids of the same material as the arena surface, while every 3 holes (= 12 holes) were opened. As in the BM1 task, each goal hole for each mouse was pseudo-randomly chosen, and an acrylic escape tunnel (width, height, and length was 9, 8.5, 30.5 cm, respectively) was connected under the goal hole at a gentle slope, 20°, so that the mice can easily run into the tunnel from the arena. The floor of the escape box was covered with paper cage bedding. A dummy tunnel was connected to the hole opposite to the goal hole. The design of the dummy tunnel is the same as the escape tunnel except that it was floorless, so mice cannot escape. We expected that strategies such as reaching the escape tunnel by exclusively seeking along the edge of the arena should be inefficient, and that mice taking such strategies should be easily identified. Twelve displays (112.5×65.5 cm; 3840×2160 pixel; DME-4K50D; DMM.com, Japan) were arranged so that these surrounded the arena at equal space. Each display presented 1 unique color, and every 2 displays presented a unique graphic. Four unique 3D objects were placed between the arena and the displays at equal space. Thus, the total 16 objects were presented as unique spatial cues. To mask environmental sounds that might allow sound localization, white noise was played from the speakers of all displays during the task. The loudness level was 55 dB in the arena. Eight room lights attached at the ceiling uniformly illuminated the arena at 500 lx. To mask scene and sound outside the arena, the BM3 apparatus described above were located in a compartment that was separated by a white round curtain.

Behavior during the trials was recorded using a GigE Vision camera (UI-5220SE; IDS Imaging Development Systems GmbH, Obersulm, Germany). The camera was mounted on the ceiling of the compartment (160 cm above the maze center) and was loaded with the ultrawide angle lens (Theia MY125M, Nittoh Inc. Nagano, Japan), such that a spatial resolution achieved 7.43 mm/pixel. Each image frame (404 × 404 pixels, with ×2 binning) was acquired at 20 Hz and displayed on a monitor outside of the compartment. All programs used for data acquisition, processing, saving, and synchronized device controls, were written in LabVIEW 2017 (National Instruments, TX, US).

#### Experimental procedure

The protocol of the BM3 consists of habituation (Day 0), training (Days 1 to 12), and the probe test (Day 13). Immediately before the trial, the experimenter moved a mouse from the homecage to one of 3 releasing points equally spaced around the arena, by a long stainless ladle covered by a lid. Then, the experimenter released the mouse from the ladle into the bottom of the lift. During this move, the mice could not see outside, as the view was obscured by the lid on the ladle. Because equilibrioceptive and proprioceptive cues might reveal certain directions, the releasing points were pseudo-randomized across trials, but common within trials. Other procedures in the habituation, training and probe test were the same with those in the BM1, except that the escape tunnel was used instead of the escape box.

### Contextual fear conditioning

The contextual fear conditioning test for Cohort S1 was performed for successive 2 days on the fear conditioning test system (O’HARA & CO.,LTD. Tokyo, Japan). On Day 1, the mice learned the association between the context and electric footshock (acquisition). The experimenter wearing a white lab coat moved mice homecage from the breeding rack to the standby rack in the experimental room, 30 min before the trial started. Immediately before the trial started, the experimenter moved each mouse from the homecage to the chamber by using the delivery cage filled by woodchip. The acquisition chamber was composed of an acrylic cube (17×15×12 cm), with a wall of black and white stripes. The trial was started after the mice entered the chamber, and lasted for 9 min. At 148 s from the trial start, an electric foot-shock (0.4 mA) was given from the grid floor for 2 s duration, and was repeated 5 times with 90 s interval. After the trial ended, the mice were returned to the homecage via the delivery cage. The soundproof box, chamber, and the grid floor were cleaned with 70% ethanol before each trial started. The brightness in the chamber and experimental room was 70 lx and 80 lx, respectively.

Twenty-four hours after the acquisition, the mice were exposed to a novel context to confirm that the generalization of the fear memory did not occur (retrieval 1). In this context, the soundproof box and the chamber was cleaned with 70% propanol. The retrieval chamber was composed of an acrylic white cube (18×11×10.5 cm) paved by latex sheets. The room light was turned off, while the concealed light was turned on, such that the brightness in the chamber and the experimental room was 40 and 8 lx, respectively. The experimenter moved each mouse directly to the retrieval chamber from the homecage in the breeding rack, then trial was started. Each trial lasted for 6 min, and no foot-shocks were given. After the trial ended, the experimenter returned each mouse directly to the homecage from the chamber. Two hours after the retrieval 1, the mice were re-exposed to the acquisition context (retrieval 2). The environment and procedure were the same as in the acquisition, except that the duration per trial was 6 min, and no foot-shocks were presented.

All behaviors in the chamber were recorded by a camera at 2 Hz. Freezing response as a conditioned response in each trial was detected when the number of pixels whose intensity changed between successive 2 frames were fewer than 30, and this state continued for more than 2 seconds. Freezing rate was calculated as a percentage of total duration of freezing response to total recording time for each mouse in each trial. We assume that contextual fear memory is formed if the freezing rate in the retrieval 2 was significantly higher than that in the retrieval 1.

### Data analysis

In the BM tasks, the mouse position coordinates for every recorded frame were estimated by either a shape adaptive mean shift or an ellipse detection algorithm implemented by the LabVIEW programs. The sequence of coordinates was transformed so that the target holes were located at the same points across mice. Conventional, strategy, and network analyses were then performed to extract the corresponding behavioral features. The definitions of these features are described in a previous study (Suzuki & Imayoshi, 2017).

The conventional analysis calculated the number of errors, time of latency, travel distance to reach the target hole for each mouse in each trial in the training phase, where the error means that the number of visits to dummy holes until the goal. Time spent around each hole was calculated at the probe test. The average of each feature across the three trials per day was calculated for each mouse. Learning curves of the BM1 and BM3 across trials were quantified by intercept and slope estimated from non-linear exponential curve fitting.

The strategy analysis was carried out to find dynamic components of spatial learning. Trajectory in each trial were categorized as any one of spatial, serial, or random strategy by algorithm-based classification. In brief, if a mouse in a trial moved directly toward the target hole and the number of visits to dummy holes was less than three, the strategy was classified as “spatial”. The “serial” strategy was assigned when the mouse sequentially approached neighboring holes until they reached the target with fewer than three quadrant crossings. All remaining behavioral patterns were assigned the “random” strategy. For quantitative definition of each strategy, see Suzuki & Imayoshi, 2017. The strategy analysis was applied to the data only in the training phase. To avoid overestimation such that the random strategy was assigned even to micromovements around the center of the arena, the trajectory data while the mice stayed within 8 cm (BM1) or 27 cm (BM3) distance of the center lift were discarded from the strategy analysis.

The network analysis for spatial navigation behavior in rodents was initially established by Weiss et al., 2012 demonstrated that the object exploration behaviors of rats could be visualized as structured networks of interconnected nodes. If it was applied for the Barnes maze task, simplifying network structures in navigation behaviors across spatial learning could be observed (Suzuki & Imayoshi, 2017). Briefly, a network is constituted from nodes and links between them. If a trajectory during navigation in an environment is expressed as a network, nodes and links can correspond to the places where certain behavior was observed, and transitions between the places, respectively. A node was generated at the coordinate clustering stopping points. When the traveling distance at least 20 successive frames was less than the threshold, one stopping coordinate was generated at the centroid of the points during the frames. The threshold was set at 4 cm, which was approximately half of the average mouse’s body length. Then, the City Clustering Algorithm recursively generated nodes using the following three steps until a convergence condition was satisfied (Rozenfeld et al., 2008; Weiss et al., 2012). First, the coordinates of a stopping point were selected and then integrated those coordinates into the nearest node when the distance between the stopping coordinates and the node was less than 4 cm. If a stop was not integrated into any nodes, the stop was treated as a new node. Second, the coordinates of all nodes were calculated, where each node was represented by the centroid of its constituent stops. Third, if all stopping coordinates were integrated into a node, the processing was stopped, otherwise returned to the first step.

A structure of a network can be quantified by a set of measures. The order was the number of nodes in the network. The degree measured the number of links connected to a node in the network. The density is the ratio of the number of links that are actually present in a network to the number of links that are theoretically possible in the network. The clustering coefficient is the probability that two neighbors of a given node are themselves neighbors. The shortest path of node *i* was quantified as the averaged number of links traversed along the shortest path between node *i* and all other nodes. Betweenness centrality is the extent to which a given node *i* lies on the shortest paths between node *s* and *t*, and is given by the sum of ratio *n_st_^i^* to *g*_st_. If node *i* lies on the shortest path from node *s* to node *t*, *n_st_^i^* is 1, otherwise 0. *g_st_* is the total number of shortest paths from node *s* to *t*. Eventually, betweenness centrality for node *i* are calculated for and averaged over all possible pairs of nodes other than *i* in the network. Closeness centrality measures the inverse of the sum of path lengths from a given node to other nodes. Degree, clustering coefficient, shortest path length, betweenness centrality and closeness centrality were calculated per node, and were averaged over nodes in each network. We assumed neither directed links nor self-links.

### Statistics

In the conventional analysis, a mixed design two-way analysis of variance (ANOVA) for number of errors, latency and travel distance in the training phase, and a mixed design two-way ANOVA for time spent around each hole in the probe test were applied. If significant differences were detected in either or both of the interaction and main effects, Tukey’s honest significant difference (HSD) test was performed as multiple comparisons. In strategy analysis and network analysis, depending on the number of groups, either the Wilcoxon rank sum test or Kruskal–Wallis test with Bonferroni correction was applied for the usage of each strategy and in network measures between groups each day, respectively. If a significant difference was found in Kruskal–Wallis test, Wilcoxon rank sum test with Bonferroni correction was applied as a *post hoc* comparison. For all statistical analyses, the significance level before Bonferroni correction was set at 0.05. For ANOVA and Wilcoxon rank sum test, η*_p_*^2^ and *r* was calculated as effect size, respectively. For strategy and network analysis, statistical results were summarized in **Statistical table 1-9**.

### Code Accessibility

All codes for data acquisition and analysis were developed in LabVIEW 2013, 2017 (National Instruments, TX, US) and MATLAB R2018a (MathWorks Inc., MA, USA) on a custom-built workstation [Windows 10, Intel(R) Core(TM) i7-7700K CPU @ 4.20 GHz, 32.0 GB RAM]. The code described in the paper is freely available online at [URL redacted for double-blind review] or **Extended data 1**.

## Results

In total, 111 mice were used in this study, and each was assigned to one of 7 cohorts in advance (**Table 1**). Each cohort had a unique history of task instances. Cohorts 1 and 2 were engaged in only 1-meter diameter Barnes maze (BM1) and 3-meter diameter Barnes maze (BM3) task, respectively. Cohort 3 was engaged only in the BM3 task as in Cohort 2, but was treated with scopolamine hydrobromide, a non-selective muscarinic receptor antagonist, at the probe test. Cohorts 4 to 6 were engaged in 2 instances. Cohort 4 was engaged in the BM3 task at the first instance, then in the BM1 task at the second instance. Cohort 5 was engaged in the modified BM1 (BM1’) task at the first instance, then in the BM1 task at the second instance, where the BM1’ was identical to the BM1 except the spatial cues and goal locations. The task instance of Cohort 6 was opposite to that of Cohort 4; the BM1 at the first instance, the BM3 at the second instance. As a control, Cohort S1 engaged in a contextual fear conditioning test at the first instance, then in the BM1 task at the second instance.

### Spatial learning within the BM3

First, we explored whether spatial learning in Barnes maze (BM) is calibrated by scale spaces, comparing the behavioral performances within the BM3 with those in the BM1. The pooled data of Cohort 1 (n = 20) and those of the first instance in Cohort 6 (n = 14) were assigned to the BM1 group, while the pooled data of Cohort 2 (n = 20), Cohort 3 (n = 16), and those of the first instance in Cohort 4 (n = 20) were assigned to the BM3 group. Because Cohort 3 was administered scopolamine hydrobromide at the probe test (see below), the data only in the training phase were used.

The diameter of the field of BM1 was 1 m (**Fig. 1A, C, E**), while that of the BM3 was 3-meter (**Fig. 1B, D, F**). In common to both BM1 and BM3, the field of the maze was the brightly illuminated open circular field, and the mice were automatically introduced to the field via a lift located at the center of the field at the start of each trial. A hole was located along the edge of the field at 30 angular intervals. Any one of the holes was a goal, and a dark escape box was attached beneath the goal hole so that mice could escape from the field. The selection of the goal hole was pseudo-randomized and counterbalanced across mice, but fixed within each mouse across the experimental phase. A unique 3D object was set at the north, south, east and west so as to surround the field of the BM1. For spatial cues in the BM3, a display presenting a unique combination of color and icon was located beyond each hole on the field, and 4 unique 3D objects were inserted between the displays asymmetrically. These spatial cues could be exploited to compute current mouse position and the destination on the field. No spatial cues were shared between the BM1 and the BM3.

The experimental phase consisted of the habituation, training, and probe test in both BM1 and BM3. In the habituation phase on Day 0, the mice were allowed to freely move the field for 10 min just once. In the training phase, the mice explored the field until they escaped to the escape box located under the goal hole. This training trial was repeated 3 times per day for each mouse. If mice acquired precise representation of the maze through the training phase, their navigation would be optimized for lower number of hole visits, shorter latency and shorter travel distance from start to goal. The training phase in the BM1 and the BM3 lasted for 6 days and 12 days, respectively. The probe test phase was conducted on the next day of the last training day, and precision of spatial representation of the maze was evaluated. The mice explored the field freely for 3 min without the escape box, as in the habituation phase. Therefore if the mice remember the exact location of the goal, they would focally search around there. For more details about the architecture and task design of BM1 and BM3, see Material and Methods.

### Lower learning rate in the BM3 than in the BM1

To evaluate learning rate across training in the BM1 and the BM3 group, 3 conventional features (**Table 1**), the number of errors, latency, and travel distance, were calculated from moving trajectories on the field per training trial, then a learning curve was determined for each mouse (**Fig. 2A-C**). To calculate the learning curves of latency and travel distance between the BM1 and BM3 in unitless form, these were normalized within individuals such that the sum of values across trials in each learning curve becomes 1 (**Fig. 2A-C**). Unnormalized latency and travel distance was displayed in Extended Data Fig.2-1. Then, each learning curve from Day 1 to 6 was fitted to an exponential function, *y = αe^-βt^* estimating optimal intercept α and decay parameter β, in nonlinear least squares method, where *t* is a trial, *y* is a value of a feature, and *e* is Euler’s number. If β was lower, the learning curve takes longer time to converge. We found that β of the learning curve was significantly lower in the BM3 than in the BM1, in all features (**Fig. 2A, B, C**). Wilcoxon rank-sum test reported that decay parameter β in the learning curve of the number of errors, latency and travel distance was significantly lower in the BM3 than in the BM1, *z* = 2.87, *p* = 0.00, *r* = 0.30, *z* = 2.11, *p* = 0.03, *r* = 0.22, *z* = 3.02, *p* = 0.00, *r* = 0.18, respectively. The daily basis learning curve was calculated for each mouse by averaging values across 3 trials within each day from Day 1 to Day 6, then compared between the BM1 and the BM3. As expected, the BM3 required a significantly higher number of errors, longer latency and longer travel distance until mice reached the goal, regardless of training days (**Fig. 2D-F**). In a mixed-design 2-way [Scale (BM1, BM3)×Day (1∼6)] ANOVA for the number of errors, latency and travel distance, the main effect of the scale was significant, *F*(1,88) = 21.11, *p* = 0.00, *η_p_*^2^ = 0.19, *F*(1,88) = 59.82, *p* = 0.00, *η*_p_*^2^* = 0.40, *F*(1,88) = 19.08, *p* = 0.00, *η_p_*^2^ = 0.18, respectively. Thus, the learning rate was lower in the BM3 than in the BM1, therefore spatial learning takes a longer time to be established in the BM3 than in the BM1.

**Figure 2.**
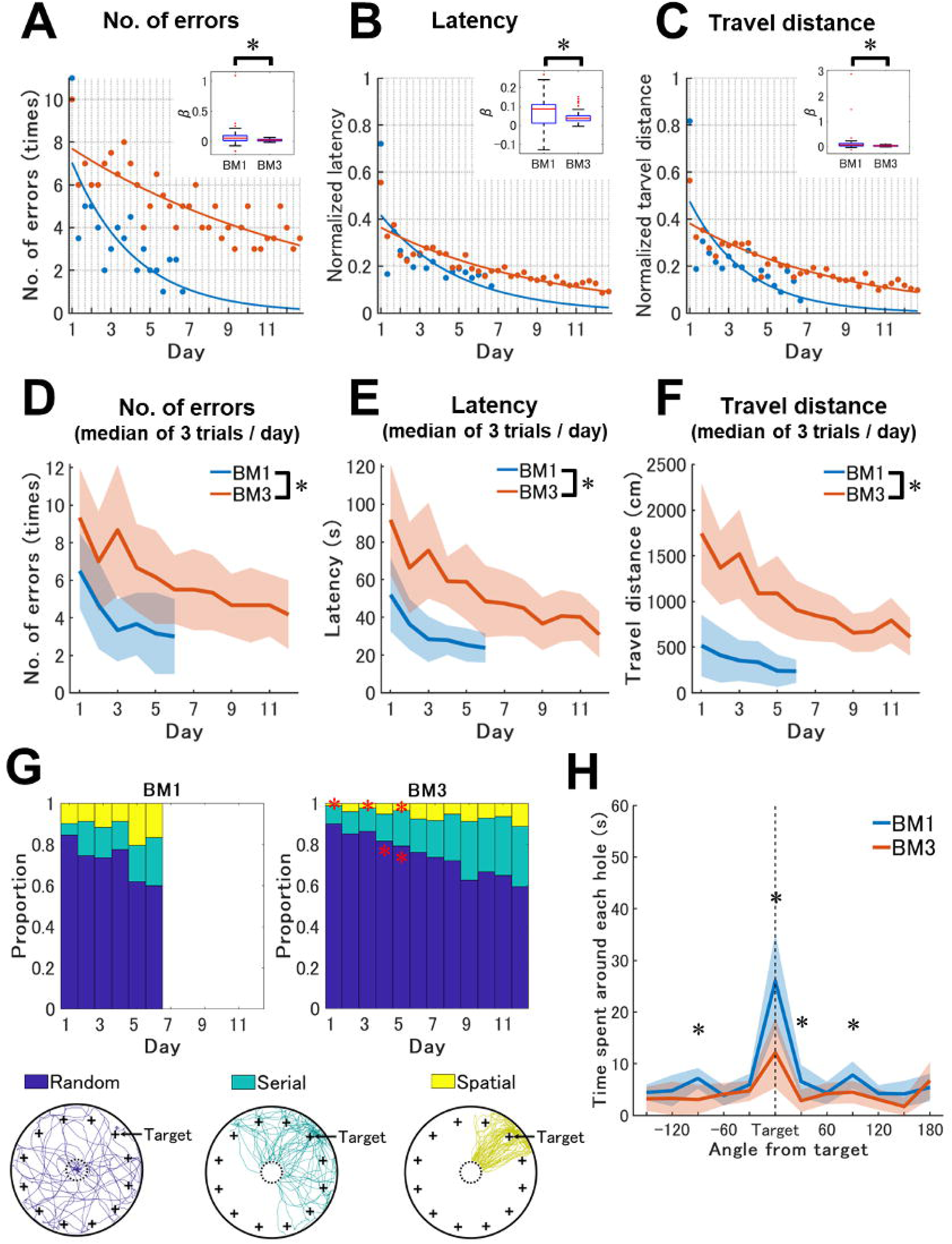
Lower learning rate and inaccurate spatial representation in the BM3 compared to the BM1. ***A–C,*** Trial-based learning rate across training periods in conventional features in the BM1 and the BM3. Three trials per day were conducted for 6 successive days in the BM1 and 11 successive days in the BM3. The measured values of number of errors (**A**), and the normalized values of latency (B) and travel distance (**C**) to reach the goals were displayed. The horizontal axis and the vertical gray grid lines of each panel indicates training days and trials, respectively. Note that both latency and travel distance were normalized so that the values range between 0 and 1. Blue- and red-colored dots or lines represent the results of BM1 (n = 34) and BM3 (n = 56), respectively. Each dot represents the median value of each group in each trial. Solid lines represent learning curves estimated by non-linear exponential curve fitting for the median of the groups. Box plots in the small insets in (***A–C***) represent distributions and comparisons of decay parameter β of the learning curves between the mouse group of BM1 and BM3. Asterisks indicate statistically significant differences. ***D–F,*** Daily basis learning curves across training periods in conventional features in the BM1 and the BM3. These scores were averaged over 3 trials per day. Blue and red solid lines represent changes in the median of measured values of number of errors (***A***) latency (***B***) and travel distance (***C***) of the BM1 and the BM3, respectively. Shaded areas indicate median absolute deviation within a mouse group each day. Asterisks indicate significant differences between the BM1 and the BM3. ***G,*** Strategy usage of the subjected mice across training days in the BM1 and the BM3. The left and right panel indicate the results of BM1 and the BM3. The vertical and horizontal axes are the proportion of the strategies and training days, respectively. In each stacked bar graph, blue, green and yellow color represent random, serial and spatial strategy, respectively. Red asterisks indicate significantly different strategy usage between the BM1 and the BM3 in a given day. Samples of each strategy observed in the BM3 are shown under the stacked bar graphs. From left, random, serial spatial strategies are shown. Each colored line represents a trajectory classified as a strategy in a trial. These samples were chosen so that the sum of travel distances of samples within each strategy were comparable between the strategies. All trajectories were transformed so that the goal is located at the right top hole, noted as “Target”. The larger circles with black solid lines represent the edges of the BM3, while the smaller circles with black dashed lines represent the start areas. “+” markers represent hole locations. ***H,*** Time spent around each hole in the probe test. The horizontal axis indicates the locations of the holes expressed as angle differences from the target. The vertical and horizontal axes are search time for the individual hole and hole location indicated by angle from the target with 30° step, respectively. “Target” shown by a black dashed line is the goal hole for each mouse. Blue and red represent the results of BM1 (n = 34) and BM3 (n = 40), respectively. The solid lines are connected between the median values in each angle of each mouse group. The shaded areas are a range of median ± median absolute deviation of the data in each angle. Asterisks indicate significant differences between the BM1 and the BM3.

Strategy analysis was performed to qualitatively evaluate how mice optimize their navigation strategies during spatial learning on the BM task (Bach et al., 1995; Barnes, 1979; Eales et al., 2014; Suzuki & Imayoshi, 2017). If the mice have optimized their navigation strategy through the training, they would move along the straight line from the start to the goal (spatial strategy). If mice have incomplete knowledge about the maze and the task (e.g. the goal hole is any one of holes located around the edges of the field), they would sequentially visit each hole toward the goal (serial strategy). If mice were almost naive for the maze and the task, they would show an indeterminate pattern in the trajectory (random strategy). Complete definition of each strategy was described in **Extended Data Table 1**. The usage of each strategy was compared between the BM1 and the BM3 within each day from Day 1 to Day 6 (**Fig. 2G**). The usage of spatial strategy on days 1, 3 and 5 was significantly lower in the BM3 than in the BM1 (**Statistical table 1, reference #13, 15, 17**, respectively), while that of random strategy on days 5 and 6 was significantly higher in the BM3 than in the BM1 (**Statistical table 1, reference #5, #6**, respectively), suggsting that the BM3 would prompt the mice to take more trial-and-error. Thus, the BM3 would require larger computational resources to optimize navigation strategies than in the BM1.

In the probe test, mixed-design 2-way [Scale (BM1, BM3)×Hole (1∼12)] ANOVA for the time spent around each hole reported a significant interaction between scale and hole, *F*(11,792) = 11.87, *p* = 0.00, *η_p_*^2^ = 0.14. Multiple comparisons detected that exploration time around the target hole was significantly longer than those around other holes in both BM1 and BM3, while those around 2 holes (target and target +30°) were significantly shorter in the BM3 than in the BM1 (**Fig. 2H**). Also, time around the target ±90° holes were significantly longer in the BM1 than in the BM3. Because a spatial cue was located beyond each target hole, the target ±90° and +180° in the BM1, the BM1 mice might preferentially search around these cues, when they observed that the goal no longer existed. Thus, retrieval of spatial representation was more inaccurate in the BM3 than in the BM1.

### Characteristic network structures of moving trajectory in the BM3

A moving trajectory of a rodent in a field can be expressed as a network with nodes and links, where a node was defined as *xy* coordinates that rodents stayed for certain time, and a link was defined as a transition between 2 nodes (Suzuki & Imayoshi, 2017; Weiss et al., 2012). The structure of a network was quantified by a set of measures (for complete description of network analysis, see **Extended Data Table 1** and Material and Methods), and shed light on novel aspects of spatial navigation and learning in rodents (Suzuki & Imayoshi, 2017; Weiss et al., 2012) Indeed, a previous study has demonstrated that network structures were simplified across spatial learning in the BM1 (Suzuki & Imayoshi, 2017). Here, we performed this network analysis to explore whether network structures diverge between the BM3 and the BM1 across training days 1 to 6.

In the training phase, the number of stops, order, degree, and shortest path length were significantly and constantly higher in the BM3 than in the BM1 in all training days (**Figure 3A, B, C & F; Statistical Table 2, references #1-18 and 33-38**). These results indicate that the networks generated in the BM3 had more nodes and links, while requiring more traverses between arbitrary 2 nodes than those in the BM1.

**Figure 3.**
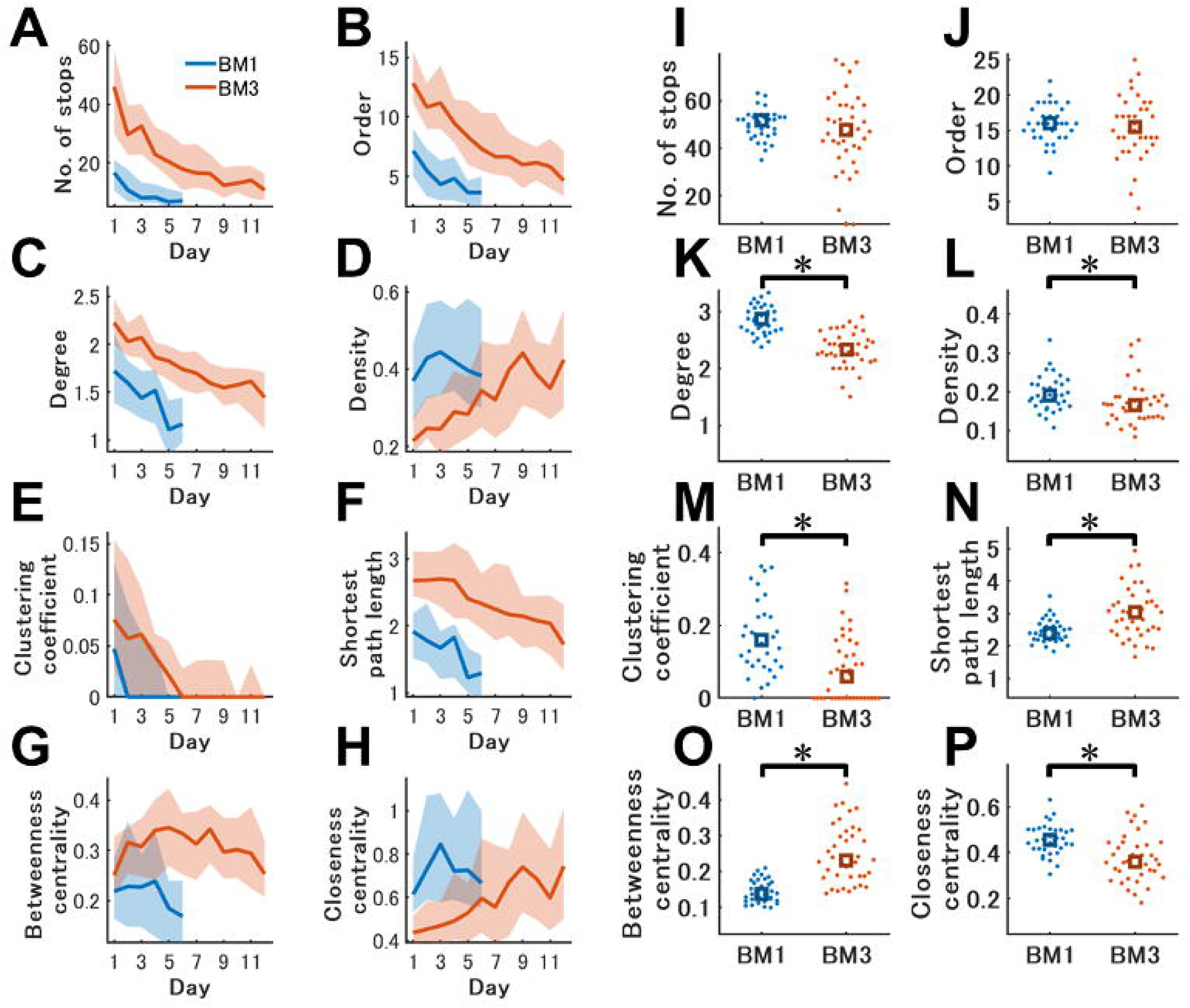
Difference of network structures between the BM1 and the BM3. ***A–H,*** Changes of network features across training in the BM1 and the BM3. Temporal changes in the graphically displayed global networks in the BM1 and BM3 during spatial learning were illustrated in **Extended data Fig. 3-1**. The vertical and horizontal axes in each panel are calculated values of each network feature and training days. These scores were averaged over 3 trials per day. Blue and red represent the results of BM1 (n = 34) and BM3 (n = 56), respectively. Solid lines represent changes in the median value of each group in each day. Shaded areas represent error ranges between 25 and 75 percentile of the data within a group each day. Statistical results on each day are shown in Statistical table 2. ***I–P,*** Network features in the probe test in the BM1 and the BM3. The vertical axis in each panel is each network measure. The blue and red represent the results of the BM1 and BM3 group, respectively. Each dot represents a mouse in either group. Squares represent median value in each group. Asterisks indicate significant differences between the BM1 and the BM3.

Density and closeness centrality was significantly higher in the BM3 than in the BM1 until Day 5, indicating that the observed links in the network in the BM3 accounted for a smaller part of theoretically possible all links, relative to those in the BM1 (**Fig. 3D, H**). The density and closeness centrality across training days would initially diverge but eventually converge between the BM1 and the BM3 (**Fig. 3D, H**). Indeed, both were significantly lower in the BM3 than in the BM1 from Day 1 to Day 5 (**Statistical table 2, references #19-23 and 47-51**). We confirmed if the measures changed across days within each group. Density and closeness centrality was comparable between Day 1 and 6 in the BM1 (**Statistical table 2, reference #25, 53**,), while that was significantly higher on Day 6 than on Day 1 in the BM3 (**Statistical table 2, reference #26, 54**). Conversely, betweenness centrality was comparable between the BM1 and the BM3 on day 1, while those were higher in the BM3 than in the BM1 on subsequent days (**Fig. 3G**). Betweenness centrality was significantly higher in the BM3 than in the BM1 from Day 2 to Day 6 (**Statistical table 2, reference #39-44**). In the BM3, the betweenness centrality represented an arch-like curve across training days, and was comparable between Day 1 and 12 (**Statistical table 2, reference #46**). In contrast, that was significantly lower on Day 6 than on Day 1 in the BM1 (**Statistical table 2, reference #45**). Betweenness centrality is a probability such that a node locates on a shortest path between any other 2 nodes in a network. One interpretation of the results is that certain places relaying any other places were constantly required across training in the BM3, while those could become unnecessary along with training in the BM1. Thus, changes of density, closeness centrality and betweenness centrality across training might depend on both scale space and learning progression.

In the probe test, we found large differences in network structures between the BM1 and the BM3. Degree, density, clustering coefficient, and closeness centrality was significantly lower in the BM3 than in the BM1 (**Fig. 3K, L, M, P**; **Statistical table 3, reference #3, 4, 5, 8**, respectively). These results indicated that the nodes were sparsely linked to others in the BM3 than in the BM1. This leads to a significantly longer shortest path in the BM3 than in the BM1 (**Fig. 3N**; **Statistical table 3, reference #6**). As in the later phase of training, betweenness centrality was significantly higher in the BM3 than in the BM1 (**Fig. 3O**; **Statistical table 3, reference #7**), suggesting the existence of more places on a shortest path between other 2 places in the BM3 than in the BM1. In contrast, the number of stops and order were comparable between the BM1 and the BM3 (**Fig. 3I, J**; **Statistical table 3, reference #1, 2,** respectively). One possible explanation is that when mice explored for a given time, the number of places where they stayed for more than 1 s is constant regardless of scale spaces, while the transition patterns between them were modulated by scale spaces.

### Limited effects of non-selective muscarinic receptor antagonist for navigation in the BM3

A previous study (Suzuki & Imayoshi, 2017) found that the mice that were treated scopolamine hydrobromide, a non-selective muscarinic acetylcholine receptor antagonist, focally searched around the goal, compared to vehicle-treated mice in the probe test in the BM1. In this study, we tested whether such effects from scopolamine hydrobromide are maintained in a large scale space. The performances in Cohort 3 (n = 16) were compared with those in the pool of Cohort 2 (n = 20) and the first instance of Cohort 6 (n = 14) (**Table 1**). The former and latter group were termed the SCOP and NO-SCOP, respectively. While both groups were engaged to the same BM1 training, the SCOP group were injected with scopolamine hydrobromide (Santa Cruz Biotechnology, TX, USA) intraperitoneally 20 min before starting the probe test. Scopolamine was prepared as a 0.3 mg/ml stock solution in 0.9% saline, so as to be 3 mg of scopolamine hydrobromide/kg of body weight. Other experimental procedures were the same between the groups.

Partially consistent with the BM1 (Suzuki & Imayoshi, 2017), the SCOP mice searched holes near the goal for a longer time, while more sparsely searching away from the goal, compared to the NO-SCOP mice (**Extended DataFig. 2-2A**). A mixed-design 2-way [Scopolamine (SCOP, NO-SCOP)×Hole (1∼12)] ANOVA for the time spent around each hole detected significant interaction between the scopolamine and hole, *F*(11,594) = 2.40, *p* = 0.01, *η_p_^2^* = 0.04. Multiple comparison testing detected that the time spent around target +30° was significantly longer in the SCOP than in the NO-SCOP whilst time spent around the opposite hole from the target was significantly shorter in the SCOP than in the NO-SCOP. Because a dummy escape tunnel was attached under the hole opposite to the goal hole, the NO-SCOP but not SCOP mice might search around it, once they observed that the true escape tunnel no longer exists at the goal hole. In contrast to the BM1 (Suzuki & Imayoshi, 2017), the SCOP mice did not show characteristic network structures compared to NO-SCOP mice in the BM3. Wilcoxon rank sum test reported that the number of stops was significantly higher in the SCOP than in the NO-SCOP, *z* = -2.81, *p* = 0.01, *r* = -0.38 (**Extended Data Fig. 2-2B**). Thus, the effects of administration of scopolamine hydrobromide on the navigation network in the BM3 partially replicated those in the BM1, suggesting that the scopolamine effect on navigation is dampened in a larger scale space than a smaller scale space.

### Spatial learning between the BM1 and the BM3

Next, we explored whether scale spaces calibrate not only the present but also subsequent spatial learning. When animals learn a task, they learn not only solutions to the immediate task, but also how to find the solutions. This meta-learning process facilitates subsequent learning in variants of the initially learned task. Thus, if this meta-learning process can be activated even for spatial learning between different scale BMs, prior learning in either BM scale (e.g. BM3) would facilitate subsequent spatial learning in the other BM scale (e.g. BM1).

### Prior learning in the BM3 facilitated subsequent spatial learning in the BM1

First, we evaluated facilitation of prior spatial learning in the BM3 on the subsequent learning in the BM1. Cohort 4 (n = 20) engaged in the BM3 task as the first instance, then the BM1 task as the second instance (**Table 1**). Cohort 5 (n = 17) was a control that engaged in the BM1’ task as the first instance then in the BM1 task as the second instance. The BM1’ was a variant of the BM1 (**Extended Data Fig. 4-1A**), and lasted for 12 days such that the number of training days matched with the BM3 task. Thus, Cohort 4 and 5 were treated as the BM3 learners and the BM1’ learners, respectively. The pool of Cohort 1 and the first instance of Cohort 6 was treated as the BM1 beginners (n = 34). The performances of BM1 were compared between the BM3 learners, BM1’ learners and the beginners. We confirmed that spatial learning could be established in the BM1’ task at the first instance of Cohort 5 (**Extended Data Fig. 4-1B**).

We found that both BM learners exhibited efficient navigation in the training of the BM1, compared to the beginners (**Fig. 4A**). In conventional analysis, mixed-design 2-way [Instance (BM3 learner, BM1’ learner, beginner)×Day (1∼6)] ANOVA for number of errors, latency and travel distance detected significant main effect of instance, *F*(2,68) = 9.03, *p* = 0.00, *η_p_^2^* = 0.21, *F*(2,68) = 3.99, *p* = 0.02, *η_p_*^2^ = 0.11, *F*(2,68) = 8.33, *p* = 0.00, *η_p_*^2^ = 0.20, respectively. Tukey’s HSD test reported that the number of errors and travel distance in the BM learners were significantly lower than the beginners. Latency in the BM1’ learners was significantly lower than the beginners.

**Figure 4.**
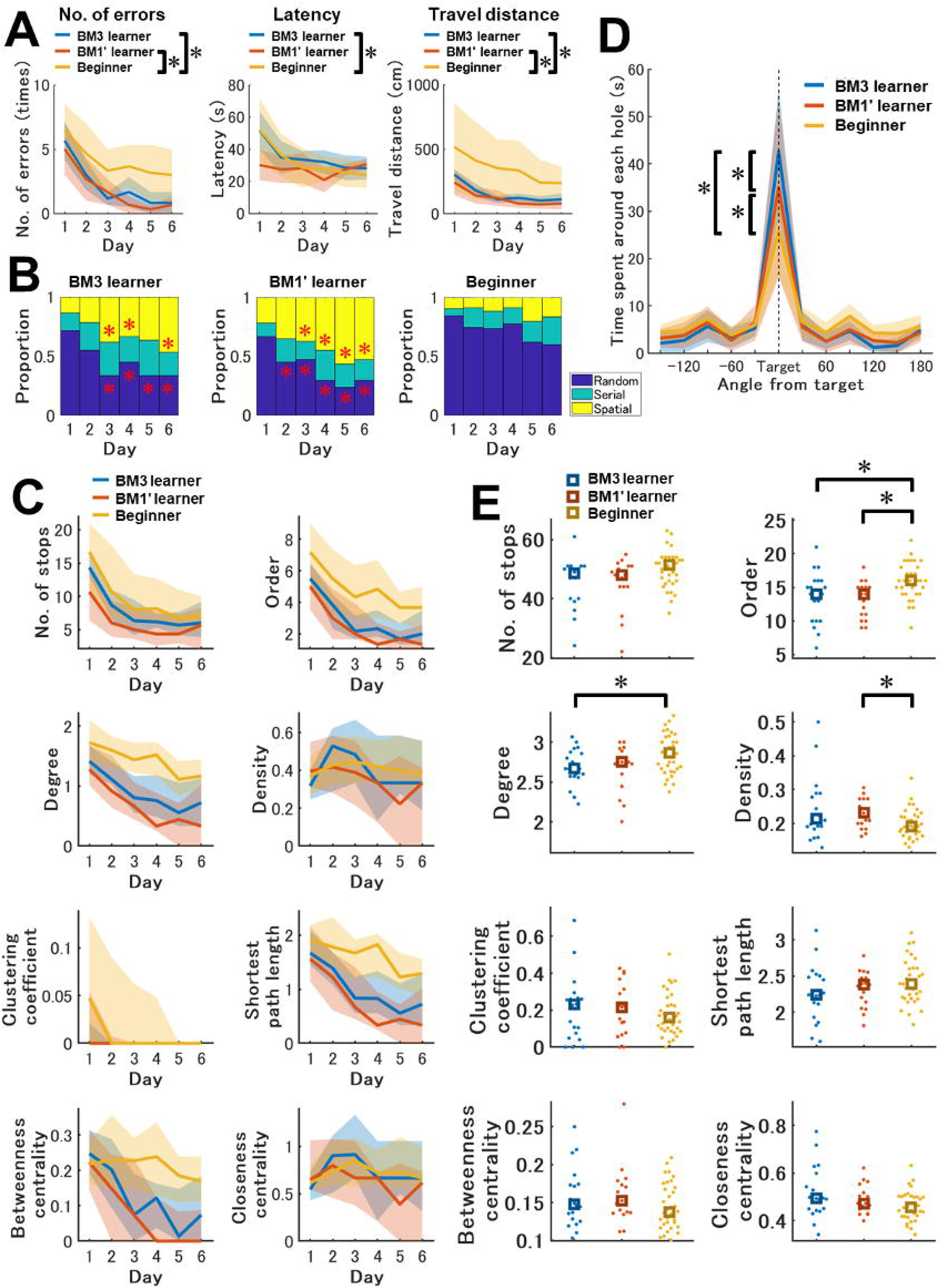
Effects of prior learning in the BM3 and BM1’ on spatial navigation and learning in the subsequent BM1 task. ***A,*** Daily basis learning curves across training periods in conventional features in the BM1. These scores were averaged over 3 trials per day. From left, the measured values of number of errors, latency and travel distance are displayed. The mouse groups of BM3 learner (n = 16) and BM1’ learner (n = 17), which experienced the BM3 and BM1’ before the BM1, respectively, were compared with the Beginner (n = 34) group. The number of errors and travel distances of the BM3 learner and the BM1’ learner were significantly lower than Beginner. Latency in the BM1’ learner was significantly shorter than Beginners. ***B,*** Daily strategy usage during BM1 training in the BM3, BM1’ learner and Beginner mouse groups. Although usages of all navigation strategies were comparable between the BM3 and the BM1’ learner in all training days, significant differences were detected in the comparison of BM3 learner and Beginner, and BM1’ learner and Beginner. Red asterisks indicate significantly different strategy usages compared to the corresponding strategy usages in the beginner on a given day. ***C,*** Temporal changes of network features in the BM1 training of BM3, BM1’ learner and Beginner. The vertical and horizontal axes in each panel are calculated values of each network feature and training days, respectively. These scores were averaged over 3 trials per day. Solid lines represent changes in the median value of each group in each day. Shaded areas represent error ranges between 25 and 75 percentile of the data within a group each day. Statistical results on each day are shown in Statistical table 5. ***D,*** Time spent around each hole in the BM1 probe test of the BM3, BM1’ learner and Beginner mouse group. The vertical and horizontal axis is time and hole location indicated by angle from the target with 30° step, respectively. “Target” expressed as a black dashed line is the goal hole for individual mice. Blue, red and yellow represent the results of Beginner, BM3 learner and the BM1’ learner, respectively. The solid lines are median values in each angle in each group. The areas are a range of median ± median absolute deviation of the data in each angle in each group. Asterisks indicate significant differences between the groups. ***E,*** Network features in the BM1 probe test compared between Beginner, BM3 learner and the BM1’ learner. The vertical axis in each panel is each network measure. The blue, red, and yellow represent the results of Beginner, BM3 learner and the BM1’ learner, respectively. Each dot represents a mouse in each group. Squares represent median value in each group. Asterisks indicate significant differences between the mouse groups.

Strategy analysis showed that the BM learners more frequently exhibited the spatial strategy than the beginner group, after Day 1 (**Fig. 4B**). The usage of spatial strategy was significantly different between instances after Day 2 (Day 3, 4, 5 and 6, **Statistical table 4, reference #30, 34, 38, 42**, respectively). *Post-hoc* comparison reported that usage of spatial strategy in the BM3 learner were significantly higher than the beginners on Day 3, 4, and 6 (**Statistical table 4, reference #33, 37, 45**, respectively), while those in the BM1’ learner were significantly higher than the beginner on after Day 2 (**Statistical table 4, reference #32, 36, 40, 44**, respectively). In contrast, the usage of random strategy was significantly different between instances after Day 1 (**Statistical table 4, reference #2, 6, 10, 14, 18**, respectively). The usage of random strategy in the BM3 learner were significantly lower than the beginners after Day 2 (**Statistical table 4, reference #9, 13, 17, 21**, respectively), while those in the BM1’ learner were significantly higher than the beginner after Day 1 (**Statistical table 4, reference #4, 8, 12, 16, 20**, respectively). All strategy usages were comparable between the BM3 and the BM1’ learner.

The network structures across training days in the BM learners resembled each other while those were distinguishable from the beginner group (**Fig. 4C**). The number of stops was significantly different between instances on Day 4 (**Statistical table 5, reference #4**). *Post-hoc* comparison reported that the number of stops in the BM1’ learner was significantly lower than the beginners (**Statistical table 5, reference #6**). Order was significantly different between instances after Day 1 (Day 2∼6, **Statistical table 5, reference #11, 15, 19, 23, 27,** respectively). Order in the BM3 learner was significantly lower than that in the beginner after Day 1 (**Statistical table 5, reference #14, 18, 22, 26, 30**, respectively), while that in the BM1’ learners was significantly lower than that in the beginners after Day 1 (**Statistical table 5, reference #13, 17, 21, 25, 29**, respectively). Degree was significantly different between instances after Day 1 (**Statistical table 5, reference #32, 36, 40, 44, 48**, respectively). Degree in the BM3 learner was significantly lower than that in the beginner after Day 1 (**Statistical table 5, reference #35, 39, 43, 47, 51**, respectively), while that in the BM1’ learners was significantly lower than that in the beginners after Day 1 (**Statistical table 5, reference #34, 38, 42, 46, 50**, respectively). Shortest path length was significantly different between instances after Day 1 (**Statistical table 5, reference #65, 69, 73, 77, 81**, respectively). Shortest path length in the BM3 learners was significantly lower than that in the beginners after Day 1 (**Statistical table 5, reference #68, 72, 76, 80, 84**, respectively), while that in the BM1’ learners was significantly lower than that in the beginners after Day 1 (**Statistical table 5, reference #67, 71, 75, 79, 83**, respectively). These results indicate that the BM learners could move between 2 nodes on the network with fewer traverses of other nodes compared to the beginners. Betweenness centrality in the BM learners was significantly lower than the beginners after Day 2, suggesting that the beginners require certain places relaying between other 2 places on the field compared to the BM learners (**Fig. 4C**). Indeed, betweenness centrality was significantly different between instances after Day 2 (**Statistical table 5, reference #87, 91, 95, 99**, respectively). Betweenness centrality in the BM3 learners was significantly lower than that in the beginners from Day 3 to Day 5 (**Statistical table 5, reference #90, 94, 98**, respectively), while that in the BM1’ learners was significantly lower than that in the beginners after Day 2 (**Statistical table 5, reference #89, 93, 97, 101,** respectively). Thus, conventional, strategy, and network analysis suggest that the BM learners could learn the BM1 task efficiently compared to the beginners.

Not predicted by our hypothesis, the BM3 learners exhibited the most accurate spatial memory in the probe test (**Fig. 4D**). Indeed, mixed-design 2-way [Instance (beginner, BM3 learner, BM1’ learner)×Hole (1∼12)] ANOVA for the time spent around each hole, significant interaction between instance and hole was detected, *F*(22,748) = 4.16, *p* = 0.00, *η_p_^2^* = 0.11. Multiple comparisons detected that the time spent around the target hole in the BM3 learners was significantly longer than all other groups, and the BM1’ learner was significantly longer than only the beginners. Thus, prior learning in the BM3 rather than in the BM1’ improves spatial representation in subsequent learning in the BM1, suggesting that prior learning in a larger scale space would facilitate subsequent spatial learning in a smaller scale space.

The network analysis revealed significant differences of exploratory networks between instances (**Fig. 4E**). Order, degree, and density were significantly different between instances (**Statistical table 6, reference #2, 6, 10**, respectively). Order in the BM1’ learner and the BM3 learner was significantly lower than the beginner (**Statistical table 6, reference #4, 5**, respectively). Degree in the BM3 learner was significantly lower than the beginner (**Statistical table 6, reference #9**). Density in the BM1’ learner was significantly higher than the beginner (**Statistical table 6, reference #12**). Thus, the BM3 learners would traverse between a limited number of places. These results support the idea that the prior spatial learning in the BM3 most facilitated subsequent spatial learning in the BM1.

Although we limitedly explored meta-learning effects within the BM paradigm, we also checked if prior learning in another behavioral paradigm such that neither environmental nor task structures were shared with the BM paradigm facilitates subsequent spatial learning in the BM1 task. Cohort S1 (n = 8) underwent a conventional contextual fear conditioning (CFC) test at the first instance (see Material & Methods), then BM1 at the second instance. The CFC learners formed contextual fear memory correctly, as Wilcoxon signed-rank test detected significantly higher freezing response in the fear acquisition context (M = 38.14, SD = 15.27), compared to that in the different context (M = 14.30, SD = 10.21), *p* = 0.02, *r* = - 0.84, *z* = -2.38. The performance in the BM1 was comparable between the CFC learners and the BM1 beginners, as there were no significant differences between the two in all conventional features (**Extended Data Fig. 4-2A**).

### Limited facilitation from the BM1 to the BM3 learning

To test whether prior learning in the BM1 facilitates subsequent learning in the BM3, the performances in the BM3 at the second instance of Cohort 6 (BM1 learner, n = 13) were compared with those in the BM3 beginner. The beginners in the training were pool of Cohort 2 (n = 20), and the first instance of Cohort 3 (n = 16) and 4 (n = 20), while those in the probe test were pool of Cohort 2 and the first instance of Cohort 4, because Cohort 3 was treated with scopolamine hydrobromide at the probe test.

In conventional analysis, the number of errors were comparable between the BM1 learners and the beginners, while latency and travel distance was significantly shorter in the BM1 than in the beginner, regardless of training days (**Fig. 5A**). Mixed-design 2-way [Instance (BM1 learners, beginners)×Day (1∼6)] ANOVA for latency and travel distance detected significant main effect of instance, *F*(1,67) = 28.72, *p* = 0.00, *η_p_^2^* = 0.30, *F*(1,67) = 24.85, *p* = 0.00, *η_p_*^2^= 0.27, respectively. These results suggest that the BM1 learners explored places other than the holes less across training days, compared to the beginner.

**Figure 5.**
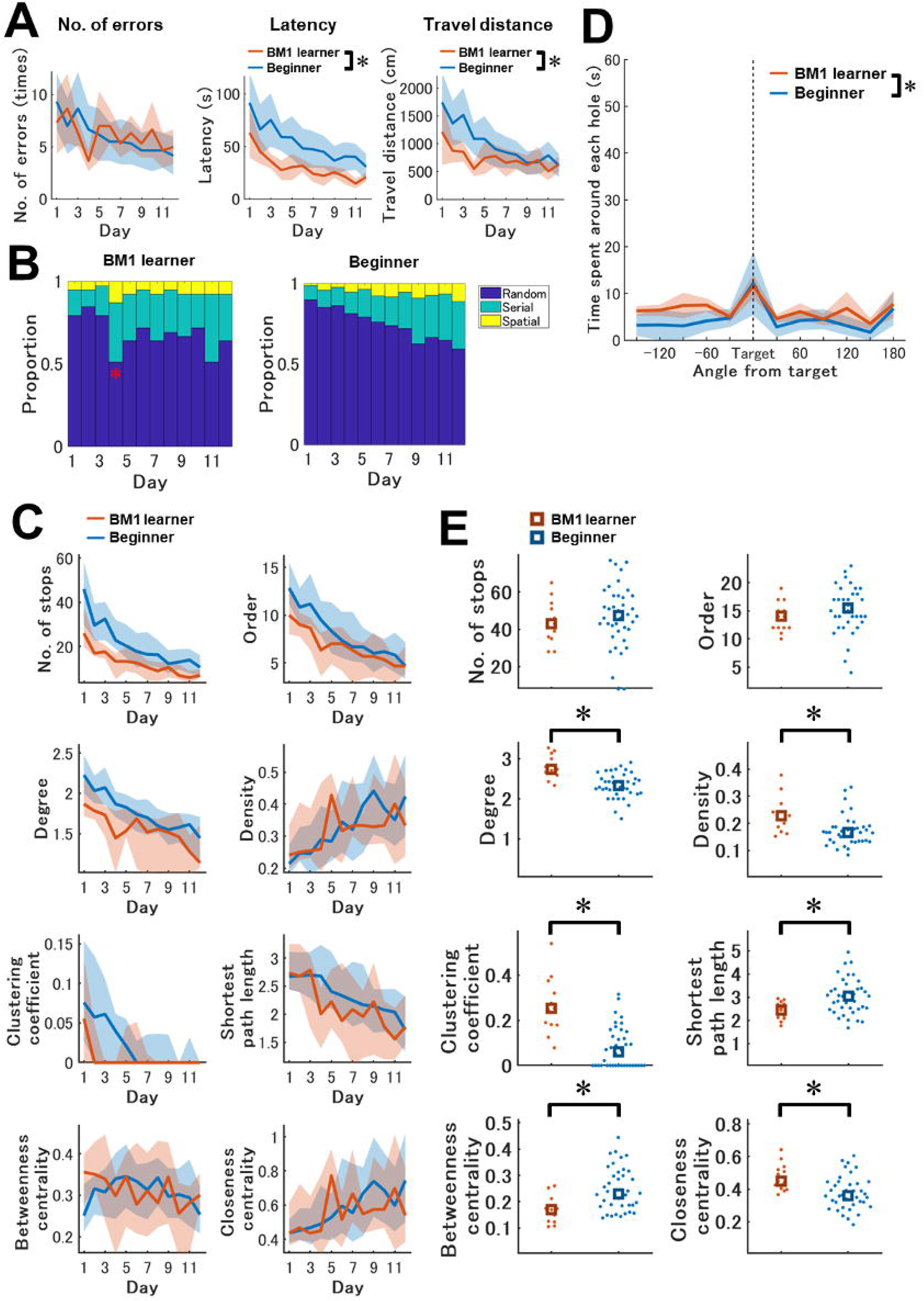
Limited effects of spatial navigation and learning in the subsequent BM3 task by prior learnings in the BM1. ***A,*** Daily basis learning curves across training periods in conventional features in the BM3. These scores were averaged over 3 trials per day. From left, the measured values of number of errors, latency and travel distance are displayed. The mouse group of the BM1 learner (n = 13), which experienced the BM1 before the BM3, was compared with the Beginner (n = 56) group. Although the significant changes were detected in latency and travel distances, no significant difference was detected in the number of errors. ***B,*** Strategy usage across training days in the BM3 and comparison between BM1 learner and Beginner. Limited changes were detected only at the Day 4 results. ***C,*** Temporal changes of network features in the BM3 training of BM1 leaner and Beginner. Statistical results on each day are shown in Statistical table 8. ***D,*** Time spent around each hole in the BM3 probe test. Only significant main effect of instance was observed; exploration time in the BM1 learner was significantly longer than that in the Beginner, regardless of hole location. ***E,*** Network features in the BM3 probe test compared between BM1 learner and Beginner. Asterisks indicate significant differences between the two groups.

In the strategy analysis, strategy usage was almost the same between the BM1 learners and the beginners (**Fig. 5B**). The usage of random strategy was significantly higher in the BM1 learners than in the beginners only on Day 4 (**Statistical table 7, reference #4**).

In network analysis, the number of stops was significantly lower in the BM1 learners than in the beginners on days 1, 3, 4, 7, 8, 11 (**Fig. 5C**; **Statistical table 8, reference #1, 3, 4, 7, 8, 11**, respectively). Though the number of stops were constantly different, order was significantly lower in the BM1 learners than in the beginners only on Day 3 and 4 (**Statistical table 8, reference #15, 16**, respectively). Degree was significantly lower in the BM1 learners than in the beginners on Day 3 and 4 (**Statistical table 8, reference #27, 28**, respectively). Clustering coefficient was significantly lower in the BM1 learners than in the beginners only on Day 3 (**Statistical table 8, reference #51**). Shortest path length was significantly lower in the BM1 learners than in the beginners on Day 4 (**Statistical table 8, reference #64**). These suggest that the beginners frequently search within certain places, compared to the BM1 learners. Other than that, strategy and network structures of spatial navigation would resemble between the BM1 learners and beginners (**Fig. 5B, C**).

In the probe test, the BM1 learners explored holes for a longer time than the beginners, regardless of the hole locations (**Fig. 5D**). In a mixed-design 2-way [Instance (beginners, BM1 learners)×Hole (1∼12)] ANOVA for the time spent around each hole, the main effect of instance was significant, *F*(1,51) = 7.23, *p* = 0.01, *η_p_*^2^ = 0.12. Multiple comparisons detected that the time spent around each hole was significantly longer in the BM1 learner than in the beginner, regardless of the hole location.

Network analysis reported that the number of stops and the order was comparable between the BM1 learners and the BM3 beginners (**Fig. 5E**; **Statistical table 9, reference #1, 2**, respectively). In contrast, degree, density, clustering coefficient, shortest path length, and closeness centrality was higher in the BM1 learner, while shortest path length was lower in the BM1 learner compared to the beginner (**Statistical table 9, reference #3, 4, 5, 6, 8,** respectively). These results suggest that the BM1 learners exhibited more various patterns of transitions between nodes in the networks (**Fig. 5E**), leading to a longer hole exploration time in the BM1 learners than in the beginners. Betweenness centrality was significantly lower in the BM1 learners than in the beginners (**Statistical table 9, reference #7**), suggesting that the beginners required certain places relaying between 2 other places (**Fig. 5E**).

## Discussion

### The BM3 requires more computational resources to establish spatial learning than the BM1

Environmental scale space is thought to differentiate the spatial navigation and learning in animals (for review, see Geva-Sagiv et al., 2015). In this study, we developed a larger scale space, 3-meter diameter Barnes maze (BM3, **Fig. 1B, D, F**), and compared the performances in the BM3 with those in a conventional scale space, 1-meter diameter Barnes maze (BM1, **Fig. 1A, C, E**), to explore how scale space calibrates spatial learning within and between BM tasks. Total 111 mice were subjected to this study, and each was assigned to any one of 7 cohorts (**Table 1**).

We first demonstrated that spatial learning in the BM3 was established. The learning curves in the conventional features of the number of errors, latency and travel distance were decreased and converged across training (**Fig. 2A-C**). The usage of optimal and suboptimal navigation strategy (i.e. spatial and serial strategy, respectively) increased, while use of naive strategy (i.e. random strategy) decreased across training (**Fig. 2G**). In the probe test, the exploration time in the goal hole was significantly longer than those in the other holes (**Fig. 2D**).

Next, we explored unique behavioral features in spatial learning in the BM3 compared to the BM1. In the training phase, all of the number of errors, latency and travel distance were constantly higher in the BM3 than in the BM1 (**Fig. 2D-F**). The learning curves of them were more gradual in the BM3 than in the BM1 (**Fig. 2A-C**), indicating that the spatial learning in the BM3 takes longer time until convergence than that in the BM1. The usage of spatial strategy was lower, while that of the random strategy was higher in the BM3 than in the BM1 (**Fig. 2G**). The BM task can be viewed as a problem to find optimal navigation strategy (i.e. spatial strategy) through trial and error using a variety of strategies. In line with this, the BM3 would demand more trial and error iterations to solve this problem. In the probe test, exploration time in the target hole was significantly shorter in the BM3 than in the BM1, while those in the other holes were virtually comparable between the BMs (**Fig. 2H**). Thus, as expected, spatial representation would be more inaccurate in the BM3 than in the BM1, since the BM3 would require more computational resources to establish learning than the BM1. However, to our knowledge, this is the first result demonstrating the effect of space scale on a goal-directed spatial learning, Barnes maze, in mice.

The increase in demand for computational resources in the BM3 could be partially explained by reducing spatial resolution of place cells in larger scale spaces. Several previous studies have reported that the size of place field per place cell in larger spaces was larger than those in smaller spaces (Fenton et al., 2008; Harland et al., 2021; Kjelstrup et al., 2008; O’Keefe & Burgess, 1996; Park et al., 2011; Rich et al., 2014). Nevertheless, in the nature of spatial navigation in animals, multiscale space representation should coexist so that animals can navigate from the order of centimeters to the order of kilometers (Geva-Sagiv et al., 2015). Indeed, a human fMRI study demonstrated a network of brain regions that were activated in either or both the navigation in large and small scale spaces (Li et al., 2021). Hence, more computational resources might be allocated in the BM3 to process multiscale spatial representations in parallel.

Previously reported effects of scopolamine hydrobromide, a non-selective muscarinic acetylcholine receptor antagonist, in the BM1 (Suzuki & Imayoshi, 2017) was limitedly replicated in the BM3 in this study. Namely, the mice treated with scopolamine in the probe test in the BM1 searched for a longer time around the goal hole, while for a shorter time around other holes far from the goal (Suzuki & Imayoshi, 2017). Such a focal search pattern was also observed in scopolamine-treated mice in the probe test of the Morris water maze (Huang et al., 2011; Lo et al., 2014). But in this study, the scopolamine-treated mice showed neither longer exploration time for the goal nor shorter exploration time for the other holes in the BM3 (**Extended Data Fig. 2-2A**). Only in the hole neighboring the goal, the scopolamine-treated mice explored for significantly longer time than in the non-treated mice (**Extended Data Fig. 2-2A**). In the BM1, significant differences between scopolamine-treated and non-treated groups were observed in all network measures except number of stops (Suzuki & Imayoshi, 2017). In the BM3, all measures except number of stops were comparable between treated and non-treated groups (**Extended Data Fig. 2-2B**). These results indicate that the effects of scopolamine might be more obvious in the BM1 than in the BM3, yet contributions of cholinergic neurons in spatial learning in larger scale spaces have not been reported previously.

We found unique network structures in navigational trajectories in the BM3 and the BM1, in both of the training and the probe test. In the training, the number of stops, order, degree and shortest path length was higher in the BM3 than in the BM1 regardless of training days (**Fig. 3A-C, F**), suggesting that the BM3 prompted the mice to generate more nodes (i.e. stopping points) on the field and transition between them regardless of learning progression. Density and closeness centrality was significantly lower in the BM3 than in the BM1 on Day 1, while that was comparable between the BMs on Day 6 (**Fig. 3D, H**). Both measures within the BM3 significantly increased from Day 1 to Day 12, while those within the BM1 were comparable between Day 1 and Day 6 (**Fig. 3D, H**). On the other hand, betweenness centrality was comparable between the BM1 and BM3 on Day 1, while that was significantly higher in the BM3 than in the BM1 on Day 6 (**Fig. 3G**). Betweenness centrality within the BM3 was comparable between Day 1 and Day 12, while that within the BM1 significantly decreased from Day 1 to Day 6 (**Fig. 3G**). Thus, density, closeness centrality and betweenness centrality would be representative measures, contrasting spatial learning in larger scale spaces with that in smaller scale spaces. In the probe test, mice in both BM1 and BM3 explored the field without the goal for 5 min. The number of stops and order was comparable between the BM1 and the BM3 (**Fig. 3I, J**), while the other network measures showed significant differences between the BMs (**Fig. 3K-P**). These results suggest that the number of network nodes generated in a given time is independent from scale space, while the transition patterns between the nodes are calibrated by scale space, as transitions between the nodes were more sparse in the BM3 than in the BM1. Betweenness centrality was significantly higher in the BM3 than in the BM1, indicating that the mice in the BM3 required more places relaying to other 2 places.

Nodes and links in a navigational network were calculated from stopping coordinates and transitions between nodes, respectively (see Material and Methods). It can be expected that the brain would represent not only geometric structures (e.g. coordinates, distances, boundaries) of the nodes, but also topological structures (e.g. degree, betweenness, closeness centrality) of them. Early psychological study suggested that rats represent topological structures of a maze of a conventional scale (Poucet & Herrmann, 2001). Recently proposed algebraic topological model predicted that the hippocampal place cell ensembles with a certain firing rate, place field size and population size are capable of encoding topological signatures of environments of conventional scales (2 × 2 m) within several minutes (Dabaghian et al., 2012). Furthermore, one electrophysiological study in rats demonstrated that topological structures of a shape-changeable track (4 m length) might be encoded in place cells in the hippocampal CA1, as these place fields were preserved even if configuration of the track was varied in Cartesian coordinates, as long as the topological structure was maintained (Dabaghian et al., 2014). For neural correlates in topological expression of larger scale space, one human fMRI study demonstrated that degree and closeness centrality at each street in London’s Soho correlated with the right posterior and right anterior hippocampal activities while participants virtually visited there (Javadi et al., 2017). In common to both smaller and larger scale spaces, these evidences suggest that topological structures or measures are expressed in the hippocampus and presumably CA1 place cells. Topological expression would reduce computational cost for spatial representation, rather than expressing them in a geometrically precise manner (Dabaghian et al., 2014). This benefit would enable a hierarchical network representation of large environments: a large environment is represented by a network such that each node is a spatial representation of a local region in the large environment and each link represents a connection between a pair of the nodes (Poucet, 1993; Wolbers & Wiener, 2014). Thus, we expect that the benefit of topological coding might be higher at navigations in larger scale spaces than in smaller scale spaces, with a consequence of a divergent neural expression of network structures in the larger scale spaces, from those in the smaller scale spaces.

### BM learning in a scale space is more facilitated by prior BM learning in equal or larger scale spaces

Next, we explored whether scale spaces calibrate not only the present but also subsequent spatial learning. When animals learn a task, they learn not only immediate task solutions *per se*, but also how to find the solutions: Harlow’s “learning set” ( Harlow, 1949). This meta-learning process leads to subsequent few-shot learning in variants of the initially learned task (for review, see Wang, 2021). Therefore, if this meta-learning process can be activated also in spatial learning between different scale BMs, prior learning in either BM scale (e.g. BM3) would facilitate subsequent spatial learning in the other BM scale (e.g. BM1).

The BM3 learners and the BM1’ learners were engaged in the BM3 task and the BM1’ task as prior learning, respectively. Then, both were tested in the BM1 task. The beginners were engaged only in the BM1. We found that prior learning in the BM tasks facilitates subsequent spatial learning in the BM1. During the training, the BM learners exhibited optimal navigation in the subsequent BM1 task compared to the beginners (**Fig. 4A, B**). Not predicted by our hypothesis, the accuracy of spatial representation in the subsequent BM1 was highest in the BM3 learners, as the exploration time around the goal hole was highest in the BM3 learners (**Fig. 4D**). Thus, prior learning in a large scale space would elaborate spatial representation in the subsequent spatial learning in a smaller scale space.

We found that this facilitation effect between BM1 and BM3 was asymmetric, as the facilitation effects from the prior learning in the BM1 to subsequent learning in the BM3 was limited. The number of errors was comparable between the BM1 learner and the beginner (**Fig. 5A**), and the usage of the navigation strategies was virtually comparable between the BM1 learners and beginners (**Fig. 5B**). Meanwhile, the latencies and travel distances were significantly lower for the BM1 learners (**Fig. 5A**). In the probe test, exploration time of holes was significantly longer in the BM1 learners than the beginners, regardless of hole locations (**Fig. 5D**). These results indicated that the BM1 learners took less time to explore places other than places around holes in the subsequent BM3 task, since they could exploit incomplete knowledge of the BM task (i.e. the goal always located at any one of peripheral holes) that was acquired through the prior learning in the BM1.

Many studies have reported meta-learning in a wide variety of tasks such as schema learning (Tse et al., 2007), multiple decision making (Rosenberg et al., 2021), structure learning (Braun et al., 2010), and spatial learning (Alonso et al., 2021; Baraduc et al., 2019; Ocampo et al., 2018). In common with these studies, the facilitation from prior learning to subsequent learning was observed if both prior and subsequent learning were performed under a variant of common task rule in virtually the same size of scale space. Our results support the idea that the BM3 and the BM1 are processed as a variant of the same BM task, as shown in the facilitation in the BM learners. However, we also expect that this facilitation is calibrated by scale spaces at each learning instance, namely if a scale space of a subsequent learning (i.e. BM1) is equal or smaller than that at a prior learning (i.e. BM3 or BM1’), the facilitation effect becomes obvious (i.e. BM3 or BM1’ learners vs. beginners in the BM1). In this case, the BM3, BM1’ and BM1 task would be processed as a variant of a BM task for the BM learners. In contrast, if a scale space of a subsequent learning (i.e. BM3) is larger than that at a prior learning (i.e. BM1), the facilitation effect is limited (i.e. BM1 learners vs. beginners in the BM3).

One possible neural circuit underlying the scale space dependent facilitation in spatial learning would involve the hippocampus and medial prefrontal cortex (mPFC). The hippocampus is activated to encode a novel context, rather than contexts that fit into a previously learned one (for review, see Alonso et al., 2020). Specifically, NMDA receptors in the hippocampal CA3 contribute to novelty detection on the CA1 place cells (Dragoi & Tonegawa, 2013). In a prior and novel BM learning, a memory engram encoding not only spatial representation of the immediate BM, but also a learning set of the BM task would be generated at the hippocampus. Then, the memory engram is transferred from the hippocampus to networks in the cortex through the consolidation process within a couple of weeks.

The mPFC would involve classifying whether a current context fits into existing ones, and suppressing hippocampal activity during memory encoding, unless the context is novel and unlike previously acquired one (van Kesteren et al., 2012). If a subsequent BM learning is a variant of the prior BM learning, a completely new memory engram would be generated little at the hippocampus, because the mPFC infers the current context as similar to the prior one, and suppresses the hippocampal activities. The existing memory engram that has been transferred to the cortex is directly activated bypassing the hippocampus, and slightly tuned to solve the task, leading to few-shot learning (Alonso et al., 2020). The retrosplenial cortex (RSC) and anterior cingulate cortex (ACC) processing decision making based on previous experiences would be additional candidates implementing the facilitation effect by prior learning (Alonso et al., 2020), as the expression of immediate early gene, Arc and Zif68, increased in the RSC and the ACC, when a previously acquired schema was activated (Tse et al., 2007; Wang et al., 2012).

Since the BM3 required more computational resources to learn the task than the BM1 (**Fig. 2**), a relatively larger memory engram should be recruited for the BM3 than for the BM1, for encoding spatial representation of the maze with a learning set of the task. If mice experienced the BM3 task as a prior learning followed by the BM1 task as a subsequent learning, a larger memory engram is generated at the hippocampus during the prior learning. Then, the memory engram is transferred from the hippocampus to the cortex, through the blank between the prior and subsequent learning. In the subsequent BM1 learning, a part of the already acquired memory engram is slightly tuned to solve the task, while information that had to be encoded as a completely new memory engram in the hippocampus became rare. Owing to this, the BM3 learners could learn the subsequent BM1 task quickly, while the beginners had to generate a new memory engram to the hippocampus. In contrast, if mice experienced the BM1 first, a smaller memory engram is generated. Hence, only tuning the existing engram would be not enough to learn the subsequent BM3 task, requiring a certain amount of new memory engram in the hippocampus. Thus, the facilitation by prior BM1 learning would be limited in the subsequent BM3 learning.

Finally, the facilitation of spatial learning between the BM1 and the BM3 would result in the differences of network structure during the training. The facilitation from the BM3 to BM1 differentiated order, degree, shortest path length and betweenness centrality after Day 1, while evened out the number of stops between the BM3 learners and the beginners (**Fig. 4C**). Conversely, the facilitation from the BM1 to BM3 evened out order, degree, shortest path length and betweenness centrality, while differentiated the number of stops between the BM1 learners and the beginners (**Fig. 5C**). Specifically, betweenness centrality would be a representative measure for a facilitation of spatial learning. In the BM3 to BM1 facilitation, decrease of betweenness centrality in the BM3 learners across training was significant and more steep than that in the beginners. In the BM1 to BM3 facilitation, such differences compared to the beginners were not observed in the BM1 learners. It is expected that brain regions implementing topological coding would be changed through prolonged exposure to the environment (Javadi et al., 2017). Our results suggest that such a topological coding in the subsequent BM space would be changed through a prior and a subsequent BM learning.

## Supporting information

Statistical table 1

Statistical table 2

Statistical table 3

Statistical table 4

Statistical table 5

Statistical table 6

Statistical table 7

Statistical table 8

Statistical table 9

## Acknowledgements

We thank Mami Matsumoto, Masako Tanaka, and Ryosuke Okuda for technical help. We appreciate Adam T. Guy PhD. for critiquing the manuscript.

---

**Extended Data Figure 2-1.**
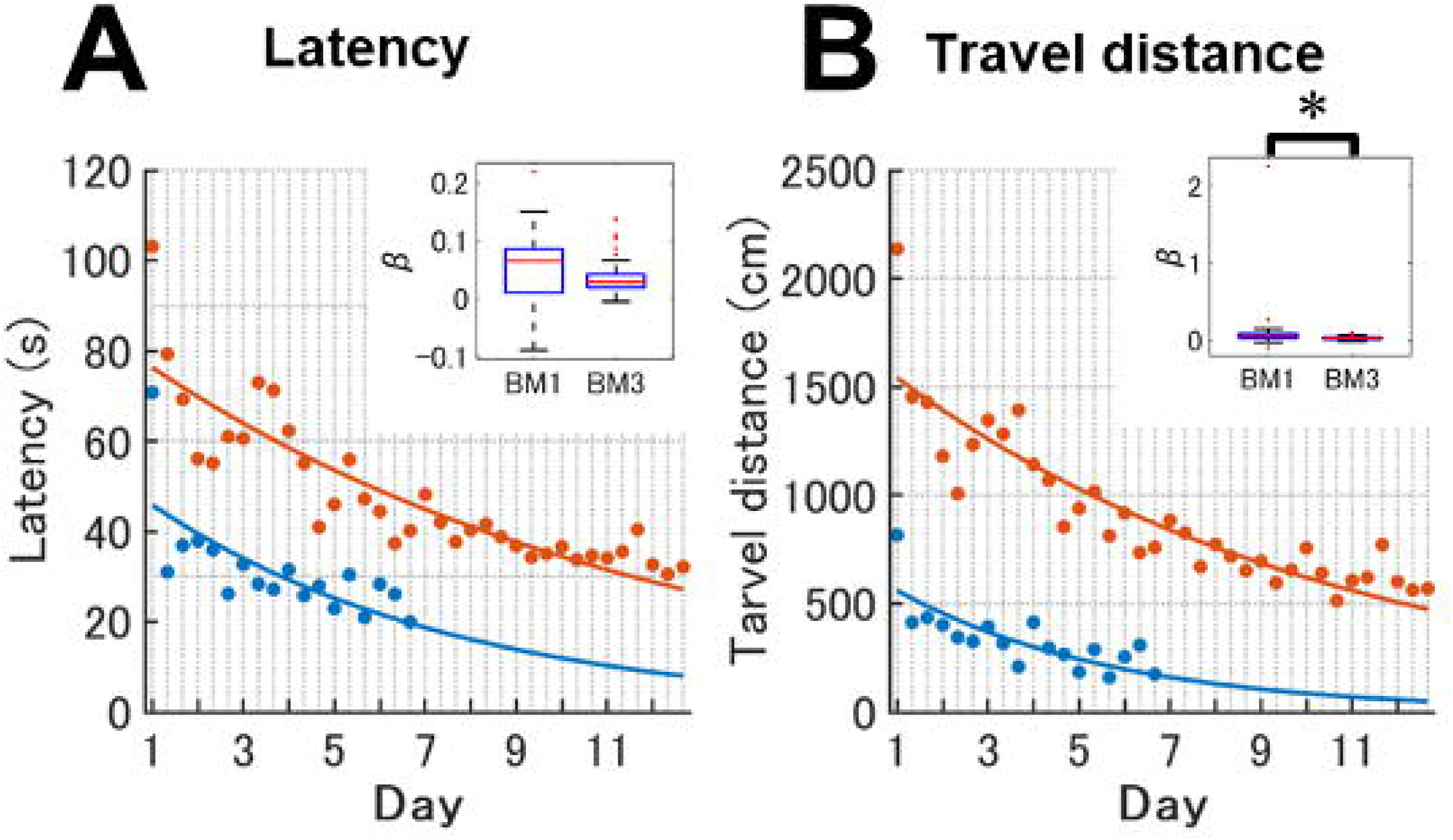
Learning curves fitted for raw values of latency and travel distance. While the BM1 and the BM3 would have different scale lower boundaries of latency and travel distance, respectively, while latency and travel distance should be compared between the BM1 and the BM3 within the same scale. So, latency and travel distance were normalized across trials so that the values range between 0 and 1 (Fig. 2B, C). ***A***, A learning curve for raw values of latency. The vertical and horizontal axis indicates latency in second and training days, respectively. The same curve fitting method was used as Fig. 2B. ***B***, A learning curve for the raw values of travel distance. The vertical and horizontal axis indicates travel distance in centimeters and training days, respectively. The same curve fitting method was used as Fig. 2C. The small inset in panel A and B is the distribution of estimated decay parameter β in the BM1 and the BM3. Asterisk indicates significant differences between the BM1 and the BM3.

**Extended Data Figure 2-2.**
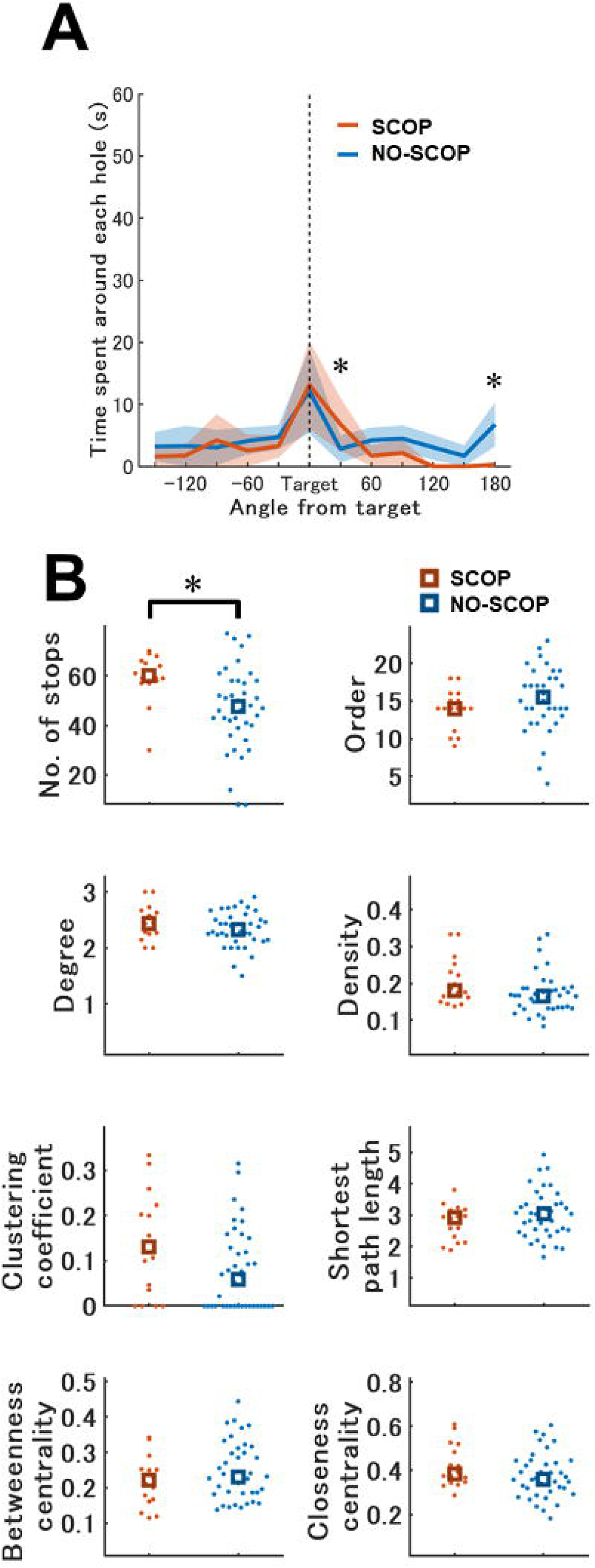
Effects of scopolamine administration on the BM3 probe test. ***A,*** Time spent around each hole in the probe test. Asterisks indicate significant differences between the two groups. Blue and red represent the NO-SCOP (n = 40) and the SCOP (n = 16) mouse groups, respectively. ***B,*** Network features in the BM3 probe test. Asterisks indicate significant differences between the two groups. Scopolamine treatment induced focal search around the target hole without obvious changes of navigation networks.

**Extended Data Figure 3-1.**
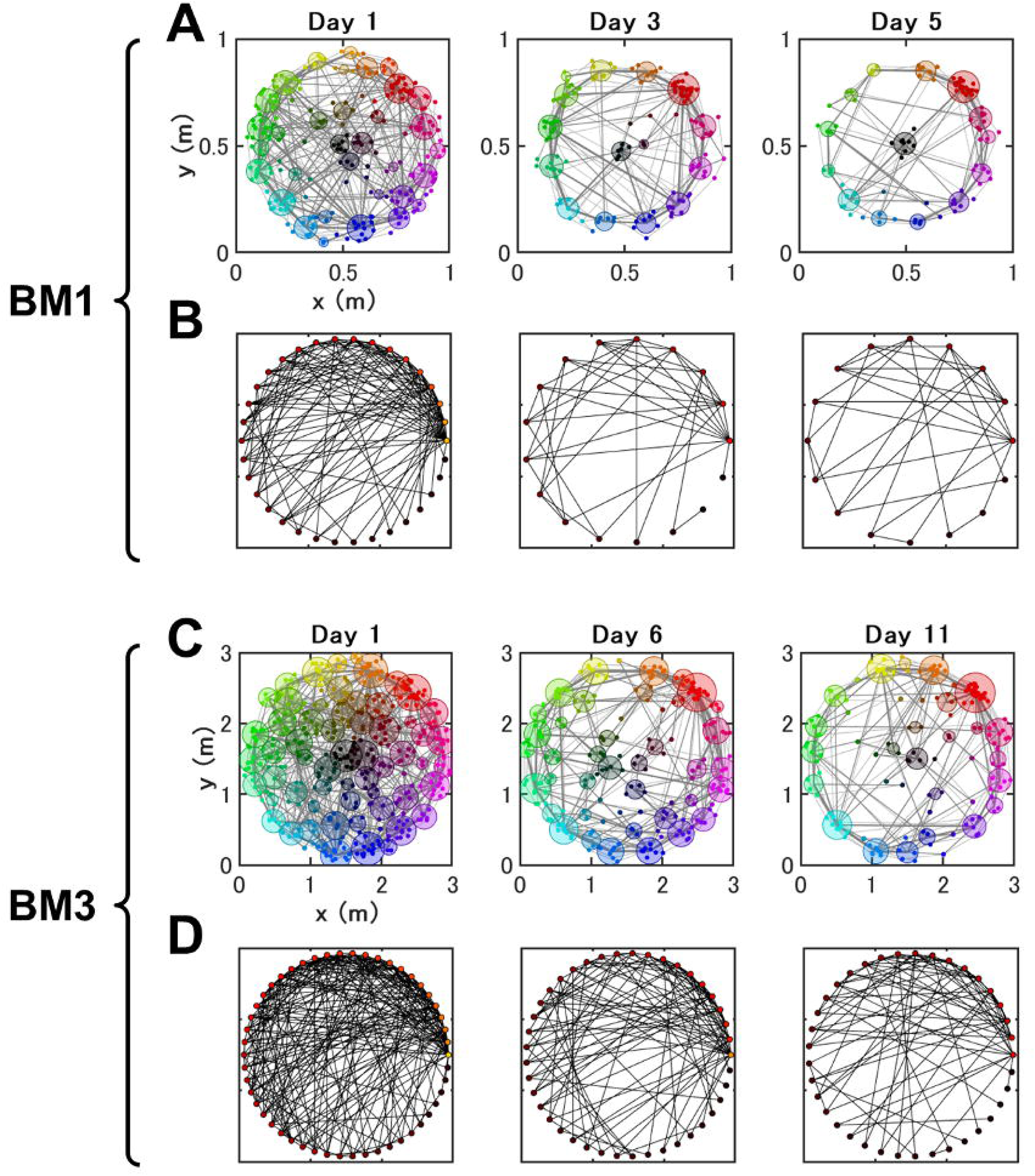
Visualization of temporal changes in the exploration networks during spatial learning in the BM1 and BM3. ***A,*** Temporal changes in the global exploration networks in the BM1 spatial learning. Network structures of exploratory behaviors formed by the dynamic node generation method are plotted (Suzuki & Imayoshi, 2017). Small colored dots and light gray lines represent nodes and links in each mouse (local network). The nodes are located on Cartesian coordinates of the BM1. Nodes are color-coded depending on their polar coordinates. To visualize the global network structure, all local networks of the mice on a single training day were projected on a single plane. Colored larger circles and dark gray lines are global nodes and links in global networks, respectively. Local nodes belong to any one of the global nodes. Likewise, a set of local links are summarized as a global link. The size of a global node is based on log-transformation of the number of nodes that belong to the global node. Likewise, the thickness of a global link is log-transformation of the number of links that belong. ***B,*** Topological expression of BM1 global networks. Global nodes were sorted by rank-order of degree and plotted on polar coordinates, so that the node with the highest degree was located at 0 degrees while the lowest one was located at 360 degrees. Circles and lines represent global nodes and links, respectively. Global nodes were ranked to any one of 30 ranks depending on its degree within each group. Then, they were colored according to the rank; the rank first node has a lot of links and is colored by white, and rank 30th node has few links and is colored by black. ***C, D,*** Temporal changes in the global networks and their topological expressions of the BM3 spatial learning were displayed.

**Extended Data Figure 4-1.**
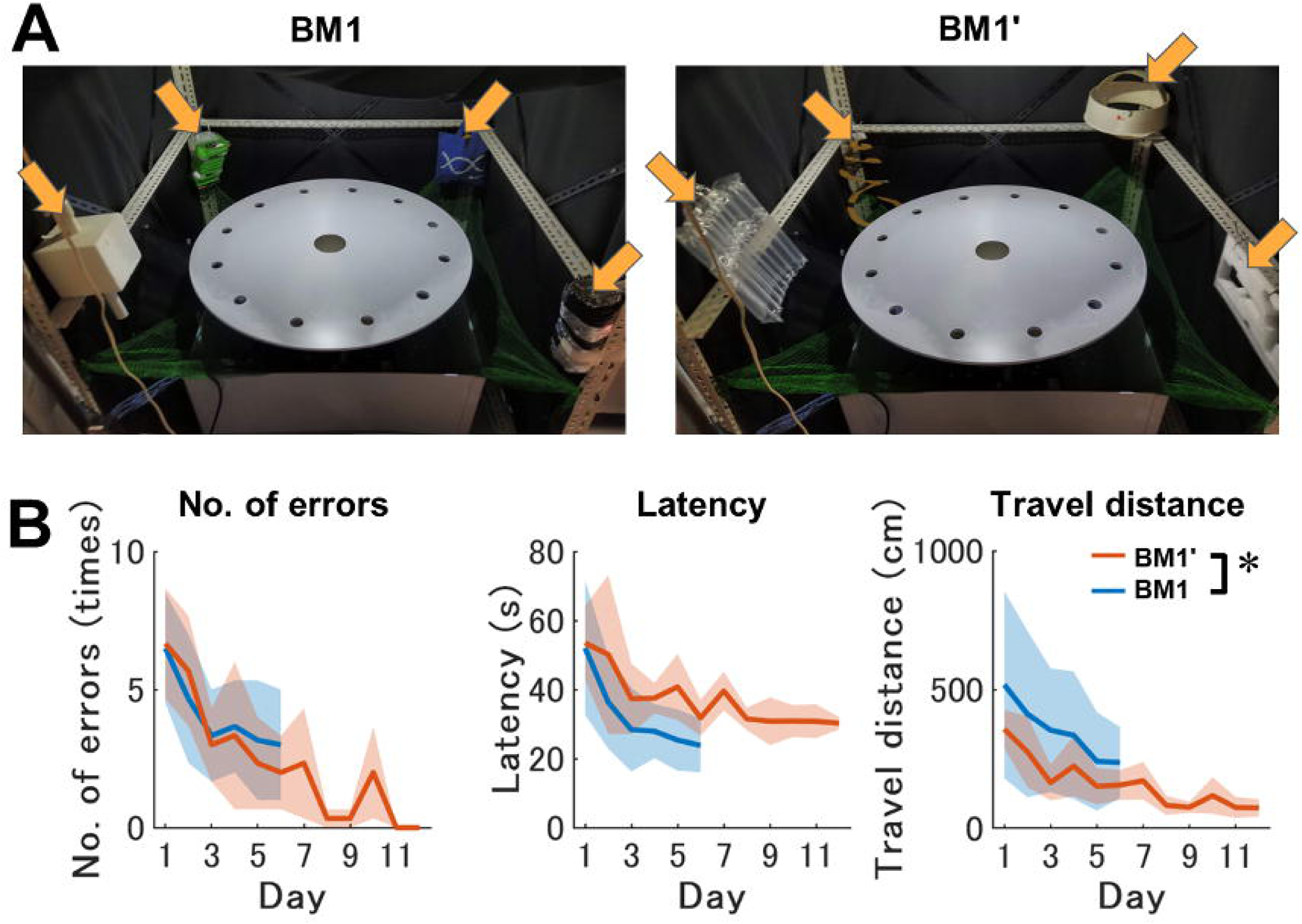
The experiment setup and spatial learning curves in BM1 and BM1’. ***A,*** Spatial cues used in the BM1 (left panel) and the BM1’ (right panel) were indicated with orange arrows. Note that all setups other than the cues were identical between the two mazes. ***B,*** Daily basis learning curves across training periods in conventional features in the BM1 and BM1’. These scores were averaged over 3 trials per day. From left, the measured values of number of errors, latency and travel distance are displayed. Only in a mixed-design 2-way [Spatial cues (BM1, BM1’)×Day (1∼6)] ANOVA for travel distance, the main effect of spatial cues was significant, *F*(1,49) = 6.72, *p* = 0.01, *η_p_*^2^ = 0.12. Asterisks indicate significant differences between the two groups.

**Extended Data Figure 4-2.**
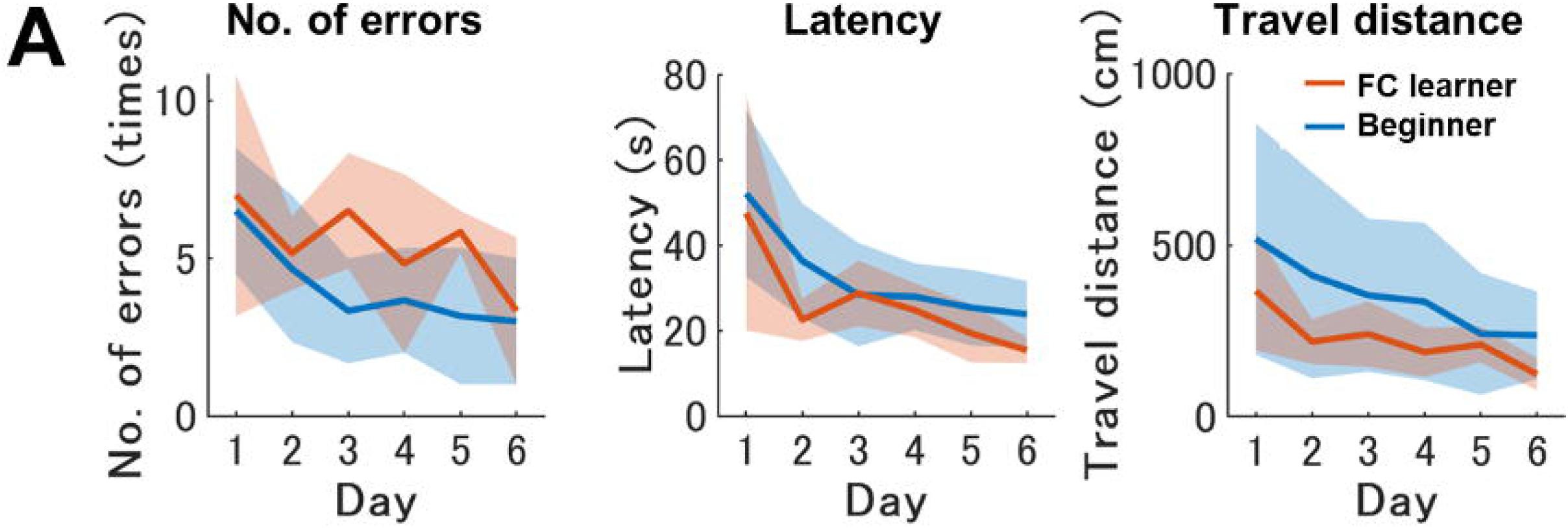
Prior fear-conditioning experience does not impact on the subsequent BM1 spatial learning. ***A,*** Daily basis learning curves of the CFC learner (n = 8) and Beginner (n = 34) across training periods in conventional features in the BM1. These scores were averaged over 3 trials per day. From left, the measured values of number of errors, latency and travel distance are displayed. Mixed-design 2-way [Instance (CFC learner, Beginner)×Day (1∼6)] ANOVA detected neither main effect of Instance nor interaction between instance and day for all features.

**Extended Data 1.** Codes for data acquisition and analysis.

