## Extended Data Table 1 for "Scale space calibrates present and subsequent spatial learning in Barnes maze in mice"

Summary for behavioral features.

| Analysis | Feature | Definition |
| --- | --- | --- |
| Conventional | No. of errors (times) | Sum of the number of visits for areas within a radius of dummy holes, where the radius is one fifth of distance between 2 holes. |
|  | Latency (s) | Latency until entry to the escape box. |
|  | Travel distance (mm) | Sum of norms of moving vector between successive 2 frames of a single trial movie. |
|  | Time spent around each hole (s) | Total duration that a mouse stayed at an area within a radius of hole, where the radius is one fifth of distance between 2 holes. |
| Strategy | Spatial | If movement history during a trial satisfied both the following conditions,  (1) Number of crossing the quadrants = 0,  (2) Number of errors ＜ 3. |
|  | Serial | If movement history during a trial satisfied all of the following conditions,  (1) Number of crossing the quadrants ＜ 3,  (2) Order of visits to holes was sequential,  (3) The strategy has not been classified to the Spatial. |
|  | Random | If movement history during a trial satisfied all of the following conditions,  (1) The strategy has been classified to neither the Spatial nor the Serial. |
| Network | No. of stops | The number of stopping coordinates in the network. |
|  | Order | The number of nodes bundling stopping coordinates in the network. |
|  | Degree | The number of links connected to a node in the network. |
|  | Density | The ratio of the number of links that are actually present links in a network to the number of links that are theoretically possible in the network. |
|  | Clustering coefficient | The probability that two neighbors of a given node are themselves neighbors. |
|  | Shortest path length | The path such that no shorter path exists in the network, and its length is quantified as the number of links traversed along the shortest path between any 2 nodes. |
|  | Betweenness centrality | The extent to which a given node lies on shortest paths between other nodes. |
|  | Closeness centrality | The inverse of the sum of path length from a given node to other nodes. |

Note. Adapted from Suzuki & Imayoshi 2017. Behavioral features analyzed in the conventional, strategy, and network analyses are summarized. Further details are described in Material and Methods.
