## Supplementary material for "Scale space calibrates present and subsequent spatial learning in Barnes maze in mice": Statistical table 1

Statistical results of strategy analysis for the BM1 and the BM3 in the training phase.

| Manuscript reference # | Figure | Measure | Comparison | Within | Data structure | Type of test | Statistic | p | Correction | ES |
| --- | --- | --- | --- | --- | --- | --- | --- | --- | --- | --- |
| 1 | Figure 2G | Random | BM1 vs. BM3 | Day 1 | No assumption | Wilcoxon rank-sum test | z = -1.11 | 0.2668 | Bonferroni (0.05 / 6 days) | r = -0.12 |
| 2 | Figure 2G | Random | BM1 vs. BM3 | Day 2 | No assumption | Wilcoxon rank-sum test | z = -1.91 | 0.0563 | Bonferroni (0.05 / 6 days) | r = -0.20 |
| 3 | Figure 2G | Random | BM1 vs. BM3 | Day 3 | No assumption | Wilcoxon rank-sum test | z = -2.01 | 0.0439 | Bonferroni (0.05 / 6 days) | r = -0.21 |
| 4 | Figure 2G | Random | BM1 vs. BM3 | Day 4 | No assumption | Wilcoxon rank-sum test | z = -0.44 | 0.6592 | Bonferroni (0.05 / 6 days) | r = -0.05 |
| 5 | Figure 2G | Random | BM1 vs. BM3 | Day 5 | No assumption | Wilcoxon rank-sum test | z = -2.68 | 0.0074* | Bonferroni (0.05 / 6 days) | r = -0.28 |
| 6 | Figure 2G | Random | BM1 vs. BM3 | Day 6 | No assumption | Wilcoxon rank-sum test | z = -2.74 | 0.0061* | Bonferroni (0.05 / 6 days) | r = -0.29 |
| 7 | Figure 2G | Serial | BM1 vs. BM3 | Day 1 | No assumption | Wilcoxon rank-sum test | z = -1.09 | 0.2748 | Bonferroni (0.05 / 6 days) | r = -0.12 |
| 8 | Figure 2G | Serial | BM1 vs. BM3 | Day 2 | No assumption | Wilcoxon rank-sum test | z = 1.66 | 0.0965 | Bonferroni (0.05 / 6 days) | r = 0.18 |
| 9 | Figure 2G | Serial | BM1 vs. BM3 | Day 3 | No assumption | Wilcoxon rank-sum test | z = 1.00 | 0.3173 | Bonferroni (0.05 / 6 days) | r = 0.11 |
| 10 | Figure 2G | Serial | BM1 vs. BM3 | Day 4 | No assumption | Wilcoxon rank-sum test | z = -0.30 | 0.7667 | Bonferroni (0.05 / 6 days) | r = -0.03 |
| 11 | Figure 2G | Serial | BM1 vs. BM3 | Day 5 | No assumption | Wilcoxon rank-sum test | z = 0.43 | 0.6658 | Bonferroni (0.05 / 6 days) | r = 0.05 |
| 12 | Figure 2G | Serial | BM1 vs. BM3 | Day 6 | No assumption | Wilcoxon rank-sum test | z = 1.49 | 0.1361 | Bonferroni (0.05 / 6 days) | r = 0.16 |
| 13 | Figure 2G | Spatial | BM1 vs. BM3 | Day 1 | No assumption | Wilcoxon rank-sum test | z = 3.20 | 0.0014* | Bonferroni (0.05 / 6 days) | r = 0.34 |
| 14 | Figure 2G | Spatial | BM1 vs. BM3 | Day 2 | No assumption | Wilcoxon rank-sum test | z = 1.39 | 0.1643 | Bonferroni (0.05 / 6 days) | r = 0.15 |
| 15 | Figure 2G | Spatial | BM1 vs. BM3 | Day 3 | No assumption | Wilcoxon rank-sum test | z = 2.85 | 0.0043* | Bonferroni (0.05 / 6 days) | r = 0.30 |
| 16 | Figure 2G | Spatial | BM1 vs. BM3 | Day 4 | No assumption | Wilcoxon rank-sum test | z = 0.82 | 0.4106 | Bonferroni (0.05 / 6 days) | r = 0.09 |
| 17 | Figure 2G | Spatial | BM1 vs. BM3 | Day 5 | No assumption | Wilcoxon rank-sum test | z = 3.50 | 0.0005* | Bonferroni (0.05 / 6 days) | r = 0.37 |
| 18 | Figure 2G | Spatial | BM1 vs. BM3 | Day 6 | No assumption | Wilcoxon rank-sum test | z = 2.40 | 0.0162 | Bonferroni (0.05 / 6 days) | r = 0.25 |

Note. Asterisks indicate statistically significant differences. ES: effect size.
