## Supplementary material for "Scale space calibrates present and subsequent spatial learning in Barnes maze in mice": Statistical table 2

Statistical results of network analysis for the BM1 and the BM3 in the training phase.

| Manuscript reference # | Figure | Measure | Comparison | Within | Data structure | Type of test | Statistic | p | Correction | ES |
| --- | --- | --- | --- | --- | --- | --- | --- | --- | --- | --- |
| 1 | Figure 3A | Number of stops | BM1 vs. BM3 | Day 1 | No assumption | Wilcoxon rank-sum test | z = -6.70 | 0.0000* | Bonferroni (0.05 / 6 days) | r = -0.71 |
| 2 | Figure 3A | Number of stops | BM1 vs. BM3 | Day 2 | No assumption | Wilcoxon rank-sum test | z = -6.50 | 0.0000* | Bonferroni (0.05 / 6 days) | r = -0.69 |
| 3 | Figure 3A | Number of stops | BM1 vs. BM3 | Day 3 | No assumption | Wilcoxon rank-sum test | z = -6.74 | 0.0000* | Bonferroni (0.05 / 6 days) | r = -0.71 |
| 4 | Figure 3A | Number of stops | BM1 vs. BM3 | Day 4 | No assumption | Wilcoxon rank-sum test | z = -6.22 | 0.0000* | Bonferroni (0.05 / 6 days) | r = -0.66 |
| 5 | Figure 3A | Number of stops | BM1 vs. BM3 | Day 5 | No assumption | Wilcoxon rank-sum test | z = -6.31 | 0.0000* | Bonferroni (0.05 / 6 days) | r = -0.67 |
| 6 | Figure 3A | Number of stops | BM1 vs. BM3 | Day 6 | No assumption | Wilcoxon rank-sum test | z = -5.51 | 0.0000* | Bonferroni (0.05 / 6 days) | r = -0.58 |
| 7 | Figure 3B | Order | BM1 vs. BM3 | Day 1 | No assumption | Wilcoxon rank-sum test | z = -6.69 | 0.0000* | Bonferroni (0.05 / 6 days) | r = -0.70 |
| 8 | Figure 3B | Order | BM1 vs. BM3 | Day 2 | No assumption | Wilcoxon rank-sum test | z = -6.34 | 0.0000* | Bonferroni (0.05 / 6 days) | r = -0.67 |
| 9 | Figure 3B | Order | BM1 vs. BM3 | Day 3 | No assumption | Wilcoxon rank-sum test | z = -6.83 | 0.0000* | Bonferroni (0.05 / 6 days) | r = -0.72 |
| 10 | Figure 3B | Order | BM1 vs. BM3 | Day 4 | No assumption | Wilcoxon rank-sum test | z = -6.09 | 0.0000* | Bonferroni (0.05 / 6 days) | r = -0.64 |
| 11 | Figure 3B | Order | BM1 vs. BM3 | Day 5 | No assumption | Wilcoxon rank-sum test | z = -6.65 | 0.0000* | Bonferroni (0.05 / 6 days) | r = -0.70 |
| 12 | Figure 3B | Order | BM1 vs. BM3 | Day 6 | No assumption | Wilcoxon rank-sum test | z = -5.77 | 0.0000* | Bonferroni (0.05 / 6 days) | r = -0.61 |
| 13 | Figure 3C | Degree | BM1 vs. BM3 | Day 1 | No assumption | Wilcoxon rank-sum test | z = -4.70 | 0.0000* | Bonferroni (0.05 / 6 days) | r = -0.50 |
| 14 | Figure 3C | Degree | BM1 vs. BM3 | Day 2 | No assumption | Wilcoxon rank-sum test | z = -5.24 | 0.0000* | Bonferroni (0.05 / 6 days) | r = -0.55 |
| 15 | Figure 3C | Degree | BM1 vs. BM3 | Day 3 | No assumption | Wilcoxon rank-sum test | z = -5.46 | 0.0000* | Bonferroni (0.05 / 6 days) | r = -0.58 |
| 16 | Figure 3C | Degree | BM1 vs. BM3 | Day 4 | No assumption | Wilcoxon rank-sum test | z = -4.39 | 0.0000* | Bonferroni (0.05 / 6 days) | r = -0.46 |
| 17 | Figure 3C | Degree | BM1 vs. BM3 | Day 5 | No assumption | Wilcoxon rank-sum test | z = -5.52 | 0.0000* | Bonferroni (0.05 / 6 days) | r = -0.58 |
| 18 | Figure 3C | Degree | BM1 vs. BM3 | Day 6 | No assumption | Wilcoxon rank-sum test | z = -4.79 | 0.0000* | Bonferroni (0.05 / 6 days) | r = -0.50 |
| 19 | Figure 3D | Density | BM1 vs. BM3 | Day 1 | No assumption | Wilcoxon rank-sum test | z = 4.66 | 0.0000* | Bonferroni (0.05 / 6 days) | r = 0.49 |
| 20 | Figure 3D | Density | BM1 vs. BM3 | Day 2 | No assumption | Wilcoxon rank-sum test | z = 4.73 | 0.0000* | Bonferroni (0.05 / 6 days) | r = 0.50 |
| 21 | Figure 3D | Density | BM1 vs. BM3 | Day 3 | No assumption | Wilcoxon rank-sum test | z = 5.73 | 0.0000* | Bonferroni (0.05 / 6 days) | r = 0.60 |
| 22 | Figure 3D | Density | BM1 vs. BM3 | Day 4 | No assumption | Wilcoxon rank-sum test | z = 3.52 | 0.0004* | Bonferroni (0.05 / 6 days) | r = 0.37 |
| 23 | Figure 3D | Density | BM1 vs. BM3 | Day 5 | No assumption | Wilcoxon rank-sum test | z = 2.78 | 0.0055* | Bonferroni (0.05 / 6 days) | r = 0.29 |
| 24 | Figure 3D | Density | BM1 vs. BM3 | Day 6 | No assumption | Wilcoxon rank-sum test | z = 1.62 | 0.1055 | Bonferroni (0.05 / 6 days) | r = 0.17 |
| 25 | Figure 3D | Density | Day 1 vs. 6 | BM1 | No assumption | Wilcoxon signed-rank test | z = -1.48 | 0.14 | N/A | r = -0.16 |
| 26 | Figure 3D | Density | Day 1 vs. 12 | BM3 | No assumption | Wilcoxon signed-rank test | z = -5.80 | 0.00 | N/A | r = -0.61 |
| 27 | Figure 3E | Clustering coefficient | BM1 vs. BM3 | Day 1 | No assumption | Wilcoxon rank-sum test | z = -1.86 | 0.0634 | Bonferroni (0.05 / 6 days) | r = -0.20 |
| 28 | Figure 3E | Clustering coefficient | BM1 vs. BM3 | Day 2 | No assumption | Wilcoxon rank-sum test | z = -2.34 | 0.0192 | Bonferroni (0.05 / 6 days) | r = -0.25 |
| 29 | Figure 3E | Clustering coefficient | BM1 vs. BM3 | Day 3 | No assumption | Wilcoxon rank-sum test | z = -2.61 | 0.0089 | Bonferroni (0.05 / 6 days) | r = -0.28 |
| 30 | Figure 3E | Clustering coefficient | BM1 vs. BM3 | Day 4 | No assumption | Wilcoxon rank-sum test | z = -1.88 | 0.0599 | Bonferroni (0.05 / 6 days) | r = -0.20 |
| 31 | Figure 3E | Clustering coefficient | BM1 vs. BM3 | Day 5 | No assumption | Wilcoxon rank-sum test | z = -2.15 | 0.0319 | Bonferroni (0.05 / 6 days) | r = -0.23 |
| 32 | Figure 3E | Clustering coefficient | BM1 vs. BM3 | Day 6 | No assumption | Wilcoxon rank-sum test | z = -0.40 | 0.6927 | Bonferroni (0.05 / 6 days) | r = -0.04 |
| 33 | Figure 3F | Shortest path length | BM1 vs. BM3 | Day 1 | No assumption | Wilcoxon rank-sum test | z = -6.29 | 0.0000* | Bonferroni (0.05 / 6 days) | r = -0.66 |
| 34 | Figure 3F | Shortest path length | BM1 vs. BM3 | Day 2 | No assumption | Wilcoxon rank-sum test | z = -5.76 | 0.0000* | Bonferroni (0.05 / 6 days) | r = -0.61 |
| 35 | Figure 3F | Shortest path length | BM1 vs. BM3 | Day 3 | No assumption | Wilcoxon rank-sum test | z = -6.68 | 0.0000* | Bonferroni (0.05 / 6 days) | r = -0.70 |
| 36 | Figure 3F | Shortest path length | BM1 vs. BM3 | Day 4 | No assumption | Wilcoxon rank-sum test | z = -5.82 | 0.0000* | Bonferroni (0.05 / 6 days) | r = -0.61 |
| 37 | Figure 3F | Shortest path length | BM1 vs. BM3 | Day 5 | No assumption | Wilcoxon rank-sum test | z = -7.00 | 0.0000* | Bonferroni (0.05 / 6 days) | r = -0.74 |
| 38 | Figure 3F | Shortest path length | BM1 vs. BM3 | Day 6 | No assumption | Wilcoxon rank-sum test | z = -6.66 | 0.0000* | Bonferroni (0.05 / 6 days) | r = -0.70 |
| 39 | Figure 3G | Betweenness centrality | BM1 vs. BM3 | Day 1 | No assumption | Wilcoxon rank-sum test | z = -2.00 | 0.0453 | Bonferroni (0.05 / 6 days) | r = -0.21 |
| 40 | Figure 3G | Betweenness centrality | BM1 vs. BM3 | Day 2 | No assumption | Wilcoxon rank-sum test | z = -2.67 | 0.0076* | Bonferroni (0.05 / 6 days) | r = -0.28 |
| 41 | Figure 3G | Betweenness centrality | BM1 vs. BM3 | Day 3 | No assumption | Wilcoxon rank-sum test | z = -3.60 | 0.0003* | Bonferroni (0.05 / 6 days) | r = -0.38 |
| 42 | Figure 3G | Betweenness centrality | BM1 vs. BM3 | Day 4 | No assumption | Wilcoxon rank-sum test | z = -3.50 | 0.0005* | Bonferroni (0.05 / 6 days) | r = -0.37 |
| 43 | Figure 3G | Betweenness centrality | BM1 vs. BM3 | Day 5 | No assumption | Wilcoxon rank-sum test | z = -6.07 | 0.0000* | Bonferroni (0.05 / 6 days) | r = -0.64 |
| 44 | Figure 3G | Betweenness centrality | BM1 vs. BM3 | Day 6 | No assumption | Wilcoxon rank-sum test | z = -6.04 | 0.0000* | Bonferroni (0.05 / 6 days) | r = -0.64 |
| 45 | Figure 3G | Betweenness centrality | Day 1 vs. 6 | BM1 | No assumption | Wilcoxon rank-sum test | z = 2.45 | 0.01 | N/A | r = -0.26 |
| 46 | Figure 3G | Betweenness centrality | Day 1 vs. 12 | BM3 | No assumption | Wilcoxon rank-sum test | z = 0.46 | 0.65 | N/A | r = 0.05 |
| 47 | Figure 3H | Closeness centrality | BM1 vs. BM3 | Day 1 | No assumption | Wilcoxon rank-sum test | z = 4.12 | 0.0000* | Bonferroni (0.05 / 6 days) | r = 0.43 |
| 48 | Figure 3H | Closeness centrality | BM1 vs. BM3 | Day 2 | No assumption | Wilcoxon rank-sum test | z = 4.41 | 0.0000* | Bonferroni (0.05 / 6 days) | r = 0.46 |
| 49 | Figure 3H | Closeness centrality | BM1 vs. BM3 | Day 3 | No assumption | Wilcoxon rank-sum test | z = 5.53 | 0.0000* | Bonferroni (0.05 / 6 days) | r = 0.58 |
| 50 | Figure 3H | Closeness centrality | BM1 vs. BM3 | Day 4 | No assumption | Wilcoxon rank-sum test | z = 3.30 | 0.0010* | Bonferroni (0.05 / 6 days) | r = 0.35 |
| 51 | Figure 3H | Closeness centrality | BM1 vs. BM3 | Day 5 | No assumption | Wilcoxon rank-sum test | z = 2.81 | 0.0050* | Bonferroni (0.05 / 6 days) | r = 0.30 |
| 52 | Figure 3H | Closeness centrality | BM1 vs. BM3 | Day 6 | No assumption | Wilcoxon rank-sum test | z = 1.79 | 0.0729 | Bonferroni (0.05 / 6 days) | r = 0.19 |
| 53 | Figure 3H | Closeness centrality | Day 1 vs. 6 | BM1 | No assumption | Wilcoxon signed-rank test | z = -1.43 | 0.15 | N/A | r = -0.15 |
| 54 | Figure 3D | Closeness centrality | Day 1 vs. 12 | BM3 | No assumption | Wilcoxon signed-rank test | z = -5.18 | 0.00 | N/A | r = -0.55 |

Note. Asterisks indicate statistically significant differences. N/A: not applicable. ES: effect size.
