## Supplementary material for "Scale space calibrates present and subsequent spatial learning in Barnes maze in mice": Statistical table 3

Statistical results of strategy analysis for the BM1 and the BM3 in the BM1 probe test.

| Manuscript reference # | Figure | Measure | Comparison | Within | Data structure | Type of test | Statistic | p | Correction | ES |
| --- | --- | --- | --- | --- | --- | --- | --- | --- | --- | --- |
| 1 | Figure 3I | Number of stops | BM1 vs. BM3 | Probe test | No assumption | Wilcoxon rank-sum test | z = 0.81 | 0.42 | N/A | r = 0.09 |
| 2 | Figure 3I | Order | BM1 vs. BM3 | Probe test | No assumption | Wilcoxon rank-sum test | z = 0.86 | 0.39 | N/A | r = 0.10 |
| 3 | Figure 3I | Degree | BM1 vs. BM3 | Probe test | No assumption | Wilcoxon rank-sum test | z = 6.03 | 0.00* | N/A | r = 0.70 |
| 4 | Figure 3I | Density | BM1 vs. BM3 | Probe test | No assumption | Wilcoxon rank-sum test | z = 2.51 | 0.01* | N/A | r = 0.29 |
| 5 | Figure 3I | Clustering coefficient | BM1 vs. BM3 | Probe test | No assumption | Wilcoxon rank-sum test | z = 3.93 | 0.00* | N/A | r = 0.46 |
| 6 | Figure 3I | Shortest path length | BM1 vs. BM3 | Probe test | No assumption | Wilcoxon rank-sum test | z = -3.36 | 0.00* | N/A | r = -0.39 |
| 7 | Figure 3J | Betweenness centrality | BM1 vs. BM3 | Probe test | No assumption | Wilcoxon rank-sum test | z = -5.61 | 0.00* | N/A | r = -0.65 |
| 8 | Figure 3J | Closeness centrality | BM1 vs. BM3 | Probe test | No assumption | Wilcoxon rank-sum test | z = 3.47 | 0.00* | N/A | r = 0.40 |

Note. Asterisks indicate statistically significant differences. N/A: not applicable. ES: effect size.
