## Supplementary material for "Scale space calibrates present and subsequent spatial learning in Barnes maze in mice": Statistical table 4

Statistical results of strategy analysis for the BM1’ learner, the BM3 learner, and the Beginner in the BM1 training.

| Manuscript reference # | Figure | Measure | Comparison | Within | Data structure | Type of test | Statistic | p | Correction | ES |
| --- | --- | --- | --- | --- | --- | --- | --- | --- | --- | --- |
| 1 | Figure 4B | Random | BM1' learner, BM3 learner, Beginner | Day 1 | No assumption | Kruskall-Wallis test | X^2^ (2) = 7.22 | 0.027 | Bonferroni (0.05 / 6 days) | N/A |
| 2 | Figure 4B | Random | BM1' learner, BM3 learner, Beginner | Day 2 | No assumption | Kruskall-Wallis test | X^2^ (2) = 9.79 | 0.007* | Bonferroni (0.05 / 6 days) | N/A |
| 3 | Figure 4B | Random | BM1' learner vs. BM3 learner | Day 2 | No assumption | Wilcoxon rank-sum test | z = -0.76 | 0.445 | Bonferroni (0.05 / 3 pairs) | r = -0.13 |
| 4 | Figure 4B | Random | BM1' learner vs. Beginner | Day 2 | No assumption | Wilcoxon rank-sum test | z = -2.95 | 0.003* | Bonferroni (0.05 / 3 pairs) | r = -0.41 |
| 5 | Figure 4B | Random | BM3 learner vs. Beginner | Day 2 | No assumption | Wilcoxon rank-sum test | z = -2.07 | 0.038 | Bonferroni (0.05 / 3 pairs) | r = -0.28 |
| 6 | Figure 4B | Random | BM1' learner, BM3 learner, Beginner | Day 3 | No assumption | Kruskall-Wallis test | X^2^ (2) = 19.40 | 0.000* | Bonferroni (0.05 / 6 days) | N/A |
| 7 | Figure 4B | Random | BM1' learner vs. BM3 learner | Day 3 | No assumption | Wilcoxon rank-sum test | z = 1.59 | 0.112 | Bonferroni (0.05 / 3 pairs) | r = 0.26 |
| 8 | Figure 4B | Random | BM1' learner vs. Beginner | Day 3 | No assumption | Wilcoxon rank-sum test | z = -2.78 | 0.005* | Bonferroni (0.05 / 3 pairs) | r = -0.39 |
| 9 | Figure 4B | Random | BM3 learner vs. Beginner | Day 3 | No assumption | Wilcoxon rank-sum test | z = -4.08 | 0.000* | Bonferroni (0.05 / 3 pairs) | r = -0.56 |
| 10 | Figure 4B | Random | BM1' learner, BM3 learner, Beginner | Day 4 | No assumption | Kruskall-Wallis test | X^2^ (2) = 24.38 | 0.000* | Bonferroni (0.05 / 6 days) | N/A |
| 11 | Figure 4B | Random | BM1' learner vs. BM3 learner | Day 4 | No assumption | Wilcoxon rank-sum test | z = -1.51 | 0.131 | Bonferroni (0.05 / 3 pairs) | r = -0.25 |
| 12 | Figure 4B | Random | BM1' learner vs. Beginner | Day 4 | No assumption | Wilcoxon rank-sum test | z = -4.49 | 0.000* | Bonferroni (0.05 / 3 pairs) | r = -0.63 |
| 13 | Figure 4B | Random | BM3 learner vs. Beginner | Day 4 | No assumption | Wilcoxon rank-sum test | z = -3.43 | 0.001* | Bonferroni (0.05 / 3 pairs) | r = -0.47 |
| 14 | Figure 4B | Random | BM1' learner, BM3 learner, Beginner | Day 5 | No assumption | Kruskall-Wallis test | X^2^ (2) = 15.50 | 0.000* | Bonferroni (0.05 / 6 days) | N/A |
| 15 | Figure 4B | Random | BM1' learner vs. BM3 learner | Day 5 | No assumption | Wilcoxon rank-sum test | z = -1.15 | 0.250 | Bonferroni (0.05 / 3 pairs) | r = -0.19 |
| 16 | Figure 4B | Random | BM1' learner vs. Beginner | Day 5 | No assumption | Wilcoxon rank-sum test | z = -3.39 | 0.001* | Bonferroni (0.05 / 3 pairs) | r = -0.47 |
| 17 | Figure 4B | Random | BM3 learner vs. Beginner | Day 5 | No assumption | Wilcoxon rank-sum test | z = -2.93 | 0.003* | Bonferroni (0.05 / 3 pairs) | r = -0.40 |
| 18 | Figure 4B | Random | BM1' learner, BM3 learner, Beginner | Day 6 | No assumption | Kruskall-Wallis test | X^2^ (2) = 15.16 | 0.001* | Bonferroni (0.05 / 6 days) | N/A |
| 19 | Figure 4B | Random | BM1' learner vs. BM3 learner | Day 6 | No assumption | Wilcoxon rank-sum test | z = -0.62 | 0.536 | Bonferroni (0.05 / 3 pairs) | r = -0.10 |
| 20 | Figure 4B | Random | BM1' learner vs. Beginner | Day 6 | No assumption | Wilcoxon rank-sum test | z = -3.23 | 0.001* | Bonferroni (0.05 / 3 pairs) | r = -0.45 |
| 21 | Figure 4B | Random | BM3 learner vs. Beginner | Day 6 | No assumption | Wilcoxon rank-sum test | z = -3.14 | 0.002* | Bonferroni (0.05 / 3 pairs) | r = -0.43 |
| 22 | Figure 4B | Serial | BM1' learner, BM3 learner, Beginner | Day 1 | No assumption | Kruskall-Wallis test | X^2^ (2) = 2.48 | 0.290 | Bonferroni (0.05 / 6 days) | N/A |
| 23 | Figure 4B | Serial | BM1' learner, BM3 learner, Beginner | Day 2 | No assumption | Kruskall-Wallis test | X^2^ (2) = 0.34 | 0.843 | Bonferroni (0.05 / 6 days) | N/A |
| 24 | Figure 4B | Serial | BM1' learner, BM3 learner, Beginner | Day 3 | No assumption | Kruskall-Wallis test | X^2^ (2) = 3.50 | 0.174 | Bonferroni (0.05 / 6 days) | N/A |
| 25 | Figure 4B | Serial | BM1' learner, BM3 learner, Beginner | Day 4 | No assumption | Kruskall-Wallis test | X^2^ (2) = 3.61 | 0.164 | Bonferroni (0.05 / 6 days) | N/A |
| 26 | Figure 4B | Serial | BM1' learner, BM3 learner, Beginner | Day 5 | No assumption | Kruskall-Wallis test | X^2^ (2) = 1.38 | 0.502 | Bonferroni (0.05 / 6 days) | N/A |
| 27 | Figure 4B | Serial | BM1' learner, BM3 learner, Beginner | Day 6 | No assumption | Kruskall-Wallis test | X^2^ (2) = 0.75 | 0.688 | Bonferroni (0.05 / 6 days) | N/A |
| 28 | Figure 4B | Spatial | BM1' learner, BM3 learner, Beginner | Day 1 | No assumption | Kruskall-Wallis test | X^2^ (2) = 3.50 | 0.174 | Bonferroni (0.05 / 6 days) | N/A |
| 29 | Figure 4B | Spatial | BM1' learner, BM3 learner, Beginner | Day 2 | No assumption | Kruskall-Wallis test | X^2^ (2) = 9.35 | 0.174 | Bonferroni (0.05 / 6 days) | N/A |
| 30 | Figure 4B | Spatial | BM1' learner, BM3 learner, Beginner | Day 3 | No assumption | Kruskall-Wallis test | X^2^ (2) = 13.70 | 0.001* | Bonferroni (0.05 / 6 days) | N/A |
| 31 | Figure 4B | Spatial | BM1' learner vs. BM3 learner | Day 3 | No assumption | Wilcoxon rank-sum test | z = -0.35 | 0.724 | Bonferroni (0.05 / 3 pairs) | r = -0.06 |
| 32 | Figure 4B | Spatial | BM1' learner vs. Beginner | Day 3 | No assumption | Wilcoxon rank-sum test | z = 2.46 | 0.014* | Bonferroni (0.05 / 3 pairs) | r = 0.34 |
| 33 | Figure 4B | Spatial | BM3 learner vs. Beginner | Day 3 | No assumption | Wilcoxon rank-sum test | z = 3.62 | 0.000* | Bonferroni (0.05 / 3 pairs) | r = 0.49 |
| 34 | Figure 4B | Spatial | BM1' learner, BM3 learner, Beginner | Day 4 | No assumption | Kruskall-Wallis test | X^2^ (2) = 15.96 | 0.000* | Bonferroni (0.05 / 6 days) | N/A |
| 35 | Figure 4B | Spatial | BM1' learner vs. BM3 learner | Day 4 | No assumption | Wilcoxon rank-sum test | z = 0.83 | 0.409 | Bonferroni (0.05 / 3 pairs) | r = 0.14 |
| 36 | Figure 4B | Spatial | BM1' learner vs. Beginner | Day 4 | No assumption | Wilcoxon rank-sum test | z = 3.48 | 0.001* | Bonferroni (0.05 / 3 pairs) | r = 0.49 |
| 37 | Figure 4B | Spatial | BM3 learner vs. Beginner | Day 4 | No assumption | Wilcoxon rank-sum test | z = 3.33 | 0.001* | Bonferroni (0.05 / 3 pairs) | r = 0.45 |
| 38 | Figure 4B | Spatial | BM1' learner, BM3 learner, Beginner | Day 5 | No assumption | Kruskall-Wallis test | X^2^ (2) = 11.27 | 0.004* | Bonferroni (0.05 / 6 days) | N/A |
| 39 | Figure 4B | Spatial | BM1' learner vs. BM3 learner | Day 5 | No assumption | Wilcoxon rank-sum test | z = 1.58 | 0.114 | Bonferroni (0.05 / 3 pairs) | r = 0.26 |
| 40 | Figure 4B | Spatial | BM1' learner vs. Beginner | Day 5 | No assumption | Wilcoxon rank-sum test | z = 3.24 | 0.001* | Bonferroni (0.05 / 3 pairs) | r = 0.45 |
| 41 | Figure 4B | Spatial | BM3 learner vs. Beginner | Day 5 | No assumption | Wilcoxon rank-sum test | z = 1.78 | 0.075 | Bonferroni (0.05 / 3 pairs) | r = 0.24 |
| 42 | Figure 4B | Spatial | BM1' learner, BM3 learner, Beginner | Day 6 | No assumption | Kruskall-Wallis test | X^2^ (2) = 16.07 | 0.000* | Bonferroni (0.05 / 6 days) | N/A |
| 43 | Figure 4B | Spatial | BM1' learner vs. BM3 learner | Day 6 | No assumption | Wilcoxon rank-sum test | z = 0.50 | 0.614 | Bonferroni (0.05 / 3 pairs) | r = 0.08 |
| 44 | Figure 4B | Spatial | BM1' learner vs. Beginner | Day 6 | No assumption | Wilcoxon rank-sum test | z = 3.27 | 0.001* | Bonferroni (0.05 / 3 pairs) | r = 0.46 |
| 45 | Figure 4B | Spatial | BM3 learner vs. Beginner | Day 6 | No assumption | Wilcoxon rank-sum test | z = 3.38 | 0.001* | Bonferroni (0.05 / 3 pairs) | r = 0.46 |

Note. Asterisks indicate statistically significant differences. N/A: not applicable. ES: effect size.
