## Supplementary material for "Scale space calibrates present and subsequent spatial learning in Barnes maze in mice": Statistical table 5

Statistical results of network analysis for the BM1’ learner, the BM3 learner, and the Beginner in the BM1 training.

| Manuscript reference # | Figure | Measure | Comparison | Within | Data structure | Type of test | Statistic | p | Correction | ES |
| --- | --- | --- | --- | --- | --- | --- | --- | --- | --- | --- |
| 1 | Figure 4C | Number of stops | BM1' learner, BM3 learner, Beginner | Day 1 | No assumption | Kruskall-Wallis test | X^2^ (2) = 8.02 | 0.018 | Bonferroni (0.05 / 6 days) | N/A |
| 2 | Figure 4C | Number of stops | BM1' learner, BM3 learner, Beginner | Day 2 | No assumption | Kruskall-Wallis test | X^2^ (2) = 6.70 | 0.035 | Bonferroni (0.05 / 6 days) | N/A |
| 3 | Figure 4C | Number of stops | BM1' learner, BM3 learner, Beginner | Day 3 | No assumption | Kruskall-Wallis test | X^2^ (2) = 8.26 | 0.016 | Bonferroni (0.05 / 6 days) | N/A |
| 4 | Figure 4C | Number of stops | BM1' learner, BM3 learner, Beginner | Day 4 | No assumption | Kruskall-Wallis test | X^2^ (2) = 10.68 | 0.005* | Bonferroni (0.05 / 6 days) | N/A |
| 5 | Figure 4C | Number of stops | BM1' learner vs. BM3 learner | Day 4 | No assumption | Wilcoxon rank-sum test | z = -1.63 | 0.102 | Bonferroni (0.05 / 3 pairs) | r = -0.27 |
| 6 | Figure 4C | Number of stops | BM1' learner vs. Beginner | Day 4 | No assumption | Wilcoxon rank-sum test | z = -2.99 | 0.003* | Bonferroni (0.05 / 3 pairs) | r = -0.42 |
| 7 | Figure 4C | Number of stops | BM3 learner vs. Beginner | Day 4 | No assumption | Wilcoxon rank-sum test | z = -1.92 | 0.055 | Bonferroni (0.05 / 3 pairs) | r = -0.26 |
| 8 | Figure 4C | Number of stops | BM1' learner, BM3 learner, Beginner | Day 5 | No assumption | Kruskall-Wallis test | X^2^ (2) = 6.86 | 0.032 | Bonferroni (0.05 / 6 days) | N/A |
| 9 | Figure 4C | Number of stops | BM1' learner, BM3 learner, Beginner | Day 6 | No assumption | Kruskall-Wallis test | X^2^ (2) = 4.07 | 0.131 | Bonferroni (0.05 / 6 days) | N/A |
| 10 | Figure 4C | Order | BM1' learner, BM3 learner, Beginner | Day 1 | No assumption | Kruskall-Wallis test | X^2^ (2) = 8.56 | 0.014 | Bonferroni (0.05 / 6 days) | N/A |
| 11 | Figure 4C | Order | BM1' learner, BM3 learner, Beginner | Day 2 | No assumption | Kruskall-Wallis test | X^2^ (2) = 13.21 | 0.001* | Bonferroni (0.05 / 6 days) | N/A |
| 12 | Figure 4C | Order | BM1' learner vs. BM3 learner | Day 2 | No assumption | Wilcoxon rank-sum test | z = -0.86 | 0.392 | Bonferroni (0.05 / 3 pairs) | r = -0.14 |
| 13 | Figure 4C | Order | BM1' learner vs. Beginner | Day 2 | No assumption | Wilcoxon rank-sum test | z = -2.99 | 0.003* | Bonferroni (0.05 / 3 pairs) | r = -0.42 |
| 14 | Figure 4C | Order | BM3 learner vs. Beginner | Day 2 | No assumption | Wilcoxon rank-sum test | z = -2.89 | 0.004* | Bonferroni (0.05 / 3 pairs) | r = -0.39 |
| 15 | Figure 4C | Order | BM1' learner, BM3 learner, Beginner | Day 3 | No assumption | Kruskall-Wallis test | X^2^ (2) = 21.66 | 0.000* | Bonferroni (0.05 / 6 days) | N/A |
| 16 | Figure 4C | Order | BM1' learner vs. BM3 learner | Day 3 | No assumption | Wilcoxon rank-sum test | z = -0.12 | 0.902 | Bonferroni (0.05 / 3 pairs) | r = -0.02 |
| 17 | Figure 4C | Order | BM1' learner vs. Beginner | Day 3 | No assumption | Wilcoxon rank-sum test | z = -3.37 | 0.001* | Bonferroni (0.05 / 3 pairs) | r = -0.47 |
| 18 | Figure 4C | Order | BM3 learner vs. Beginner | Day 3 | No assumption | Wilcoxon rank-sum test | z = -4.19 | 0.000* | Bonferroni (0.05 / 3 pairs) | r = -0.57 |
| 19 | Figure 4C | Order | BM1' learner, BM3 learner, Beginner | Day 4 | No assumption | Kruskall-Wallis test | X^2^ (2) = 27.74 | 0.000* | Bonferroni (0.05 / 6 days) | N/A |
| 20 | Figure 4C | Order | BM1' learner vs. BM3 learner | Day 4 | No assumption | Wilcoxon rank-sum test | z = -1.81 | 0.070 | Bonferroni (0.05 / 3 pairs) | r = -0.30 |
| 21 | Figure 4C | Order | BM1' learner vs. Beginner | Day 4 | No assumption | Wilcoxon rank-sum test | z = -4.54 | 0.000* | Bonferroni (0.05 / 3 pairs) | r = -0.64 |
| 22 | Figure 4C | Order | BM3 learner vs. Beginner | Day 4 | No assumption | Wilcoxon rank-sum test | z = -3.84 | 0.000* | Bonferroni (0.05 / 3 pairs) | r = -0.52 |
| 23 | Figure 4C | Order | BM1' learner, BM3 learner, Beginner | Day 5 | No assumption | Kruskall-Wallis test | X^2^ (2) = 21.83 | 0.000* | Bonferroni (0.05 / 6 days) | N/A |
| 24 | Figure 4C | Order | BM1' learner vs. BM3 learner | Day 5 | No assumption | Wilcoxon rank-sum test | z = -0.98 | 0.329 | Bonferroni (0.05 / 3 pairs) | r = -0.16 |
| 25 | Figure 4C | Order | BM1' learner vs. Beginner | Day 5 | No assumption | Wilcoxon rank-sum test | z = -3.90 | 0.000* | Bonferroni (0.05 / 3 pairs) | r = -0.55 |
| 26 | Figure 4C | Order | BM3 learner vs. Beginner | Day 5 | No assumption | Wilcoxon rank-sum test | z = -3.67 | 0.000* | Bonferroni (0.05 / 3 pairs) | r = -0.50 |
| 27 | Figure 4C | Order | BM1' learner, BM3 learner, Beginner | Day 6 | No assumption | Kruskall-Wallis test | X^2^ (2) = 13.86 | 0.001* | Bonferroni (0.05 / 6 days) | N/A |
| 28 | Figure 4C | Order | BM1' learner vs. BM3 learner | Day 6 | No assumption | Wilcoxon rank-sum test | z = -1.20 | 0.231 | Bonferroni (0.05 / 3 pairs) | r = -0.20 |
| 29 | Figure 4C | Order | BM1' learner vs. Beginner | Day 6 | No assumption | Wilcoxon rank-sum test | z = -3.29 | 0.001* | Bonferroni (0.05 / 3 pairs) | r = -0.46 |
| 30 | Figure 4C | Order | BM3 learner vs. Beginner | Day 6 | No assumption | Wilcoxon rank-sum test | z = -2.62 | 0.009* | Bonferroni (0.05 / 3 pairs) | r = -0.36 |
| 31 | Figure 4C | Degree | BM1' learner, BM3 learner, Beginner | Day 1 | No assumption | Kruskall-Wallis test | X^2^ (2) = 8.63 | 0.013 | Bonferroni (0.05 / 6 days) | N/A |
| 32 | Figure 4C | Degree | BM1' learner, BM3 learner, Beginner | Day 2 | No assumption | Kruskall-Wallis test | X^2^ (2) = 16.81 | 0.000* | Bonferroni (0.05 / 6 days) | N/A |
| 33 | Figure 4C | Degree | BM1' learner vs. BM3 learner | Day 2 | No assumption | Wilcoxon rank-sum test | z = -1.13 | 0.258 | Bonferroni (0.05 / 3 pairs) | r = -0.19 |
| 34 | Figure 4C | Degree | BM1' learner vs. Beginner | Day 2 | No assumption | Wilcoxon rank-sum test | z = -3.29 | 0.001* | Bonferroni (0.05 / 3 pairs) | r = -0.46 |
| 35 | Figure 4C | Degree | BM3 learner vs. Beginner | Day 2 | No assumption | Wilcoxon rank-sum test | z = -3.33 | 0.001* | Bonferroni (0.05 / 3 pairs) | r = -0.45 |
| 36 | Figure 4C | Degree | BM1' learner, BM3 learner, Beginner | Day 3 | No assumption | Kruskall-Wallis test | X^2^ (2) = 22.40 | 0.000* | Bonferroni (0.05 / 6 days) | N/A |
| 37 | Figure 4C | Degree | BM1' learner vs. BM3 learner | Day 3 | No assumption | Wilcoxon rank-sum test | z = -0.52 | 0.603 | Bonferroni (0.05 / 3 pairs) | r = -0.09 |
| 38 | Figure 4C | Degree | BM1' learner vs. Beginner | Day 3 | No assumption | Wilcoxon rank-sum test | z = -3.79 | 0.000* | Bonferroni (0.05 / 3 pairs) | r = -0.53 |
| 39 | Figure 4C | Degree | BM3 learner vs. Beginner | Day 3 | No assumption | Wilcoxon rank-sum test | z = -3.93 | 0.000* | Bonferroni (0.05 / 3 pairs) | r = -0.54 |
| 40 | Figure 4C | Degree | BM1' learner, BM3 learner, Beginner | Day 4 | No assumption | Kruskall-Wallis test | X^2^ (2) = 29.42 | 0.000* | Bonferroni (0.05 / 6 days) | N/A |
| 41 | Figure 4C | Degree | BM1' learner vs. BM3 learner | Day 4 | No assumption | Wilcoxon rank-sum test | z = -1.84 | 0.066 | Bonferroni (0.05 / 3 pairs) | r = -0.30 |
| 42 | Figure 4C | Degree | BM1' learner vs. Beginner | Day 4 | No assumption | Wilcoxon rank-sum test | z = -4.68 | 0.000* | Bonferroni (0.05 / 3 pairs) | r = -0.66 |
| 43 | Figure 4C | Degree | BM3 learner vs. Beginner | Day 4 | No assumption | Wilcoxon rank-sum test | z = -3.95 | 0.000* | Bonferroni (0.05 / 3 pairs) | r = -0.54 |
| 44 | Figure 4C | Degree | BM1' learner, BM3 learner, Beginner | Day 5 | No assumption | Kruskall-Wallis test | X^2^ (2) = 17.11 | 0.000* | Bonferroni (0.05 / 6 days) | N/A |
| 45 | Figure 4C | Degree | BM1' learner vs. BM3 learner | Day 5 | No assumption | Wilcoxon rank-sum test | z = -1.17 | 0.240 | Bonferroni (0.05 / 3 pairs) | r = -0.19 |
| 46 | Figure 4C | Degree | BM1' learner vs. Beginner | Day 5 | No assumption | Wilcoxon rank-sum test | z = -3.62 | 0.000* | Bonferroni (0.05 / 3 pairs) | r = -0.51 |
| 47 | Figure 4C | Degree | BM3 learner vs. Beginner | Day 5 | No assumption | Wilcoxon rank-sum test | z = -3.00 | 0.003* | Bonferroni (0.05 / 3 pairs) | r = -0.41 |
| 48 | Figure 4C | Degree | BM1' learner, BM3 learner, Beginner | Day 6 | No assumption | Kruskall-Wallis test | X^2^ (2) = 13.22 | 0.001* | Bonferroni (0.05 / 6 days) | N/A |
| 49 | Figure 4C | Degree | BM1' learner vs. BM3 learner | Day 6 | No assumption | Wilcoxon rank-sum test | z = -1.05 | 0.295 | Bonferroni (0.05 / 3 pairs) | r = -0.17 |
| 50 | Figure 4C | Degree | BM1' learner vs. Beginner | Day 6 | No assumption | Wilcoxon rank-sum test | z = -3.09 | 0.002* | Bonferroni (0.05 / 3 pairs) | r = -0.43 |
| 51 | Figure 4C | Degree | BM3 learner vs. Beginner | Day 6 | No assumption | Wilcoxon rank-sum test | z = -2.74 | 0.006* | Bonferroni (0.05 / 3 pairs) | r = -0.37 |
| 52 | Figure 4C | Density | BM1' learner, BM3 learner, Beginner | Day 1 | No assumption | Kruskall-Wallis test | X^2^ (2) = 3.45 | 0.178 | Bonferroni (0.05 / 6 days) | N/A |
| 53 | Figure 4C | Density | BM1' learner, BM3 learner, Beginner | Day 2 | No assumption | Kruskall-Wallis test | X^2^ (2) = 0.70 | 0.706 | Bonferroni (0.05 / 6 days) | N/A |
| 54 | Figure 4C | Density | BM1' learner, BM3 learner, Beginner | Day 3 | No assumption | Kruskall-Wallis test | X^2^ (2) = 2.58 | 0.275 | Bonferroni (0.05 / 6 days) | N/A |
| 55 | Figure 4C | Density | BM1' learner, BM3 learner, Beginner | Day 4 | No assumption | Kruskall-Wallis test | X^2^ (2) = 5.87 | 0.053 | Bonferroni (0.05 / 6 days) | N/A |
| 56 | Figure 4C | Density | BM1' learner, BM3 learner, Beginner | Day 5 | No assumption | Kruskall-Wallis test | X^2^ (2) = 5.09 | 0.079 | Bonferroni (0.05 / 6 days) | N/A |
| 57 | Figure 4C | Density | BM1' learner, BM3 learner, Beginner | Day 6 | No assumption | Kruskall-Wallis test | X^2^ (2) = 3.37 | 0.186 | Bonferroni (0.05 / 6 days) | N/A |
| 58 | Figure 4C | Clustering coefficient | BM1' learner, BM3 learner, Beginner | Day 1 | No assumption | Kruskall-Wallis test | X^2^ (2) = 9.31 | 0.009 | Bonferroni (0.05 / 6 days) | N/A |
| 59 | Figure 4C | Clustering coefficient | BM1' learner, BM3 learner, Beginner | Day 2 | No assumption | Kruskall-Wallis test | X^2^ (2) = 7.83 | 0.020 | Bonferroni (0.05 / 6 days) | N/A |
| 60 | Figure 4C | Clustering coefficient | BM1' learner, BM3 learner, Beginner | Day 3 | No assumption | Kruskall-Wallis test | X^2^ (2) = 7.37 | 0.025 | Bonferroni (0.05 / 6 days) | N/A |
| 61 | Figure 4C | Clustering coefficient | BM1' learner, BM3 learner, Beginner | Day 4 | No assumption | Kruskall-Wallis test | X^2^ (2) = 8.64 | 0.013 | Bonferroni (0.05 / 6 days) | N/A |
| 62 | Figure 4C | Clustering coefficient | BM1' learner, BM3 learner, Beginner | Day 5 | No assumption | Kruskall-Wallis test | X^2^ (2) = 6.74 | 0.034 | Bonferroni (0.05 / 6 days) | N/A |
| 63 | Figure 4C | Clustering coefficient | BM1' learner, BM3 learner, Beginner | Day 6 | No assumption | Kruskall-Wallis test | X^2^ (2) = 2.75 | 0.253 | Bonferroni (0.05 / 6 days) | N/A |
| 64 | Figure 4C | Shortest path length | BM1' learner, BM3 learner, Beginner | Day 1 | No assumption | Kruskall-Wallis test | X^2^ (2) = 2.11 | 0.347 | Bonferroni (0.05 / 6 days) | N/A |
| 65 | Figure 4C | Shortest path length | BM1' learner, BM3 learner, Beginner | Day 2 | No assumption | Kruskall-Wallis test | X^2^ (2) = 11.49 | 0.003* | Bonferroni (0.05 / 6 days) | N/A |
| 66 | Figure 4C | Shortest path length | BM1' learner vs. BM3 learner | Day 2 | No assumption | Wilcoxon rank-sum test | z = -1.13 | 0.259 | Bonferroni (0.05 / 3 pairs) | r = -0.19 |
| 67 | Figure 4C | Shortest path length | BM1' learner vs. Beginner | Day 2 | No assumption | Wilcoxon rank-sum test | z = -2.92 | 0.004* | Bonferroni (0.05 / 3 pairs) | r = -0.41 |
| 68 | Figure 4C | Shortest path length | BM3 learner vs. Beginner | Day 2 | No assumption | Wilcoxon rank-sum test | z = -2.47 | 0.013* | Bonferroni (0.05 / 3 pairs) | r = -0.34 |
| 69 | Figure 4C | Shortest path length | BM1' learner, BM3 learner, Beginner | Day 3 | No assumption | Kruskall-Wallis test | X^2^ (2) = 19.10 | 0.000* | Bonferroni (0.05 / 6 days) | N/A |
| 70 | Figure 4C | Shortest path length | BM1' learner vs. BM3 learner | Day 3 | No assumption | Wilcoxon rank-sum test | z = -0.41 | 0.680 | Bonferroni (0.05 / 3 pairs) | r = -0.07 |
| 71 | Figure 4C | Shortest path length | BM1' learner vs. Beginner | Day 3 | No assumption | Wilcoxon rank-sum test | z = -3.37 | 0.001* | Bonferroni (0.05 / 3 pairs) | r = -0.47 |
| 72 | Figure 4C | Shortest path length | BM3 learner vs. Beginner | Day 3 | No assumption | Wilcoxon rank-sum test | z = -3.76 | 0.000* | Bonferroni (0.05 / 3 pairs) | r = -0.51 |
| 73 | Figure 4C | Shortest path length | BM1' learner, BM3 learner, Beginner | Day 4 | No assumption | Kruskall-Wallis test | X^2^ (2) = 25.76 | 0.000* | Bonferroni (0.05 / 6 days) | N/A |
| 74 | Figure 4C | Shortest path length | BM1' learner vs. BM3 learner | Day 4 | No assumption | Wilcoxon rank-sum test | z = -1.86 | 0.063 | Bonferroni (0.05 / 3 pairs) | r = -0.31 |
| 75 | Figure 4C | Shortest path length | BM1' learner vs. Beginner | Day 4 | No assumption | Wilcoxon rank-sum test | z = -4.34 | 0.000* | Bonferroni (0.05 / 3 pairs) | r = -0.61 |
| 76 | Figure 4C | Shortest path length | BM3 learner vs. Beginner | Day 4 | No assumption | Wilcoxon rank-sum test | z = -3.71 | 0.000* | Bonferroni (0.05 / 3 pairs) | r = -0.50 |
| 77 | Figure 4C | Shortest path length | BM1' learner, BM3 learner, Beginner | Day 5 | No assumption | Kruskall-Wallis test | X^2^ (2) = 20.32 | 0.000* | Bonferroni (0.05 / 6 days) | N/A |
| 78 | Figure 4C | Shortest path length | BM1' learner vs. BM3 learner | Day 5 | No assumption | Wilcoxon rank-sum test | z = -1.25 | 0.211 | Bonferroni (0.05 / 3 pairs) | r = -0.21 |
| 79 | Figure 4C | Shortest path length | BM1' learner vs. Beginner | Day 5 | No assumption | Wilcoxon rank-sum test | z = -3.92 | 0.000* | Bonferroni (0.05 / 3 pairs) | r = -0.55 |
| 80 | Figure 4C | Shortest path length | BM3 learner vs. Beginner | Day 5 | No assumption | Wilcoxon rank-sum test | z = -3.30 | 0.001* | Bonferroni (0.05 / 3 pairs) | r = -0.45 |
| 81 | Figure 4C | Shortest path length | BM1' learner, BM3 learner, Beginner | Day 6 | No assumption | Kruskall-Wallis test | X^2^ (2) = 12.53 | 0.002* | Bonferroni (0.05 / 6 days) | N/A |
| 82 | Figure 4C | Shortest path length | BM1' learner vs. BM3 learner | Day 6 | No assumption | Wilcoxon rank-sum test | z = -1.06 | 0.289 | Bonferroni (0.05 / 3 pairs) | r = -0.17 |
| 83 | Figure 4C | Shortest path length | BM1' learner vs. Beginner | Day 6 | No assumption | Wilcoxon rank-sum test | z = -3.14 | 0.002* | Bonferroni (0.05 / 3 pairs) | r = -0.44 |
| 84 | Figure 4C | Shortest path length | BM3 learner vs. Beginner | Day 6 | No assumption | Wilcoxon rank-sum test | z = -2.48 | 0.013* | Bonferroni (0.05 / 3 pairs) | r = -0.34 |
| 85 | Figure 4C | Betweenness centrality | BM1' learner, BM3 learner, Beginner | Day 1 | No assumption | Kruskall-Wallis test | X^2^ (2) = 1.37 | 0.504 | Bonferroni (0.05 / 6 days) | N/A |
| 86 | Figure 4C | Betweenness centrality | BM1' learner, BM3 learner, Beginner | Day 2 | No assumption | Kruskall-Wallis test | X^2^ (2) = 6.80 | 0.033 | Bonferroni (0.05 / 6 days) | N/A |
| 87 | Figure 4C | Betweenness centrality | BM1' learner, BM3 learner, Beginner | Day 3 | No assumption | Kruskall-Wallis test | X^2^ (2) = 16.61 | 0.000* | Bonferroni (0.05 / 6 days) | N/A |
| 88 | Figure 4C | Betweenness centrality | BM1' learner vs. BM3 learner | Day 3 | No assumption | Wilcoxon rank-sum test | z = 0.00 | 1.000 | Bonferroni (0.05 / 3 pairs) | r = 0.00 |
| 89 | Figure 4C | Betweenness centrality | BM1' learner vs. Beginner | Day 3 | No assumption | Wilcoxon rank-sum test | z = -3.02 | 0.002* | Bonferroni (0.05 / 3 pairs) | r = -0.42 |
| 90 | Figure 4C | Betweenness centrality | BM3 learner vs. Beginner | Day 3 | No assumption | Wilcoxon rank-sum test | z = -3.59 | 0.000* | Bonferroni (0.05 / 3 pairs) | r = -0.49 |
| 91 | Figure 4C | Betweenness centrality | BM1' learner, BM3 learner, Beginner | Day 4 | No assumption | Kruskall-Wallis test | X^2^ (2) = 19.58 | 0.000* | Bonferroni (0.05 / 6 days) | N/A |
| 92 | Figure 4C | Betweenness centrality | BM1' learner vs. BM3 learner | Day 4 | No assumption | Wilcoxon rank-sum test | z = -1.47 | 0.140 | Bonferroni (0.05 / 3 pairs) | r = -0.24 |
| 93 | Figure 4C | Betweenness centrality | BM1' learner vs. Beginner | Day 4 | No assumption | Wilcoxon rank-sum test | z = -4.03 | 0.000* | Bonferroni (0.05 / 3 pairs) | r = -0.56 |
| 94 | Figure 4C | Betweenness centrality | BM3 learner vs. Beginner | Day 4 | No assumption | Wilcoxon rank-sum test | z = -2.93 | 0.003* | Bonferroni (0.05 / 3 pairs) | r = -0.40 |
| 95 | Figure 4C | Betweenness centrality | BM1' learner, BM3 learner, Beginner | Day 5 | No assumption | Kruskall-Wallis test | X^2^ (2) = 21.48 | 0.000* | Bonferroni (0.05 / 6 days) | N/A |
| 96 | Figure 4C | Betweenness centrality | BM1' learner vs. BM3 learner | Day 5 | No assumption | Wilcoxon rank-sum test | z = -0.65 | 0.517 | Bonferroni (0.05 / 3 pairs) | r = -0.11 |
| 97 | Figure 4C | Betweenness centrality | BM1' learner vs. Beginner | Day 5 | No assumption | Wilcoxon rank-sum test | z = -4.01 | 0.000* | Bonferroni (0.05 / 3 pairs) | r = -0.56 |
| 98 | Figure 4C | Betweenness centrality | BM3 learner vs. Beginner | Day 5 | No assumption | Wilcoxon rank-sum test | z = -3.46 | 0.001* | Bonferroni (0.05 / 3 pairs) | r = -0.47 |
| 99 | Figure 4C | Betweenness centrality | BM1' learner, BM3 learner, Beginner | Day 6 | No assumption | Kruskall-Wallis test | X^2^ (2) = 9.98 | 0.007* | Bonferroni (0.05 / 6 days) | N/A |
| 100 | Figure 4C | Betweenness centrality | BM1' learner vs. BM3 learner | Day 6 | No assumption | Wilcoxon rank-sum test | z = -0.91 | 0.362 | Bonferroni (0.05 / 3 pairs) | r = -0.15 |
| 101 | Figure 4C | Betweenness centrality | BM1' learner vs. Beginner | Day 6 | No assumption | Wilcoxon rank-sum test | z = -2.81 | 0.005* | Bonferroni (0.05 / 3 pairs) | r = -0.39 |
| 102 | Figure 4C | Betweenness centrality | BM3 learner vs. Beginner | Day 6 | No assumption | Wilcoxon rank-sum test | z = -2.18 | 0.029 | Bonferroni (0.05 / 3 pairs) | r = -0.30 |
| 103 | Figure 4C | Closeness centrality | BM1' learner, BM3 learner, Beginner | Day 1 | No assumption | Kruskall-Wallis test | X^2^ (2) = 3.06 | 0.217 | Bonferroni (0.05 / 6 days) | N/A |
| 104 | Figure 4C | Closeness centrality | BM1' learner, BM3 learner, Beginner | Day 2 | No assumption | Kruskall-Wallis test | X^2^ (2) = 0.42 | 0.811 | Bonferroni (0.05 / 6 days) | N/A |
| 105 | Figure 4C | Closeness centrality | BM1' learner, BM3 learner, Beginner | Day 3 | No assumption | Kruskall-Wallis test | X^2^ (2) = 2.44 | 0.296 | Bonferroni (0.05 / 6 days) | N/A |
| 106 | Figure 4C | Closeness centrality | BM1' learner, BM3 learner, Beginner | Day 4 | No assumption | Kruskall-Wallis test | X^2^ (2) = 3.43 | 0.180 | Bonferroni (0.05 / 6 days) | N/A |
| 107 | Figure 4C | Closeness centrality | BM1' learner, BM3 learner, Beginner | Day 5 | No assumption | Kruskall-Wallis test | X^2^ (2) = 4.73 | 0.094 | Bonferroni (0.05 / 6 days) | N/A |
| 108 | Figure 4C | Closeness centrality | BM1' learner, BM3 learner, Beginner | Day 6 | No assumption | Kruskall-Wallis test | X^2^ (2) = 2.36 | 0.308 | Bonferroni (0.05 / 6 days) | N/A |

Note. Asterisks indicate statistically significant differences. N/A: not applicable. ES: effect size.
