## Supplementary material for "Scale space calibrates present and subsequent spatial learning in Barnes maze in mice": Statistical table 6

Statistical results of network analysis for the BM1’ learner, the BM3 learner, and the Beginner in the BM1 probe test.

| Manuscript reference # | Figure | Measure | Comparison | Within | Data structure | Type of test | Statistic | p | Correction | ES |
| --- | --- | --- | --- | --- | --- | --- | --- | --- | --- | --- |
| 1 | Figure 4E | Number of stops | BM1' learner, BM3 learner, Beginner | Probe test | No assumption | Kruskall-Wallis test | X^2^ (2) = 5.61 | 0.06 | N/A | N/A |
| 2 | Figure 4E | Order | BM1' learner, BM3 learner, Beginner | Probe test | No assumption | Kruskall-Wallis test | X^2^ (2) = 12.99 | 0.00* | N/A | N/A |
| 3 | Figure 4E | Order | BM1' learner vs. BM3 learner | Probe test | No assumption | Wilcoxon rank-sum test | z = -0.11 | 0.91 | Bonferroni (0.05 / 3 instances) | r = -0.02 |
| 4 | Figure 4E | Order | BM1' learner vs. Beginner | Probe test | No assumption | Wilcoxon rank-sum test | z = -3.23 | 0.00* | Bonferroni (0.05 / 3 instances) | r = -0.45 |
| 5 | Figure 4E | Order | BM3 learner vs. Beginner | Probe test | No assumption | Wilcoxon rank-sum test | z = -2.67 | 0.01* | Bonferroni (0.05 / 3 instances) | r = -0.36 |
| 6 | Figure 4E | Degree | BM1' learner, BM3 learner, Beginner | Probe test | No assumption | Kruskall-Wallis test | X^2^ (2) = 7.17 | 0.03* | N/A | N/A |
| 7 | Figure 4E | Degree | BM1' learner vs. BM3 learner | Probe test | No assumption | Wilcoxon rank-sum test | z = 0.63 | 0.53 | Bonferroni (0.05 / 3 instances) | r = 0.10 |
| 8 | Figure 4E | Degree | BM1' learner vs. Beginner | Probe test | No assumption | Wilcoxon rank-sum test | z = -1.70 | 0.09 | Bonferroni (0.05 / 3 instances) | r = -0.24 |
| 9 | Figure 4E | Degree | BM3 learner vs. Beginner | Probe test | No assumption | Wilcoxon rank-sum test | z = -2.51 | 0.01* | Bonferroni (0.05 / 3 instances) | r = -0.34 |
| 10 | Figure 4E | Density | BM1' learner, BM3 learner, Beginner | Probe test | No assumption | Kruskall-Wallis test | X^2^ (2) = 7.24 | 0.03* | N/A | N/A |
| 11 | Figure 4E | Density | BM1' learner vs. BM3 learner | Probe test | No assumption | Wilcoxon rank-sum test | z = 0.11 | 0.92 | Bonferroni (0.05 / 3 instances) | r = 0.02 |
| 12 | Figure 4E | Density | BM1' learner vs. Beginner | Probe test | No assumption | Wilcoxon rank-sum test | z = 2.50 | 0.01* | Bonferroni (0.05 / 3 instances) | r = 0.35 |
| 13 | Figure 4E | Density | BM3 learner vs. Beginner | Probe test | No assumption | Wilcoxon rank-sum test | z = 1.89 | 0.06 | Bonferroni (0.05 / 3 instances) | r = 0.26 |
| 14 | Figure 4E | Clustering coefficient | BM1' learner, BM3 learner, Beginner | Probe test | No assumption | Kruskall-Wallis test | X^2^ (2) = 0.57 | 0.75 | N/A | N/A |
| 15 | Figure 4E | Shortest path length | BM1' learner, BM3 learner, Beginner | Probe test | No assumption | Kruskall-Wallis test | X^2^ (2) = 4.06 | 0.13 | N/A | N/A |
| 16 | Figure 4E | Betweenness centrality | BM1' learner, BM3 learner, Beginner | Probe test | No assumption | Kruskall-Wallis test | X^2^ (2) = 3.04 | 0.22 | N/A | N/A |
| 17 | Figure 4E | Closeness centrality | BM1' learner, BM3 learner, Beginner | Probe test | No assumption | Kruskall-Wallis test | X^2^ (2) = 4.69 | 0.10 | N/A | N/A |

Note. Asterisks indicate statistically significant differences. N/A: not applicable. ES: effect size.
