## Supplementary material for "Scale space calibrates present and subsequent spatial learning in Barnes maze in mice": Statistical table 7

Statistical results of strategy analysis for the BM1 learner and the Beginner in the BM3 training phase.

| Manuscript reference # | Figure | Measure | Comparison | Within | Data structure | Type of test | Statistic | p | Correction | Effect size |
| --- | --- | --- | --- | --- | --- | --- | --- | --- | --- | --- |
| 1 | Figure 5B | Random | BM1 learner vs. Beginner | Day 1 | No assumption | Wilcoxon rank-sum test | z = -1.79 | 0.073 | Bonferroni (0.05 / 12 days) | r = -0.22 |
| 2 | Figure 5B | Random | BM1 learner vs. Beginner | Day 2 | No assumption | Wilcoxon rank-sum test | z = -0.30 | 0.768 | Bonferroni (0.05 / 12 days) | r = -0.04 |
| 3 | Figure 5B | Random | BM1 learner vs. Beginner | Day 3 | No assumption | Wilcoxon rank-sum test | z = -0.98 | 0.329 | Bonferroni (0.05 / 12 days) | r = -0.12 |
| 4 | Figure 5B | Random | BM1 learner vs. Beginner | Day 4 | No assumption | Wilcoxon rank-sum test | z = -3.21 | 0.001* | Bonferroni (0.05 / 12 days) | r = -0.39 |
| 5 | Figure 5B | Random | BM1 learner vs. Beginner | Day 5 | No assumption | Wilcoxon rank-sum test | z = -2.08 | 0.038 | Bonferroni (0.05 / 12 days) | r = -0.25 |
| 6 | Figure 5B | Random | BM1 learner vs. Beginner | Day 6 | No assumption | Wilcoxon rank-sum test | z = -0.09 | 0.927 | Bonferroni (0.05 / 12 days) | r = -0.01 |
| 7 | Figure 5B | Random | BM1 learner vs. Beginner | Day 7 | No assumption | Wilcoxon rank-sum test | z = -1.09 | 0.274 | Bonferroni (0.05 / 12 days) | r = -0.13 |
| 8 | Figure 5B | Random | BM1 learner vs. Beginner | Day 8 | No assumption | Wilcoxon rank-sum test | z = -0.28 | 0.778 | Bonferroni (0.05 / 12 days) | r = -0.03 |
| 9 | Figure 5B | Random | BM1 learner vs. Beginner | Day 9 | No assumption | Wilcoxon rank-sum test | z = 0.33 | 0.739 | Bonferroni (0.05 / 12 days) | r = 0.04 |
| 10 | Figure 5B | Random | BM1 learner vs. Beginner | Day 10 | No assumption | Wilcoxon rank-sum test | z = 0.84 | 0.398 | Bonferroni (0.05 / 12 days) | r = 0.10 |
| 11 | Figure 5B | Random | BM1 learner vs. Beginner | Day 11 | No assumption | Wilcoxon rank-sum test | z = -1.50 | 0.135 | Bonferroni (0.05 / 12 days) | r = -0.18 |
| 12 | Figure 5B | Random | BM1 learner vs. Beginner | Day 12 | No assumption | Wilcoxon rank-sum test | z = 0.54 | 0.591 | Bonferroni (0.05 / 12 days) | r = 0.06 |
| 13 | Figure 5B | Serial | BM1 learner vs. Beginner | Day 1 | No assumption | Wilcoxon rank-sum test | z = 1.05 | 0.296 | Bonferroni (0.05 / 12 days) | r = 0.13 |
| 14 | Figure 5B | Serial | BM1 learner vs. Beginner | Day 2 | No assumption | Wilcoxon rank-sum test | z = 0.00 | 1.000 | Bonferroni (0.05 / 12 days) | r = 0.00 |
| 15 | Figure 5B | Serial | BM1 learner vs. Beginner | Day 3 | No assumption | Wilcoxon rank-sum test | z = 0.91 | 0.365 | Bonferroni (0.05 / 12 days) | r = 0.11 |
| 16 | Figure 5B | Serial | BM1 learner vs. Beginner | Day 4 | No assumption | Wilcoxon rank-sum test | z = 2.69 | 0.007 | Bonferroni (0.05 / 12 days) | r = 0.32 |
| 17 | Figure 5B | Serial | BM1 learner vs. Beginner | Day 5 | No assumption | Wilcoxon rank-sum test | z = 1.54 | 0.122 | Bonferroni (0.05 / 12 days) | r = 0.19 |
| 18 | Figure 5B | Serial | BM1 learner vs. Beginner | Day 6 | No assumption | Wilcoxon rank-sum test | z = 0.54 | 0.587 | Bonferroni (0.05 / 12 days) | r = 0.07 |
| 19 | Figure 5B | Serial | BM1 learner vs. Beginner | Day 7 | No assumption | Wilcoxon rank-sum test | z = 1.42 | 0.154 | Bonferroni (0.05 / 12 days) | r = 0.17 |
| 20 | Figure 5B | Serial | BM1 learner vs. Beginner | Day 8 | No assumption | Wilcoxon rank-sum test | z = 0.24 | 0.807 | Bonferroni (0.05 / 12 days) | r = 0.03 |
| 21 | Figure 5B | Serial | BM1 learner vs. Beginner | Day 9 | No assumption | Wilcoxon rank-sum test | z = -0.32 | 0.750 | Bonferroni (0.05 / 12 days) | r = -0.04 |
| 22 | Figure 5B | Serial | BM1 learner vs. Beginner | Day 10 | No assumption | Wilcoxon rank-sum test | z = -0.71 | 0.479 | Bonferroni (0.05 / 12 days) | r = -0.09 |
| 23 | Figure 5B | Serial | BM1 learner vs. Beginner | Day 11 | No assumption | Wilcoxon rank-sum test | z = 1.56 | 0.118 | Bonferroni (0.05 / 12 days) | r = 0.19 |
| 24 | Figure 5B | Serial | BM1 learner vs. Beginner | Day 12 | No assumption | Wilcoxon rank-sum test | z = -0.16 | 0.875 | Bonferroni (0.05 / 12 days) | r = -0.02 |
| 25 | Figure 5B | Spatial | BM1 learner vs. Beginner | Day 1 | No assumption | Wilcoxon rank-sum test | z = 1.61 | 0.107 | Bonferroni (0.05 / 12 days) | r = 0.19 |
| 26 | Figure 5B | Spatial | BM1 learner vs. Beginner | Day 2 | No assumption | Wilcoxon rank-sum test | z = 0.26 | 0.793 | Bonferroni (0.05 / 12 days) | r = 0.03 |
| 27 | Figure 5B | Spatial | BM1 learner vs. Beginner | Day 3 | No assumption | Wilcoxon rank-sum test | z = 0.05 | 0.959 | Bonferroni (0.05 / 12 days) | r = 0.01 |
| 28 | Figure 5B | Spatial | BM1 learner vs. Beginner | Day 4 | No assumption | Wilcoxon rank-sum test | z = 1.92 | 0.055 | Bonferroni (0.05 / 12 days) | r = 0.23 |
| 29 | Figure 5B | Spatial | BM1 learner vs. Beginner | Day 5 | No assumption | Wilcoxon rank-sum test | z = 1.17 | 0.242 | Bonferroni (0.05 / 12 days) | r = 0.14 |
| 30 | Figure 5B | Spatial | BM1 learner vs. Beginner | Day 6 | No assumption | Wilcoxon rank-sum test | z = -0.38 | 0.701 | Bonferroni (0.05 / 12 days) | r = -0.05 |
| 31 | Figure 5B | Spatial | BM1 learner vs. Beginner | Day 7 | No assumption | Wilcoxon rank-sum test | z = -0.13 | 0.894 | Bonferroni (0.05 / 12 days) | r = -0.02 |
| 32 | Figure 5B | Spatial | BM1 learner vs. Beginner | Day 8 | No assumption | Wilcoxon rank-sum test | z = -0.05 | 0.961 | Bonferroni (0.05 / 12 days) | r = -0.01 |
| 33 | Figure 5B | Spatial | BM1 learner vs. Beginner | Day 9 | No assumption | Wilcoxon rank-sum test | z = -0.06 | 0.950 | Bonferroni (0.05 / 12 days) | r = -0.01 |
| 34 | Figure 5B | Spatial | BM1 learner vs. Beginner | Day 10 | No assumption | Wilcoxon rank-sum test | z = 0.12 | 0.906 | Bonferroni (0.05 / 12 days) | r = 0.01 |
| 35 | Figure 5B | Spatial | BM1 learner vs. Beginner | Day 11 | No assumption | Wilcoxon rank-sum test | z = 0.26 | 0.792 | Bonferroni (0.05 / 12 days) | r = 0.03 |
| 36 | Figure 5B | Spatial | BM1 learner vs. Beginner | Day 12 | No assumption | Wilcoxon rank-sum test | z = -0.65 | 0.514 | Bonferroni (0.05 / 12 days) | r = -0.08 |

Note. Asterisks indicate statistically significant differences. ES: effect size
