## Supplementary material for "Scale space calibrates present and subsequent spatial learning in Barnes maze in mice": Statistical table 8

Statistical results of network analysis for the BM1 learner and the Beginner in the BM3 training phase.

| Manuscript reference # | Figure | Measure | Comparison | Within | Data structure | Type of test | Statistic | p | Correction | ES |
| --- | --- | --- | --- | --- | --- | --- | --- | --- | --- | --- |
| 1 | Figure 5C | Number of stops | BM1 learner vs. Beginner | Day 1 | No assumption | Kruskall-Wallis test | z = -2.88 | 0.004* | Bonferroni (0.05 / 12 days) | r = -0.35 |
| 2 | Figure 5C | Number of stops | BM1 learner vs. Beginner | Day 2 | No assumption | Kruskall-Wallis test | z = -2.80 | 0.005 | Bonferroni (0.05 / 12 days) | r = -0.34 |
| 3 | Figure 5C | Number of stops | BM1 learner vs. Beginner | Day 3 | No assumption | Kruskall-Wallis test | z = -3.35 | 0.001* | Bonferroni (0.05 / 12 days) | r = -0.40 |
| 4 | Figure 5C | Number of stops | BM1 learner vs. Beginner | Day 4 | No assumption | Kruskall-Wallis test | z = -4.49 | 0.000* | Bonferroni (0.05 / 12 days) | r = -0.54 |
| 5 | Figure 5C | Number of stops | BM1 learner vs. Beginner | Day 5 | No assumption | Kruskall-Wallis test | z = -2.35 | 0.019 | Bonferroni (0.05 / 12 days) | r = -0.28 |
| 6 | Figure 5C | Number of stops | BM1 learner vs. Beginner | Day 6 | No assumption | Kruskall-Wallis test | z = -2.22 | 0.027 | Bonferroni (0.05 / 12 days) | r = -0.27 |
| 7 | Figure 5C | Number of stops | BM1 learner vs. Beginner | Day 7 | No assumption | Kruskall-Wallis test | z = -2.93 | 0.003* | Bonferroni (0.05 / 12 days) | r = -0.35 |
| 8 | Figure 5C | Number of stops | BM1 learner vs. Beginner | Day 8 | No assumption | Kruskall-Wallis test | z = -3.02 | 0.003* | Bonferroni (0.05 / 12 days) | r = -0.36 |
| 9 | Figure 5C | Number of stops | BM1 learner vs. Beginner | Day 9 | No assumption | Kruskall-Wallis test | z = -1.81 | 0.070 | Bonferroni (0.05 / 12 days) | r = -0.22 |
| 10 | Figure 5C | Number of stops | BM1 learner vs. Beginner | Day 10 | No assumption | Kruskall-Wallis test | z = -2.57 | 0.010 | Bonferroni (0.05 / 12 days) | r = -0.31 |
| 11 | Figure 5C | Number of stops | BM1 learner vs. Beginner | Day 11 | No assumption | Kruskall-Wallis test | z = -3.51 | 0.000* | Bonferroni (0.05 / 12 days) | r = -0.42 |
| 12 | Figure 5C | Number of stops | BM1 learner vs. Beginner | Day 12 | No assumption | Kruskall-Wallis test | z = -2.18 | 0.029 | Bonferroni (0.05 / 12 days) | r = -0.26 |
| 13 | Figure 5C | Order | BM1 learner vs. Beginner | Day 1 | No assumption | Kruskall-Wallis test | z = -2.09 | 0.037 | Bonferroni (0.05 / 12 days) | r = -0.25 |
| 14 | Figure 5C | Order | BM1 learner vs. Beginner | Day 2 | No assumption | Kruskall-Wallis test | z = -1.92 | 0.055 | Bonferroni (0.05 / 12 days) | r = -0.23 |
| 15 | Figure 5C | Order | BM1 learner vs. Beginner | Day 3 | No assumption | Kruskall-Wallis test | z = -3.20 | 0.001* | Bonferroni (0.05 / 12 days) | r = -0.39 |
| 16 | Figure 5C | Order | BM1 learner vs. Beginner | Day 4 | No assumption | Kruskall-Wallis test | z = -3.79 | 0.000* | Bonferroni (0.05 / 12 days) | r = -0.46 |
| 17 | Figure 5C | Order | BM1 learner vs. Beginner | Day 5 | No assumption | Kruskall-Wallis test | z = -1.77 | 0.077 | Bonferroni (0.05 / 12 days) | r = -0.21 |
| 18 | Figure 5C | Order | BM1 learner vs. Beginner | Day 6 | No assumption | Kruskall-Wallis test | z = -1.13 | 0.259 | Bonferroni (0.05 / 12 days) | r = -0.14 |
| 19 | Figure 5C | Order | BM1 learner vs. Beginner | Day 7 | No assumption | Kruskall-Wallis test | z = -1.58 | 0.114 | Bonferroni (0.05 / 12 days) | r = -0.19 |
| 20 | Figure 5C | Order | BM1 learner vs. Beginner | Day 8 | No assumption | Kruskall-Wallis test | z = -1.98 | 0.047 | Bonferroni (0.05 / 12 days) | r = -0.24 |
| 21 | Figure 5C | Order | BM1 learner vs. Beginner | Day 9 | No assumption | Kruskall-Wallis test | z = -0.32 | 0.747 | Bonferroni (0.05 / 12 days) | r = -0.04 |
| 22 | Figure 5C | Order | BM1 learner vs. Beginner | Day 10 | No assumption | Kruskall-Wallis test | z = -1.13 | 0.259 | Bonferroni (0.05 / 12 days) | r = -0.14 |
| 23 | Figure 5C | Order | BM1 learner vs. Beginner | Day 11 | No assumption | Kruskall-Wallis test | z = -2.13 | 0.033 | Bonferroni (0.05 / 12 days) | r = -0.26 |
| 24 | Figure 5C | Order | BM1 learner vs. Beginner | Day 12 | No assumption | Kruskall-Wallis test | z = -0.14 | 0.890 | Bonferroni (0.05 / 12 days) | r = -0.02 |
| 25 | Figure 5C | Degree | BM1 learner vs. Beginner | Day 1 | No assumption | Kruskall-Wallis test | z = -2.28 | 0.023 | Bonferroni (0.05 / 12 days) | r = -0.27 |
| 26 | Figure 5C | Degree | BM1 learner vs. Beginner | Day 2 | No assumption | Kruskall-Wallis test | z = -2.22 | 0.027 | Bonferroni (0.05 / 12 days) | r = -0.27 |
| 27 | Figure 5C | Degree | BM1 learner vs. Beginner | Day 3 | No assumption | Kruskall-Wallis test | z = -3.74 | 0.000* | Bonferroni (0.05 / 12 days) | r = -0.45 |
| 28 | Figure 5C | Degree | BM1 learner vs. Beginner | Day 4 | No assumption | Kruskall-Wallis test | z = -3.47 | 0.001* | Bonferroni (0.05 / 12 days) | r = -0.42 |
| 29 | Figure 5C | Degree | BM1 learner vs. Beginner | Day 5 | No assumption | Kruskall-Wallis test | z = -2.40 | 0.016 | Bonferroni (0.05 / 12 days) | r = -0.29 |
| 30 | Figure 5C | Degree | BM1 learner vs. Beginner | Day 6 | No assumption | Kruskall-Wallis test | z = -1.42 | 0.156 | Bonferroni (0.05 / 12 days) | r = -0.17 |
| 31 | Figure 5C | Degree | BM1 learner vs. Beginner | Day 7 | No assumption | Kruskall-Wallis test | z = -1.86 | 0.063 | Bonferroni (0.05 / 12 days) | r = -0.22 |
| 32 | Figure 5C | Degree | BM1 learner vs. Beginner | Day 8 | No assumption | Kruskall-Wallis test | z = -1.65 | 0.099 | Bonferroni (0.05 / 12 days) | r = -0.20 |
| 33 | Figure 5C | Degree | BM1 learner vs. Beginner | Day 9 | No assumption | Kruskall-Wallis test | z = -1.01 | 0.315 | Bonferroni (0.05 / 12 days) | r = -0.12 |
| 34 | Figure 5C | Degree | BM1 learner vs. Beginner | Day 10 | No assumption | Kruskall-Wallis test | z = -1.19 | 0.234 | Bonferroni (0.05 / 12 days) | r = -0.14 |
| 35 | Figure 5C | Degree | BM1 learner vs. Beginner | Day 11 | No assumption | Kruskall-Wallis test | z = -2.24 | 0.025 | Bonferroni (0.05 / 12 days) | r = -0.27 |
| 36 | Figure 5C | Degree | BM1 learner vs. Beginner | Day 12 | No assumption | Kruskall-Wallis test | z = -0.91 | 0.365 | Bonferroni (0.05 / 12 days) | r = -0.11 |
| 37 | Figure 5C | Density | BM1 learner vs. Beginner | Day 1 | No assumption | Kruskall-Wallis test | z = 1.39 | 0.165 | Bonferroni (0.05 / 12 days) | r = 0.17 |
| 38 | Figure 5C | Density | BM1 learner vs. Beginner | Day 2 | No assumption | Kruskall-Wallis test | z = 0.53 | 0.597 | Bonferroni (0.05 / 12 days) | r = 0.06 |
| 39 | Figure 5C | Density | BM1 learner vs. Beginner | Day 3 | No assumption | Kruskall-Wallis test | z = 1.40 | 0.160 | Bonferroni (0.05 / 12 days) | r = 0.17 |
| 40 | Figure 5C | Density | BM1 learner vs. Beginner | Day 4 | No assumption | Kruskall-Wallis test | z = -0.28 | 0.776 | Bonferroni (0.05 / 12 days) | r = -0.03 |
| 41 | Figure 5C | Density | BM1 learner vs. Beginner | Day 5 | No assumption | Kruskall-Wallis test | z = 1.63 | 0.102 | Bonferroni (0.05 / 12 days) | r = 0.20 |
| 42 | Figure 5C | Density | BM1 learner vs. Beginner | Day 6 | No assumption | Kruskall-Wallis test | z = -0.70 | 0.485 | Bonferroni (0.05 / 12 days) | r = -0.08 |
| 43 | Figure 5C | Density | BM1 learner vs. Beginner | Day 7 | No assumption | Kruskall-Wallis test | z = 0.75 | 0.452 | Bonferroni (0.05 / 12 days) | r = 0.09 |
| 44 | Figure 5C | Density | BM1 learner vs. Beginner | Day 8 | No assumption | Kruskall-Wallis test | z = -0.65 | 0.514 | Bonferroni (0.05 / 12 days) | r = -0.08 |
| 45 | Figure 5C | Density | BM1 learner vs. Beginner | Day 9 | No assumption | Kruskall-Wallis test | z = -1.10 | 0.272 | Bonferroni (0.05 / 12 days) | r = -0.13 |
| 46 | Figure 5C | Density | BM1 learner vs. Beginner | Day 10 | No assumption | Kruskall-Wallis test | z = -1.17 | 0.240 | Bonferroni (0.05 / 12 days) | r = -0.14 |
| 47 | Figure 5C | Density | BM1 learner vs. Beginner | Day 11 | No assumption | Kruskall-Wallis test | z = 0.20 | 0.842 | Bonferroni (0.05 / 12 days) | r = 0.02 |
| 48 | Figure 5C | Density | BM1 learner vs. Beginner | Day 12 | No assumption | Kruskall-Wallis test | z = -1.64 | 0.101 | Bonferroni (0.05 / 12 days) | r = -0.20 |
| 49 | Figure 5C | Clustering coefficient | BM1 learner vs. Beginner | Day 1 | No assumption | Kruskall-Wallis test | z = -1.46 | 0.144 | Bonferroni (0.05 / 12 days) | r = -0.18 |
| 50 | Figure 5C | Clustering coefficient | BM1 learner vs. Beginner | Day 2 | No assumption | Kruskall-Wallis test | z = -1.98 | 0.048 | Bonferroni (0.05 / 12 days) | r = -0.24 |
| 51 | Figure 5C | Clustering coefficient | BM1 learner vs. Beginner | Day 3 | No assumption | Kruskall-Wallis test | z = -3.30 | 0.001* | Bonferroni (0.05 / 12 days) | r = -0.40 |
| 52 | Figure 5C | Clustering coefficient | BM1 learner vs. Beginner | Day 4 | No assumption | Kruskall-Wallis test | z = -2.74 | 0.006 | Bonferroni (0.05 / 12 days) | r = -0.33 |
| 53 | Figure 5C | Clustering coefficient | BM1 learner vs. Beginner | Day 5 | No assumption | Kruskall-Wallis test | z = -0.99 | 0.320 | Bonferroni (0.05 / 12 days) | r = -0.12 |
| 54 | Figure 5C | Clustering coefficient | BM1 learner vs. Beginner | Day 6 | No assumption | Kruskall-Wallis test | z = 0.19 | 0.853 | Bonferroni (0.05 / 12 days) | r = 0.02 |
| 55 | Figure 5C | Clustering coefficient | BM1 learner vs. Beginner | Day 7 | No assumption | Kruskall-Wallis test | z = -0.58 | 0.559 | Bonferroni (0.05 / 12 days) | r = -0.07 |
| 56 | Figure 5C | Clustering coefficient | BM1 learner vs. Beginner | Day 8 | No assumption | Kruskall-Wallis test | z = -1.64 | 0.101 | Bonferroni (0.05 / 12 days) | r = -0.20 |
| 57 | Figure 5C | Clustering coefficient | BM1 learner vs. Beginner | Day 9 | No assumption | Kruskall-Wallis test | z = -2.15 | 0.032 | Bonferroni (0.05 / 12 days) | r = -0.26 |
| 58 | Figure 5C | Clustering coefficient | BM1 learner vs. Beginner | Day 10 | No assumption | Kruskall-Wallis test | z = -0.47 | 0.640 | Bonferroni (0.05 / 12 days) | r = -0.06 |
| 59 | Figure 5C | Clustering coefficient | BM1 learner vs. Beginner | Day 11 | No assumption | Kruskall-Wallis test | z = -1.63 | 0.102 | Bonferroni (0.05 / 12 days) | r = -0.20 |
| 60 | Figure 5C | Clustering coefficient | BM1 learner vs. Beginner | Day 12 | No assumption | Kruskall-Wallis test | z = -0.66 | 0.507 | Bonferroni (0.05 / 12 days) | r = -0.08 |
| 61 | Figure 5C | Shortest path length | BM1 learner vs. Beginner | Day 1 | No assumption | Kruskall-Wallis test | z = 0.56 | 0.575 | Bonferroni (0.05 / 12 days) | r = 0.07 |
| 62 | Figure 5C | Shortest path length | BM1 learner vs. Beginner | Day 2 | No assumption | Kruskall-Wallis test | z = 0.37 | 0.713 | Bonferroni (0.05 / 12 days) | r = 0.04 |
| 63 | Figure 5C | Shortest path length | BM1 learner vs. Beginner | Day 3 | No assumption | Kruskall-Wallis test | z = -0.91 | 0.361 | Bonferroni (0.05 / 12 days) | r = -0.11 |
| 64 | Figure 5C | Shortest path length | BM1 learner vs. Beginner | Day 4 | No assumption | Kruskall-Wallis test | z = -3.02 | 0.003* | Bonferroni (0.05 / 12 days) | r = -0.36 |
| 65 | Figure 5C | Shortest path length | BM1 learner vs. Beginner | Day 5 | No assumption | Kruskall-Wallis test | z = -1.20 | 0.228 | Bonferroni (0.05 / 12 days) | r = -0.15 |
| 66 | Figure 5C | Shortest path length | BM1 learner vs. Beginner | Day 6 | No assumption | Kruskall-Wallis test | z = -0.93 | 0.353 | Bonferroni (0.05 / 12 days) | r = -0.11 |
| 67 | Figure 5C | Shortest path length | BM1 learner vs. Beginner | Day 7 | No assumption | Kruskall-Wallis test | z = -1.17 | 0.243 | Bonferroni (0.05 / 12 days) | r = -0.14 |
| 68 | Figure 5C | Shortest path length | BM1 learner vs. Beginner | Day 8 | No assumption | Kruskall-Wallis test | z = -1.80 | 0.073 | Bonferroni (0.05 / 12 days) | r = -0.22 |
| 69 | Figure 5C | Shortest path length | BM1 learner vs. Beginner | Day 9 | No assumption | Kruskall-Wallis test | z = 0.08 | 0.939 | Bonferroni (0.05 / 12 days) | r = 0.01 |
| 70 | Figure 5C | Shortest path length | BM1 learner vs. Beginner | Day 10 | No assumption | Kruskall-Wallis test | z = -1.65 | 0.099 | Bonferroni (0.05 / 12 days) | r = -0.20 |
| 71 | Figure 5C | Shortest path length | BM1 learner vs. Beginner | Day 11 | No assumption | Kruskall-Wallis test | z = -1.91 | 0.056 | Bonferroni (0.05 / 12 days) | r = -0.23 |
| 72 | Figure 5C | Shortest path length | BM1 learner vs. Beginner | Day 12 | No assumption | Kruskall-Wallis test | z = 0.13 | 0.896 | Bonferroni (0.05 / 12 days) | r = 0.02 |
| 73 | Figure 5C | Betweenness centrality | BM1 learner vs. Beginner | Day 1 | No assumption | Kruskall-Wallis test | z = 2.40 | 0.016 | Bonferroni (0.05 / 12 days) | r = 0.29 |
| 74 | Figure 5C | Betweenness centrality | BM1 learner vs. Beginner | Day 2 | No assumption | Kruskall-Wallis test | z = 1.90 | 0.058 | Bonferroni (0.05 / 12 days) | r = 0.23 |
| 75 | Figure 5C | Betweenness centrality | BM1 learner vs. Beginner | Day 3 | No assumption | Kruskall-Wallis test | z = 1.24 | 0.217 | Bonferroni (0.05 / 12 days) | r = 0.15 |
| 76 | Figure 5C | Betweenness centrality | BM1 learner vs. Beginner | Day 4 | No assumption | Kruskall-Wallis test | z = -1.50 | 0.135 | Bonferroni (0.05 / 12 days) | r = -0.18 |
| 77 | Figure 5C | Betweenness centrality | BM1 learner vs. Beginner | Day 5 | No assumption | Kruskall-Wallis test | z = -0.54 | 0.586 | Bonferroni (0.05 / 12 days) | r = -0.07 |
| 78 | Figure 5C | Betweenness centrality | BM1 learner vs. Beginner | Day 6 | No assumption | Kruskall-Wallis test | z = -0.30 | 0.765 | Bonferroni (0.05 / 12 days) | r = -0.04 |
| 79 | Figure 5C | Betweenness centrality | BM1 learner vs. Beginner | Day 7 | No assumption | Kruskall-Wallis test | z = -0.52 | 0.602 | Bonferroni (0.05 / 12 days) | r = -0.06 |
| 80 | Figure 5C | Betweenness centrality | BM1 learner vs. Beginner | Day 8 | No assumption | Kruskall-Wallis test | z = -0.81 | 0.420 | Bonferroni (0.05 / 12 days) | r = -0.10 |
| 81 | Figure 5C | Betweenness centrality | BM1 learner vs. Beginner | Day 9 | No assumption | Kruskall-Wallis test | z = 0.62 | 0.534 | Bonferroni (0.05 / 12 days) | r = 0.07 |
| 82 | Figure 5C | Betweenness centrality | BM1 learner vs. Beginner | Day 10 | No assumption | Kruskall-Wallis test | z = -1.35 | 0.177 | Bonferroni (0.05 / 12 days) | r = -0.16 |
| 83 | Figure 5C | Betweenness centrality | BM1 learner vs. Beginner | Day 11 | No assumption | Kruskall-Wallis test | z = -1.15 | 0.250 | Bonferroni (0.05 / 12 days) | r = -0.14 |
| 84 | Figure 5C | Betweenness centrality | BM1 learner vs. Beginner | Day 12 | No assumption | Kruskall-Wallis test | z = 1.28 | 0.200 | Bonferroni (0.05 / 12 days) | r = 0.15 |
| 85 | Figure 5C | Closeness centrality | BM1 learner vs. Beginner | Day 1 | No assumption | Kruskall-Wallis test | z = 0.73 | 0.466 | Bonferroni (0.05 / 12 days) | r = 0.09 |
| 86 | Figure 5C | Closeness centrality | BM1 learner vs. Beginner | Day 2 | No assumption | Kruskall-Wallis test | z = 0.21 | 0.836 | Bonferroni (0.05 / 12 days) | r = 0.02 |
| 87 | Figure 5C | Closeness centrality | BM1 learner vs. Beginner | Day 3 | No assumption | Kruskall-Wallis test | z = 0.44 | 0.662 | Bonferroni (0.05 / 12 days) | r = 0.05 |
| 88 | Figure 5C | Closeness centrality | BM1 learner vs. Beginner | Day 4 | No assumption | Kruskall-Wallis test | z = -0.67 | 0.504 | Bonferroni (0.05 / 12 days) | r = -0.08 |
| 89 | Figure 5C | Closeness centrality | BM1 learner vs. Beginner | Day 5 | No assumption | Kruskall-Wallis test | z = 1.56 | 0.119 | Bonferroni (0.05 / 12 days) | r = 0.19 |
| 90 | Figure 5C | Closeness centrality | BM1 learner vs. Beginner | Day 6 | No assumption | Kruskall-Wallis test | z = -0.97 | 0.330 | Bonferroni (0.05 / 12 days) | r = -0.12 |
| 91 | Figure 5C | Closeness centrality | BM1 learner vs. Beginner | Day 7 | No assumption | Kruskall-Wallis test | z = 0.75 | 0.452 | Bonferroni (0.05 / 12 days) | r = 0.09 |
| 92 | Figure 5C | Closeness centrality | BM1 learner vs. Beginner | Day 8 | No assumption | Kruskall-Wallis test | z = -0.77 | 0.438 | Bonferroni (0.05 / 12 days) | r = -0.09 |
| 93 | Figure 5C | Closeness centrality | BM1 learner vs. Beginner | Day 9 | No assumption | Kruskall-Wallis test | z = -0.97 | 0.330 | Bonferroni (0.05 / 12 days) | r = -0.12 |
| 94 | Figure 5C | Closeness centrality | BM1 learner vs. Beginner | Day 10 | No assumption | Kruskall-Wallis test | z = -1.01 | 0.315 | Bonferroni (0.05 / 12 days) | r = -0.12 |
| 95 | Figure 5C | Closeness centrality | BM1 learner vs. Beginner | Day 11 | No assumption | Kruskall-Wallis test | z = 0.10 | 0.921 | Bonferroni (0.05 / 12 days) | r = 0.01 |
| 96 | Figure 5C | Closeness centrality | BM1 learner vs. Beginner | Day 12 | No assumption | Kruskall-Wallis test | z = -1.78 | 0.075 | Bonferroni (0.05 / 12 days) | r = -0.21 |

Note. Asterisks indicate statistically significant differences. ES: effect size.
