## Supplementary material for "Scale space calibrates present and subsequent spatial learning in Barnes maze in mice": Statistical table 9

Statistical results of network analysis for the BM1 learner and the Beginner in the BM3 training phase.

| Manuscript reference # | Figure | Measure | Comparison | Within | Data structure | Type of test | Statistic | p | Correction | ES |
| --- | --- | --- | --- | --- | --- | --- | --- | --- | --- | --- |
| 1 | Figure 5E | Number of stops | BM1 learner vs. Beginner | Probe test | No assumption | Wilcoxon rank-sum test | z = -1.00 | 0.32 | N/A | r = -0.14 |
| 2 | Figure 5E | Order | BM1 learner vs. Beginner | Probe test | No assumption | Wilcoxon rank-sum test | z = -1.45 | 0.15 | N/A | r = -0.20 |
| 3 | Figure 5E | Degree | BM1 learner vs. Beginner | Probe test | No assumption | Wilcoxon rank-sum test | z = 3.97 | 0.00* | N/A | r = 0.55 |
| 4 | Figure 5E | Density | BM1 learner vs. Beginner | Probe test | No assumption | Wilcoxon rank-sum test | z = 2.90 | 0.00* | N/A | r = 0.40 |
| 5 | Figure 5E | Clustering coefficient | BM1 learner vs. Beginner | Probe test | No assumption | Wilcoxon rank-sum test | z = 4.40 | 0.00* | N/A | r = 0.60 |
| 6 | Figure 5E | Shortest path length | BM1 learner vs. Beginner | Probe test | No assumption | Wilcoxon rank-sum test | z = -2.58 | 0.01* | N/A | r = -0.35 |
| 7 | Figure 5E | Betweenness centrality | BM1 learner vs. Beginner | Probe test | No assumption | Wilcoxon rank-sum test | z = -2.95 | 0.00* | N/A | r = -0.40 |
| 8 | Figure 5E | Closeness centrality | BM1 learner vs. Beginner | Probe test | No assumption | Wilcoxon rank-sum test | z = 2.84 | 0.00* | N/A | r = 0.39 |

Note. Asterisks indicate statistically significant differences. N/A: not applicable. ES: effect size.
