## Extended Data 1 for "Scale space calibrates present and subsequent spatial learning in Barnes maze in mice": analyze_cnv_v1.m

```
% analyze conventional indices
function cnvObj = analyze_cnv_v1(paramObj, tbl, order)


varnames = tbl.Properties.VariableNames;
ind = [find(strcmp(varnames,'Day')),find(strcmp(varnames,'SN')),find(strcmp(varnames,'Trial'))];
tbl = sortrows(tbl, ind);

groups = unique(tbl.Group);
trials = unique(tbl.Trial);

cnvObj = struct('tbl_trn_daily',{}, 'tbl_trn_sum',{}, 'tbl_trn_stat',{},...
    'tbl_prb_daily',{}, 'tbl_prb_sum',{}, 'tbl_prb_stat',{});
featurenames = {'no_of_errors', 'latency', 'travel_distance', 'approach'};


%% get daily mean @ training

if all(ismember(tbl.Phase,{'training','retraining','reversal_training'}))
    
    data = tbl(ismember(tbl.Phase,{'training','retraining','reversal_training'}), ['Day','Group','SN',featurenames(1:3)]);    
    SN = unique(tbl.SN);
    daynums = unique(data.Day);
    
    C = mat2cell(data{:,featurenames(1:3)}, ones(size(data,1)/length(trials),1)*length(trials), 3);
    C = cellfun(@(x) mean(x,1,'omitnan'), C, 'uniform',0);
    C = vertcat(C{:});    
    
    C = array2table(C); C.Properties.VariableNames = featurenames(1:3);
    C = [data(1:length(trials):end, {'Day','Group','SN'}), C];
    
    tbl_trn_daily = C;
    cnvObj(1).tbl_trn_daily = tbl_trn_daily;

    
    
    %% make cross table (original) @ training
    tbl_trn_sum = cell(3*length(groups)*2, length(daynums)+3);
    str_cnvs = repmat(featurenames(1:3)', length(groups)*2,1);
    str_groups = repmat(groups, 3*2,1);
    str_params = repmat({'mu';'ci'}, 3*length(groups),1);
    tbl_trn_sum(:,1:2) = sortrows([str_cnvs, sort(str_groups)],1);
    tbl_trn_sum(:,3) = str_params;
    tbl_trn_sum = cell2table(tbl_trn_sum);
    tbl_trn_sum.Properties.VariableNames =...
        ['Feature', 'Group', 'Statistic', strcat('Day_',cellfun(@num2str,num2cell(daynums),'uniform',0))'];
    mu4sum = strcmp(tbl_trn_sum.Statistic, 'mu');
    
    for i = 1:length(groups)
        for j = 1:length(daynums)
            ind = strcmp(tbl_trn_daily.Group, groups{i})&...
                (tbl_trn_daily.Day==daynums(j));
            
            group4sum = strcmp(tbl_trn_sum.Group, groups{i});
            daynames = strcat('Day_',num2str(daynums(j)));
            
            data = tbl_trn_daily{ind, featurenames(1:3)};
            
            Cij_mu = num2cell(median(data,'omitnan'));
            Cij_ci = num2cell(1.96.*(std(data)./sqrt(sum(ind))));
            
            for k = 1:3
                cnvs4sum = strcmp(tbl_trn_sum.Feature, featurenames{k});
                row = (cnvs4sum.*group4sum.*mu4sum)==1;
                tbl_trn_sum(row, daynames) = {Cij_mu(k)};
                row = (cnvs4sum.*group4sum.*~mu4sum)==1;
                tbl_trn_sum(row, daynames) = {Cij_ci(k)};
            end
            
        end
    end
    
    Lev = [find(strcmp(tbl_trn_sum.Feature,featurenames{1})),...
        find(strcmp(tbl_trn_sum.Feature,featurenames{2})),...
        find(strcmp(tbl_trn_sum.Feature,featurenames{3}))]; % •À‚×‘Ö‚¦ƒCƒ“ƒfƒbƒNƒX
    
    tbl_trn_sum = tbl_trn_sum(Lev,:);
    cnvObj(2).tbl_trn_sum = tbl_trn_sum;
    
    
    
    %% statistical analysis @ training
    if order{'Conventional','Stats'}
        
        [~,~,G] = unique(tbl_trn_daily.Group);
        [~,~,D] = unique(tbl_trn_daily.Day);
        NumSbjDay = arrayfun(@(x) length(find(D==x)), unique(D), 'Uniform',0);
        NumSbjDay = unique(cell2mat(NumSbjDay));
        
        tbl_trn_stat = struct('tbl_anova',{},...
            'tbl_mult_Dxgn',{}, 'tbl_mult_Gxdn',{},...
            'tbl_mult_G',{}, 'tbl_mult_D',{});
        tbl_anova = cell2table(cell(0,7));
        mult_Dxgn = cell(0); % for interaction
        mult_Gxdn = cell(0);
        mult_G = cell(0); % for main effect
        mult_D = cell(0);
        
        for i = 1:3
            
            Data = sortrows([G, D, tbl_trn_daily{:, featurenames(i)}],[2,1]);
            X = [Data(:,[3,1,2]),repmat(1:NumSbjDay, 1,length(daynums))'];
            [~,~,~,~,p] = mixed_between_within_anova(X,1);
            p = [p{:}]; p(isnan(p)) = 0;
            
            % ANOVA
            [Report,~,~,~,~,~,~,~] = anova_2bw(Data);
            Report = Report([2,4:8], :);
            z = cell(size(Report,1), 7);
            
            for j = 1:size(Report, 1)
                y = strsplit(Report(j, :));
                y(cellfun('isempty',y)) = [];
                z(j,1:length(y)) = y;
            end
            a = ~cellfun(@isempty, z(:,strcmp(z(1,:),'p')));
            b = strcmp(z(a,strcmp(z(1,:),'p')), 'NaN'); b = b(2:end);
            
            z(strcmp(z,'NaN')) = cellfun(@num2str,num2cell(p(b)),'uniform',0);
            Z = cell2table(z(2:end,:));

            z(1,strcmp(z(1,:),'pEta^2')) = {'SquaredpEta'};
            Z.Properties.VariableNames = z(1,:);
            tbl_anova.Properties.VariableNames = z(1,:);
            tbl_anova = [tbl_anova; Z];
            
            
            % multiple comparison
            if p(3)<.05 % for interaction
                
                for Fct = 1:2
                    
                    [~,~,~,~,~,p,~,~] = sme_2bw(Data,Fct);
                    %                 [~,~,~,~,~,p,~,~] = sme_2ww(Data,Fct);
                    daynames = strcat('Day_', cellfun(@num2str,num2cell(daynums),'uniform',0));
                    LevNames = cell(max([length(daynames),length(groups)]),2);
                    LevNames(1:length(groups),1) = groups;
                    LevNames(1:length(daynames),2) = daynames;
                    
                    p(isnan(p)) = 0;
                    
                    if any(p<.05)
                        Lev = find(p<.05);
                        for k = 1:length(Lev)
                            
                            [Report,~,~,Result5,~,~] = tukey_sme_2bw(Data,Fct,Lev(k));
                            FctLev = strtrim(Report(1,:));
                            marker = cell(size(Result5)); marker(triu(Result5)) = {'*'};
                            
                            if Fct==1
                                r = [repmat(featurenames(i),length(groups),1),...
                                    repmat(daynames(Lev(k)),length(groups),1), groups];
                                mult_Dxgn = [mult_Dxgn; [r,marker;cell(1,size([r,marker],2))]];
                            else
                                r = [repmat(featurenames(i),length(daynames),1),...
                                    repmat(groups(Lev(k)),length(daynames),1), daynames];
                                mult_Gxdn = [mult_Gxdn; [r,marker;cell(1,size([r,marker],2))]];
                            end
                            
                        end
                    end
                end
                
            elseif any(p(1:2)<.05) % for main effect
                
                SigMainEffect = find(p(1:2)<.05);
                for Fct = 1:length(SigMainEffect)
                    [~,~,~,Result5,~,~] = tukey_2bw(Data,SigMainEffect(Fct));
                    marker = cell(size(Result5)); marker(triu(Result5)) = {'*'};
                    daynames = strcat('Day_',cellfun(@num2str,num2cell(daynums),'uniform',0));
                    
                    
                    switch SigMainEffect(Fct)
                        case 1
                            r = [repmat(featurenames(i),length(groups),1), groups];
                            mult_G = [mult_G; [r,marker;cell(1,size([r,marker],2))]];
                        case 2
                            r = [repmat(featurenames(i),length(daynames),1), daynames];
                            mult_D = [mult_D; [r,marker;cell(1,size([r,marker],2))]];
                    end
                    
                end
            end
            
            
        end
        
        rownames = cell(3*5,1); rownames(1:5:end) = featurenames(1:3);
        rownames = table(rownames);
        rownames.Properties.VariableNames = {'Feature'};
        tbl_anova = [rownames,tbl_anova];
        tbl_trn_stat(1).tbl_anova = tbl_anova;
        
        tbl_mult_Dxgn = cell2table(mult_Dxgn);
        if ~isempty(tbl_mult_Dxgn)
            tbl_mult_Dxgn.Properties.VariableNames =...
                ['Feature', 'Day', 'Group', groups'];
            tbl_trn_stat(2).tbl_mult_Dxgn = tbl_mult_Dxgn;
        else
            tbl_trn_stat(2).tbl_mult_Dxgn = [];
        end
        
        
        tbl_mult_Gxdn = cell2table(mult_Gxdn);
        if ~isempty(tbl_mult_Gxdn)
            tbl_mult_Gxdn.Properties.VariableNames =...
                ['Feature', 'Group', 'Day', daynames'];
            tbl_trn_stat(3).tbl_mult_Gxdn = tbl_mult_Gxdn;
        else
            tbl_trn_stat(3).tbl_mult_Gxdn = [];
        end
        
        
        tbl_mult_G = cell2table(mult_G);
        if ~isempty(tbl_mult_G)
            tbl_mult_G.Properties.VariableNames =...
                ['Feature', 'Group', groups'];
            tbl_trn_stat(4).tbl_mult_G = tbl_mult_G;
        else
            tbl_trn_stat(4).tbl_mult_G = [];
        end
        
        
        tbl_mult_D = cell2table(mult_D);
        if ~isempty(tbl_mult_D)
            tbl_mult_D.Properties.VariableNames =...
                ['Feature', 'Day', daynames'];
            tbl_trn_stat(5).tbl_mult_D = tbl_mult_D;
        else
            tbl_trn_stat(5).tbl_mult_D = [];
        end
        
        
        cnvObj(3).tbl_trn_stat = tbl_trn_stat;
        fprintf('\nStatistics......DONE\n')
        
    else
        cnvObj(3).tbl_trn_stat = [];
        fprintf('\nStatistics......SKIP\n')
    end
end


%% get daily mean @ probe test
if any(strcmp(tbl.Phase, 'probe_test'))
    
    data = tbl(strcmp(tbl.Phase, 'probe_test'), ['SN', 'Group', featurenames(end)]);
    Sbj = unique(data.SN);
    tbl_prb_daily = cell(length(Sbj)*paramObj.nHoles, 3);
    Lev = table2cell(data(:,{'SN','Group'}));
    Lev = sortrows(repmat(Lev, paramObj.nHoles,1), 1);
    Lev = [Lev, num2cell(repmat((1:paramObj.nHoles)',length(Sbj),1))];
    tbl_prb_daily(:,1:3) = Lev;
    tbl_prb_daily = cell2table(tbl_prb_daily);
    tbl_prb_daily.Properties.VariableNames = {'SN', 'Group', 'Holes'};
    D = cell(length(Sbj)*paramObj.nHoles, 1);
    for i = 1:sum(strcmp(tbl.Phase, 'probe_test'))
        iSbj = tbl_prb_daily.SN==data{i,'SN'};
        iD = data{i,featurenames{end}}; iD = num2cell(iD{:});
        D(iSbj) = iD;
    end
    D = cell2table(D);
    D.Properties.VariableNames = featurenames(end);
    tbl_prb_daily = [tbl_prb_daily, D];
    cnvObj(4).tbl_prb_daily = tbl_prb_daily;
    
    
    %% make cross table (original) @ probe test
    Lev = cell(length(groups)*2, 2);
    Lev(:,1:2) =...
        [sort(repmat(groups,2,1)), repmat({'mu';'ci'},length(groups),1)];
    Lev = cell2table(Lev);
    Lev.Properties.VariableNames = {'Group', 'Statistic'};
    iMu = strcmp(Lev.Statistic, 'mu');
    
    D = cell(length(groups)*2, paramObj.nHoles);
    str_holes = strcat('Hole_', strsplit(num2str(1:paramObj.nHoles)));
    for i = 1:length(groups)
        iSbj = strcmp(tbl_prb_daily.Group, groups{i});
        data = tbl_prb_daily{iSbj, featurenames(end)};
        data = reshape(data, paramObj.nHoles, length(data)/paramObj.nHoles)';
        
        mu = num2cell(mean(data));
        ci = num2cell(1.96.*(std(data)./sqrt(sum(iSbj)/paramObj.nHoles)));
        
        iGrp = strcmp(Lev.Group, groups{i});
        D(iGrp.*iMu==1, :) = mu;
        D(iGrp.*~iMu==1, :) = ci;
    end
    
    D = cell2table(D);
    D.Properties.VariableNames = str_holes;
    tbl_prb_sum = [Lev,D];
    cnvObj(5).tbl_prb_sum = tbl_prb_sum;
    
    
    %% statistical analysis @ probe test
    if order{'Conventional','Stats'}
        
        [~,~,G] = unique(tbl_prb_daily.Group);
        [~,~,H] = unique(tbl_prb_daily.Holes);
        
        NumSbj = arrayfun(@(x)length(find(H==x)), unique(H), 'Uniform',0);
        NumSbj = unique(cell2mat(NumSbj));
        
        tbl_prb_stat = struct('tbl_anova',{},...
            'tbl_mult_Hxgn',{}, 'tbl_mult_Gxhn',{},...
            'tbl_mult_G',{}, 'tbl_mult_H',{});
        tbl_anova = cell2table(cell(0,7));
        mult_Hxgn = cell(0); % for interaction
        mult_Gxhn = cell(0);
        mult_G = cell(0); % for main effect
        mult_H = cell(0);
        
        Data = sortrows([G, H, tbl_prb_daily{:,featurenames(end)}], 2);
        
        X = [Data(:,[3,1,2]),repmat(1:NumSbj, 1,paramObj.nHoles)'];
        [~,~,~,~,p] = mixed_between_within_anova(X,1);
        p = [p{:}]; p(isnan(p)) = 0;
        
        % ANOVA
        [Report,~,~,~,~,~,~,~] = anova_2bw(Data);
        
        Report = Report([2,4:8], :);
        z = cell(size(Report,1),7);
        
        for j = 1:size(Report, 1)
            y = strsplit(Report(j, :));
            x = find(cellfun('isempty',y)); % ‹ó‚ÌƒZƒ‹‚ð’T‚·
            y(x) = [];
            z(j,1:length(y)) = y;
        end
        a = ~cellfun(@isempty, z(:,strcmp(z(1,:),'p')));
        b = strcmp(z(a,strcmp(z(1,:),'p')), 'NaN'); b = b(2:end);
        
        z(strcmp(z,'NaN')) = cellfun(@num2str, num2cell(p(b)), 'uniform',0);
        z(1,strcmp(z(1,:),'pEta^2')) = {'SquaredpEta'};
        Z = cell2table(z(2:end,:));
        Z.Properties.VariableNames = z(1,:);
        tbl_anova.Properties.VariableNames = z(1,:);
        tbl_anova = [tbl_anova; Z];
        
        
        % multiple comparison
        if p(3)<.05 % for interaction
            for Fct = 1:2
                
                [~,~,~,~,~,p,~,~] = sme_2bw(Data,Fct);
                p(isnan(p)) = 0;
                holenames = strcat('Hole_', strtrim(cellstr(num2str((1:paramObj.nHoles)'))));
                LevNames = cell(max([length(holenames),length(groups)]),2);
                LevNames(1:length(groups),1) = groups;
                LevNames(1:length(holenames),2) = holenames;
                
                if any(p<.05)
                    Lev = find(p<.05);
                    for k = 1:length(Lev)
                        
                        [Report,~,~,Result5,~,~] = tukey_sme_2bw(Data,Fct,Lev(k));
                        FctLev = strtrim(Report(1,:));
                        marker = cell(size(Result5)); marker(triu(Result5)) = {'*'};
                        if Fct==1
                            r = [repmat(featurenames(end),length(groups),1),...
                                repmat(holenames(Lev(k)),length(groups),1), groups];
                            mult_Hxgn = [mult_Hxgn; [r,marker;cell(1,size([r,marker],2))]];
                        else
                            r = [repmat(featurenames(end),length(holenames),1),...
                                repmat(groups(Lev(k)),length(holenames),1), holenames];
                            mult_Gxhn = [mult_Gxhn; [r,marker;cell(1,size([r,marker],2))]];
                        end
                        
                    end
                end
            end
            
        elseif any(p(1:2)<.05) % for main effect
            
            SigMainEffect = find(p(1:2)<.05);
            for Fct = 1:length(SigMainEffect);
                
                [~,~,~,Result5,~,~] = tukey_2bw(Data, SigMainEffect(Fct));
                marker = cell(size(Result5)); marker(triu(Result5)) = {'*'};
                holenames = strcat('Hole_',strtrim(cellstr(num2str((1:paramObj.nHoles)'))));
                
                switch SigMainEffect(Fct)
                    case 1
                        r = [repmat(featurenames(end),length(groups),1), groups];
                        mult_G = [mult_G; [r,marker;cell(1,size([r,marker],2))]];
                    case 2
                        r = [repmat(featurenames(end),length(holenames),1), holenames];
                        mult_H = [mult_H; [r,marker;cell(1,size([r,marker],2))]];
                end
                
            end
        end
        
        tbl_prb_stat(1).tbl_anova = tbl_anova;
        
        tbl_mult_Hxgn = cell2table(mult_Hxgn);
        if ~isempty(tbl_mult_Hxgn)
            tbl_mult_Hxgn.Properties.VariableNames = ['Feature', 'Hole', 'Group', groups'];
            tbl_prb_stat(2).tbl_mult_Hxgn = tbl_mult_Hxgn;
        else
            tbl_prb_stat(2).tbl_mult_Hxgn = [];
        end
        
        
        tbl_mult_Gxhn = cell2table(mult_Gxhn);
        if ~isempty(tbl_mult_Gxhn)
            tbl_mult_Gxhn.Properties.VariableNames = ['Feature', 'Group', 'Hole', holenames'];
            tbl_prb_stat(3).tbl_mult_Gxhn = tbl_mult_Gxhn;
        else
            tbl_prb_stat(3).tbl_mult_Gxhn = [];
        end
        
        
        tbl_mult_G = cell2table(mult_G);
        if ~isempty(tbl_mult_G)
            tbl_mult_G.Properties.VariableNames = ['Feature', 'Group', groups'];
            tbl_prb_stat(4).tbl_mult_G = tbl_mult_G;
        else
            tbl_prb_stat(4).tbl_mult_G = [];
        end
        
        
        tbl_mult_H = cell2table(mult_H);
        if ~isempty(tbl_mult_H)
            tbl_mult_H.Properties.VariableNames = ['Feature', 'Hole', holenames'];
            tbl_prb_stat(5).tbl_mult_H = tbl_mult_H;
        else
            tbl_prb_stat(5).tbl_mult_H = [];
        end
        
        cnvObj(6).tbl_prb_stat = tbl_prb_stat;
        fprintf('\nStatistics......DONE\n')
    else
        cnvObj(6).tbl_prb_stat = [];
        fprintf('\nStatistics......SKIP\n')
    end
    
else
    fields = fieldnames(cnvObj); fields = fields(4:end);
    cnvObj = rmfield(cnvObj,fields);
end
```
