## Extended Data 1 for "Scale space calibrates present and subsequent spatial learning in Barnes maze in mice": analyze_ntw_med_170215.m

```
function ntwObj = analyze_ntw_med_170215(paramObj, tbl)

varnames = tbl.Properties.VariableNames;
ind = [find(strcmp(varnames,'Day')),find(strcmp(varnames,'SN')),find(strcmp(varnames,'Trial'))];
tbl = sortrows(tbl, ind);

groups = unique(tbl.Group);
trials = unique(tbl.Trial);


featurenames = {'xy_1', 'n_1', 'l_1', 'm_1', 'CWS_1', 'rho_1', 'x_1', 'cl_1',...
    'xy_2', 'NumLinks_2', 'o_2', 'n_2', 'l_2', 'm_2', 'CWS_2', 'rho_2', 'x_2', 'cl_2'};
featurenames = featurenames(~ismember(featurenames, {'xy_1','xy_2','NumLinks_2'})); % ÊuÀWÆem[hÌ¶ÌNð­


%% get daily mean @ training
if all(ismember(tbl.Phase,{'training','retraining','reversal_training'}))
    
    ntwObj = struct('tbl_trn_daily',{}, 'tbl_trn_sum',{});
    
    data = tbl(ismember(tbl.Phase,{'training','retraining','reversal_training'}),...
        ['Day', 'Group', 'SN', featurenames]);
    daynums = unique(data.Day);
    
    C = mat2cell(data{:,featurenames}, ones(size(data,1)/length(trials),1)*length(trials), length(featurenames));
    C = cellfun(@mean, C, num2cell(ones(length(C),1)), 'uniform',0); C = vertcat(C{:});
    C = array2table(C); C.Properties.VariableNames = featurenames;
    C = [data(1:length(trials):end, {'Day','Group','SN'}), C];
    
    tbl_trn_daily = C;
    ntwObj(1).tbl_trn_daily = tbl_trn_daily;
    
    
    
    %% make cross table (original) @ training
    tbl_trn_sum = cell(length(featurenames)*length(groups)*2, length(daynums)+3);
    str_ntws = repmat(featurenames', length(groups)*2,1);
    str_groups = repmat(groups, length(featurenames)*2,1);
    str_params = repmat({'mu';'ci'}, length(featurenames)*length(groups),1);
    tbl_trn_sum(:,1:2) = sortrows([str_ntws, sort(str_groups)]);
    tbl_trn_sum(:,3) = str_params;
    tbl_trn_sum = cell2table(tbl_trn_sum);
    tbl_trn_sum.Properties.VariableNames =...
        ['Feature', 'Group', 'Statistic', strcat('Day_',cellfun(@num2str,num2cell(daynums'),'uniform',0))];
    mu4sum = strcmp(tbl_trn_sum.Statistic, 'mu');
    
    for i = 1:length(groups)
        for j = 1:length(daynums)
            ind = strcmp(tbl_trn_daily.Group, groups{i})&...
                (tbl_trn_daily.Day==daynums(j));
            
            group4sum = strcmp(tbl_trn_sum.Group, groups{i});
            daynames = strcat('Day_',num2str(daynums(j)));
            
            data = tbl_trn_daily{ind, featurenames};
            Cij_mu = num2cell(median(data));
            Cij_ci = num2cell(1.96.*(std(data)./sqrt(sum(ind))));
            
            for k = 1:length(featurenames)
                cnvs4sum = strcmp(tbl_trn_sum.Feature, featurenames{k});
                row = (cnvs4sum.*group4sum.*mu4sum)==1;
                tbl_trn_sum(row, daynames) = {Cij_mu(k)};
                row = (cnvs4sum.*group4sum.*~mu4sum)==1;
                tbl_trn_sum(row, daynames) = {Cij_ci(k)};
            end
            
        end
    end
    
    ind = [];
    for i = 1:length(featurenames)
        ind = [ind, find(strcmp(tbl_trn_sum.Feature,featurenames{i}))]; % À×Ö¦CfbNX
    end
    tbl_trn_sum = tbl_trn_sum(ind,:);
    ntwObj(2).tbl_trn_sum = tbl_trn_sum;
    
    
    
else
    %% get daily mean @ probe test
    ntwObj = struct('tbl_prb_daily',{}, 'tbl_prb_sum',{});
    tbl_prb_daily = tbl(strcmp(tbl.Phase, 'probe_test'), ['SN', 'Group', featurenames]);
    ntwObj(1).tbl_prb_daily = tbl_prb_daily;
    
    
    
    %% get cross table @ probe test
    ind = cell(length(groups)*2, 2);
    ind(:,1:2) =...
        [sort(repmat(groups,2,1)), repmat({'mu';'ci'},length(groups),1)];
    ind = cell2table(ind);
    ind.Properties.VariableNames = {'Group', 'Statistic'};
    iMu = strcmp(ind.Statistic, 'mu');
    
    D = cell(length(groups)*2, length(featurenames));
    for i = 1:length(groups)
        iSbj = strcmp(tbl_prb_daily.Group, groups{i});
        data = tbl_prb_daily{iSbj, featurenames};
        
        if iscell(data)
            mu = num2cell(median(cell2num(data)));
            ci = num2cell(1.96.*(std(cell2num(data))./sqrt(sum(iSbj)/paramObj.nHoles)));
        else
            mu = num2cell(median(data));
            ci = num2cell(1.96.*(std(data)./sqrt(sum(iSbj)/paramObj.nHoles)));
        end
        
        
        iGrp = strcmp(ind.Group, groups{i});
        D(iGrp.*iMu==1, :) = mu;
        D(iGrp.*~iMu==1, :) = ci;
    end
    
    D = cell2table(D);
    D.Properties.VariableNames = featurenames;
    tbl_prb_sum = [ind,D];
    ntwObj(2).tbl_prb_sum = tbl_prb_sum;
    
    
    
end
```
