## Extended Data 1 for "Scale space calibrates present and subsequent spatial learning in Barnes maze in mice": analyze_str_v0.m

```
% analyze conventional indices
function strObj = analyze_str_v0(tbl)

varnames = tbl.Properties.VariableNames;
ind = [find(strcmp(varnames,'Day')),find(strcmp(varnames,'SN')),find(strcmp(varnames,'Trial'))];
tbl = sortrows(tbl, ind);

groups = unique(tbl.Group);
trials = unique(tbl.Trial);

strObj = struct('tbl_trn_daily',{}, 'tbl_trn_sum',{});
featurenames = {'Random', 'Serial', 'Spatial', 'Perseveration'};

if any(strcmp(tbl.Properties.VariableNames,'Reversal'))
    if all(tbl.Reversal)
        featurenames = featurenames(1:4);
    else
        featurenames = featurenames(1:3);
    end
else
    featurenames = featurenames(1:3);
end


%% get daily sum @ training
if all(ismember(tbl.Phase,{'training','retraining','reversal_training'}))
    
    
    data = tbl(ismember(tbl.Phase,{'training','retraining','reversal_training'}),...
        ['Day', 'Group', 'SN', featurenames]);
    daynums = unique(data.Day);
    
    C = mat2cell(data{:,featurenames}, ones(size(data,1)/length(trials),1)*length(trials), length(featurenames));
    C = cellfun(@sum, C, num2cell(ones(length(C),1)), 'uniform',0); C = vertcat(C{:});
    C = array2table(C); C.Properties.VariableNames = featurenames;
    C = [data(1:length(trials):end, {'Day','Group','SN'}), C];
    
    tbl_trn_daily = C;
    strObj(1).tbl_trn_daily = tbl_trn_daily;       

    
    
    %% make cross table @ training
    tbl_trn_sum = cell(length(featurenames)*length(groups), length(daynums)+3);
    strnames = repmat(featurenames', length(groups),1);
    str_groups = sort(repmat(groups, length(featurenames),1));
    str_params = repmat({'sum'}, length(featurenames)*length(groups),1);
    tbl_trn_sum(:,1:2) = sortrows([strnames,str_groups],2);
    tbl_trn_sum(:,3) = str_params;
    tbl_trn_sum = cell2table(tbl_trn_sum);
    tbl_trn_sum.Properties.VariableNames =...
        ['Feature','Group','Statistic', strcat('Day_',cellfun(@num2str,num2cell(daynums),'uniform',0))'];
    sig4sum = strcmp(tbl_trn_sum.Statistic, 'sum');
    
    for i = 1:length(groups)
        for j = 1:length(daynums)
            
            ind = strcmp(tbl_trn_daily.Group, groups{i})&...
                (tbl_trn_daily.Day==daynums(j));
            
            group4sum = strcmp(tbl_trn_sum.Group, groups{i});
            dayname = strcat('Day_',num2str(daynums(j)));
            
            data = tbl_trn_daily{ind, featurenames};
            Cij_sum = num2cell(sum(data));
            
            for k = 1:length(featurenames)
                strs4sum = strcmp(tbl_trn_sum.Feature, featurenames{k});
                row = (strs4sum.*group4sum.*sig4sum)==1;
                tbl_trn_sum(row, dayname) = {Cij_sum(k)};
            end
            
        end
    end

    
    try
        tbl.Reversal;
    catch        
        Reversal = false(size(tbl,1),1); Reversal = table(Reversal);
        k = find(strcmp(tbl.Properties.VariableNames,'Trial'));
        tbl = [tbl(:,1:k),Reversal,tbl(:,k+1:end)];
    end
    
    if all(tbl.Reversal)
        ind = [find(strcmp(tbl_trn_sum.Feature,featurenames{1})),...
            find(strcmp(tbl_trn_sum.Feature,featurenames{2})),...
            find(strcmp(tbl_trn_sum.Feature,featurenames{3})),...
            find(strcmp(tbl_trn_sum.Feature,featurenames{4}))]; % •À‚×‘Ö‚¦ƒCƒ“ƒfƒbƒNƒX
    else
        ind = [find(strcmp(tbl_trn_sum.Feature,featurenames{1})),...
            find(strcmp(tbl_trn_sum.Feature,featurenames{2})),...
            find(strcmp(tbl_trn_sum.Feature,featurenames{3}))]; % •À‚×‘Ö‚¦ƒCƒ“ƒfƒbƒNƒX
    end


    tbl_trn_sum = tbl_trn_sum(ind,:);
    strObj(2).tbl_trn_sum = tbl_trn_sum;
    
end
```
