## Extended Data 1 for "Scale space calibrates present and subsequent spatial learning in Barnes maze in mice": feature_ntw_d_v0.m

```
function NetworkFeatures = feature_ntw_d_v0(parameters,NetworkFeatures,n,XY)

%% Behavioral network
% Stopping CorrdinatesÌZo
%{
    (1) Avni, et al., 2006, Behav Brain Res,
    (2) Huldot by Michael Lieberthal, Weiss et al., 2012, PLoS ONE
%}

T = size(XY,1);

allfeatures = NetworkFeatures.Properties.VariableNames;
dynamicfeatures = allfeatures(~cellfun(@isempty,strfind(allfeatures,'_2')));
bin = parameters.fps; % time window (default = 5 frame)

fin = 0;
initframe = 1;
step = 1;
StopPos = [];
while ~fin
    while ~fin
        a = XY(initframe:initframe+step, :);
        b = mat2cell(diff(a), ones(step,1),2);
        d = cellfun(@norm,b);
        fin = or(sum(d)>parameters.thresh4CCA, (initframe+step-1)>=T-1);
        step = step + 1;
    end
    
    if step-1>=bin
        StopPos = [StopPos; mean(a)];
    end
    fin = (initframe+step-1)>=T-1;
    initframe = initframe + step - 1;
    step = 1;
end


if size(StopPos,1)==0 % stop ªÈ¢êCÊ0ÌOt
    
    NetworkFeatures{n,dynamicfeatures(3:end)} = 0;
    NetworkFeatures.xy_2{n} = [];
    NetworkFeatures.A_2{n} = [];
    
elseif size(StopPos,1)==1 % stopping coordinates ª2¢ÌêCÊ1ÌÓÌÈ¢Ot
    
    NetworkFeatures.xy_2{n} = StopPos; % xy
    NetworkFeatures.A_2{n} = 1;
    NetworkFeatures.o_2(n) = 1; % Total number of stop
    NetworkFeatures.n_2(n) = 1; % Order
    NetworkFeatures.l_2(n) = 0; % Shortest path length (l)
    NetworkFeatures.m_2(n) = 0; % Degree/Connectivity (k)
    NetworkFeatures.CWS_2(n) = 0; % Clustering coefficient (C)
    NetworkFeatures.rho_2(n) = 0; % Network density (d)
    NetworkFeatures.x_2(n) = 0; % betweenness centrality (w)
    NetworkFeatures.cl_2(n) = 0; % closeness centrality (cl)
    
else % m[hª2ÂÈã éêÍð¢
    
    [NodePos,stop2node,~] = CCA(StopPos, parameters.thresh4CCA);
    NodePos = unique(NodePos, 'rows','stable');
    [~,~,node_indx] = unique(stop2node); % stopping coordinates ªÇÌ node ÉèÄçêÄ¢é©
    
    
    %% e Network parameter ÌZo
    
    % Total number of stop
    NetworkFeatures.o_2(n) = size(StopPos,1);
    
    % xy
    NetworkFeatures.xy_2{n} = NodePos;
    
    % transition
    NetworkFeatures.A_2{n} = node_indx;
    
    % Order
    NetworkFeatures.n_2(n) = size(NodePos,1);
    
    
    if size(NodePos,1) == 1 % ãÌm[hª1ÂÌê
        
        NetworkFeatures.l_2(n) = 0; % Shortest path length (l)
        NetworkFeatures.m_2(n) = 0; % Degree/Connectivity (k)
        NetworkFeatures.CWS_2(n) = 0; % Clustering coefficient (C)
        NetworkFeatures.rho_2(n) = 0; % Network density (d)
        NetworkFeatures.x_2(n) = 0; % Network density (bc)
        NetworkFeatures.cl_2(n) = 0; % Network density (cc)
        
    else
        
        DG = zeros(max(node_indx));
        DG(sub2ind(size(DG), node_indx(1:end-1), node_indx(2:end))) = 1;
        DG = triu(DG)+tril(DG)';
        DG = DG + DG';
        D = diag(ones(size(DG,1),1)); D = ~D;
        DG = DG.*D; % ×Úsñ
        
        [s,t] = find(DG);
        DG = sparse(s, t, ones(length(find(DG)),1), size(DG,1), size(DG,2));
        
        
        uqnode_indx = unique(node_indx);
        AllLinkComb = nchoosek(uqnode_indx, 2); % all pairs of source and destination node
        
        % Shortest path length (l)
        %{
                LüOtÌÅZoHð_CNXgÌû@Åßé
                http://goo.gl/c60yN7
        %}
        minPathLength = zeros(size(AllLinkComb,1), 1);
        for i = 1:size(AllLinkComb,1)
            [Dist, ~, ~] = graphshortestpath(DG, AllLinkComb(i,1), AllLinkComb(i,2),...
                'Directed', false,...
                'Method', 'Dijkstra');
            minPathLength(i) = Dist;
        end
        
        minPathLength = minPathLength(~isinf(minPathLength)); % InfÌ
        NetworkFeatures.l_2(n) = mean(minPathLength);
        
        
        % Betweenness
        w = node_betweenness_slow(DG);
        NetworkFeatures.x_2(n) = mean(w);
        
        
        % closeness centrality
        cl = closeness(DG);
        cl = cl.*size(NodePos,1); % Newman, 2010
        NetworkFeatures.cl_2(n) = mean(cl);
        
        
        % Degree/Connectivity (k)
        %{
                êÂÌm[h©çÌNÌ½Ï
        %}
        NetworkFeatures.m_2(n) = size([s,t],1)/size(NodePos,1);
        
        
        % Clustering coefficient (C)
        %{
                iÉ×ÚµÄ¢ékÂÌm[hÌÈ©Ìj, mÌQÂðIñ¾Æ«ÉC
                »ÌQÂª×ÚµÄ¢é©Ç¤©Ì
                iÆjÆmªOp`ÉÈéñ
        %}
        NetworkFeatures.CWS_2(n) = mean(clustering(full(DG)));
           
        
        % Network density (d)
        %{
                 ém[h©çÂ\ÈNÌÉÎ·éÀÛÌN
        %}
        NumPossibleLinks = 2*nchoosek(size(NodePos,1), 2);
        NetworkFeatures.rho_2(n) = size([s,t],1)/NumPossibleLinks;
        
        
    end
    
    
end
```
