## Extended Data 1 for "Scale space calibrates present and subsequent spatial learning in Barnes maze in mice": feature_ntw_s_v0.m

```
function NetworkFeatures = feature_ntw_s_v0(parameters,NetworkFeatures,n,XY)


%% real-coord network
%{
    S©çÀWÜÅÌxNgvÌå«³ÆC
    S©çÜÅÌxNghÆÌÈ·p
    1. Node x 12 @ Outer
    2. Node x 1  @ Center
    3. Node x 12 @ Pericenter
%}


T = size(XY,1);
nodes = zeros(T, 5); % [x, y, node ID, node weight, link weight]
for i = 1:T
    
    % S©çÀWÜÅÌxNgvÌå«³
    edges = [parameters.CenterEdge; parameters.CenterEdge*2];
    vOX = XY(i, :)-parameters.O;
    nv = repmat(norm(vOX), length(edges), 1);
    SortedTransit = nv-edges > 0;
    
    % vÆS©çÜÅÌxNghÆÌÈ·p
    vOX = repmat(vOX, parameters.nHoles, 1);
    vOH = parameters.holes - repmat(parameters.O, parameters.nHoles, 1);
    InnerProduct = zeros(1, T-1);
    CrossProduct = zeros(1, T-1);
    cos = zeros(1, parameters.nHoles);
    theta = zeros(1, parameters.nHoles);
    
    for j = 1:parameters.nHoles
        InnerProduct(j) = dot(vOH(j, :), vOX(j, :));
        CrossProduct(j) = vOH(j, 1)*vOX(j, 2) - vOH(j, 2)*vOX(j, 1);
        
        vOH_norm = norm(vOH(j, :));
        vOX_norm = norm(vOX(j, :));
        cos(j) = InnerProduct(j)/(vOH_norm*vOX_norm);
        theta(j) = acos(cos(j))*180/pi;
    end
    
    node_indx = find(theta==min(theta), 1);
    
    
    % »è
    if and(SortedTransit(1), SortedTransit(2)) % outer layer
        nodes(i, 1:2) = parameters.holes(node_indx, :);
        nodes(i, 3) = node_indx;
    elseif xor(SortedTransit(1), SortedTransit(2)) % middle layer
        nodes(i, 1:2) = parameters.middle_nodes(node_indx, :);
        nodes(i, 3) = node_indx + parameters.nHoles;
    elseif not(SortedTransit) % inner layer
        nodes(i, 1:2) = parameters.O;
        nodes(i, 3) = parameters.nNodes;
    else
        nodes(i, 1:2) = [NaN, NaN];
        nodes(i, 3) = NaN;
    end
    
    
end

nodes = nodes(~isnan(nodes(:, 3)), :);


PossibleLinks = parameters.PossibleLinks;
linkdiff = abs(diff(PossibleLinks,1,2));


% Â\ÈNÌièÝ
%icentar2midnode, midnode2midnode, midnode2hole, hole2holej
o2mid = all([any(PossibleLinks==25,2),...
    any(PossibleLinks<13,2)], 2);
mid2mid = all([and(PossibleLinks(:,1)<13,PossibleLinks(:,2)<13),...
    ismember(linkdiff,[1,11])], 2);
mid2hole = and(and(PossibleLinks(:,1)<13,PossibleLinks(:,2)>=13),...
    linkdiff==12);
hole2hole = all([and(PossibleLinks(:,1)>=13,PossibleLinks(:,2)>=13),...
    all(PossibleLinks~=25,2),...
    ismember(linkdiff,[1,11])], 2);


PossibleLinks = PossibleLinks(any([mid2mid,mid2hole,hole2hole,o2mid],2),:);
PossibleLinks = sortrows([PossibleLinks;fliplr(PossibleLinks)]);
G = sparse(PossibleLinks(:,1), PossibleLinks(:,2), ones(size(PossibleLinks,1),1));
% ×ÚµÄ¢È¢m[hÖÌÚ®ðo&C³
x = [];
for i = 1:size(nodes,1)-1    
    [~, path, ~] = graphshortestpath(G,...
        nodes(i,3), nodes(i+1,3),...
        'Directed', false,...
        'Method', 'Dijkstra');    
    x = [x,path];
end
areas = [parameters.middle_nodes; parameters.holes; parameters.O];
nodes = [areas(x,:), x', zeros(length(x),3)];
T2 = size(nodes,1);


% em[hÖÌiüñÌZo
uqNodes = unique(nodes(:, 3), 'stable');
[NumUqNodes, ~] = size(uqNodes);
NodeWT = zeros(1, parameters.nNodes);

for i = 1:NumUqNodes
    NodeEntry = find(nodes(:, 3)==uqNodes(i));
    nNodeEntry = length(NodeEntry);
    nodes(NodeEntry, 4) = nNodeEntry;
    NodeWT(uqNodes(i)) = nNodeEntry;
end


% eNÌgpñÌZo
state = [nodes(1:end-1, 3), nodes(2:end, 3)]; % transition from node A (src) to node B (destination)
GetTransit = zeros(T2, 1); % transition o
GetTransit(1:end-1) = state(:, 1)~=state(:, 2);
nodes(:, 5) = GetTransit;
transit = state(logical(GetTransit), :);
[nTransit, ~] = size(transit);
LinkWT = zeros(nTransit, 1); % No. of Links

for i = 1:nTransit
    forward =  ismember(transit,...
        repmat(transit(i, :), nTransit, 1),'rows'); % node A ==> node B
    backward = ismember(transit,...
        repmat(fliplr(transit(i, :)), nTransit, 1),'rows'); % node A <== node B
    LinkWT(or(forward, backward)) = sum(or(forward, backward));
end

[SortedTransit, ~, ic] = unique(transit);
B = 1:length(SortedTransit);
transit_indx = reshape(B(ic), nTransit, 2); % TransitÌÊCfbNX


%% e Network parameter ÌZo
% Total number of nodes
NetworkFeatures.xy_1(n) = {nodes};
NetworkFeatures.n_1(n) = NumUqNodes;

if NumUqNodes<2
    
    NetworkFeatures.l_1(n) = 0;
    NetworkFeatures.m_1(n) = 0;
    NetworkFeatures.CWS_1(n) = 0;
    NetworkFeatures.rho_1(n) = 0;
    NetworkFeatures.x_1(n) = 0;
    NetworkFeatures.cl_1(n) = 0;
    
else
    % Shortest path length (l)
    %{
        LüOtÌÅZoHð_CNXgÌû@Åßé
        http://goo.gl/c60yN7
    %}
    uqNode_indx = 1:NumUqNodes;
    FullLinks = nchoosek(uqNode_indx, 2); % pairs of source and destination node
    
    UndirLinks = unique(transit_indx, 'rows', 'stable');
    UndirLinks = [UndirLinks; fliplr(UndirLinks)]; % ³üN
    UndirLinks = unique(UndirLinks, 'rows');
%     NumUndirLinks = size(UndirLinks,1);
    
    DG = sparse(UndirLinks(:,1), UndirLinks(:,2), ones(size(UndirLinks,1),1)); % ×Úsñ
    L = zeros(size(FullLinks,1), 1);
    
    %{
    lbg[N DG É¶Ý·ém[h©çC
    [FullLinks(i,1), FullLinks(i,2)] ÅyAm[hðwèµC
    »ÌÅZoH·ðßé
    %}
    for i = 1:size(FullLinks,1)
        [dist, ~, ~] = graphshortestpath(DG, FullLinks(i,1), FullLinks(i,2),...
            'Directed', false,...
            'Method', 'Dijkstra'); 
        L(i) = dist;
    end
    % h = view(biograph(DG,[],'ShowWeights','on')) % lbg[Ng|W[mF
    L = L(~isinf(L)); % InfÌ
    NetworkFeatures.l_1(n) = mean(L);
    
    
    % betweenness centrality
    w = node_betweenness_slow(DG);
    NetworkFeatures.x_1(n) = mean(w);
    
    % closeness centrality
    C = closeness(DG);
    NetworkFeatures.cl_1(n) = mean(C);
    
    
    % Clustering coefficient (C)
    %{
        iÉ×ÚµÄ¢ékÂÌm[hÌÈ©Ìj, mÌQÂðIñ¾Æ«ÉC
        »ÌQÂª×ÚµÄ¢é©Ç¤©Ì
        iÆjÆmªOp`ÉÈéñ
    %}
    NetworkFeatures.CWS_1(n) = mean(clustering(full(DG)));
        
    % Network density (d)
    %{
         ém[h©çÂ\ÈNÌÉÎ·éÀÛÌN
    %}
    NumPossibleLinks = 2*nchoosek(NumUqNodes, 2);
    NetworkFeatures.rho_1(n) = size(UndirLinks,1)/NumPossibleLinks;
    
    % Degree/Connectivity (k)
    %{
    êÂÌm[h©çÌNÌ½Ï
    %}
    NetworkFeatures.m_1(n) = size(UndirLinks,1)/NumUqNodes;
    
end


% Kâµ½j[NÈm[h
% AllVisitNodes = [AllVisitNodes; uqNodes];


% em[hÖÌKâñ
% AllNodeWTs(n, :) = NodeWT;


% gpµ½j[NÈN
% [A, ~, ~] = unique(sortrows([transit, LinkWT]), 'rows');
% AllLinksWTs = [AllLinksWTs; A];
```
