## Extended Data 1 for "Scale space calibrates present and subsequent spatial learning in Barnes maze in mice": feature_str_v1.m

```
function StrategyFeatures = feature_str_v1(parameters,options,LOG,StrategyFeatures,n,OnlineTracking,XY)

%% Calculate Strategy Indices
%{
Nat Commun. 2014 Aug 19;5:4701. doi: 10.1038/ncomms5701.
The MK2/3 cascade regulates AMPAR trafficking and cognitive flexibility.
Eales KL, Palygin O, O'Loughlin T, Rasooli-Nejad S, Gaestel M, M?ller J, Collins DR, Pankratov Y, Corr?a SA.
%}

T = size(XY,1);

holes = parameters.holes;
startframe = 1;

%{
75.63 pix:2 mm/pix (1mBM)
283.61 pix:7.5 mm/pix (3mBM)
32 pix:240 mm (core radius of 3mBM)
%}
threshold = parameters.bwHoles/2.5;

% X^[gGAÌf[^ðJEgµÈ¢
if OnlineTracking
    d = 0;
    while (d<=threshold)&&(startframe<T)
        d = norm(XY(startframe,:) - parameters.O);
        startframe = startframe + 1;
    end
end

XY = XY(startframe:end, :);
T = size(XY,1);

% quadrant »è
p = XY-ones(T,1)*parameters.O;
q = zeros(T,1);
for i = 1:T
    signxy = [sign(p(i,1)), sign(p(i,2))];
    if (signxy(1)==1)&&(signxy(2)==1)
        q(i) = 1;
    elseif (signxy(1)==1)&&(signxy(2)==-1)
        q(i) = 4;
    elseif (signxy(1)==-1)&&(signxy(2)==1)
        q(i) = 2;
    elseif (signxy(1)==-1)&&(signxy(2)==-1)
        q(i) = 3;
    end
end


% Av[`µ½ÌíÞ
threshold = 80/parameters.spr;
XH = zeros(T, parameters.nHoles);
ApproachHoles = zeros(T,1);
for i = 1:parameters.nHoles
    for j = 1:T
        XH(j,i) = norm(XY(j,:) - holes(i,:));
    end
    ApproachHoles(XH(:,i)<=threshold) = i;
end
Approach_indx = unique(ApproachHoles(ApproachHoles~=0), 'stable');

if isempty(Approach_indx)
    Approach_indx = 0;
end


% ½]OÌS[ÌTõ
XO = zeros(T,1);
O = mean(XY);
for i = 1:T
    XO(i) = norm(XY(i,:) - O);
end
stdU = parameters.RAD; % threshold

% Av[`µ½ðnñÉJEg
if isempty(ApproachHoles)
    serial_flag = 0;
else
    k = [1;(diff(ApproachHoles)~=0)];
    UniqueApproachHoles = ApproachHoles(logical(k));
    UniqueApproachHoles(UniqueApproachHoles==0) = [];
end


switch options.maze_dia
    case 1
        if LOG.Reversal(n)==1
            spatial_flag = all([all(ismember(Approach_indx,[6,7,8])),...
                all(q==1)]);
            perseveration_flag = and(mean(XO)/stdU<.45, mean(XH(:,1))/stdU<.40);
            serial_flag = all([all(ismember(abs(diff(UniqueApproachHoles)),[1,11])),...
                length(unique(q))<=2,...
                ~any(ismember(q,[0,3]))]);
        else
            spatial_flag = all([all(ismember(Approach_indx,[12,1,2])),...
                all(q==1)]);
            perseveration_flag = 0;
            serial_flag = all([all(ismember(abs(diff(UniqueApproachHoles)),[1,11])),...
                length(unique(q))<=2,...
                ~any(ismember(q,[0,3]))]);
        end
    case 3
        if LOG.Reversal(n)==1
            spatial_flag = all([all(ismember(Approach_indx,[6,7,8])),...
                all(q==1)]);
            perseveration_flag = and(mean(XO)/stdU<.45, mean(XH(:,1))/stdU<.40);
            serial_flag = all([all(ismember(abs(diff(UniqueApproachHoles)),[1,11])),...
                length(unique(q))<=2,...
                ~any(ismember(q,[0,3]))]);
        else
            spatial_flag = all([all(ismember(Approach_indx,[12,1,2])),...
                all(q==1)]);
            perseveration_flag = 0;
            serial_flag = all([all(ismember(abs(diff(UniqueApproachHoles)),[1,11])),...
                length(unique(q))<=2,...
                ~any(ismember(q,[0,3]))]);
        end
end


% Algorithm-based classification (cf. Garthe, et al., 2009)
if spatial_flag
    StrategyFeatures.Spatial(n) = 1;
elseif perseveration_flag
    StrategyFeatures.Perseveration(n) = 1;
elseif serial_flag
    StrategyFeatures.Serial(n) = 1;
else
    StrategyFeatures.Random(n) = 1;
end
```
