## Extended Data 1 for "Scale space calibrates present and subsequent spatial learning in Barnes maze in mice": FeatureExtraction_v0.m

```
function [ConventionalFeatures, StrategyFeatures, NetworkFeatures, ExtraFeatures] =...
    FeatureExtraction_v0(parameters, options, LOG,...
    ConventionalFeatures, StrategyFeatures, NetworkFeatures, ExtraFeatures)

N = size(LOG, 1);

for n = 1:N
    %% load individual data
    Day = LOG.Day(n);
    trial = LOG.Trial(n);
    sn = LOG.SN(n);
    goal = LOG.Goal(n);
    phase = LOG.Phase(n);
    group = LOG.Group(n);
    OnlineTracking = LOG.OnlineTracking(n);
    
    
    if strcmp(phase, 'habituation')
        fprintf('\nDay :[%d], Trial [%d], No.[%d], Group [%s], Phase [%s].......SKIP.\n',...
            Day, trial, sn, group{:}, phase{:});
        continue
    end
    
    file_path = fullfile(...
        parameters.PrjDir,...
        phase{:},...
        sprintf('d%d', Day),...
        sprintf('t%d', trial),...
        sprintf('#%d', sn));
    
    if exist([file_path,'\xy.txt'],'file')==0
        continue
    end
    
    data = dlmread([file_path,'\xy.txt']); % t@CÌwè
    
    fprintf('\nDay [%d], Trial [%d], No.[%d], Group [%s], Phase [%s]......START.\n',...
        Day, trial, sn, group{:}, phase{:});
    
    
    
    %% Get position
    if OnlineTracking % online tracking ¬÷Ìê
        initframe = 10;
        if strcmp(phase, 'probe_test') % probe test Ìê
            endframe = parameters.fps*150;
        else % training Ìê
            % S[ãÌNaNðµÄI¹t[ðßé
            tf = ~any(isnan(data(:,3:4)),2)&~all(data(:,3:4)==0,2);
            endframe = find(tf, 1, 'last');
        end
        data = data(initframe:endframe, :);
        x = data(:,3); y = data(:,4); xy = [x,y]; % ÀWZo
        
    else % online tracking ¸sÌê
        nan_ind = any(isnan(data(:,1:end-1)),2);
        if strcmp(phase, 'probe_test') % probe test Ìê
            endframe = parameters.fps*150;
        else % training Ìê
            endframe = find(~nan_ind, 1, 'last');
        end
        x1 = data(:,1); x2 = data(:,3); y1 = data(:,2); y2 = data(:,4); % ÀWZo
        xy = [(x1-x2)/2+x2, (y1-y2)/2+y2];
        xy = xy(1:endframe, :); % S[ãÌNaNðµÄI¹t[ðßé
    end
    
    options.timeout = size(xy,1)>parameters.fps*600;
    xy(parameters.fps*600:end,:) = []; % cutoff over 10 min
    
    XY = xy - parameters.RAD;
    
    if options.maze_dia==1
        switch goal{:} % dSÀWÌ½]
            case 'N'
                XY = [XY(:,1), -XY(:,2)];
            case 'E'
                XY = [XY(:,1), XY(:,2)];
            case 'W'
                XY = [-XY(:,1), -XY(:,2)];
            case 'S'
                XY = [-XY(:,1), XY(:,2)];
        end
    else
        switch goal{:} % dSÀWÌ½]
            case 'N'
                XY = [-XY(:,1), XY(:,2)];
            case 'E'
                XY = [-XY(:,1), -XY(:,2)];
            case 'W'
                XY = [XY(:,1), XY(:,2)];
            case 'S'
                XY = [XY(:,1), -XY(:,2)];
        end
    end
    
    XY = XY + parameters.RAD;
    
    nan_ind = or(isnan(XY(:,1)),isnan(XY(:,2)));
    zero_ind = (XY(:,1)==0)&(XY(:,2)==0);
    discard_ind = nan_ind|zero_ind;
    a = XY;
    a(discard_ind, :) = [];
    [b,~] = size(a);
    
    % Oêl(R[i[ðgbNµÄ¢éàÌ)ð
    outlier_ind = zeros(b,1);
    for i = 1:b
        outlier_ind(i) = norm(a(i,:)-parameters.O);
    end
    a(outlier_ind>parameters.RAD,:) = [];
    XY = a;
    T = size(XY,1);
    
    
    %%%%%%%%%%%%%%%%%%%%%%%%%%%%%%%%%%%%%%%%%%%%%%%%%%%%%%%%%%%%%%%%%%%%%%
    % X^[gGAðoÄ©çðÍ·éê
    %%%%%%%%%%%%%%%%%%%%%%%%%%%%%%%%%%%%%%%%%%%%%%%%%%%%%%%%%%%%%%%%%%%%%%
    if and(options.offset, ~strcmp(phase,'probe_test'))
        a = arrayfun(@(x,y) norm([x,y]-parameters.O),XY(:,1),XY(:,2));
        a = find(a>(parameters.bwHoles/2.5));
        if ~isempty(a)
            XY = XY(a:end,:);
            T = size(XY,1);
        end
    end
    
    
    %%%%%%%%%%%%%%%%%%%%%%%%%%%%%%%%%%%%%%%%%%%%%%%%%%%%%%%%%%%%%%%%%%%%%%
    % ^[QbgÅ ? s Èãâ~µÄ¢ét[ðJbg·éê
    %%%%%%%%%%%%%%%%%%%%%%%%%%%%%%%%%%%%%%%%%%%%%%%%%%%%%%%%%%%%%%%%%%%%%%
    if and(options.cutoff>0, ~strcmp(phase,'probe_test'))
                
        threshold = parameters.bwHoles/2.5;
        
        target = parameters.holes(1,:); target = repmat(target, T,1);
        d = mat2cell(XY-target, ones(T,1),2);
        d = cellfun(@norm, d);
        
        fin = 0;
        initframe = find(d<threshold,1,'first'); endframe = 1;
        if and(initframe~=T, T>options.cutoff*parameters.fps)
            while ~fin
                while ~fin
                    fin = or(and(d(initframe+endframe,:)<threshold,...
                        endframe>options.cutoff*parameters.fps),...
                        initframe+endframe>=T-1);
                    endframe = endframe + 1;
                end
                initframe = initframe + endframe - 1;
                endframe = 1;
            end
        else
            initframe = T+1;
        end
        endframe = initframe-1;
        XY(endframe:end,:) = [];
    end
      
        
   
    
    
    %% Network analysis
    if options.order{'Network','Analysis'}
        NetworkFeatures = feature_ntw_s_v0(parameters,NetworkFeatures,n,XY);
        disp('Network analysis (static node generation)....DONE.');
        NetworkFeatures = feature_ntw_d_v0(parameters,NetworkFeatures,n,XY);
        disp('Network analysis (dynamic node generation)...DONE.');
    else
        disp('Network analysis.............................SKIP.');
    end
    
    
    
    %% Strategy Indices
    if options.order{'Strategy','Analysis'}&&ismember(phase,{'training','retraining','reversal_training'})
        StrategyFeatures = feature_str_v1(parameters,options,LOG,StrategyFeatures,n,OnlineTracking,XY);
        disp('Searching strategy analysis..................DONE.');
    else
        disp('Searching strategy analysis..................SKIP.');
    end
    
    
    
    %% Conventional Indices
    if options.order{'Conventional','Analysis'}
        ConventionalFeatures =...
            feature_cnv_v2(parameters, options, ConventionalFeatures, n, phase, XY);
        disp('Conventional analysis........................DONE.');
    else
        disp('Conventional analysis........................SKIP.');
    end
    
    
    
    %% Extra Indices
    if options.order{'Extra','Analysis'}
        % ExtraFeatures = feature_extra_150812
        disp('Extra analysis...............................DONE.');
    else
        disp('Extra analysis...............................SKIP.');
    end
    
    
    
end
```
