## Extended Data 1 for "Scale space calibrates present and subsequent spatial learning in Barnes maze in mice": main_v0.m

```
clear;
close all
clc


%% set parameters
%%%%%%%%%%%%%%%%%%%%%%%%%%%%%%%%%%%%%%%%%%%%%%%%%%%%%%%%%%%%%%%%%%%%%%%%%%%
start_path = 'D:\documents\data';
options.maze_dia = 1; % BM type
options.offset = true; % offset; 32 in BM3, 41 in BM1
options.cutoff = 30; % cutoff = duration (s) till micro-movements around goals are supposed as goal entry
options.champs = []; % selet certain serial numbers to visualize whose local network
options.order = order_v0();
options.timeout = false;
options.view_network_graph = true;
%%%%%%%%%%%%%%%%%%%%%%%%%%%%%%%%%%%%%%%%%%%%%%%%%%%%%%%%%%%%%%%%%%%%%%%%%%%


%% load data
addpath(genpath(pwd))

switch options.maze_dia
    case 1
        options.step = 1; % x-axis view step (day)
        parameters = parameters_BM1; % make an object binding parameters
    case 3
        options.step = 2; % x-axis view step (day)
        parameters = parameters_BM3; % make an object binding parameters
end


if all(options.order.Load)
    
    [FileName,PathName] = uigetfile('*.mat', 'Select the LOG file', start_path);
    
    k = strfind(PathName,'\');
    PrjDir = PathName(1:k(end-1)-1);    
    TrnDir = dir(fullfile(PrjDir, 'training'));
    PrbDir = dir(fullfile(PrjDir, 'probe_test'));
    DatDir = [{TrnDir.name}, {PrbDir.name}];
    
    load(fullfile(PathName,FileName));    
    parameters.PrjDir = PrjDir;
    parameters.DatDir = DatDir;
    parameters.groups = unique(tbl.Group);
    parameters.SN = unique(str2double(tbl.SN));  
    
else    
    
    [FileName,PathName] = uigetfile('*.csv', 'Select the LOG file', start_path);
    
    k = strfind(PathName,'\');
    PrjDir = PathName(1:k(end-1)-1);    
    TrnDir = dir(fullfile(PrjDir, 'training'));
    PrbDir = dir(fullfile(PrjDir, 'probe_test'));
    DatDir = [{TrnDir.name}, {PrbDir.name}];
    
    LOG = readtable(fullfile(PathName,FileName), 'Delimiter',',', 'ReadVariableNames',1);
    
    N = size(LOG,1);
    
    varnames = LOG.Properties.VariableNames;
    if ~any(strcmp(varnames,'Reversal'))
        a = array2table(zeros(N,1)); a.Properties.VariableNames = {'Reversal'};
        b = find(strcmp(varnames,'Trial'));
        LOG = [LOG(:,1:b), a, LOG(:,b+1:end)];
    end    
    
    parameters.PrjDir = PrjDir;
    parameters.DatDir = DatDir;
    parameters.groups = unique(LOG.Group);
    parameters.SN = unique(LOG.SN);
    
    ConventionalFeatures = table;
    ConventionalFeatures.xy = cell(N,1);
    ConventionalFeatures.approach = cell(N,1);
    ConventionalFeatures{:,{'no_of_errors','latency','travel_distance'}} = nan(N,3);
    
    StrategyFeatures = table;
    StrategyFeatures{:,{'Random','Serial','Spatial','Perseveration'}} = zeros(N,4);
    
    NetworkFeatures = table;
    NetworkFeatures.xy_1 = cell(N,1);
    NetworkFeatures{:,{'n_1','l_1','m_1','CWS_1','rho_1','x_1','cl_1'}} = nan(N,7);
    NetworkFeatures{:,{'xy_2','A_2'}} = cell(N,2);
    NetworkFeatures{:,{'o_2','n_2','l_2','m_2','CWS_2','rho_2','x_2','cl_2'}} = nan(N,8);
    
    features = {'Sequence', 'travel_distance_prb', 'init_distrib'};
    ExtraFeatures = cell2table(cell(N,length(features))); % Extra
    ExtraFeatures.Properties.VariableNames = features;

    
    %% feature extraction
    [ConventionalFeatures, StrategyFeatures, NetworkFeatures, ExtraFeatures] =...
        FeatureExtraction_v0(parameters, options, LOG,...
        ConventionalFeatures, StrategyFeatures, NetworkFeatures, ExtraFeatures);
    
    tbl = [LOG,...
        ConventionalFeatures,...
        StrategyFeatures,...
        NetworkFeatures,...
        ExtraFeatures];    
    
end


%% postprocessing
varnames = tbl.Properties.VariableNames;
ind = zeros(length(varnames),1);
for i = 1:length(varnames)
    t = tbl.(i);
    if iscell(t)
        if all(cellfun(@ischar, t))
            if all(~cellfun(@isempty, cellfun(@str2num,t,'uniform',0)))
                tbl.(i) = cell2mat(cellfun(@str2num,t,'uniform',0));
            end
        elseif all(cellfun(@isnumeric, t))
            if all(cellfun(@length,t)==1)
                tbl.(i) = vertcat(t{:});
            end
        end
    end
end

tbl = sortrows(tbl, find(ismember(varnames,{'Day','Trial','SN'})));
if ~all(options.order.Load)
    phase = unique(tbl.Phase); phase = phase{:}; phase(phase=='_') = [];
    save([PathName,'tbl_',phase,'.mat'],'tbl')
end


%% summary conventional results
if options.order{'Conventional','Analysis'} % output data of conventional indices
    summary_cnv_v0(parameters,options,tbl);
else
    fprintf('\nAnalyze conventional indices......SKIP\n')
end


%% summary strategy results
if options.order{'Strategy','Analysis'}&&~ismember('probe_test',unique(tbl.Phase))
    summary_str_v0(parameters,options,tbl)
else
    fprintf('\nAnalyze strategy indices......SKIP\n')
end


%% summary network results
if options.order{'Network','Analysis'}
    summary_ntw_v0(parameters,options,tbl)
else
    fprintf('\nAnalyze network indices......SKIP\n')
end


%% summary extra results
if options.order{'Extra','Analysis'}
    summary_ext_v0(parameters,options,tbl)
else
    fprintf('\nAnalyze network indices......SKIP\n')
end


%%
fprintf('\nFinish.\n\n')
```
