## Extended Data 1 for "Scale space calibrates present and subsequent spatial learning in Barnes maze in mice": multiple_comparison_bonferoni_v0.m

```
clear;
close all
clc


%% training
src = 'F:\meta_learning\X-BM1\BM1\training\bundle\';
raw_file_name = 'NetworkScore.xls';
stats_file_name = 'ntw_stats.txt';
phase_ind = 0;


raw = readtable([src,raw_file_name]);
[groups,~,ig] = unique(raw.Group,'stable');
num_groups = length(groups);
C = nchoosek(1:num_groups,2);

if phase_ind==1    
    d = table(zeros(size(raw,1),1));
    d.Properties.VariableNames = {'Day'};
    raw = [d,raw];
end
feat = raw.Properties.VariableNames(4:end);
[Days,~,id] = unique(raw.Day); 
num_days = length(Days);


stats = readtable([src,stats_file_name]);
network_feat_ind = repmat(1:length(feat),1,length(Days));


for i = 1:length(feat)
    for j = 1:num_days        
        if stats.p((network_feat_ind==i)&(stats.Day==Days(j))')>=(.05/num_days)
            continue
        end
        for k = 1:size(C,1)
            tf = all([raw.Day==Days(j),ig==C(k,1)],2);
            A = raw{tf, feat{i}};
            tf = all([raw.Day==Days(j),ig==C(k,2)],2);
            B = raw{tf, feat{i}};
            [p,~,summary] = ranksum(A,B);
            if p<(0.05/(nchoosek(num_groups,2)))
                fprintf('%s, day %d, %s (%0.2f) vs %s (%0.2f), z = %0.2f, r = %0.2f, *.\n',...
                    feat{i},Days(j), groups{C(k,1)}, median(A),...
                    groups{C(k,2)}, median(B),...
                    summary.zval, summary.zval/sqrt(length(A)+length(B)))
            else
                fprintf('%s, day %d, %s (%0.2f) vs %s (%0.2f), z = %0.2f, r = %0.2f, n.s.\n',...
                    feat{i},Days(j), groups{C(k,1)}, median(A),...
                    groups{C(k,2)}, median(B),...
                    summary.zval, summary.zval/sqrt(length(A)+length(B)))
            end
            
        end        
    end
end

% writetable(tbl,[src,''])
```
