## Extended Data 1 for "Scale space calibrates present and subsequent spatial learning in Barnes maze in mice": adj2adjL.m

```
% Converts an adjacency graph representation to an adjacency list
% Valid for a general (directed, not simple) network model, but edge
% weights get lost in the conversion.
% INPUT: an adjacency matrix, NxN, N - # of nodes
% OUTPUT: cell structure for adjacency list: x{i_1}=[j_1,j_2 ...]
% GB, October 1, 2009

function L = adj2adjL(adj)

L=cell(length(adj),1);

for i=1:length(adj); L{i}=find(adj(i,:)>0); end
```
