## Extended Data 1 for "Scale space calibrates present and subsequent spatial learning in Barnes maze in mice": CCA.m

```
function [unions,nb,na] = CCA(individuals, criterion)


% criterion = 20; % unifying criterion
unions = individuals;

na = 1:size(unions,1);
nb = na;
pairs = nchoosek(nb, 2); % j[Nm[hÔÌSÄÌgÝí¹

X = 1;
ub = [];
uc = [];

while any(X)
    
    dist = unions(pairs(:,1), :) - unions(pairs(:,2), :);
    dist = mat2cell(dist, ones(size(dist,1),1), 2);
    dist = cellfun(@norm, dist);
    
    
    % m[hÔÌ£ªèlÈºÆÈém[hÌyAðo
    X = and(dist<=criterion, dist>0);
    Y = pairs(X,:);
    
    % ·ÅÉ¢¸ê©Ìm[hÉÜÜêÄ¢ém[hð
    [Cy,~,~] = unique(Y(:,2), 'stable');
    Y(ismember(Y(:,1),Cy),:) = [];
    
    g = unique(Y(:,1)); % m[hÌCfbNXðo
    NM = length(g); % ·ém[hÌ
    
    % m[hÌNX^[ÆdSÀWðZo
    for k = 1:NM
        
        M = Y(Y(:,1)==g(k), 2); % üÓm[h
        p = zeros(length(M)+1, 2);
        p(1,:) = unions(g(k), :);
        p(2:end,:) = unions(M, :);
        ua = [g(k); M];
        ub = [ub; ua]; % Sm[hÌCfbNX
        uc = [uc; repmat(k,length(ua),1)]; % Sm[hÌC³³ê½CfbNX
        
        nb(ua) = min(ua);
        na(ua) = k;
        unions(ua, :) = repmat(mean(p), [length(ua), 1]);
        
    end
    
end

[~, free, ~] = setxor(nb, ub); % ³êÄ¢È¢m[hÌCfbNX
[~, ia, ~] = unique(ub, 'last'); % ÅIIÈSm[hÌCfbNX
uc = uc(ia); % ÅIIÈSm[hÌC³³ê½CfbNX
free_indx = 1:size(individuals,1);
[~, free_indx, ~] = setxor(free_indx, uc); % Sm[hÌCfbNXð­
free_indx = free_indx(1:length(free));
na(free) = free_indx;
```
