## Extended Data 1 for "Scale space calibrates present and subsequent spatial learning in Barnes maze in mice": CCA_visualize.m

```
function [G,Gib] = CCA_visualize(lnode_xy, criterion)

% criterion = paramObj.thresh4CCA;

N = size(lnode_xy,1);

G = lnode_xy;
Gib = zeros(N,1);
stop = 0;
while ~stop

    Gc = mat2cell(G,ones(N,1),2);
    Gia = Gib;
    
    % calculate distance between local nodes
    D = cellfun(@(x) ones(N,1)*x-G, Gc, 'uniform',0);
    D = mat2cell(vertcat(D{:}), ones(N^2,1), 2);
    D = cellfun(@(x) norm(x)<=criterion, D);
    D = reshape(D, N, N);
    
    D(diag(ones(N,1))==1) = 0;
    [r,c] = find(D);    
    
    n = 1:N;
    Gib(~ismember(n,c)) = n(~ismember(n,c));
    
    for i = 1:N
        ind = [unique(c(c==i)); nonzeros(r(c==i))];
        G(ind,:) = repmat(mean(lnode_xy(ind,:),1), length(ind),1);
        Gib(ind) = i;
        c(ismember(c,ind)) = 0;
        r(ismember(r,ind)) = 0;
    end
    
    stop = all(Gia==Gib); % if locations of all stops/local nodes are converged, then stop

end
```
