## Extended Data 1 for "Scale space calibrates present and subsequent spatial learning in Barnes maze in mice": closeness.m

```
% Computes the closeness centrality for every vertex: 1/sum(dist to all other nodes)
% For disconnected graphs can use: sum_over_t(2^-d(i,t)), idea Dangalchev (2006)
% C(i)=sum(2.^(-d)) if graph is disconnected, but sum w/o d(i)
% INPUTs: graph representation (adjacency matrix nxn)
% OUTPUTs: vector of centralities, nx1
% Source: social networks literature
% Other routines used: simple_dijkstra.m 
% GB, Last updated: October 9, 2009

function C=closeness(adj)

C=zeros(length(adj),1);  % initialize closeness vector

for i=1:length(adj); C(i)=1/sum( simple_dijkstra(adj,i) ); end
```
