## Extended Data 1 for "Scale space calibrates present and subsequent spatial learning in Barnes maze in mice": clustering.m

```
function coeff = clustering(A, type)
% CLUSTERING   Calculates the clustering coefficient for each node from an adjacency matrix.
%   The clustering coefficient for each node in the graph is calculated
%   from the given adjacency matrix. If the type is given, then the
%   adjacency matrix is assumed to represent a graph of that type (either
%   directed or undirected). If the type is not given, the graph is assumed
%   to be undirected if the adjaceny matrix is symmetric, and directed
%   otherwise.
%       
%       USAGE:
%           coeff = clustering(A)
%           coeff = clustering(A, 'directed')
%           coeff = clustering(A, 'undirected')
%       
%       coeff
%           The column vector containing the clustering coefficient of each node.
%
%       A
%           The adjacency matrix.
%
%       type = 'directed'/'undirected'
%           The type of graph the adjacency matrix represents. If not
%           given, the graph is assumed to be undirected if it is
%           symmetric.

n = size(A,1);

if (nargin>1)
    if strcmp(type,'directed')
        digraph = true;
    elseif strcmp(type,'undirected')
        digraph = false;
    else
        error('Type must be either "directed" or "undirected"')
    end
else
    if all(all(A == A'))
        digraph = false;
    else
        digraph = true;
    end
end

if digraph
    c = sum((A^2) .* A, 2);
else
    c = sum((A^3) .* eye(n), 2);
end

% Calculate the out degree of the nodes
out = sum(A,2);

% Calculate the clustering coefficient
s = warning('off','MATLAB:divideByZero');
coeff = c ./ (out .* (out - 1));
warning(s);

% Remove the Inf's from the possible divide by 0
coeff(out == 0) = 0;
coeff(out == 1) = 0;
```
