## Extended Data 1 for "Scale space calibrates present and subsequent spatial learning in Barnes maze in mice": degrees.m

```
% Compute the total degree, in-degree and out-degree of a graph based on
% the adjacency matrix; should produce weighted degrees, if the input matrix is weighted
% INPUTS: adjacency matrix
% OUTPUTS: degree, indegree and outdegree sequences
% GB, Last Updated: October 2, 2009

function [deg,indeg,outdeg]=degrees(adj)

indeg = sum(adj);
outdeg = sum(adj');

if isdirected(adj)
  deg = indeg + outdeg; % total degree

else   % undirected graph: indeg=outdeg
  deg = indeg + diag(adj)';  % add self-loops twice, if any

end
```
