## Extended Data 1 for "Scale space calibrates present and subsequent spatial learning in Barnes maze in mice": diameter.m

```
% The longest shortest path between any two nodes nodes in the network
% INPUTS: adjacency matrix, adj
% OUTPUTS: network diameter, diam
% Other routines used: simple_dijkstra.m
% GB, Last updated: June 8, 2010

function diam = diameter(adj)

diam=0;
for i=1:size(adj,1)
    d=simple_dijkstra(adj,i);
    diam = max([max(d),diam]);
end
```
