## Extended Data 1 for "Scale space calibrates present and subsequent spatial learning in Barnes maze in mice": dijkstra.m

```
% INPUTS: adj - adjacency matrix, s - source node, target - target node
% OUTPUTS: distance, d and path, P (from s to target)
% Note: if target==[], then dist and P include all distances and paths from s
% Other routines used: adj2adjL.m, purge.m
% GB, Last Updated: Dec 22, 2009

function [dist,P]=dijkstra(adj,s,target)

% initialize distances ==========================
n=length(adj);            % number of nodes
adjL=adj2adjL(adj);       % list of neighbors

dist=inf(1,n);
dist(s)=0;

previous=[1:n; inf(1,n)]';  % {i: inf}, i=1:n, inf -> not assigned
S=cell(1,n); % shortest path sequence


Q=[1:n]; % all unvisited vertices, entire graph
while length(Q)>0 % while not empty
    % get min dist member among unvisited vertices
    [mindist,min_ind]=min(dist(Q));
    u=Q(min_ind);
    
    % termination condition - save source-u path
    S{u}=[];
    t=u;
    while not(isempty(find(previous(:,1)==t)))  % t in previous.keys():
        % insert u at the beginning of S
        S{u}=[t S{u}];
        t=previous(t,2);
    end
    if length(target)>0 & u==target
        dist=dist(u); P=S{u};
        return
    end            
    
    % =========================================
    Q=purge(Q,u);  % remove u from Q
    for v=1:length(adjL{u})   % across all neighbors of u
        v=adjL{u}(v);        
        alt=dist(u)+adj(u,v);
        if alt < dist(v)
            dist(v)=alt;
            previous(v,2)=u;
        end
    end
end

P=S;
```
