## Extended Data 1 for "Scale space calibrates present and subsequent spatial learning in Barnes maze in mice": dot_matrix_plot.m

```
% Draws the matrix as a column/row sorted square dot-matrix pattern
% INPUTs: adj - adjacency matrix representation of the graph
% OUTPUTs: plot
% Note: Change colors and marker types in lines 41, 48, 55 and 62
% Other routines used: degrees.m, sort_nodes_by_max_neighbor_degree.m,
%                      eigencentrality.m, newman_eigenvector_method.m,
%                      node_betweenness_faster.m
% GB, Last Updated: May 2, 2006, modified April 28, 2011

function [] = dot_matrix_plot(adj)

n = size(adj,1);
%markersize=ceil(n/50); % scale for plotting purposes
markersize=3;
[deg,indeg,outdeg]=degrees(adj);

Yd=sort_nodes_by_max_neighbor_degree(adj); % degree centrality
[betw, Yb] = sort(node_betweenness_faster(adj)); % node betweenness centrality
[EC,Yec]=sort(eigencentrality(adj)); % eigen-centrality

% sort by module
modules=newman_eigenvector_method(adj);
% sort modules by length
mL=zeros(1,length(modules));
for i=1:length(modules); mL(i)=length(modules{i}); end
[mS,Yms]=sort(mL);

% sort nodes by degree inside modules
Ym=[];
for mm=1:length(modules)
    module=modules{Yms(mm)};
    deg_module=deg(module);
    [ds,Yds]=sort(deg_module);
    module_sorted=module(Yds);
    for xx=1:length(module_sorted)
        Ym=[Ym module_sorted(xx)];
    end
end
    

set(gcf,'Color',[1 1 1])
subplot(2,2,1)
spy(adj(Yd,Yd),'ks',markersize)
xlabel('ordered by degree','FontWeight','bold')
axis([0 n 0 n]);
set(gca,'YDir','normal')
axis square

subplot(2,2,2)
spy(adj(Yb,Yb),'ks',markersize)
xlabel('ordered by betweenness','FontWeight','bold')
axis([0 n 0 n]);
set(gca,'YDir','normal')
axis square

subplot(2,2,3)
spy(adj(Yec,Yec),'ks',markersize)
xlabel('ordered by eigen-centrality','FontWeight','bold')
axis([0 n 0 n]);
set(gca,'YDir','normal')
axis square

subplot(2,2,4)
spy(adj(Ym,Ym),'ks',markersize)
xlabel('ordered by module','FontWeight','bold')
axis([0 n 0 n]);
set(gca,'YDir','normal')
axis square
```
