## Extended Data 1 for "Scale space calibrates present and subsequent spatial learning in Barnes maze in mice": draw_circ_graph.m

```
% Draw a circular graph with links and nodes in order of degree
% Strategy: position vertices in a regular n-polygon
% INPUTs: adj - adjacency matrix
% OUTPUTs: a figure
% Other routines used: degrees.m
% GB, February 21, 2006

function [] = draw_circ_graph(adj)

n = size(adj,1); % number of nodes
[degs,~,~]=degrees(adj);
[~, Y] = sort(degs);   % Y - sorted nodal indices
angl = 2*pi/n; % rotation angle

for k=1:n
  x(Y(k)) = real(exp(angl*(k-1)*i));
  y(Y(k)) = imag(exp(angl*(k-1)*i));
end

for k=1:n
  plot(x(k),y(k),'ko')
  text(1.1*x(k),1.1*y(k),strcat('v',num2str(k)));
  hold off; hold on;
end

edges=find(adj>0);
set(gcf,'Color',[1,1,1])

for e=1:length(edges)
    [ii,jj]=ind2sub([n,n],edges(e));
    line([x(ii) x(jj)],[y(ii) y(jj)],'Color','k');
    hold off; hold on;
end
axis off;
```
