## Extended Data 1 for "Scale space calibrates present and subsequent spatial learning in Barnes maze in mice": eigencentrality.m

```
% The ith component of the eigenvector corresponding to the greatest 
% eigenvalue gives the centrality score of the ith node in the network.
% INPUTs: adjacency matrix
% OUTPUTs: eigen(-centrality) vector
% GB, Last Updated: October 14, 2009

function x=eigencentrality(adj)

[V,D]=eig(adj);
[max_eig,ind]=max(diag(D));
x=V(:,ind);
```
