## Extended Data 1 for "Scale space calibrates present and subsequent spatial learning in Barnes maze in mice": el2geom.m

```
% Plot geometry based on extended edgelist
% INPUTS: extended edgelist el[i,:]=[n1 n2 m x1 y1 x2 y2]
% OUTPUTS: geometry plot, higher-weight links are thicker and lighter
% Note 1: m - edge weight; (x1,y1) are the Euclidean coordinates of n1, (x2,y2) - n2 resp. 
% Note 2: Easy to change colors and corresponding edge weight coloring
% Example input:
%       [el,p] = newmangastner(1000,0.5);
%       elnew = [];
%       for e=1:size(el,1)
%           elnew = [elnew; el(e,1), el(e,2), randi(8), p(el(e,1),1), p(el(e,1),2), p(el(e,2),1), p(el(e,2),2)];
%       end
%       el2geom(elnew)
% GB, last updated: april 24, 2007

function []=el2geom(el)

set(gcf,'Color',[1 1 1])
map=colormap('hot');

for i=1:size(el,1)
    
    % plot line between two nodes
    x1=el(i,4); y1=el(i,5);
    x2=el(i,6); y2=el(i,7);

    % edge weights ==============
    if el(i,3)<2
        color=map(8,:);
        linew=1;
    elseif el(i,3)>=2 & el(i,3)<3
        color=map(2*8,:);
        linew=2;
    elseif el(i,3)>=3 & el(i,3)<4
        color=map(3*8,:);
        linew=3;
    elseif el(i,3)>=4 & el(i,3)<5
        color=map(4*8,:);
        linew=4;
    elseif el(i,3)>=5 & el(i,3)<6
        color=map(5*8,:);
        linew=5;
    elseif el(i,3)>=6 & el(i,3)<7
        color=map(6*8,:);
        linew=6;
    elseif el(i,3)>=7 & el(i,3)<8
        color=map(7*8,:);
        linew=7;
    elseif el(i,3)>=8
        color=map(8*8,:);
        linew=8;
    end
    line([x1 x2],[y1 y2],'LineWidth',linew,'Color',color)

    hold off; hold on;

end
axis equal
```
