## Extended Data 1 for "Scale space calibrates present and subsequent spatial learning in Barnes maze in mice": isdirected.m

```
% Using the matrix transpose function
% INPUTS: adjacency matrix
% OUTPUTS: boolean variable
% GB, Last updated: October 1, 2009

function S=isdirected(adj)

S = true;
if adj==transpose(adj); S = false; end

% one-liner alternative: S=not(issymmetric(adj));
```
