## Extended Data 1 for "Scale space calibrates present and subsequent spatial learning in Barnes maze in mice": kmin_neighbors.m

```
% Finds the number of "kmin"-neighbors (k links away at a minimum) for every node
% If nodes are k-links away due to loops (so they appear as m-neighbours, m<k), they are not counted
% INPUTS: adjacency matrix, node index, k - number of links
% OUTPUTS: vector of "kmin"-neighbors indices
% GB, May 16, 2011

function kneigh = kmin_neighbors(adj,ind,k)

close_neighbors=[];

adjk = adj;
for i=1:k-1
  close_neighbors = [close_neighbors find(adjk(ind,:)>0)];
  adjk = adjk*adj;
end

kneigh = setdiff(find(adjk(ind,:)>0),[close_neighbors ind]);
```
