## Extended Data 1 for "Scale space calibrates present and subsequent spatial learning in Barnes maze in mice": kneighbors.m

```
% Finds the number of k-neighbors (k links away) for every node
% INPUTS: adjacency matrix, node index, k - number of links
% OUTPUTS: vector of k-neighbors indices
% GB, May 3, 2006

function kneigh = kneighbors(adj,ind,k)

adjk = adj;
for i=1:k-1; adjk = adjk*adj; end;

kneigh = find(adjk(ind,:)>0);
```
