## Extended Data 1 for "Scale space calibrates present and subsequent spatial learning in Barnes maze in mice": LOCGEN.m

```
%%%%%%%%%%%%%%%%%%%%%%%%%%%%%%%%%%%%%%%%%%%%%%%%%%%%%%%%%%%%%%%%
% Author            :   Qian Shi 
% Date Created      :   Nov 2008
% Date Updated      :   22nd Dec 2008
% Title             :   random topology generator with location constrains
% Copyright: This code is copyrighted by the ResiliNets group at the
% University of Kansas. Please contact {shiqian,jabbar,jpgs}@ittc.ku.edu
% for further details. https://wiki.ittc.ku.edu/resilinets/Topology_Modelling
%%%%%%%%%%%%%%%%%%%%%%%%%%%%%%%%%%%%%%%%%%%%%%%%%%%%%%%%%%%%%%%%
clc;clear all;
npoints=input('number of nodes:');
disp('LOCATION CONSTRAINED TOPOLOGY GENERATOR');
disp('available model:');
disp('1.Sprint corenetwork');
disp('2.Geant Network topology')
disp('3.Pure Random');
disp('4.Exponential');
disp('5.Locality');
disp('6.Wax-man');
disp('7.Doar-Leslie(Waxman2)');
disp('8.Barabasi-Albert');
adj_matr=zeros(npoints,npoints);
Topology_generator=input('model to use:');
%%%%%%%%%%%%%%%%%%%%%%%%%%%%%%%%%%%%%%%%%%%%%%%%%%%%
%Sample Topology of Sprint Network
%%%%%%%%%%%%%%%%%%%%%%%%%%%%%%%%%%%%%%%%%%%%%%%%%%%%
if (Topology_generator==1)
    npoints=27;
    nd_coord=[57.65,47.6;57.55,47.25;58.7,38.59;58.7,37.97;58,37.35;
          61.6,34.1;62.1,33.8;75.22,41.13;75,39.7;83.28,32.95;
          83.25,32.78;85.45,39.12;85.62,38.91;92.35,41.85;93.2,39.83;
          95.58,33.75;98.62,28.54;100.83,35.48;102.51,39.04;102.67,38.95;
          103,38.9;103.3,39.23;104.95,39.95;105.95,40.13;106,40.71;
          107.41,42.1;108.94,42.36;];
    disp('Sprint Topology');
    adj_matr=zeros(27,27);
    adj_matr(1,2) = 1;
    adj_matr(1,4) = 1;
    adj_matr(1,14) = 1;
    adj_matr(1,10) = 1;  
    adj_matr(1,15) = 1;
    adj_matr(2,8) = 1;
    adj_matr(2,22) = 1;
    adj_matr(2,15) = 1;     
    adj_matr(2,7) = 1;
    adj_matr(2,5) = 1;
    adj_matr(3,4) = 1;
    adj_matr(4,14) = 1;     
    adj_matr(4,25) = 1;
    adj_matr(4,23) = 1;
    adj_matr(2,4) = 1;
    adj_matr(4,5) = 1;     
    adj_matr(4,7) = 1;
    adj_matr(5,14) = 1;
    adj_matr(5,22) = 1;
    adj_matr(5,7) = 1;
    adj_matr(6,7) = 1;  
    adj_matr(7,14) = 1;
    adj_matr(7,10) = 1;
    adj_matr(7,25) = 1;
    adj_matr(7,22) = 1;     
    adj_matr(7,16) = 1;
    adj_matr(7,13) = 1;
    adj_matr(8,9) = 1;
    adj_matr(8,14) = 1;     
    adj_matr(8,25) = 1;
    adj_matr(8,10) = 1;
    adj_matr(8,16) = 1;
    adj_matr(8,13) = 1;     
    adj_matr(10,13) = 1;
    adj_matr(10,16) = 1;
    adj_matr(10,14) = 1;
    adj_matr(10,23) = 1;
    adj_matr(10,25) = 1;
    adj_matr(10,11) = 1;
    adj_matr(12,13) = 1;     
    adj_matr(13,14) = 1;
    adj_matr(13,15) = 1;
    adj_matr(13,16) = 1;
    adj_matr(13,17) = 1;     
    adj_matr(13,25) = 1;
    adj_matr(13,23) = 1;
    adj_matr(13,22) = 1;
    adj_matr(14,16) = 1;     
    adj_matr(14,17) = 1;
    adj_matr(14,22) = 1;
    adj_matr(14,25) = 1;
    adj_matr(14,26) = 1;
    adj_matr(15,25) = 1;
    adj_matr(16,17) = 1;
    adj_matr(16,18) = 1;     
    adj_matr(16,22) = 1;
    adj_matr(16,25) = 1;
    adj_matr(17,25) = 1;
    adj_matr(18,22) = 1;     
    adj_matr(19,20) = 1;
    adj_matr(20,21) = 1;
    adj_matr(21,22) = 1;
    adj_matr(22,23) = 1;     
    adj_matr(22,25) = 1;
    adj_matr(23,26) = 1;
    adj_matr(24,25) = 1;
    adj_matr(25,26) = 1;
    adj_matr(26,27) = 1;
     adj_matr1=adj_matr;
end

if (Topology_generator==2)
    npoints=34;
    nd_coord=[-21.5100,64.0900;-6.1500,53.2000;-0.1000,51.3000;-9.0800,38.4300;
           -3.4100,40.2400;2.2000,48.5200;4.5400,52.2200;4.2000,50.5000;
            6.0900,49.3600;10.4500,59.5500;12.3500,55.4000;18.0300,59.2000;
           24.5800, 60.1000;13.2100,52.2900;14.2600,50.0500;21.0000,52.1500;
           17.0700,48.0900;16.2000,48.1300;8.3200,47.2300;9.1200,45.2800;
           14.3100,46.0300;15.5800,45.4800;19.0500,47.3000;26.0600,44.2600;
           14.3100,35.5400;23.4300,37.5800;28.5000,40.5800;33.2200,35.1000;
           34.4600,32.0400;24.4500,59.2500;24.0600,56.5700;25.1900,54.4100;
           37.3500,55.4500; 23.20,42.45;];
       adj_matr(34,34)=0;
       disp('Geant Topology');
       adj_matr(1,11) = 1;
       adj_matr(2,3) = 1;
       adj_matr(3,8) = 1;  
       adj_matr(3,6) = 1;
       adj_matr(3,4) = 1;
       adj_matr(4,5) = 1;     
       adj_matr(5,6) = 1;
       adj_matr(5,20) = 1;
       adj_matr(5,19) = 1;
       adj_matr(6,9) = 1;     
       adj_matr(6,19) = 1;
       adj_matr(7,11) = 1;
       adj_matr(7,14) = 1;
       adj_matr(7,8) = 1;  
       adj_matr(7,25) = 1;
       adj_matr(9,14) = 1; 
       adj_matr(10,11) = 1;
       adj_matr(10,12) = 1;
       adj_matr(11,12) = 1;
       adj_matr(11,33) = 1;
       adj_matr(11,30) = 1;
       adj_matr(11,14) = 1;
       adj_matr(12,13) = 1;  
       adj_matr(14,15) = 1;
       adj_matr(14,16) = 1;     
       adj_matr(14,18) = 1;
       adj_matr(14,19) = 1;
       adj_matr(14,33) = 1;
       adj_matr(14,29) = 1;
       adj_matr(15,16) = 1;
       adj_matr(15,17) = 1; 
       adj_matr(16,32) = 1;
       adj_matr(17,23) = 1;
       adj_matr(17,18) = 1;
       adj_matr(18,20) = 1;
       adj_matr(18,21) = 1;
       adj_matr(18,26) = 1;
       adj_matr(19,20) = 1;
       adj_matr(20,25) = 1;
       adj_matr(20,26) = 1;     
       adj_matr(20,28) = 1;
       adj_matr(21,22) = 1;
       adj_matr(22,23) = 1;
       adj_matr(23,24) = 1;
       adj_matr(23,34) = 1;
       adj_matr(24,34) = 1;
       adj_matr(24,27) = 1;
       adj_matr(26,28) = 1;
       adj_matr(27,34) = 1;
       adj_matr(30,31) = 1;
       adj_matr(31,32) = 1;
       adj_matr1=adj_matr;
end
%%%%%%%%%%%%%%%%%%%%%%%%%%%%%%%%%%%%%%%%%%%%%%%%%%%%%%%%%%%%%%
%Calculating the real distance in Kilometre between node pairs 
%%%%%%%%%%%%%%%%%%%%%%%%%%%%%%%%%%%%%%%%%%%%%%%%%%%%%%%%%%%%%% 
if (Topology_generator~=1 &Topology_generator~=2)
    choice=input('Do you want to use 1:Sprint, 2:Geant or 2:your own coordinates? ');
    if( choice==1)
        nd_coord=[57.65,47.6;57.55,47.25;58.7,38.59;58.7,37.97;58,37.35;
          61.6,34.1;62.1,33.8;75.22,41.13;75,39.7;83.28,32.95;
          83.25,32.78;85.45,39.12;85.62,38.91;92.35,41.85;93.2,39.83;
          95.58,33.75;98.62,28.54;100.83,35.48;102.51,39.04;102.67,38.95;
          103,38.9;103.3,39.23;104.95,39.95;105.95,40.13;106,40.71;
          107.41,42.1;108.94,42.36;];
    end
    
    if (choice ==2)
        nd_coord=[-21.5100,64.0900;-6.1500,53.2000;-0.1000,51.3000;-9.0800,38.4300;
           -3.4100,40.2400;2.2000,48.5200;4.5400,52.2200;4.2000,50.5000;
            6.0900,49.3600;10.4500,59.5500;12.3500,55.4000;18.0300,59.2000;
           24.5800, 60.1000;13.2100,52.2900;14.2600,50.0500;21.0000,52.1500;
           17.0700,48.0900;16.2000,48.1300;8.3200,47.2300;9.1200,45.2800;
           14.3100,46.0300;15.5800,45.4800;19.0500,47.3000;26.0600,44.2600;
           14.3100,35.5400;23.4300,37.5800;28.5000,40.5800;33.2200,35.1000;
           34.4600,32.0400;24.4500,59.2500;24.0600,56.5700;25.1900,54.4100;
           37.3500,55.4500; 23.20,42.45;];
    end
    if(choice==3)
    nd_coord=input('Enter the Coordinate of nodes ');
    end
end

long_rep = repmat(nd_coord(:, 1), 1, npoints);
    la_rep = repmat(nd_coord(:, 2), 1, npoints);
  for x=1:1:npoints
         for y=1:1:npoints
           delta_la=la_rep(y,x)-la_rep(x,x);
           delta_long=long_rep(y,x)-long_rep(x,x);

             b=sin((delta_la*pi/180)/2)^2+cos(la_rep(x,x)*pi/180)*cos((la_rep(y,x))*pi/180)*(sin((delta_long*pi/180)/2))^2;
             c=atan2(sqrt(b),sqrt(1-b));

          dist_matr(y,x)=6371*2*c;
         end
     end
    dist_matr=sparse(triu(dist_matr));
    
if (Topology_generator==3)
    disp('using Pure random generator:');
   
        
    alpha = input ('link probability:');    
    % generate the adjacency matrix
    runi = sprand(dist_matr);
    adj_matr = (runi>0) & (runi < alpha)
    adj_matr1=full(adj_matr);

end

if (Topology_generator==4)
    disp('Using exponential generator');
    alpha = input ('link probability:');
    prob_matr = alpha*spfun('exp', ...
                   -dist_matr./(max(max(dist_matr))-dist_matr));
               % generate the adjacency matrix
               runi = sprand(dist_matr);
               adj_matr = (runi>0) & (runi < prob_matr)
                adj_matr1=full(adj_matr);
end

if (Topology_generator==5)
    disp('Locality generator');
    alpha = input('parameter for short link probability :'); % parameter for the local link probability
    beta = input('parameter for long link probability :'); % parameter for the long link probability
    gama = input('distance threshhold:');
    %generate the Adjacency matrix
    runi = sprand(dist_matr);
    adj_matr = ((runi>0) & (runi < alpha) & (dist_matr <= gama))|((runi>0) & (runi < beta) & (dist_matr >= gama))
     adj_matr1=full(adj_matr);
end

if (Topology_generator==6)
    disp('using Waxman generator');
    alpha = input('parameter for link probability a:'); % parameter for the link probability
    beta = input('parameter for link probability b:'); % parameter for the link probability

 % create the matrix of probabilities
    prob_matr = alpha*spfun('exp', ...
                   -dist_matr./(beta*max(max(dist_matr))));
 % generate the adjacency matrix
    runi = sprand(dist_matr);
    adj_matr = (runi>0) & (runi < prob_matr)
     adj_matr1=full(adj_matr);
end

if (Topology_generator==7)
    disp('using DoarLeslie generator');
    alpha = input('parameter for link probability a:'); % parameter for the link probability
    beta = input('parameter for link probability b:'); % parameter for the link probability
    n = npoints;
    e = input('desired degree e :');
    k = input('constant to satisfy the probability');
    prob_matr = (alpha*k*e/n)*spfun('exp', ...
                   -dist_matr./(beta*max(max(dist_matr))));
    % generate the adjacency matrix
    runi = sprand(dist_matr);
    adj_matr = (runi>0) & (runi < prob_matr)
     adj_matr1=full(adj_matr);
end

if (Topology_generator==8)
    disp('Using Albert-Barabasi (power-law) generator(to be finished)');
     disp('using Waxman to connect backbone Network ');
    alpha = input('parameter for link probability a:'); % parameter for the link probability
    beta = input('parameter for link probability b:'); % parameter for the link probability
    region=input('Network region')
   prob_matr = alpha*spfun('exp', ...
                   -dist_matr./(beta*max(max(dist_matr))));
    runi = sprand(dist_matr);
    adj_matr = (runi>0) & (runi < prob_matr);
    %%%%%%%%%%%%%%%%%%%%%%%%%
    %adding power law nodes
    %%%%%%%%%%%%%%%%%%%%%%%%%%%
    new_nodes = input('number of nodes to add: ');
    region = input('region to put new nodes:');      
    new_nd_coord=rand(new_nodes,1)*region/2+region/2;
    new_nd_coord=rand(new_nodes,2)*region/2+region/4;
    nd_coord_new=[nd_coord;new_nd_coord];
    degree=zeros(1,npoints);
    total_degree=0;
     for x=2:1:npoints
                 for y=1:1:x-1
                     if (adj_matr(y,x)==1) 
                         degree(1,y)=degree(1,y)+1;
                         degree(1,x)=degree(1,x)+1;
                     end
                 end
             end    
    total_degree=sum(degree);
%%%%%%%%%%%%%%%%%%%%%%%%%%%
%calculating waxman distance for new nodes
%%%%%%%%%%%%%%%%%%%%%%%%%%%    
    long_rep = nd_coord_new(:, 1);
    la_rep = nd_coord_new(:, 2);
   for xx=npoints+1:1:npoints+new_nodes; 
       new_distm_matr=0;
       for zz=1:1:xx-1   
           delta_la=la_rep(xx,1)-la_rep(zz,1);
           delta_long=long_rep(xx,1)-long_rep(zz,1);

             b=sin((delta_la*pi/180)/2)^2+cos(la_rep(zz,1)*pi/180)*cos((la_rep(zz,1))*pi/180)*(sin((delta_long*pi/180)/2))^2;
             c=atan2(sqrt(b),sqrt(1-b));

          new_dist_matr(1,zz)=6371*2*c;
     
     end
    new_dist_matr
    max_dist=0;abc=0;
 [max_dist,abc]=max(new_dist_matr);
 new_prob_matr=alpha*exp(-new_dist_matr./(beta*max_dist));  
          total_new_prob_matr=sum(new_prob_matr);
  
        preferencial_prob=new_prob_matr.*degree./(total_new_prob_matr*total_degree)
% generate the adjacency matrix
[max,connect]=max(preferencial_prob)
adj_matr(connect,xx)=1;degree(1,xx)=1;degree(1,connect)=degree(1,connect)+1;
total_degree=sum(degree);
   end
end
%%%%%%%%%%%%%%%%%%%%%%%%%%%%%%%%%%%%%%%%%%%
%Calculate cost of network generated
%%%%%%%%%%%%%%%%%%%%%%%%%%%%%%%%%%%%%%%%%%%
fix_cost=input('input the fix cost for each link');
cost_per_mile=input('input the coast per mile for links')
Cost=0;
for x=2:1:npoints
    for y=1:1:x-1        
        if (adj_matr(y,x)==1) 
            Cost=Cost+fix_cost+cost_per_mile*dist_matr(y,x);
        end  
    end
end
disp('Cost of Network:')
disp(Cost);


 figure(1);
 clf;
 hold on;
 plot(nd_coord(:, 1), nd_coord(:, 2),'ok');
 gplot(adj_matr, nd_coord,'k');
 title('Topology of Network');
 hold off;
%%%%%%%%%%%%%%%%%%%%%%%%%%%%
%write adjacency matrix and node coordinates to files for challenge model
%%%%%%%%%%%%%%%%%%%%%%%%%%%%
%%%%%%%%%%%%%%%%%%%%%%%%%%%%%
 dlmwrite('adjacency_matrix.txt',adj_matr1,'delimiter', '\t');
 dlmwrite('node_coordinate.txt',nd_coord,'delimiter', '\t');
```
