## Extended Data 1 for "Scale space calibrates present and subsequent spatial learning in Barnes maze in mice": make_graph_v0.m

```
clear;
close all
clc


load('G:\nVista\OF\X.mat')
X = X.history{7}{:,end};
[~,~,X] = unique(X);
X(X==1) = [];
X([false;diff(X)==0]) = [];

X = [X(1:end-1),X(2:end)];

LinearInd = sub2ind([max(unique(X)),max(unique(X))],X(:,1),X(:,2));
X = zeros(max(unique(X)));
X(LinearInd) = 1;
X(1,:) = []; X(:,1) = [];

colormap jet
G = graph(X,'upper');
p = plot(G);
p.MarkerSize = 10;
p.EdgeColor = 'k';
p.LineWidth = 2;
p.NodeLabel = {};

% h.EdgeColor = 'k';

G.Nodes.NodeColors = degree(G);
% G.Nodes.NodeColors = centrality(G,'betweenness');
p.NodeCData = G.Nodes.NodeColors;
colorbar

axis off
axis square

set(gca,'FontSize',24)
```
