## Extended Data 1 for "Scale space calibrates present and subsequent spatial learning in Barnes maze in mice": newmangastner.m

```
% Implements the Newman-Gastner model for spatially distributed networks
% Source: Newman, Gastner, "Shape and efficiency in spatial distribution networks"
% Note 1: minimize: wij = dij + beta x (dj0)
% Note 2: easy to change to input point coordinates, instead of generate randomly
% Inputs: n - number of points/nodes, beta - parameter [0,1], plt - 'on'/'off'
% Outputs: graph (edgelist), point coordinates and plot [optional]
% GB, Last Updated: May 23, 2007

function [el,points]=newmangastner(n,beta,plt)

% create random point coordinates
points = zeros(n-1,2);
rad = rand(1,n-1);
theta = 2*pi*rand(1,n-1);
for i=1:n-1; points(i,:) = rad(i)*[cos(theta(i)),sin(theta(i))]; end
points=[[0 0]; points];               % add zero

% order points in closeness to 0
for i=1:n; normp(i)=norm(points(i,:)); end
[normps,ind]=sort(normp);
points=points(ind,:);

% calculate distances between all points (can also use pdist)
L=zeros(n);
for i=1:n
    for j=i+1:n
        L(i,j)=norm(points(i,:)-points(j,:));
        L(j,i)=L(i,j);
    end
    L(i,i)=Inf;
end

if nargin>2 & strcmp(plt,'on')  % if plot is 'on'
  set(gcf,'Color',[1 1 1])
  plot(points(:,1),points(:,2),'.','Color',[1,1,1])
  hold off; hold on;
  axis off
end

% connect points consequtively
el=[];
for i=2:n
    % current node is "i"
    w=L(i,1:i-1)+beta*normps(1:i-1);
    [minv,minj]=min(w);
    el=[el; i minj 1];
    
    if nargin>2 & strcmp(plt,'on')  % if plot is 'on'
      line([points(i,1) points(minj,1)],[points(i,2) points(minj,2)],'Color',[0.5,0.5,0.5])
      hold off; hold on;
      drawnow
    end
end
```
