## Extended Data 1 for "Scale space calibrates present and subsequent spatial learning in Barnes maze in mice": node_betweenness_faster.m

```
% Betweenness centrality measure: number of shortest paths running though a vertex 
% Compute for all vertices, using Dijkstra's algorithm, using 'number of shortest paths through a node' definition
% Note: Valid for a general (connected) graph.
% INPUTS: adjacency (distances) matrix (nxn)
% OUTPUTS: betweeness vector for all vertices (nx1)
% Other routines used: dijkstra.m
% GB, December 22, 2009

function betw = node_betweenness_faster(adj)

n = length(adj);
spaths=inf(n,n);
adjk = adj;

% calculate number of shortest paths
for k=1:n-1
  
  for i=1:n
    for j=1:n
      
      if adjk(i,j)>0; spaths(i,j)=min([spaths(i,j) adjk(i,j)]); end
    
    end
  end
  
  adjk=adjk*adj;
 
end


betw = zeros(1,n);  
for i=1:n
    [dist,P]=dijkstra(adj,i,[]);
    for j=1:n
      
        if dist(j)<=1; continue; end   % either i=j or i,j are 1 edge apart
        betw(P{j}(2:dist(j))) = betw(P{j}(2:dist(j))) + 1/spaths(i,j);
        
    end
end

betw=betw/nchoosek(n,2);   % further normalize by the number of all node pairs
```
