## Extended Data 1 for "Scale space calibrates present and subsequent spatial learning in Barnes maze in mice": node_betweenness_slow.m

```
% Betweenness centrality measure: number of shortest paths running though a

% vertex. Compute for all vertices.

% Note: Valid for a general graph. Using 'number of shortest paths through a node' definition

% INPUTS: adjacency (distances) matrix (nxn)

% OUTPUTS: betweeness vector for all vertices (nx1)

%

% GB, October 13, 2009


function betw = node_betweenness_slow(adj)


n = numnodes(adj);

spaths=inf(n,n);


% calculate number of shortest paths

for k=1:n-1

  adjk=adj^k;

  for i=1:n

    for j=1:n

      if adjk(i,j)>0

        spaths(i,j)=min([spaths(i,j) adjk(i,j)]);

      end

    end

  end

end


betw = zeros(1,n);  

for i=1:n

    [J_st,route_st,J,route]=shortest_pathDP(adj,i,i,n);

    for j=1:n

        if i==j

            continue

        end

        %[J_st,route_st]=shortest_pathDP(adj,i,j,n);

        [J_ji,step_ind] = min(J(:,j));

        route_ji = [j, route(step_ind,j).path];

        betw(route_ji(2:length(route_ji)-1)) = betw(route_ji(2:length(route_ji)-1)) + 1/spaths(j,i);

    end

end


betw=betw/nchoosek(n,2);
```
