## Extended Data 1 for "Scale space calibrates present and subsequent spatial learning in Barnes maze in mice": numnodes.m

```
% Returns the number of nodes, given an adjacency list
% also works for an adjacency matrix
% INPUTs: adjacency list: {i:j_1,j_2 ..}
% OUTPUTs: number of nodes
% GB, February 19, 2006

function n = numnodes(L)

n = length(L);
```
