## Extended Data 1 for "Scale space calibrates present and subsequent spatial learning in Barnes maze in mice": pdf_cdf_rank.m

```
% Compute the pdf, cdf and rank distributions for a sequence of values
% INPUTS: sequence of values: x, size 1xn, 'plot' - 'on' or 'off'
% OUTPUTS: pdf, cdf and rank distribution values
% Note: pdf = frequency, cdf = cumulative frequency, rank = log-log scale of the sorted sequence
% GB, Last Updated: June 27, 2007

function [xpdf,ypdf,xcdf,ycdf,logk,logx]=pdf_cdf_rank(x,plt)

xx=unique(x);

bin = 100; % arbitrary, change if xx has few values
if length(xx)<100; bin = 10; end;


for ii=1:numel(xx)
  xcdf(ii) = xx(ii); ycdf(ii) = length(find(x<=xx(ii)))/numel(x);
  
  % how many x's fall in the interval [xx(ii)-0.5*numel(xx)/bin,xx(ii)+0.5*numel(xx)/bin]
  xpdf(ii) = xx(ii); ypdf(ii) = length(find(abs(xx(ii)-x)<=0.5*numel(xx)/bin))/numel(x); 
end

x=-sort(-x);
logk=log(1:length(x));
logx=log(x);

if strcmp(plt,'on')
     set(gcf,'color',[1,1,1])
     subplot(1,3,1)
     plot(xpdf,ypdf,'k.')
     title('pdf')
     axis('tight')
     subplot(1,3,2)
     plot(xcdf,ycdf,'k.')
     title('cdf')
     axis('tight')
     subplot(1,3,3)
     plot(logk,logx,'k.')
     title('rank')
     axis('tight')
end
```
