## Extended Data 1 for "Scale space calibrates present and subsequent spatial learning in Barnes maze in mice": purge.m

```
% Removes a subset from a set, but preserves order of elements
% Similar to setdiff - which sorts the elements
% INPUTs: original set A, subset to remove B
% OUTPUTs: set Anew = A-B
% GB, Last updated: October 12, 2009

function Anew = purge(A,B)

Anew = [];
for a=1:numel(A); 
  if isempty(find(B==A(a))); Anew=[Anew, A(a)]; end
end
```
