## Extended Data 1 for "Scale space calibrates present and subsequent spatial learning in Barnes maze in mice": radial_plot.m

```
% Plots nodes radially out from a given center. Equidistant nodes
% have the same radius, but different angles. Works best as a quick
% visualization for trees, or very sparse graphs. 
% Note 1: No spring-energy method implemented.
% Note 2: If a center node is not specified, the nodes are ordered by
% sum of neighbor degrees, and the node with highest sum is plotted
% in the center.
% Note 3: The graph has to be connected.
% Note 4: To change the color scheme, modify lines: 84, 90, 96 and 110
% Inputs: adjacency matrix, and center node (optional)
% Outputs: plot
% Other routines used: sort_nodes_by_sum_neighbor_degrees.m,
%                      adj2adjL.m, diameter.m, kmin_neighbors.m
% Last updated: May 19, 2011, GB

function [] = radial_plot(adj,i0)

d=diameter(adj);
if d==Inf; fprintf('this graph is not connected, choose a connected graph\n'); return; end

if nargin==1; I=sort_nodes_by_sum_neighbor_degrees(adj); i0=I(1); end;

L=adj2adjL(adj);

circles={};   % per circle
circle{i0}=0; % per node

for k=1:d
  kneigh = kmin_neighbors(adj,i0,k);
  if not(isempty(kneigh));
    circles{k}=kneigh;
    for nei=1:length(kneigh); circle{kneigh(nei)}=k; end
  end;
end
  
for i=1:size(adj,1); descendants{i} = 0; end;

for k=length(circles)-1:-1:1
  
  for i=1:length(circles{k})
    node = circles{k}(i);
    neighs=L{node};
    
    for nei=1:length(L{node})
      nei = L{node}(nei);
      if sum(find(circles{k+1}==nei))>0; descendants{node}=descendants{node}+descendants{nei}+1; end
    end
  end
  
end

descendants{i0}=descendants{i0} + length(L{i0});
for i=1:length(L{i0}); descendants{i0}=descendants{i0} + descendants{L{i0}(i)}; end;

start_angle{i0}=0;
range{i0}=2*pi;

queue=[i0]; visited=[];

while not(isempty(queue))
  
  node=queue(1); queue=queue(2:length(queue)); visited=[visited node];
  
  % select neighbours that have not been visited
  neigh=[];
  for ll=1:length(L{node})
    if sum(find(visited==L{node}(ll)))==0 & sum(find(queue==L{node}(ll)))==0; neigh=[neigh L{node}(ll)]; end;
  end
  
  if isempty(neigh); continue; end;
  
  range{neigh(1)} = range{node}*(descendants{neigh(1)}+1)/descendants{node};
  start_angle{neigh(1)}=start_angle{node};
  if sum(find(queue==neigh(1)))==0; queue=[queue neigh(1)]; end;

  for j=2:length(neigh)
    range{neigh(j)} = range{node}*(descendants{neigh(j)}+1)/descendants{node};
    start_angle{neigh(j)} = start_angle{neigh(j-1)} + range{neigh(j-1)};
    if sum(find(queue==neigh(j)))==0; queue=[queue neigh(j)]; end;
  end
    
end

set(gcf,'Color',[1 1 1],'Colormap',hot);
map=colormap('hot');

% plot with the angle and radius (circle) information
for i=1:size(adj,1)
  x = circle{i}*cos(start_angle{i}+range{i}/2);
  y = circle{i}*sin(start_angle{i}+range{i}/2);
  plot(x,y,'.','Color',map(mod(5*circle{i},length(map))+1,:)); hold off; hold on;
end

xs = cos([0:1:30]*2*pi/30);
ys = sin([0:1:30]*2*pi/30);
for c=1:length(circles)
  plot(c*xs,c*ys,':','Color',[244/255,244/255,244/255]);
  hold off; hold on;
end

% add the links between nodes
for i=1:size(adj,1)
  for j=i+1:size(adj,1)
    if adj(i,j)>0
      xi=circle{i}*cos(start_angle{i}+range{i}/2);
      yi=circle{i}*sin(start_angle{i}+range{i}/2);
      
      xj=circle{j}*cos(start_angle{j}+range{j}/2);
      yj=circle{j}*sin(start_angle{j}+range{j}/2);
      
      line([xi xj],[yi yj],'Color',map(mod(5*circle{i},length(map))+1,:))
    end
  end
end
    
axis tight
axis off
```
