## Extended Data 1 for "Scale space calibrates present and subsequent spatial learning in Barnes maze in mice": shortest_pathDP.m

```
% Shortest path algorithm using Dynamic Programming
% Valid for directed/undirected network
% Disclaimer: if links have weights, they are treated as distances
% INPUTs: L - (cost/path lengths matrix), s - (start/source node), t - (end/destination node)
% OUTPUTS: 
%       route - sequence of nodes on optimal path, at current stage
%       ex: route(i,j) - best route from j to destination in (i) steps
%       Jo - optimal cost function (path length)
% Source: D. P. Bertsekas, Dynamic Programming and Optimal Control, Athena Scientific, 2005 (3rd edition)
% GB, Last Updated: March 9, 2006

function [J_st,route_st,J,route]=shortest_pathDP(L,s,t,steps)

n = size(L,2);

L(find(L==0))=Inf;  % make all zero distances equal to infinity

for i=1:n
  J(steps,i) = L(i,t); 
  route(steps,i).path = [t];
end

% find min for every i: Jk(i)=min_j(L(i,j)+Jk+1(j))
for p=1:steps-1
  k=steps-p; % recurse backwards
  
  for i=1:n
    %fprintf('stage %2i, node %2i \n',k,i)
    [J(k,i),ind_j] =  min(L(i,:)+J(k+1,:));
    route(k,i).path = [ind_j, route(k+1,ind_j).path];
  end
  
end

[J_st,step_ind] = min(J(:,s));
route_st = [s, route(step_ind,s).path];
J=J(sort(1:n,'descend'),:);
route=route(sort(1:n,'descend'),:);
```
