## Extended Data 1 for "Scale space calibrates present and subsequent spatial learning in Barnes maze in mice": simple_dijkstra.m

```
% Implements a simple version of the Dijkstra shortest path algorithm
% Returns the distance from a single vertex to all others, doesn't save the path
% INPUTS: adjacency matrix (adj), start node (s)
% OUTPUTS: shortest path length from start node to all other nodes
% Note: works with a weighted/directed matrix
% GB, Last Updated: December 13, 2004

function d = simple_dijkstra(adj,s)

n=length(adj);
d = inf*ones(1,n); % distance s-all nodes
d(s) = 0;    % s-s distance
T = 1:n;    % node set with shortest paths not found

while not(isempty(T))
    [dmin,ind] = min(d(T));
    for j=1:length(T)
        if adj(T(ind),T(j))>0 & d(T(j))>d(T(ind))+adj(T(ind),T(j))
            d(T(j))=d(T(ind))+adj(T(ind),T(j));
        end
    end 
    T = setdiff(T,T(ind));
end
```
