## Extended Data 1 for "Scale space calibrates present and subsequent spatial learning in Barnes maze in mice": sort_nodes_by_max_neighbor_degree.m

```
% Sort nodes by degree, and where there's equality, by maximum neighbor degree
% Ideas from Guo, Chen, Zhou, "Fingerprint for Network Topologies"
% INPUTS: adjacency matrix, 0s and 1s
% OUTPUTS: sorted sequence from 1 to n, where n is the number of rows/cols of the adjacency
% Other routines used: degrees.m, kneighbors.m

function I=sort_nodes_by_max_neighbor_degree(adj)

[deg,~,~]=degrees(adj); % compute all degrees, use "deg" assuming symmetry

degmat=zeros(size(adj,1),2);  % a nx2 matrix

for x=1:size(adj,1)   % across all nodes
    degmat(x,1)=deg(x);
    if deg(x)==0;  degmat(x,2)=0; continue; end
    nei_inds=kneighbors(adj,x,1);
    degmat(x,2)=max(deg(nei_inds));
    
end

[sortmat,I]=sortrows(degmat);
```
