## Extended Data 1 for "Scale space calibrates present and subsequent spatial learning in Barnes maze in mice": sort_nodes_by_sum_neighbor_degrees.m

```
% Sort nodes by degree, and where there's equality, by sum of neighbor degrees and then neighbors' neighbors degree and so on
% Ideas from s-max algorithm by Li et al 2005 "Towards a theory of scale-free graphs"
% and Guo, Chen, Zhou, "Fingerprint for Network Topologies"
% INPUTS: adjacency matrix, 0s and 1s
% OUTPUTS: sorted sequence from 1 to n, where n is the number of rows/cols of the adjacency
% Other routines used: degrees.m, kneighbors.m

function I=sort_nodes_by_sum_neighbor_degrees(adj)

[deg,~,~]=degrees(adj); % compute all degrees, use "deg" assuming symmetry

degmat=zeros(size(adj,1),size(adj,1));  % nxn matrix [index i,sum(deg of neighbors of index i, <=j links away)]

for x=1:size(adj,1)   % across all nodes
    degmat(x,1)=deg(x);
    
    % across all threshold distances, 2 to n-1 [should really be 2 to diameter, if the diameter is known]
    for kk=2:size(adj,1); degmat(x,kk)=sum( deg(kneighbors(adj,x,kk-1)) ); end

end

[sortmat,I]=sortrows(degmat);
I=I(end:-1:1);
```
