## Extended Data 1 for "Scale space calibrates present and subsequent spatial learning in Barnes maze in mice": order_v0.m

```
function requests = order_v0()


f = figure('Position',[250,250,500,300]);


pm = uicontrol(f,'Style','popupmenu',...
    'String',{'Run all','Load'},...
    'Value',1,'Position',[35,270,100,20]);

t = uicontrol(f,'Style','text',...
    'String','Select run all processing including feature extraction or load data table',...
    'Position',[150,260,200,30]);


bg = uibuttongroup(f,'Title','Your requests',...
    'Position',[.07, .15, .9, .7]);

tb = uicontrol(f,'Style','togglebutton',...
    'String','OK',...
    'Value',0,'Position',[370 60 100 30]);


xywh_colname = [126,200,80,30];

cnv = uicontrol(f,'Style','text',...
    'String','Conventional',...
    'Position',xywh_colname);
str = uicontrol(f,'Style','text',...
    'String','Strategy',...
    'Position',xywh_colname+[90,0,0,0]);
ntw = uicontrol(f,'Style','text',...
    'String','Network',...
    'Position',xywh_colname+[2*90,0,0,0]);
ext = uicontrol(f,'Style','text',...
    'String','Extra',...
    'Position',xywh_colname+[3*90,0,0,0]);


dy = 25;
xywh_rawname = [70,178,50,20];
analysis = uicontrol(f,'Style','text',...
    'String','Analysis',...
    'Position',xywh_rawname);
output = uicontrol(f,'Style','text',...
    'String','Output',...
    'Position',xywh_rawname-[0,dy,0,0]);
stats = uicontrol(f,'Style','text',...
    'String','Stats',...
    'Position',xywh_rawname-[0,2*dy,0,0]);
view = uicontrol(f,'Style','text',...
    'String','View',...
    'Position',xywh_rawname-[0,3*dy,0,0]);


xywh_chkbox = [148,136,20,20];
a1 = uicontrol(bg,'Style','checkbox',...
    'Value',1,'Position',xywh_chkbox-[0,0,0,0]);
a2 = uicontrol(bg,'Style','checkbox',...
    'Value',1,'Position',xywh_chkbox+[79,0,0,0]-[0,0,0,0]);
a3 = uicontrol(bg,'Style','checkbox',...
    'Value',1,'Position',xywh_chkbox+[79+89,0,0,0]-[0,0,0,0]);
a4 = uicontrol(bg,'Style','checkbox',...
    'Value',0,'Position',xywh_chkbox+[79+89+83,0,0,0]-[0,0,0,0]);


b1 = uicontrol(bg,'Style','checkbox',...
    'Value',1,'Position',xywh_chkbox-[0,dy,0,0]);
b2 = uicontrol(bg,'Style','checkbox',...
    'Value',1,'Position',xywh_chkbox+[79,0,0,0]-[0,dy,0,0]);
b3 = uicontrol(bg,'Style','checkbox',...
    'Value',1,'Position',xywh_chkbox+[79+89,0,0,0]-[0,dy,0,0]);
b4 = uicontrol(bg,'Style','checkbox',...
    'Value',0,'Position',xywh_chkbox+[79+89+83,0,0,0]-[0,dy,0,0]);


c1 = uicontrol(bg,'Style','checkbox',...
    'Value',1,'Position',xywh_chkbox-[0,2*dy,0,0]);
c2 = uicontrol(bg,'Style','checkbox',...
    'Value',1,'Position',xywh_chkbox+[79,0,0,0]-[0,2*dy,0,0]);
c3 = uicontrol(bg,'Style','checkbox',...
    'Value',1,'Position',xywh_chkbox+[79+89,0,0,0]-[0,2*dy,0,0]);
c4 = uicontrol(bg,'Style','checkbox',...
    'Value',0,'Position',xywh_chkbox+[79+89+83,0,0,0]-[0,2*dy,0,0]);


d1 = uicontrol(bg,'Style','checkbox',...
    'Value',1,'Position',xywh_chkbox-[0,3*dy,0,0]);
d2 = uicontrol(bg,'Style','checkbox',...
    'Value',1,'Position',xywh_chkbox+[79,0,0,0]-[0,3*dy,0,0]);
d3 = uicontrol(bg,'Style','checkbox',...
    'Value',1,'Position',xywh_chkbox+[79+89,0,0,0]-[0,3*dy,0,0]);
d4 = uicontrol(bg,'Style','checkbox',...
    'Value',0,'Position',xywh_chkbox+[79+89+83,0,0,0]-[0,3*dy,0,0]);


a = 0;
while ~a
    drawnow
    a = tb.Value;
end


rownames = {'Conventional','Strategy','Network','Extra'};
colnames = {'Load','Analysis','Output','Stats','View'};
requests = [a1.Value, a2.Value, a3.Value, a4.Value;...
    b1.Value, b2.Value, b3.Value, b4.Value;...
    c1.Value, c2.Value, c3.Value, c4.Value;...
    d1.Value, d2.Value, d3.Value, d4.Value]';
if pm.Value==1
    x = zeros(size(requests,1),1);    
else
    x = ones(size(requests,1),1);
end
requests = [x,requests];

requests = array2table(requests);
requests.Properties.VariableNames = colnames;
requests.Properties.RowNames = rownames;


close all
```
