## Extended Data 1 for "Scale space calibrates present and subsequent spatial learning in Barnes maze in mice": parameters_BM1.m

```
function parameters = parameters_BM1()

% field parameters
WIDTH = 980; % mm
nHoles = 12; % Ì
nMidNodes = 12; % Ôm[hÌ
nNodes = nHoles + nMidNodes + 1; % m[hÌ
PossibleLinks = nchoosek(1:nNodes, 2);
nPossibleLinks = 2*nchoosek(nNodes, 2);


% recording parameters
fps = 20;
pix = 500; % pixel 
spr = WIDTH/pix; % óÔðxCx mm / pix
DIA = WIDTH/spr; % tB[hÌ¼aCpix\¦
RAD = DIA/2; % tB[hÌ¼aCpix\¦
O = [RAD,RAD]; % tB[hÌSÀW
dia = 40/spr; % Ì¼aCpix\¦
diaInitPos = 130/spr;
rad = dia/2; % Ì¼aCpix\¦
Edge2Holes = 90/spr; % tB[hÌOü©çÌ~üÜÅÌÅZ£Cpix\¦
Center2Holes = RAD - (Edge2Holes + rad); % tB[hÌS©çÌSÜÅÌ£Cpix\¦
Center2Holes_H = Center2Holes + rad; % tB[hÌS©çÌãÜÅÌ£Cpix\¦
Center2Holes_L = Center2Holes - rad; % tB[hÌS©çÌºÜÅÌ£Cpix\¦
diaStartBox = 130/spr; %

thresh4CCA = 80/spr;


%eÌÀWðvZ
%{
GA = (250, 250)©ç(1, 1)ÖÌxNgðC
30¸Âñ]³¹½ÌxNgÆÀWÆå«³ðÝè
x' = x*cos(theta) - y*sin(theta)
y' = x*sin(theta) + y*cos(theta)
%}
theta = (0:30:330)';
holes = zeros(length(theta), 2);
holes(:,1) = ((cosd(theta)-sind(theta))*...
    (Center2Holes/sqrt(2))+RAD)';
holes(:,2) = ((sind(theta)+cosd(theta))*...
    (Center2Holes/sqrt(2))+RAD)';
bwHoles = sqrt(Center2Holes^2-(Center2Holes*sind(75))^2) * 2; % distance b/w holes


%Ôm[hÌÀWðvZ
%{
GA = (250, 250)©ç(1, 1)ÖÌxNgðC
30¸Âñ]³¹½ÌxNgÆÀWÆå«³ðÝè
x' = x*cos(theta) - y*sin(theta)
y' = x*sin(theta) + y*cos(theta)
%}
CenterEdge = (Center2Holes-(bwHoles/2))/2; % Sm[hÌ~ü
Center2MidNodes = CenterEdge + CenterEdge/2;
theta = (0:30:330)';
middle_nodes = zeros(length(theta), 2);
middle_nodes(:, 1) = ((cosd(theta)-sind(theta))*...
    (Center2MidNodes/sqrt(2))+RAD)';
middle_nodes(:, 2) = ((sind(theta)+cosd(theta))*...
    (Center2MidNodes/sqrt(2))+RAD)';
bwMidNodes = sqrt(Center2MidNodes^2-(Center2MidNodes*sind(75))^2) * 2;


% make structure of parameter
c = who;
params_cell = cell(length(c),1);
for i = 1:length(c)
    if strcmp(c{i}, 'c')
        continue
    end
    params_cell(i) = {eval(c{i})};
end
parameters = cell2struct(params_cell, c, 1);
```
