## Extended Data 1 for "Scale space calibrates present and subsequent spatial learning in Barnes maze in mice": parameters_BM3.m

```
function parameters = parameters_BM3()

% field parameters
WIDTH = 3000; % mm
nHoles = 12; % Ì
nMidNodes = 12; % Ôm[hÌ
nNodes = nHoles + nMidNodes + 1; % m[hÌ
PossibleLinks = nchoosek(1:nNodes, 2);
nPossibleLinks = 2*nchoosek(nNodes, 2);

fps = 20;
pix = 400; % pixel 
spr = WIDTH / pix; % óÔðxCx mm / pix
DIA = WIDTH / spr; % tB[hÌ¼aCpix\¦
RAD = DIA / 2; % tB[hÌ¼aCpix\¦
O = [RAD,RAD]; % tB[hÌSÀW
dia = 160 / spr; % Ì¼aCpix\¦
diaInitPos = 130 / spr;
rad = dia / 2; % Ì¼aCpix\¦
Edge2Holes = 125 / spr; % tB[hÌOü©çÌ~üÜÅÌÅZ£Cpix\¦
Center2Holes = RAD - (Edge2Holes + rad); % tB[hÌS©çÌSÜÅÌ£Cpix\¦
Center2Holes_H = Center2Holes + rad; % tB[hÌS©çÌãÜÅÌ£Cpix\¦
Center2Holes_L = Center2Holes - rad; % tB[hÌS©çÌºÜÅÌ£Cpix\¦

thresh4CCA = 30; % default = 80/2/spr


%eÌÀWðvZ
%{
LabVIEW Ì¨ÌovOÅo³ê½eÌÀWðvñèÅ\[g
%}
listing = dir('*.txt');
listing = {listing.name}; listing = listing(end);
holes = dlmread(listing{:},'\t');
holes = circshift(holes,[-4,0]);
holes = holes(:,2:3);
w = repmat([0.86,0.86,0.86,0.87,...
    0.89,0.9,0.9,0.9,...
    0.89,0.88,0.87,0.86],2,1)';
p = (holes-O).*w+O;

bwHoles = sqrt(Center2Holes^2-(Center2Holes*sind(75))^2) * 2; % distance b/w holes

direct = [0,0, pix,0, pix,pix, 0,pix];

d = repmat(holes,1,4)-repmat(direct,size(holes,1),1);
d = mat2cell(d, ones(size(holes,1),1), ones(4,1).*2);
d = cellfun(@norm, d);
[~,GoalHoleNum] = min(d);


CenterEdge = (Center2Holes-(bwHoles/2))/2; % Sm[hÌ~ü

%Ôm[hÌÀWðvZ
%{
center ©ç holes Ì a {ÌÊuÉ¶¬³ê½m[h
%}
a = 0.5;
middle_nodes = a.*holes;
middle_nodes = middle_nodes + ones(size(middle_nodes,1),1)*(O-mean(middle_nodes));


holes = p;


% make structure of parameter
c = who;
params_cell = cell(length(c),1);
for i = 1:length(c)
    if strcmp(c{i}, 'c')
        continue
    end
    params_cell(i) = {eval(c{i})};
end
parameters = cell2struct(params_cell, c, 1);
```
