## Extended Data 1 for "Scale space calibrates present and subsequent spatial learning in Barnes maze in mice": README.md

# Barnes Maze Analysis

## Requirement
* The codes were developed and tested on MATLAB R2018 and R2020 on Windows 10.
* Excel 2016 or later will be required if you want Excel outputs.


## Usage
(setup) Download, unzip, move, or clone the source code folder ***BM-main.zip***
(setup) A data folder should have the structure of ***demo***.


1. Edit ***set parameters*** section of ***main_v0.m***.
2. Run ***main_v0.m***. A popup window will appear.
3. Fill checkboxes. Do NOT choose ***Extra***. Then select the OK button.
    * ***Analysis***; fill checkboxes at columns of behavioral features, ***Conventional***, ***Strategy*** and ***Network*** that you want to analyze.
    * ***Output***; fill checkboxes at columns of behavioral features, ***Conventional***, ***Strategy*** and ***Network*** that you want to get spreadsheet output. If your PC does not have Excel, do NOT fill any checkboxes, as the codes write data to Excel files.
    * ***Stats***; fill checkboxes at columns of behavioral features, ***Conventional***, ***Strategy*** and ***Network*** that you want to perform statistical tests. If your PC does not have Excel, do NOT fill the checkbox of ***Conventional***, as the codes write data to Excel files.
    * ***View***; fill checkboxes at columns of behavioral features, ***Conventional***, ***Strategy*** and ***Network*** that you want to get figures.

 
## Author

* name
* affiliation
* e-mail
 
## License
A part of codes in the ***network_analysis*** folder are modified version of [MIT_network_toolbox](https://github.com/cliffordlab/MIT_network_toolbox.git)

A part of codes in the ***stats*** folder are modified version of [Mixed (Between/Within Subjects) ANOVA Version 1.1.0.0](https://uk.mathworks.com/matlabcentral/fileexchange/27080-mixed-between-within-subjects-anova), and [sto (ver. 0.30)](https://rnpsychology.org/sto/)
