## Extended Data 1 for "Scale space calibrates present and subsequent spatial learning in Barnes maze in mice": mixed_between_within_anova.m

```
function [SSQs, DFs, MSQs, Fs, Ps]=mixed_between_within_anova(X,suppress_output)
% simple function for mixed (between- and within-subjects) ANOVA
% 
% Based loosely on BWAOV2 (http://www.mathworks.com/matlabcentral/fileexchange/5579-bwaov2) by Antonio Trujillo-Ortiz
%  (insofar as I used that function to figure out the basic equations, as it is apparently very hard to find documentation
%  on mixed-model ANOVAs on the Internet). However, the code is all original.
% 
% The major advantage of this function over the existing BWAOV2 is that it corrects a bug that occurs when the groups
%  have unequal numbers of subjects, as pointed out in the Matlab Central File Exchange by Jaewon Hwang. The code is also,
%  at least in my opinion, much cleaner.
% 
% At present this function only supports mixed models with a single between-subjects factor and a single within-subjects
%  (repeated measures) factor, each with as many levels as you like. I would be happy to add more bells and whistles in
%  future editions, such as the ability to define multiple factors, apply Mauchly's test and add in non-sphericity 
%  corrections when necessary, etc. I'm a better programmer than I am a statistician, though, so if anyone out there would 
%  like to lend a hand with the math (e.g., feed me the equations for these features), I'd be more than happy to implement 
%  them. Email matthew DOT r DOT johnson AT aya DOT yale DOT edu
% 
% Also feel free to modify this file for your own purposes and upload your changes to the Matlab Central File Exchange if 
%  you like.
% 
% I have checked this function against the example data in David M. Lane's HyperStat online textbook, which is the same 
%  data that breaks BWAOV2 (http://davidmlane.com/hyperstat/questions/Chapter_14.html, question 6). I have also checked it
%  against SPSS and gotten identical results. However, I haven't tested every possible case so bugs may remain. Use at 
%  your own risk. If you find bugs and let me know, I'm happy to try to fix them.
% 
% ===============
%      USAGE
% ===============
% 
% Inputs:
% 
% X: design matrix with four columns (future versions may allow different input configurations)
%     - first column  (i.e., X(:,1)) : all dependent variable values
%     - second column (i.e., X(:,2)) : between-subjects factor (e.g., subject group) level codes (ranging from 1:L where 
%         L is the # of levels for the between-subjects factor)
%     - third column  (i.e., X(:,3)) : within-subjects factor (e.g., condition/task) level codes (ranging from 1:L where 
%         L is the # of levels for the within-subjects factor)
%     - fourth column (i.e., X(:,4)) : subject codes (ranging from 1:N where N is the total number of subjects)
% 
% suppress_output: defaults to 0 (meaning it displays the ANOVA table as output). If you don't want to display the table,
%  just pass in a non-zero value
% 
% Outputs:
% 
% SSQs, DFs, MSQs, Fs, Ps : Sum of squares, degrees of freedom, mean squares, F-values, P-values. All the same values 
%  that are shown in the table if you choose to display it. All will be cell matrices. Values within will be in the same 
%  order that they are shown in the output table.
% 
% Enjoy! -MJ


if nargin < 1, 
   error('No input');
end;

if nargin < 2 || isempty(suppress_output)
    suppress_output=0;
end

all_dvs=X(:,1);
all_bs_labels=X(:,2);
all_ws_labels=X(:,3);
all_subj_labels=X(:,4);

bs_levels=sort(unique(all_bs_labels));
ws_levels=sort(unique(all_ws_labels));
subj_levels=sort(unique(all_subj_labels));
n_bs_levels=length(bs_levels);
n_ws_levels=length(ws_levels);
n_subjects=length(subj_levels);

if any(bs_levels(:)~=(1:n_bs_levels)') || any(ws_levels(:)~=(1:n_ws_levels)') || any(subj_levels(:)~=(1:n_subjects)')
    error('Levels of factors (including subject labels) must be numbered 1:L (where L is the # of levels for that factor');
end
for i=1:n_bs_levels
    for j=1:n_ws_levels
        this_cell_inds=find(all_bs_labels==i & all_ws_labels==j);
        if isempty(this_cell_inds)
            error('At least one empty cell found');
        end
        n_subs_per_cell(i,j)=length(this_cell_inds); %#ok<AGROW>
        cell_totals(i,j)=sum(all_dvs(this_cell_inds)); %#ok<AGROW>
    end
    
    if any(n_subs_per_cell(i,:)~=n_subs_per_cell(i,1))
        error('At least one subject missing at least one repeated measure (or is possibly entered more than once)');
        %technically this is not a failsafe check, as it could be that subject ! has only conditions A & B,
        %  whereas subject 2 has only condition C, which still gives equal values for all the conditions.
        %  We'll double-check for that circumstance below
    end
end

for k=1:n_subjects
    this_subj_inds=find(all_subj_labels==k);
    if length(this_subj_inds)~=n_ws_levels
        %our second check for this issue
        error('At least one subject missing at least one repeated measure (or is possibly entered more than once)');
    end
    subj_totals(k)=sum(all_dvs(this_subj_inds)); %#ok<AGROW>
end

correction_term = sum(all_dvs)^2 / length(all_dvs);
SStot = sum(all_dvs.^2) - correction_term;
%don't really need this for calculations, but can uncomment if we want to print
% DFtot = length(all_dvs) - 1; %total degrees of freedom

%subject "factor" (i.e. differences in subject means)
SSsub = sum(subj_totals .^ 2)/n_ws_levels - correction_term;

%between-subjects factor
SStmp=[];
for i=1:n_bs_levels
    SStmp(i)=(sum(cell_totals(i,:))^2) / sum(n_subs_per_cell(i,:)); %#ok<AGROW>
end
SSbs    = sum(SStmp) - correction_term;
DFbs    = n_bs_levels - 1;
MSbs    = SSbs / DFbs;
%error terms for between-subjects factor
ERRbs   = SSsub - SSbs;
DFERRbs = n_subjects - n_bs_levels;
MSERRbs = ERRbs / DFERRbs;

%correction with harmonic mean of cell sizes if cell sizes are not all equal
n_subs_hm=harmmean(n_subs_per_cell(:));
cell_totals_hm = (cell_totals ./ n_subs_per_cell) * n_subs_hm;
correction_term_hm = sum(cell_totals_hm(:))^2 / (n_subs_hm * n_bs_levels * n_ws_levels);
n_subs_per_cell_hm = ones(n_bs_levels,n_ws_levels) * n_subs_hm;

%within-subjects factor
SStmp=[];
for j=1:n_ws_levels
    SStmp(j)=(sum(cell_totals_hm(:,j))^2) ./ sum(n_subs_per_cell_hm(:,j)); %#ok<AGROW>
end
SSws  = sum(SStmp) - correction_term_hm;
DFws  = n_ws_levels - 1;
MSws  = SSws / DFws;

%uncorrected version of within-subjects factor for calculating interaction
SStmp=[];
for j=1:n_ws_levels
    SStmp(j)=(sum(cell_totals(:,j))^2) ./ sum(n_subs_per_cell(:,j)); %#ok<AGROW>
end
SSws_unc = sum(SStmp) - correction_term;

%interaction of between-subjects and within-subjects factor
SStmp = sum((cell_totals(:) .^ 2) ./ n_subs_per_cell(:));
SSint = SStmp - SSbs - SSws_unc - correction_term;
DFint = DFbs * DFws;
MSint = SSint / DFint;

%error terms (for both within-subjects factor and interaction)
ERRws   = SStot - SSbs - ERRbs - SSws_unc - SSint;
DFERRws = DFERRbs * DFws;
MSERRws = ERRws / DFERRws;

%F-values
Fbs  = MSbs  / MSERRbs;
Fws  = MSws  / MSERRws;
Fint = MSint / MSERRws;

%P-values
Pbs  = 1-fcdf(Fbs, DFbs, DFERRbs);
Pws  = 1-fcdf(Fws, DFws, DFERRws);
Pint = 1-fcdf(Fint,DFint,DFERRws);

if ~suppress_output
    disp(' ');
    disp(' ');
    fprintf(1,'     Source                          SSq        df       MSq         F         p        \n');
    fprintf(1,'-------------------------------------------------------------------------------------\n');
    fprintf(1,'     Between-subjects factor   %9.3f %9i %9.3f %9.3f %9.7f\n',SSbs,DFbs,MSbs,Fbs,Pbs);
    fprintf(1,'     Between-subjects error    %9.3f %9i %9.3f\n',ERRbs,DFERRbs,MSERRbs);
    fprintf(1,'     Within-subjects factor    %9.3f %9i %9.3f %9.3f %9.7f\n',SSws,DFws,MSws,Fws,Pws);
    fprintf(1,'     Within x between int.     %9.3f %9i %9.3f %9.3f %9.7f\n',SSint,DFint,MSint,Fint,Pint);
    fprintf(1,'     Within-subjects error     %9.3f %9i %9.3f\n',ERRws,DFERRws,MSERRws);
    disp(' ');
    disp(' ');
end

SSQs = { SSbs; ERRbs;   SSws; SSint; ERRws   };
DFs  = { DFbs; DFERRbs; DFws; DFint; DFERRws };
MSQs = { MSbs; MSERRbs; MSws; MSint; MSERRws };
Fs   = { Fbs;  [];      Fws;  Fint;  []      };
Ps   = { Pbs;  [];      Pws;  Pint;  []      };
```
