## Extended Data 1 for "Scale space calibrates present and subsequent spatial learning in Barnes maze in mice": anova_1wm.m

```
function [Report,Level,M,N,F,p,MS,df,ES] = anova_1wm(Data)
% 
% anova_1wm:  対応のある1要因分散分析（データは集計表形式）
% 
% [Report,Level,M,N,F,p,MS,df,ES] = anova_1wm(Data)
%    Data    対象となるデータ（集計表形式）
%    Report  分散分析表
%    Level   水準番号の一覧
%    M       各水準の平均値
%    N       サンプル数（e.g.,参加者数）
%    F       F値
%    p       p値
%    MS      平均平方
%    df      自由度
%    ES      効果量（pEta^2:偏イータ2乗）
%
% 入力データ（Data）は、サンプル数×水準数の行列です。
% すなわち、同一行内の数値は対応したデータです。
% 4名の参加者による3水準のデータの例：
%       Data = [
%          12.3   18.9   15.5
%          15.1   21.2   16.1
%           9.9   23.5   12.0
%          11.3   20.9   14.7
%       ];
% 
% see also:   anova_1w   anova_1b   tukey_1wm
% 
% (2015/09/10, by R. NIIMI)

Report = ['']; Level = []; M = []; N = []; F = []; p = []; MS = []; df = []; ES = [];

if ~(size(Data,1)>1 & size(Data,2)>1)
    disp('  [anova_1wm]:  Data must be a matrix.');
    return;
end

[N NL] = size(Data);
Nall = N * NL;
M = mean(Data);
X = sum(sum(Data))^2 / (N * NL);
AS = sum(sum(Data.^2));
A = sum(sum(Data).^2) / N;
S = sum(sum(Data').^2) / NL;

SS1 = A - X;
SSs = S - X;
SSe = AS - A - S + X;
SSt = AS - X;
df1 = NL-1;
dfs = N-1;
dfe = df1*dfs;
dft = Nall-1;
MS1 = SS1/df1;
MSs = SSs/dfs;
MSe = SSe/dfe;
F1 = MS1/MSe;
Fs = MSs/MSe;
p1 = p4F(F1,df1,dfe);
ps = p4F(Fs,dfs,dfe);
ES = SS1/(SS1+SSe);

temp = [
    length('SS') length('df') length('MS') length('F') length('p') length('pEta^2');
    length(num2str(SS1)) length(num2str(df1)) length(num2str(MS1)) length(num2str(F1)) length(num2str(p1)) length(num2str(ES));
    length(num2str(SSs)) length(num2str(dfs)) length(num2str(MSs)) length(num2str(Fs)) length(num2str(ps)) 0 ;
    length(num2str(SSe)) length(num2str(dfe)) length(num2str(MSe)) 0 0 0 ;
    length(num2str(SSt)) length(num2str(dft)) 0 0 0 0 ;
];
temp = [max(temp); max(temp); max(temp); max(temp); max(temp);] - temp;
sp = '   ';
Report(2,:) = ['   Source' sp 'SS' blanks(temp(1,1)) sp 'df' blanks(temp(1,2)) sp 'MS' blanks(temp(1,3)) sp 'F' blanks(temp(1,4)) sp 'p' blanks(temp(1,5)) sp 'pEta^2' blanks(temp(1,6))];
Report(1,:) = ['   ' char(ones(1,size(Report,2)-3)*45)];
Report(3,:) = Report(1,:);
Report(4,:) = ['   Factor' sp blanks(temp(2,1)) num2str(SS1) sp blanks(temp(2,2)) num2str(df1) sp blanks(temp(2,3)) num2str(MS1) sp blanks(temp(2,4)) num2str(F1) sp blanks(temp(2,5)) num2str(p1) sp blanks(temp(2,6)) num2str(ES)];
Report(5,:) = ['   Sample' sp blanks(temp(3,1)) num2str(SSs) sp blanks(temp(3,2)) num2str(dfs) sp blanks(temp(3,3)) num2str(MSs) sp blanks(temp(3,4)) num2str(Fs) sp blanks(temp(3,5)) num2str(ps) sp blanks(temp(3,6))];
Report(6,:) = ['   Error ' sp blanks(temp(4,1)) num2str(SSe) sp blanks(temp(4,2)) num2str(dfe) sp blanks(temp(4,3)) num2str(MSe) sp blanks(sum(temp(4,4:6))) sp sp];
Report(7,:) = Report(1,:);
Report(8,:) = ['   Total ' sp blanks(temp(5,1)) num2str(SSt) sp blanks(temp(5,2)) num2str(dft) sp blanks(sum(temp(5,3:6))) sp sp sp];
Report(9,:) = Report(1,:);

SS = [SS1 SSs SSe SSt];
df = [df1 dfs dfe dft];
MS = [MS1 MSs MSe];
F = [F1 Fs];
p = [p1 ps];
Level = [1:NL];
```
