## Extended Data 1 for "Scale space calibrates present and subsequent spatial learning in Barnes maze in mice": anova_2bb.m

```
function [Report,Level,M,N,F,p,MS,df,ES] = anova_2bb(Data)
% 
% anova_2bb:  2要因分散分析、2要因とも対応のない場合
% 
% [Report,Level,M,N,F,p,MS,df,ES] = anova_2bb(Data)
%    Data    対象となるデータ（3列の行列）
%    Report  分散分析表
%    Level   水準番号の一覧（セルアレイ）
%    M       各条件の平均値
%    N       サンプル数
%    F       F値
%    p       p値
%    MS      平均平方
%    df      自由度
%    ES      効果量（pEta^2:偏イータ2乗）
% 
% 入力データ（Data）は、1列目に要因Aの水準番号、2列目に要因Bの
% 水準番号、そして3列目に測定値が書かれた行列。すなわち、1行が
% 1サンプルです。
% 要因A（2水準）×要因B（3水準）のデータの例：
%       Data = [
%          1    1    12.3
%          1    1    15.1
%          1    1     9.9
%          1    1    11.3
%          1    2    18.9
%          1    2    21.2
%          1    2    23.5
%          1    3    15.5
%          1    3    16.1
%          1    3    12.0
%          1    3    14.7
%          0    1    19.3
%          0    1    20.7
%          0    1    19.0
%          0    1    18.1
%          0    1    17.9
%          0    2    21.8
%          0    2    32.6
%          0    2    27.4
%          0    2    29.0
%          0    3    23.8
%          0    3    23.2
%          0    3    16.7
%          0    3    22.1
%       ];
% 水準番号は整数（負も可）。MやNは、水準番号の小さい順になっていま
% す。なお、各水準のサンプル数が等しくない場合には、非加重平均値
% を用いてSSを計算します。
% 
% see also:   anova_2ww   anova_2bw   sme_2bb   tukey_2bb
% 
% (2015/09/10, by R. NIIMI)

Report = ['']; Level = []; M = []; N = []; F = []; p = []; MS = []; df = []; ES = [];

if ~(size(Data,1)>1 & size(Data,2)==3)
    disp('  [anova_2bb]:  Data must be a N by 3 matrix.');
    return;
end

Nall = size(Data,1);
Level{1} = min(Data(:,1));
for k=min(Data(:,1))+1:max(Data(:,1))
   if sum(Data(:,1)==k) > 0
       Level{1} = [Level{1} k];
   end
end
Level{2} = min(Data(:,2));
for k=min(Data(:,2))+1:max(Data(:,2))
   if sum(Data(:,2)==k) > 0
       Level{2} = [Level{2} k];
   end
end
Dat = [];
NL = [length(Level{1}) length(Level{2})]; % 水準数
for k=1:NL(1)
    for l=1:NL(2)
        I = find(Data(:,1)==Level{1}(k) & Data(:,2)==Level{2}(l));
        N(k,l) = length(I);
        M(k,l) = sum(Data(I,3));
%        Dat(:,l,k) = Data(I,3); % データを3次元（S*B*A）の集計表形式にする
    end
end

ABS = sum(Data(:,3).^2);
AB = sum(sum((M.^2)./N));
M = M./N; % 各セルの平均値
G = sum(sum(M));
X = G*G / (NL(1)*NL(2));
A = NL(2) * sum(mean(M').^2);
B = NL(1) * sum(mean(M).^2);

% AB = sum(sum(M.^2));
Ntil = NL(1)*NL(2) / sum(sum(1./N));

SS = [
    Ntil*(A-X); % SSa
    Ntil*(B-X); % SSb
    Ntil*(sum(sum(M.^2))-A-B+X); % SSab
    ABS - AB; % SSwc
];

dfa = NL(1) - 1;
dfb = NL(2) - 1;
dfab = dfa * dfb;
dfwc = Nall - NL(1)*NL(2);
df = [dfa dfb dfab dfwc]';

MS = SS ./ df;

F = [
    MS(1) / MS(4); % Fa
    MS(2) / MS(4); % Fb
    MS(3) / MS(4); % Fab
];

p = [
    p4F(F(1),dfa,dfwc); % pa
    p4F(F(2),dfb,dfwc); % pb
    p4F(F(3),dfab,dfwc); % pab
];

ES = SS(1:3) ./ (SS(1:3) + SS(4));

Report = [
    'Source    ';
    'A         ';
    'B         ';
    'A*B       ';
    'Error     ';
];
sp = char(ones(5,3)*32);
Report = [Report sp strvcat('SS',num2str(SS)) sp strvcat('df',num2str(df)) sp strvcat('MS',num2str(MS))];
Report = [Report sp strvcat('F',num2str(F),' ')];
Report = [Report sp strvcat('p',num2str(p),' ')];
Report = [Report sp strvcat('pEta^2',num2str(ES),' ')];
temp = char(ones(1,size(Report,2))*45);
Report = [temp; Report(1,:); temp; Report(2:5,:); temp];
Report = [char(ones(8,3)*32) Report];

SS = SS';
df = df';
MS = MS';
F = F';
p = p';
ES = ES';
```
