## Extended Data 1 for "Scale space calibrates present and subsequent spatial learning in Barnes maze in mice": anova_2bw.m

```
function [Report,Level,M,N,F,p,MS,df,ES] = anova_2bw(Data)
% 
% anova_2bw:  2要因分散分析、混合計画の場合
% [Report,Level,M,N,F,p,MS,df,ES] = anova_2bw(Data)
%    Data    対象となるデータ（3列の行列）
%    Report  分散分析表
%    Level   水準番号の一覧（セルアレイ）
%    M       各条件の平均値
%    N       サンプル数
%    F       F値
%    p       p値
%    MS      平均平方
%    df      自由度
%    ES      効果量（pEta^2:偏イータ2乗）
% 
% 1つの要因に対応があり、もう1つの要因に対応がない場合（混合
% 計画）の2要因分散分析を行います。
% 
% 入力データ（Data）は、1列目に対応のない要因（要因A）の水準
% 番号、2列目に対応のある要因（要因B）の水準番号、そして3列目
% に測定値が書かれた行列。すなわち、1行が1サンプルです。
% 要因A（3水準）×要因B（2水準）のデータの例：
% （11名の参加者が要因Aの各水準に4名、4名、3名のように割り当て
% られ、各参加者は要因Bの2水準の両方の実験に参加した、と想定）
%       Data = [
%          1    1    12.3
%          1    1    15.1
%          1    1     9.9
%          1    1    11.3
%          2    1    18.9
%          2    1    21.2
%          2    1    23.5
%          2    1    15.5
%          3    1    16.1
%          3    1    12.0
%          3    1    14.7
%          1    0    19.3
%          1    0    20.7
%          1    0    19.0
%          1    0    21.8
%          2    0    32.6
%          2    0    27.4
%          2    0    29.0
%          2    0    23.8
%          3    0    23.2
%          3    0    16.7
%          3    0    22.1
%       ];
% 要因Bの各水準における行の順序は対応させて下さい。すなわち、上例の
% 1行目と12行目、2行目と13行目は同一の参加者による（対応した）デー
% タです。一方、1行目と5行目は対応していないデータです。
% 
% 水準番号は整数（負も可）。MやNは、この水準番号の小さい順になってい
% ます。なお、各水準のサンプル数が等しくない場合には、非加重平均値
% を用いてSSを計算します。
% 
% see also:   anova_2bb   anova_2ww   sme_2bw   tukey_2bw
% 
% (2015/09/10, by R. NIIMI)

Report = ['']; Level = []; M = []; N = []; F = []; p = []; MS = []; df = []; ES = [];

if ~(size(Data,1)>1 & size(Data,2)==3)
    disp('  [anova_2bw]:  Data must be a N by 3 matrix.');
    return;
end

Nall = size(Data,1);
Level{1} = min(Data(:,1));
for k=min(Data(:,1))+1:max(Data(:,1))
   if sum(Data(:,1)==k) > 0
       Level{1} = [Level{1} k];
   end
end
Level{2} = min(Data(:,2));
for k=min(Data(:,2))+1:max(Data(:,2))
   if sum(Data(:,2)==k) > 0
       Level{2} = [Level{2} k];
   end
end
Dat = [];
Alert = 0;
NL = [length(Level{1}) length(Level{2})]; % 水準数
for k=1:NL(2)
    I = find(Data(:,2)==Level{2}(k));
    if k>1
        if length(I)~=size(Dat,1) % 要因Bの各水準のデータ数が一致しているかチェック
            Alert = 1;
        else
            Dat(:,:,k) = Data(I,:);
            if size(Dat,1)~=sum(Dat(:,1,k-1)==Dat(:,1,k)) % 対応したデータ同士で要因Aの水準が一致しているかチェック
                Alert = 1;
            end
        end
    else
  %      temp = Data(I,1);
        Dat = Data(I,:);
    end        
    for l=1:NL(1)
        I = find(Data(:,1)==Level{1}(l) & Data(:,2)==Level{2}(k));
        N(l,k) = length(I);
        M(l,k) = sum(Data(I,3));
    end
end

if Alert
    disp('  [anova_2bw]:  Factor B must be a repated-measures factor.');
    return;
end

ABS = sum(Data(:,3).^2);
AS = sum(sum(Dat(:,3,:),3).^2)/NL(2);
AB = sum(sum((M.^2)./N));
A = sum((sum(M').^2) ./ sum(N'));
M = M ./ N; % 各セルの平均値
Gp = sum(sum(M));
Xp = Gp^2 / (NL(1)*NL(2));
Ap = sum(mean(M,2).^2) * NL(2);
Bp = sum(mean(M).^2) * NL(1);
ABp = sum(sum(M.^2));
Ntil = NL(1) / sum(1./mean(N'));

SS = [
    Ntil * (Ap - Xp); % SSa
    AS - A; % SSs(a)
    Ntil * (Bp - Xp); % SSb
    Ntil * (ABp - Ap - Bp + Xp); % SSab
    ABS - AB - AS + A; % SSbs(a)
];

df = [
    NL(1) - 1; % dfa
    sum(mean(N')) - NL(1); % dfs(a)
    NL(2) - 1; % dfb
    (NL(1)-1) * (NL(2)-1); % dfab
    (sum(mean(N')) - NL(1)) * (NL(2)-1); % dfbs(a)
];

MS = SS ./ df;

F = [
    MS(1)/MS(2); % Fa
    MS(3)/MS(5); % Fb
    MS(4)/MS(5); % Fab
];

p = [
    p4F(F(1),df(1),df(2)); % pa
    p4F(F(2),df(3),df(5)); % pb
    p4F(F(3),df(4),df(5)); % pab
];

ES = [
    SS(1) / (SS(1) + SS(2));
    SS(3) / (SS(3) + SS(5));
    SS(4) / (SS(4) + SS(5));
];

Report = [
    'Source     ';
    'A          ';
    'Sample(A)  ';
    'B          ';
    'A*B        ';
    'B*Sample(A)';
];
sp = char(ones(6,3)*32);
Report = [Report sp strvcat('SS',num2str(SS)) sp strvcat('df',num2str(df)) sp strvcat('MS',num2str(MS))];
Report = [Report sp strvcat('F',num2str(F(1)),' ',num2str(F(2:3)),' ')];
Report = [Report sp strvcat('p',num2str(p(1)),' ',num2str(p(2:3)),' ')];
Report = [Report sp strvcat('pEta^2',num2str(ES(1)),' ',num2str(ES(2:3)),' ')];
temp = char(ones(1,size(Report,2))*45);
Report = [temp; Report(1,:); temp; Report(2:6,:); temp];
Report = [char(ones(9,3)*32) Report];

SS = SS';
df = df';
MS = MS';
F = F';
p = p';
ES = ES';
```
