## Extended Data 1 for "Scale space calibrates present and subsequent spatial learning in Barnes maze in mice": anova_2ww.m

```
function [Report,Level,M,N,F,p,MS,df,ES] = anova_2ww(Data)
% 
% anova_2ww:  2要因分散分析、2要因とも対応のある場合（データはリスト形式）
% 
% [Report,Level,M,N,F,p,MS,df,ES] = anova_2ww(Data)
%    Data    対象となるデータ（3列の行列）
%    Report  分散分析表
%    Level   水準番号の一覧（セルアレイ）
%    M       各条件の平均値
%    N       サンプル数（e.g.,参加者数）
%    F       F値
%    p       p値
%    MS      平均平方
%    df      自由度
%    ES      効果量（pEta^2:偏イータ2乗）
% 
% 入力データ（Data）は、1列目に要因Aの水準番号、2列目に要因Bの
% 水準番号、そして3列目に測定値が書かれた行列。すなわち、1行が
% 1サンプルです。
% 4名の参加者による要因A（2水準）×要因B（3水準）のデータの例：
%       Data = [
%          1    1    12.3
%          1    1    15.1
%          1    1     9.9
%          1    1    11.3
%          1    2    18.9
%          1    2    21.2
%          1    2    23.5
%          1    2    20.9
%          1    3    15.5
%          1    3    16.1
%          1    3    12.0
%          1    3    14.7
%          0    1    19.3
%          0    1    20.7
%          0    1    19.0
%          0    1    18.1
%          0    2    21.8
%          0    2    32.6
%          0    2    27.4
%          0    2    29.0
%          0    3    23.8
%          0    3    23.2
%          0    3    16.7
%          0    3    22.1
%       ];
% 各水準における行の順序は対応させて下さい。すなわち、上例の1行目, 5行
% 目, 9行目, 13行目, 17行目, 21行目は同一の参加者による（対応した）デー
% タです。水準番号は整数（負も可）。MやNは、水準番号の小さい順になって
% います。
% 
% see also:   anova_2wwm   anova_2bw   sme_2ww   tukey_2ww
% 
% (2015/09/10, by R. NIIMI)

Report = ['']; Level = []; M = []; N = []; F = []; p = []; MS = []; df = []; ES = [];

if ~(size(Data,1)>1 & size(Data,2)==3)
    disp('  [anova_2ww]:  Data must be a N by 3 matrix.');
    return;
end

Nall = size(Data,1);
Level{1} = min(Data(:,1));
for k=min(Data(:,1))+1:max(Data(:,1))
   if sum(Data(:,1)==k) > 0
       Level{1} = [Level{1} k];
   end
end
Level{2} = min(Data(:,2));
for k=min(Data(:,2))+1:max(Data(:,2))
   if sum(Data(:,2)==k) > 0
       Level{2} = [Level{2} k];
   end
end
Dat = [];
NL = [length(Level{1}) length(Level{2})]; % 水準数
N = Nall / (NL(1) * NL(2));
for k=1:NL(1)
    for l=1:NL(2)
        I = find(Data(:,1)==Level{1}(k) & Data(:,2)==Level{2}(l));
        M(k,l) = sum(Data(I,3));
        Dat(:,l,k) = Data(I,3); % データを3次元（S*B*A）の集計表形式にする
    end
end

ABS = sum(sum(sum(Dat.^2)));
X = sum(sum(sum(Dat))).^2 / (N*NL(1)*NL(2));
A = sum(sum(M').^2) / (N*NL(2));
B = sum(sum(M).^2) / (N*NL(1));
AB = sum(sum(M.^2)) / N;
S = sum(sum(sum(Dat,2),3).^2) / (NL(1)*NL(2));
AS = sum(sum(sum(Dat,2).^2)) / NL(2);
BS = sum(sum(sum(Dat,3).^2)) / NL(1);

SS = [
    S - X; % SSs
    A - X; % SSa
    AS - A - S + X; % SSas
    B - X; % SSb
    BS - B - S + X; % SSbs
    AB - A - B + X; % SSab
    ABS - AB - AS - BS + A + B + S - X; % SSabs
    ABS - X; % SSt
];

dfs = N - 1;
dfa = NL(1) - 1;
dfas = dfa * dfs;
dfb = NL(2) - 1;
dfbs = dfb * dfs;
dfab = dfa * dfb;
dfabs = dfas * dfb;
dft = Nall - 1;
df = [dfs dfa dfas dfb dfbs dfab dfabs dft]';

MS = SS ./ df;

F = [
    MS(2) / MS(3); % Fa
    MS(4) / MS(5); % Fb
    MS(6) / MS(7); % Fab
];

p = [
    p4F(F(1),dfa,dfas); % pa
    p4F(F(2),dfb,dfbs); % pb
    p4F(F(3),dfab,dfabs); % pab
];

ES = [
    SS(2) / (SS(2) + SS(3));
    SS(4) / (SS(4) + SS(5));
    SS(6) / (SS(6) + SS(7));
];

M = M / N; % 各セルの平均値

Report = [
    'Source    ';
    'Sample    ';
    'A         ';
    'A*Sample  ';
    'B         ';
    'B*Sample  ';
    'A*B       ';
    'A*B*Sample';
    'Total     ';
];
sp = char(ones(9,3)*32);
Report = [Report sp strvcat('SS',num2str(SS)) sp strvcat('df',num2str(df))];
Report = [Report sp strvcat('MS',num2str(MS(1:7)),' ')];
Report = [Report sp strvcat('F',' ',num2str(F(1)),' ',num2str(F(2)),' ',num2str(F(3)),' ',' ')];
Report = [Report sp strvcat('p',' ',num2str(p(1)),' ',num2str(p(2)),' ',num2str(p(3)),' ',' ')];
Report = [Report sp strvcat('pEta^2',' ',num2str(ES(1)),' ',num2str(ES(2)),' ',num2str(ES(3)),' ',' ')];
temp = char(ones(1,size(Report,2))*45);
Report = [temp; Report(1,:); temp; Report(2:8,:); temp; Report(9,:); temp];
Report = [char(ones(13,3)*32) Report];

SS = SS';
df = df';
MS = MS';
F = F';
p = p';
ES = ES';
```
