## Extended Data 1 for "Scale space calibrates present and subsequent spatial learning in Barnes maze in mice": anova_2wwm.m

```
function [Report,Level,M,N,F,p,MS,df,ES] = anova_2wwm(Data)
% 
% anova_2wwm:  2要因分散分析、2要因とも対応のある場合（データは集計表形式）
% 
% [Report,Level,M,N,F,p,MS,df,ES] = anova_2wwm(Data)
%    Data    対象となるデータ（集計表形式）
%    Report  分散分析表
%    Level   水準番号の一覧（セルアレイ）
%    M       各水準の平均値
%    N       サンプル数（e.g.,参加者数）
%    F       F値
%    p       p値
%    MS      平均平方
%    df      自由度
%    ES      効果量（pEta^2:偏イータ2乗）
% 
% 入力データ（Data）は、サンプル数×要因Bの水準数×要因Aの水準
% 数となる、3次元の行列です。同一行内の数値は対応したデータです。
% 
% 4名の参加者による要因A（2水準）×要因B（3水準）のデータの例：
%       Data(:,:,1) = [
%           19.3   21.8   23.8
%           20.7   32.6   23.2
%           19.0   27.4   16.7
%           18.1   29.0   22.1
%       ];
%       Data(:,:,2) = [
%           12.3   18.9   15.5
%           15.1   21.2   16.1
%            9.9   23.5   12.0
%           11.3   20.9   14.7
%       ];
% 
% see also:   anova_2ww   sme_2wwm
% 
% (2015/09/10, by R. NIIMI)

Report = ['']; Level = []; M = []; N = []; F = []; p = []; MS = []; df = []; ES = [];

if ~((size(Data,1)>1 & size(Data,2)>1) & size(Data,3)>1)
    disp('  [anova_2wwm]:  Data must be a 3-dimensional matrix.');
    return;
end

N = size(Data,1);
NL = [size(Data,3) size(Data,2)]; % 水準数
Nall = N * NL(1) * NL(2);
for k=1:NL(1)
    for l=1:NL(2)
        M(k,l) = sum(Data(:,l,k));
    end
end

ABS = sum(sum(sum(Data.^2)));
X = sum(sum(sum(Data))).^2 / (N*NL(1)*NL(2));
A = sum(sum(M').^2) / (N*NL(2));
B = sum(sum(M).^2) / (N*NL(1));
AB = sum(sum(M.^2)) / N;
S = sum(sum(sum(Data,2),3).^2) / (NL(1)*NL(2));
AS = sum(sum(sum(Data,2).^2)) / NL(2);
BS = sum(sum(sum(Data,3).^2)) / NL(1);

SS = [
    S - X; % SSs
    A - X; % SSa
    AS - A - S + X; % SSas
    B - X; % SSb
    BS - B - S + X; % SSbs
    AB - A - B + X; % SSab
    ABS - AB - AS - BS + A + B + S - X; % SSabs
    ABS - X; % SSt
];

dfs = N - 1;
dfa = NL(1) - 1;
dfas = dfa * dfs;
dfb = NL(2) - 1;
dfbs = dfb * dfs;
dfab = dfa * dfb;
dfabs = dfas * dfb;
dft = Nall - 1;
df = [dfs dfa dfas dfb dfbs dfab dfabs dft]';

MS = SS ./ df;

F = [
    MS(2) / MS(3); % Fa
    MS(4) / MS(5); % Fb
    MS(6) / MS(7); % Fab
];

p = [
    p4F(F(1),dfa,dfas); % pa
    p4F(F(2),dfb,dfbs); % pb
    p4F(F(3),dfab,dfabs); % pab
];

ES = [
    SS(2) / (SS(2) + SS(3));
    SS(4) / (SS(4) + SS(5));
    SS(6) / (SS(6) + SS(7));
];

M = M / N; % 各セルの平均値

Report = [
    'Source    ';
    'Sample    ';
    'A         ';
    'A*Sample  ';
    'B         ';
    'B*Sample  ';
    'A*B       ';
    'A*B*Sample';
    'Total     ';
];
sp = char(ones(9,3)*32);
Report = [Report sp strvcat('SS',num2str(SS)) sp strvcat('df',num2str(df))];
Report = [Report sp strvcat('MS',num2str(MS(1:7)),' ')];
Report = [Report sp strvcat('F',' ',num2str(F(1)),' ',num2str(F(2)),' ',num2str(F(3)),' ',' ')];
Report = [Report sp strvcat('p',' ',num2str(p(1)),' ',num2str(p(2)),' ',num2str(p(3)),' ',' ')];
Report = [Report sp strvcat('pEta^2',' ',num2str(ES(1)),' ',num2str(ES(2)),' ',num2str(ES(3)),' ',' ')];
temp = char(ones(1,size(Report,2))*45);
Report = [temp; Report(1,:); temp; Report(2:8,:); temp; Report(9,:); temp];
Report = [char(ones(13,3)*32) Report];

SS = SS';
df = df';
MS = MS';
F = F';
p = p';
ES = ES';
Level{1} = [1:NL(1)];
Level{2} = [1:NL(2)];
```
