## Extended Data 1 for "Scale space calibrates present and subsequent spatial learning in Barnes maze in mice": aprime.m

```
function [A] = aprime(HR,FA)
% 
% aprime: A'の計算
% 
% [A] = aprime(HR,FA)
%    A          A'値
%    HR         ヒット率
%    FA         誤警報率
% 
% A'を計算します。HRとFAはスカラーまたは等しい要素数の
% ベクトルです。HRとFAがベクトルの場合、Aも同じ要素数の
% ベクトルになります。
% 
% パフォーマンスがチャンスレベル未満の場合（HR < FA の
% 場合）は、Aaronson & Watts (1987), Psychological
%  Bulletin, vol. 102, pp. 439-442 の式(9)に従って計算
% します。
% 
% see also:   dprime   fitroc
% 
% (2009/04/14, by R. NIIMI)

A = [];

Check = find((HR<0) | (HR>1));
if 1-isempty(Check)
    disp('  [aprime]:  HR should be 0~1.');
    return;
end
Check = find((FA<0) | (FA>1));
if 1-isempty(Check)
    disp('  [aprime]:  FA should be 0~1.');
    return;
end


S = (HR>=FA);
HR2 = HR.*S + FA.*(1-S);
FA2 = FA.*S + HR.*(1-S);
S = S - (1-S);

A = 0.5 + S.* ((HR2-FA2).*(1+HR2-FA2)) ./ (4*HR2.*(1-FA2));
```
