## Extended Data 1 for "Scale space calibrates present and subsequent spatial learning in Barnes maze in mice": asintr.m

```
function [Result] = asintr(Data)
% 
% asintr: 変数の逆正弦変換
% 
% [Result] = asintr(Data)
%    Data    対象となるデータ（行列可）
%    Result  変換後のデータ
% 
% 逆正弦変換します。平方根とってarcsineする。それだけ。
% Data内の数値は、0〜1の範囲内でなければなりません。
% 
% (2008/10/09, by R. NIIMI)

Result = [];
if (min(min(Data)) < 0) | (max(max(Data)) > 1)
    disp('  [asintr]:  Data must be comprised of numbers 0~1.');
    Result = [];
    return;
end
Result = asin(sqrt(Data));
```
