## Extended Data 1 for "Scale space calibrates present and subsequent spatial learning in Barnes maze in mice": binom_c.m

```
function [Pc] = binom_c(X,N,P)
% 
% binom_c: 二項分布の累積確率
% 
% [Pc] = binom_c(X,N,P)
%    Pc    二項分布における、0〜Xまでの累積確率
%    X     事象の生起回数（スカラー）
%    N     試行回数（スカラー）
%    P     事象の生起確率（スカラー）
% 
% この関数は、N回の試行を繰り返したときに、生起確率Pの事象の生起回数が
% 0以上X以下である確率 Pc を計算します。二項検定などに利用できます。
% たとえば二者択一の実験課題を30試行実施した場合、まったくランダムに
% 選択肢を選ぶとすると正答する確率は P=0.5（つまりチャンスレベル）であ
% り、正答数が0〜20回である確率 Pc は binom_c(20,30,0.5) により 0.9786 
% となります。NとXが等しい場合には Pc=1 です。
% 
% see also:   binocdf (Statistics Toolbox)
% 
% (2009/01/23, by R. NIIMI)

Pc = [];

if 1-isscalar(N)
    disp('  [binom_c]:  N must be a scalar.');
    return;
end
if 1-isscalar(X)
    disp('  [binom_c]:  X must be a scalar.');
    return;
end
if 1-isscalar(P)
    disp('  [binom_c]:  P must be a scalar.');
    return;
end
if (P<0) | (P>1)
    disp('  [binom_c]:  P must be a proportion (0~1).');
    return;
end
if N<0 
    disp('  [binom_c]:  N must be 0 or larger.');
    return;
end
if X<0 
    disp('  [binom_c]:  X must be 0 or larger.');
    return;
end
if X>N
    disp('  [binom_c]:  X cannot exceed N.');
    return;
end

if X==N
    Pc = 1;
else
    Flag = (X>(N-X));
    if Flag
        temp = [N:-1:(X+1)];
    else
        temp = [0:X];
    end

    BC = [N:-1:1] ./ [1:N];
    BC = [1 BC];
    BC = BC(1:length(temp));
    BC = cumsum(log10(BC));
    Pc = BC + (log10(P)*temp) + (log10(1-P)*(N-temp));
    Pc = sum(10.^Pc);
    
    if Flag
        Pc = 1-Pc;
    end
end
```
