## Extended Data 1 for "Scale space calibrates present and subsequent spatial learning in Barnes maze in mice": bonf.m

```
function [Result]=bonf(p,Alpha,varargin)
% 
% bonf: ボンフェローニ法による多重比較
% 
% [Result] = bonf(p,Alpha,Method)
%    p       全対比較のp値（ベクトル）
%    Alpha   有意水準
%    Method  方法を指定します。
%             0 -> ボンフェローニの方法
%             1 -> ホルムの方法
%    Result   検定結果（1:有意, 0:n.s.）
% 
% ボンフェローニ法による多重比較を行います。pには、対象と
% なる対比較すべてのp値をあらかじめ計算し、ベクトルで与え
% ます。例えば4水準の間での多重比較では6つの対比較を行う
% ので、pは6要素です。
% 
% Alphaは有意水準で、スカラーです。例えば有意水準5パーセ
% ントならば、0.05とします。
% 
% オプション引数Methodで方法を選択できます。Methodが0の
% とき、個々の対比較に対する有意水準としてAlphaを対比較数
% で割った値を用います。Methodが1のときには、ホルムの方法
% を用います。省略すると、デフォルトとして0が用いられます。
% 
% Resultはpと同じ長さのベクトルで、検定の結果有意となった
% p値に対応する要素が1になっています。
% 
% see also:   bonf_ttest_2   bonf_ttest_2w   bonf_ttest_2p
%             bonf_chisqtest_ind
% 
% (2009/02/16, by R. NIIMI)

% clear;
% p = [28 31 26 7 19 2 14]/1000;
% Alpha = 0.05;
% Method = 0;

Result = [];
if 1-isvector(p)
    disp('  [bonf]:  p must be a vector.');
    return;
end
if 1-isscalar(Alpha)
    disp('  [bonf]:  Alpha must be a scalar.');
    return;
end
if (sum(p<0)+sum(p>1))>0
    disp('  [bonf]:  p must be 0~1.');
    return;
end
if ((Alpha<0) | (Alpha>1))
    disp('  [bonf]:  Alpha must be 0~1.');
    return;
end
Method = 0;
if ~isempty(varargin)
    if ~isempty(varargin{1})
        Method = varargin{1}(1);
    end
end
if sum(Method==[0 1])==0
    disp('  [bonf]:  Method must be 0 or 1.');
    return;
end    

Flag = 0;
if size(p,1)>1
    p = p';
    Flag = 1;
end
K = length(p);
p = [p' [1:K]'];
p = sortrows(p);
F = p(:,2)';
p = p(:,1)';
Result = zeros(1,K);

if Method==0 % standard Bonferroni
    A = Alpha/K;
    Result(F) = (p <= A);
end

if Method==1 % Holm
    A = Alpha./[K:-1:1];
    Result = (p>A);
    temp = find(Result);
    if isempty(temp) % All p is less than A
        Result = ones(1,K);
    else
        Result = ([1:K] < min(temp));
        Result(F) = Result;
        p(F) = p;
        A(F) = A;
    end
end
if Flag
    Result = Result';
end
```
