## Extended Data 1 for "Scale space calibrates present and subsequent spatial learning in Barnes maze in mice": bonf_chisqtest_ind.m

```
function [Report,Result,Xsq,p,df] = bonf_chisqtest_ind(Data,Alpha,varargin)
% 
% bonf_chisqtest_ind: ボンフェローニ法による多重比較、対比較はchisqtest_ind
% 
% [Report,Result,Xsq,p,df] = bonf_chisqtest_ind(Data,Alpha,Method,Yates)
%    Data    対象となるデータ（観測度数）
%    Alpha   有意水準
%    Method  方法を指定します。
%             0 -> ボンフェローニの方法
%             1 -> ホルムの方法
%    Yates    イエーツの連続性の修正をするには1を与えて下さい
%    Report  検定結果の表（カイ2乗値の表、* 有意差あり）
%    Result  検定結果（1:有意, 0:n.s.）
%    Xsq     各対比較の観測されたカイ2乗統計量
%    p       各対比較のp値
%    df      各対比較の自由度
% 
% 対比較に関数chisqtest_ind（独立性のカイ2乗検定）を用いて、
% ボンフェローニ法による多重比較を行います。
% 
% Data は chisqtest_ind と同じく、2つの要因によるクロス集計
% 表（分割表）形式の観測度数データを与えて下さい。例えば、
% 要因A（2カテゴリ）と要因B（3カテゴリ）の間の独立性の検定
% では、Data は2行×3列の行列です。リスト形式のデータを集計
% 表形式に変換するには list2matrix を利用して下さい。
% 
% この多重比較は、要因Aについての比率が、要因Bのどのカテゴリ
% 間で有意に異なるかを検討します。上の例であれば、要因Bは3カ
% テゴリなので、合計3つの対比較を行います。要因Aと要因Bを入
% れ換えるには、Dataを転置（Data'）して与えて下さい。
% 
% Alphaは有意水準で、スカラーです。例えば有意水準5パーセ
% ントならば、0.05とします。
% 
% オプション引数Methodで方法を選択できます。Methodが0の
% とき、個々の対比較に対する有意水準としてAlphaを対比較数
% で割った値を用います。Methodが1のときには、ホルムの方法
% を用います。省略すると、デフォルトとして0が用いられます。
%
% 引数 Yates は省略可能です（省略すると修正は行われません）。
% 
% Result, Xsq, p, df は、要因Bのカテゴリ数×要因Bのカテゴリ
% 数の行列です。
% 
% この関数は、関数 bonf, chisqtest_ind を利用します。
% 
% see also:   bonf   chisqtest_ind
% 
% (2009/02/19, by R. NIIMI)

Report = []; Result = []; Xsq = []; p = []; df = [];

if 1-isscalar(Alpha)
    disp('  [bonf_chisqtest_ind]:  Alpha must be a scalar.');
    return;
end
if ((Alpha<0) | (Alpha>1))
    disp('  [bonf_chisqtest_ind]:  Alpha must be 0~1.');
    return;
end
Method = 0;
Yates = 0;
if ~isempty(varargin)
    if ~isempty(varargin{1})
        Method = varargin{1}(1);
    end
    if length(varargin)>1
        if ~isempty(varargin{2})
            Yates = varargin{2}(1);
        end
    end
end
if sum(Method==[0 1])==0
    disp('  [bonf_chisqtest_ind]:  Method must be 0 or 1.');
    return;
end    
if sum(Yates==[0 1])==0
    disp('  [bonf_chisqtest_ind]:  Yates must be 0 or 1.');
    return;
end
if ndims(Data)>2
    disp('  [bonf_chisqtest_ind]:  Data must be a 2-dimensional matrix.');
    return;
end

NL = size(Data,2);
Level = [1:NL];
% M = mean(Data);
% Md = (ones(NL,1) * M) - (M' * ones(1,NL));
F1 = ones(NL,1) * [1:NL];
F2 = [1:NL]'*ones(1,NL);
F = find(F1 < F2);
F1 = F1(F);
F2 = F2(F);
Xsq = zeros(NL);
p = zeros(NL);
df = zeros(NL);
for I = 1:length(F)
    [Report,Xsq(F(I)),p(F(I)),df(F(I))] = chisqtest_ind(Data(:,[F1(I) F2(I)]),Yates);
end
temp = bonf(p(F),Alpha,Method);
Result = zeros(NL);
Result(F) = temp;
Result = Result + Result';
Xsq = Xsq + Xsq';
df = df + df';
p = p + p';

Sp = char(ones(NL,3)*32);
Report = [Sp char(ones(NL,1)*91) num2str(Level') char(ones(NL,1)*93)];
Report = [blanks(size(Report,2)); Report];
Sp = [blanks(3); Sp];
for j=1:NL
    NowCol = num2str(Xsq(:,j));
    temp = size(NowCol,2) - (length(num2str(Level(j)))+2);
    if temp>=0
        NowCol = [blanks(temp) '[' num2str(Level(j)) ']'; NowCol];
    else
        NowCol = ['[' num2str(Level(j)) ']'; [char(ones(NL,-temp)*32) NowCol]]; 
    end
    temp = [0; Result(:,j)];
    for i=1:NL+1
        Ast(i,:) = [char(ones(1,temp(i))*42)  blanks(2-temp(i))];
    end
    Report = [Report Sp NowCol Ast];
end
if Yates==0.5
    Report = strvcat(Report,' ','   (Yates'' continuity correction was applied.)');
end
```
