## Extended Data 1 for "Scale space calibrates present and subsequent spatial learning in Barnes maze in mice": bonf_ttest_2.m

```
function [Report,Result,Md,t,p,df] = bonf_ttest_2(Data,Alpha,varargin)
% 
% bonf_ttest_2: ボンフェローニ法による多重比較、対比較はttest_2
% 
% [Report,Result,Md,t,p,df] = bonf_ttest_2(Data,Alpha,Method)
%    Data    対象となるデータ（2列の行列）
%    Alpha   有意水準
%    Method  方法を指定します。
%             0 -> ボンフェローニの方法
%             1 -> ホルムの方法
%    Report  検定結果の表（平均値の差の表、* 有意差あり）
%    Result  検定結果（1:有意, 0:n.s.）
%    Md      平均値の差の表
%    t       各対比較のt値
%    p       各対比較のp値
%    df      各対比較の自由度
% 
% 対比較に関数ttest_2（対応のない2標本のt検定、分散が等質な
% 場合）を用いて、ボンフェローニ法による多重比較を行います。
% 
% Data は1列目に水準番号、2列目に測定値が書かれた行列で、
% anova_1b 等と同様のリスト形式のデータです。2要因以上の
% リスト形式データは受け付けません。
% 
% Alphaは有意水準で、スカラーです。例えば有意水準5パーセ
% ントならば、0.05とします。
% 
% オプション引数Methodで方法を選択できます。Methodが0の
% とき、個々の対比較に対する有意水準としてAlphaを対比較数
% で割った値を用います。Methodが1のときには、ホルムの方法
% を用います。省略すると、デフォルトとして0が用いられます。
%
% Result, Md, t, p, df は、水準数×水準数の行列です。これ
% らの変数の行および列の順序は、水準番号の値の小さい順序と
% なっています。
% 
% この関数は、関数 bonf, list2cell, ttest_2 を利用します。
% 
% see also:   bonf_ttest_2w   bonf_ttest_2p   bonf   ttest_2
% 
% (2009/02/19, by R. NIIMI)

Report = []; Result = []; Md = []; t = []; p = []; df = [];

if 1-isscalar(Alpha)
    disp('  [bonf_ttest_2]:  Alpha must be a scalar.');
    return;
end
if ((Alpha<0) | (Alpha>1))
    disp('  [bonf_ttest_2]:  Alpha must be 0~1.');
    return;
end
Method = 0;
if ~isempty(varargin)
    if ~isempty(varargin{1})
        Method = varargin{1}(1);
    end
end
if sum(Method==[0 1])==0
    disp('  [bonf_ttest_2]:  Method must be 0 or 1.');
    return;
end    

DataCell = []; Level = {[]};
Nall = size(Data,1);
NF = size(Data,2)-1;
if NF ~= 1
    disp('  [bonf_ttest_2]:  Data must be comprised of 2 columns.');
    return;
end

[DataCell,Level,N] = list2cell(Data);
Level = Level{1};
NL = length(N);
M = [];
for L = 1:NL
    M(L) = mean(DataCell{L});
end
Md = (ones(NL,1) * M) - (M' * ones(1,NL));
F1 = ones(NL,1) * [1:NL];
F2 = [1:NL]'*ones(1,NL);
F = find(F1 < F2);
F1 = F1(F);
F2 = F2(F);
p = zeros(NL);
t = zeros(NL);
df = zeros(NL);
for I = 1:length(F)
    [Report,t(F(I)),p(F(I)),df(F(I)),Ntemp] = ttest_2(DataCell{F1(I)},DataCell{F2(I)});
end
temp = bonf(p(F),Alpha,Method);
Result = zeros(NL);
Result(F) = temp;
Result = Result + Result';
t = t - t';
df = df + df';
p = p + p';

Sp = char(ones(NL,3)*32);
Report = [Sp char(ones(NL,1)*91) num2str(Level') char(ones(NL,1)*93)];
Report = [blanks(size(Report,2)); Report];
Sp = [blanks(3); Sp];
for j=1:NL
    NowCol = num2str(Md(:,j));
    temp = size(NowCol,2) - (length(num2str(Level(j)))+2);
    if temp>=0
        NowCol = [blanks(temp) '[' num2str(Level(j)) ']'; NowCol];
    else
        NowCol = ['[' num2str(Level(j)) ']'; [char(ones(NL,-temp)*32) NowCol]]; 
    end
    temp = [0; Result(:,j)];
    for i=1:NL+1
        Ast(i,:) = [char(ones(1,temp(i))*42)  blanks(2-temp(i))];
    end
    Report = [Report Sp NowCol Ast];
end
```
