## Extended Data 1 for "Scale space calibrates present and subsequent spatial learning in Barnes maze in mice": chisqtest_fit.m

```
function [Report,Xsq,p,df] = chisqtest_fit(Data,varargin)
% 
% chisqtest_fit: 適合度のカイ2乗検定
% 
% [Report,Xsq,p,df] = chisqtest_fit(Data,Ex,Yates)
%    Data     データ（観測度数）
%    Ex       期待度数もしくは期待比率
%    Yates    イエーツの連続性の修正をするには1を与えて下さい
%    Report   検定結果
%    Xsq      観測されたカイ2乗統計量
%    p        観測されたp値
%    df       自由度
% 
% ピアソンのカイ2乗統計量による適合度の検定を行います。Data に
% は観測度数を、Ex には期待度数もしくは期待比率を与えて下さい。
% 両者は等しい要素数のベクトルである必要があります。
% 
% Ex は度数・比率・比のいずれで指定しても可です。例えば、観測
% 数が100で、カテゴリが3つ、期待比率が[0.45 0.30 0.25]のとき、
% Ex に度数（ [45 30 25] ）、比率（ [0.45 0.30 0.25] ）、そして
% 比（ [9 6 5] ）のいずれを指定しても同じ計算を行います。
% 
% Ex が省略された場合、期待度数が全てのカテゴリで等しい値
% （ Dataの平均値 ）であるという帰無仮説について検定をします。
% 
% 引数 Yates は省略可能です（省略すると修正は行われません）。
% Ex を省略して Yates を指定するには、chisqtest_fit(Data,[],1)
% のように Ex に空配列を指定して下さい。
% 
% この関数は、関数 p4chisq.m を利用します。
% 
% see also:   chisqtest_ind
% 
% (2008/12/25, by R. NIIMI)

Report = ['']; Xsq = []; p = []; df = [];

N = sum(Data);
Nc = length(Data);
Ex = [];
Yates = 0;
if length(varargin)>0
    Ex = varargin{1};
    if length(varargin)>1
        if varargin{2}==1
            Yates = 0.5;
        end
    end
end
if isempty(Ex)
    Ex = N / Nc * ones(1,Nc);
end

if 1-isvector(Data)
    disp('  [chisqtest_fit]:  Data must be a vector.');
    return;
end
if 1-isvector(Ex)
    disp('  [chisqtest_fit]:  Ex must be a vector.');
    return;
elseif length(Ex)~=Nc
    disp('  [chisqtest_fit]:  Data and Ex must be matched in the number of elements.');
    return;
elseif sum(Ex)~=sum(Data)
    Ex = Ex/sum(Ex)*N;
end
if size(Data,1)>1
    Data = Data';
end
if size(Ex,1)>1
    Ex = Ex';
end

Xsq = sum(((abs(Data-Ex)-Yates).^2)./Ex);
df = Nc-1;
p = p4chisq(Xsq,df);
Report = ['  [chisqtest_fit]:  chi-square(' num2str(df) ',N=' num2str(N) ') = ' num2str(Xsq) ',  p = ' num2str(p)];
if Yates==0.5
    Report = strvcat(Report,'                    (Yates'' continuity correction was applied.)');
end
```
