## Extended Data 1 for "Scale space calibrates present and subsequent spatial learning in Barnes maze in mice": chisqtest_ind.m

```
function [Report,Xsq,p,df] = chisqtest_ind(Data,varargin)
% 
% chisqtest_ind: 独立性のカイ2乗検定
%
% [Report,Xsq,p,df] = chisqtest_ind(Data,Yates)
%    Data     データ（観測度数）
%    Yates    イエーツの連続性の修正をするには1を与えて下さい
%    Report   検定結果
%    Xsq      観測されたカイ2乗統計量
%    p        観測されたp値
%    df       自由度
% 
% ピアソンのカイ2乗統計量による独立性の検定を行います。Data には
% 2つの要因によるクロス集計表（分割表）形式の観測度数データを与え
% てください。例えば、要因A（2カテゴリ）と要因B（3カテゴリ）の間
% の独立性の検定では、Data は2行×3列の行列です。リスト形式のデー
% タを集計表形式に変換するには list2matrix を利用して下さい。
% 
% 引数 Yates は省略可能です（省略すると修正は行われません）。
% 
% 期待度数が1を下回るセルがあるか、期待度数が5を下回るセルが全セル
% の20％以上ある場合には、警告メッセージが出ます。
% 
% この関数は、関数 p4chisq.m を利用します。
% 
% see also:   chisqtest_fit   bonf_chisqtest_ind   list2matrix
% 
% (2009/02/16, by R. NIIMI)

Report = ['']; Xsp = []; p = []; df = [];

if (ndims(Data)~=2) | (min(size(Data))<2)
    disp('  [chisqtest_ind]:  Data must be 2-dimensional (N by M) matrix.');
    return;
end

Yates = 0;
if 1-isempty(varargin)
    if varargin{1}==1
        Yates = 0.5;
    end
end

N = sum(sum(Data));
Ex = (sum(Data,2) * sum(Data,1)) / N;
if (sum(sum(Ex<1))>0) | (sum(sum(Ex<5))>=(prod(size(Data))/5))
    disp('  [chisqtest_ind]:  Warning! The expected frequencies are very small in some cells.');
end
Xsq = sum(sum(((abs(Data-Ex)-Yates).^2)./Ex));
df = prod(size(Data)-1);
p = p4chisq(Xsq,df);
Report = ['  [chisqtest_ind]:  chi-square(' num2str(df) ',N=' num2str(N) ') = ' num2str(Xsq) ',  p = ' num2str(p)];
if Yates==0.5
    Report = strvcat(Report,'                    (Yates'' continuity correction was applied.)');
end
```
