## Extended Data 1 for "Scale space calibrates present and subsequent spatial learning in Barnes maze in mice": Contents.m

```
% sto version 0.30 (2015/09/22)
% 
% [t-Test]
%   ttest_1      1標本のt検定
%   ttest_2      対応のない2標本のt検定、分散が等質な場合
%   ttest_2w     対応のない2標本のt検定、分散が等質でない場合（ウェルチの検定）
%   ttest_2p     対応のある2標本のt検定
%   bonf_ttest_2    ボンフェローニ法による多重比較、対比較はttest_2
%   bonf_ttest_2w   ボンフェローニ法による多重比較、対比較はttest_2w
%   bonf_ttest_2p   ボンフェローニ法による多重比較、対比較はttest_2p
% 
% [One-Factoral ANOVA]
%   anova_1b     対応のない1要因分散分析
%   anova_1w     対応のある1要因分散分析（データはリスト形式）
%   anova_1wm    対応のある1要因分散分析（データは集計表形式）
%   tukey_1b     対応のない1要因分散分析における、TukeyのHSD法による多重比較
%   tukey_1w     対応のある1要因分散分析における、TukeyのHSD法による多重比較
%               （データはリスト形式）
%   tukey_1wm    対応のある1要因分散分析における、TukeyのHSD法による多重比較
%               （データは集計表形式）
% 
% [Two-Factoral ANOVA]
%   anova_2bb    2要因分散分析、2要因とも対応のない場合
%   anova_2bw    2要因分散分析、混合計画の場合
%   anova_2ww    2要因分散分析、2要因とも対応のある場合（データはリスト形式）
%   anova_2wwm   2要因分散分析、2要因とも対応のある場合（データは集計表形式）
%   tukey_2bb    2要因分散分析（2要因とも対応のない場合）での、TukeyのHSD法に
%                よる多重比較
%   tukey_2bw    2要因分散分析（混合計画の場合）での、TukeyのHSD法による多重
%                比較
%   tukey_2ww    2要因分散分析（2要因とも対応のある場合）での、TukeyのHSD法に
%                よる多重比較（データはリスト形式）
%   tukey_2wm    2要因分散分析（2要因とも対応のある場合）での、TukeyのHSD法に
%                よる多重比較（データは集計表形式）
%   sme_2bb      2要因分散分析（2要因とも対応のない場合）での単純主効果の検定
%   sme_2bw      2要因分散分析（混合計画の場合）での単純主効果の検定)
%   sme_2ww      2要因分散分析（2要因とも対応のある場合）での単純主効果の検定
%               （データはリスト形式）
%   sme_2wwm     2要因分散分析（2要因とも対応のある場合）での単純主効果の検定
%               （データは集計表形式）
%   tukey_sme_2bb   2要因分散分析（2要因とも対応のない場合）での、単純主効果に
%                   ついてのTukeyのHSD法による多重比較
%   tukey_sme_2bw   2要因分散分析（混合計画の場合）での、単純主効果についての
%                   TukeyのHSD法による多重比較
%   tukey_sme_2ww   2要因分散分析（2要因とも対応のある場合）での、単純主効果に
%                   ついてのTukeyのHSD法による多重比較（データはリスト形式）
%   tukey_sme_2wwm  2要因分散分析（2要因とも対応のある場合）での、単純主効果に
%                   ついてのTukeyのHSD法による多重比較（データは集計表形式）
% 
% [Pearson's R]
%   pr              ピアソンの積率相関係数Rの算出と無相関検定、直線回帰
%   prtest_1        1つの相関係数の仮説値との差の検定
%   prtest_2        複数の相関係数の差の検定
%   r2z             フィッシャーのz変換
%   z2r             フィッシャーのz変換の逆
% 
% [Chi-Squarel Tests]
%   chisqtest_fit        適合度のカイ2乗検定
%   chisqtest_ind        独立性のカイ2乗検定
%   bonf_chisqtest_ind   ボンフェローニ法による多重比較、対比較はchisqtest_ind
% 
% [Other Hypothesis Tests]
%   Utest           マン・ホイットニーのU検定
%   Htest           クラスカル・ウォリスのH検定
%   Ftest           F検定（2つの分散の差の検定）
%   binom_c         二項分布の累積確率
% 
% [Signal Detection Theory]
%   dprime          d'の計算
%   fitroc          ROC曲線のあてはめとd'の推定
%   roc             ROC曲線上で誤警報率に対応するヒット率の計算
%   rocinv          ROC曲線上でヒット率に対応する誤警報率の計算
%   aprime          A'の計算
% 
% [Utilities]
%   asintr       変数の逆正弦変換
%   matrix2list  集計表形式のデータをリスト形式に変換
%   list2matrix  リスト形式のデータを集計表形式に変換
%   list2cell    リスト形式のデータをセル配列に変換
%   sumoflist    リスト形式のデータの集計
%   findoutlier  リスト形式のデータから外れ値を検出する
% 
% [Other Functions]
%   p4ztest      z検定において、与えられたz値が臨界値となるようなp値（両側確率）
%   p4ttest      t検定において、与えられたt値が臨界値となるようなp値（両側確率）
%   p4F          与えられたF値が臨界値となるようなp値（上側確率）
%   p4chisq      与えられたカイ2乗値が臨界値となるようなp値（上側確率）
%   studrange_5  スチューデント化された範囲の5%点（TukeyのHSD法におけるq値）
%   studrange_1  スチューデント化された範囲の1%点（TukeyのHSD法におけるq値）
%   bonf         ボンフェローニ法による多重比較
%   num2rank     数値に順位をつける
%   interpNev    関数のNeville補間
% 
% [Help]
%   helpsto      readme.txtファイルの中身を表示します
% 
```
