## Extended Data 1 for "Scale space calibrates present and subsequent spatial learning in Barnes maze in mice": dprime.m

```
function [d,b,c] = dprime(HR,FA,varargin)
% 
% dprime: d'の計算
% 
% [d,b,c] = dprime(HR,FA,sdratio,wannaplot)
%    d          d'値（感度の指標）
%    b          β値（バイアスの指標）
%    c          判断基準（criterion）の位置
%    HR         ヒット率
%    FA         誤警報率
%    sdratio    N分布の標準偏差に対するSN分布の標準偏差の比
%    wannaplot  ROC曲線をプロットするには1を指定して下さい
% 
% ヒット率（hit rate, HR）と誤警報率（false-alarm rate, FA）
% から感度の指標d'を計算します。HRとFAはスカラーまたは等しい
% 要素数のベクトルです。HRとFAがベクトルの場合、d, b, cの第n
% 要素はHR(n)とFA(n)に対する計算結果です。
% 
% N分布とSN分布には正規分布を用います。sdratioには、両分布の
% 標準偏差の比（N分布の標準偏差を1としたときのSN分布の標準偏差）
% を指定して下さい。省略された場合には、1が指定されたものとみな
% します（つまり、両分布は等分散であると仮定します）。
% 
% dは、N分布の平均とSN分布の平均との距離（感度）を、N分布の標準
% 偏差を単位として表したものです。両分布が等分散でないという仮
% 定の下では、このdの値がSN分布の標準偏差を単位としていないもの
% であることに注意して下さい。なお、SN分布の標準偏差を単位として
% 表した感度は d/sdratio で求められます。
% 
% bは判断基準におけるN分布関数の値とSN分布関数の値の比（尤度比）
% であり、信号検出理論におけるβ（バイアスの指標）です。判断基準
% が両分布の交点に置かれている場合、bは1です。
% 
% cは判断基準のN分布における位置をz得点で表わした値です。N分布と
% SN分布が等分散の場合、判断基準のSN分布における位置は c-d です。
% 
% 計算結果に基づいてROC曲線をプロットするには、wannaplotに1を与
% えて下さい。省略するとプロットされません。HRとFAがベクトルの
% 場合、複数のROC曲線が重ねてプロットされます。
% 
% 複数の誤警報率とヒット率の組み合わせに対して単一のROC曲線をあ
% てはめ、d'を推定するには、fitrocを使って下さい。
% 
% see also:   aprime   fitroc   roc   rocinv
% 
% (2008/12/12, by R. NIIMI)

d = []; b = []; c = [];

Check = find((HR<0) | (HR>1));
if 1-isempty(Check)
    disp('  [dprime]:  HR should be 0~1.');
    return;
end
Check = find((FA<0) | (FA>1));
if 1-isempty(Check)
    disp('  [dprime]:  FA should be 0~1.');
    return;
end

sdratio = 1;
wannaplot = 0;
if length(varargin)>0
    sdratio = varargin{1};
    if length(sdratio)>1
        disp('  [dprime]:  sdratio should be a scalar.');
        return;
    end
    if sdratio<=0
        disp('  [dprime]:  sdratio should be larger than zero.');
        return;
    end
    if isempty(sdratio)
        sdratio = 1;
    end
    if length(varargin)>1
        wannaplot = varargin{2}(1);
    end
    if isempty(wannaplot)
        wannaplot = 0;
    end
end

Zn  = erfinv((1-FA)*2-1) * sqrt(2); % z-value of the criterion in terms of N distribution
Zsn = erfinv((1-HR)*2-1) * sqrt(2); % z-value of the criterion in terms of S+N distribution
d = Zn - (Zsn * sdratio);

b = exp((Zsn.^2)/(-2)) ./ exp((Zn.^2)/(-2));
c = Zn;
% c = (Zsn + Zn) / 2;

if wannaplot==1
    plotFA  = [0.001 0.005 ([1:99]/100) 0.995 0.999];
    plotZn  = erfinv((1-plotFA)*2-1) * sqrt(2);
    plotZsn = d'*ones(1,103) - ones(length(d),1)*plotZn;
    plotZsn = plotZsn / sdratio;
    plotHR  = (erf(plotZsn/sqrt(2))+1)/2;
    plotFA = [0 plotFA 1];
    plotHR = [zeros(length(d),1) plotHR ones(length(d),1)];
    plot(plotFA,plotHR,'r-'), hold on;
    plot(FA,HR,'b.'), hold on;
    plot([0 1],[0 1],'k-');
    axis square;
    xlim([0 1]); ylim([0 1]);
    xlabel('False Alarm'); ylabel('Hit');
end
```
