## Extended Data 1 for "Scale space calibrates present and subsequent spatial learning in Barnes maze in mice": findoutlier.m

```
function [Result,S] = findoutlier(Data,C1,C2,C1type,C2type)
% 
% findoutlier:  リスト形式のデータから外れ値を検出する
% 
% [Result,S] = findoutlier(Data,C1,C2,C1type,C2type)
%    Data     対象となるデータ（リスト形式）
%    C1       上側しきい値
%    C2       下側しきい値
%    C1type   上側しきい値のタイプ
%               （1=絶対値，2=平均+/-固定値，3=平均+/-C1*SD）
%    C2type   下側しきい値のタイプ
%               （1=絶対値，2=平均+/-固定値，3=平均+/-C2*SD）
%    Result   外れ値の行に1、非外れ値の行に0が入力された1列の行列
%    S        外れ値に関する集計結果
% 
% リスト形式のデータを集計して、外れ値に該当する行に1、そうでない
% 行に0が代入された、Dataと同じ行数の1列の行列Resultを返します。
% Dataの中の外れ値に該当する行を抽出するには、
%    Data(find(Result),:) 
% のようにして下さい。逆に、非外れ値の行を抽出するには、
%    Data(find(1-Result),:)
% のようにしてください。
% 
% しきい値のタイプ C1type / C2type が1の場合、C1およびC2の値そ
% のものをしきい値として用います。 C1type / C2type が2の場合、
% 各セルの平均値にC1を足した値を上側しきい値、C2を引いた値を下
% 側しきい値とします。C1type / C2type が3の場合、各セルの平均値
% +各セルの標準偏差*C1を上側しきい値、各セルの平均値-各セルの標
% 準偏差*C2を下側しきい値とします。また、C1やC2に空配列[]を指定
% すると、外れ値検出を行いません。
%   例1：平均+50を超える値を外れ値とする
%           findoutlier(Data,50,[],2,[])
%   例2：200msより短いか、平均+3SDより長い反応時間を外れ値とする
%           findoutlier(Data,3,200,3,1)
% 
% 変数Sはセル数の行を持つ行列で、外れ値処理の集計結果です。例えば
% 2要因（2水準×3水準）のデータについて処理した結果は、
% 
%    S = [
%       0     0    6    1    1    50.2438    28.7562
%       0     5    6    1    2    43.5587    29.7746
%       0    10    5    1    0    57.6783    26.9884
%       1     0    6    1    1    56.0912    41.9088
%       1     5    4    2    1    47.5511    34.1632
%       1    10    4    1    2    40.4792    29.8065
%    ];
% 
% のようになります。左列から要因Aの水準番号、要因Bの水準番号、
% 非外れ値の数、上側外れ値の数、下側外れ値の数、上側しきい値、
% 下側しきい値を示しています。例えば要因A=0，要因B=10のセルで
% は、上側外れ値が1つあり、下側外れ値はなく、5つのデータが残っ
% たことがわかります。なお、C1やC2に空配列が指定された場合や、
% データが存在しないセルでは、上例の4列目以降は0になります。
% 
% see also:   sumoflist
% 
% (2008/10/10, by R. NIIMI)

Result = []; S = [];

if ~(isscalar(C1) & isscalar(C2))
    disp('  [findoutlier]:  C1 and C2 must be scalar.');
    return;
end
if ~(isscalar(C1type) & isscalar(C2type))
    disp('  [findoutlier]:  C1type and C2type must be scalar.');
    return;
end
if (sum(C1type==[1 2 3]) * sum(C2type==[1 2 3]))==0
    disp('  [findoutlier]:  Please specify 1, 2, or 3 as C1type and C2 type.');
    return;
end

Nall = size(Data,1);
NF = size(Data,2)-1; % 要因数
if NF <= 1
    disp('  [findoutlier]:  Data must contain 2 or more columns.');
    return;
end

% 水準番号の抽出
Level = [];
for F = 1:NF
    Level{F} = min(Data(:,F));
    for k=min(Data(:,F))+1:max(Data(:,F))
        if sum(Data(:,F)==k) > 0
           Level{F} = [Level{F} k];
        end
    end
    NL(F) = length(Level{F}); % 水準数
end

Ncell = prod(NL); % セル数
S = [];
for F = 1:NF-1
    temp = mod([0:Ncell-1],prod(NL(F:NF)))+1;
    temp = ceil(temp/prod(NL((F+1):NF)));
    S = [S temp'];
end
temp = mod([0:Ncell-1],NL(NF))+1;
S = [S temp'];
for F = 1:NF
    S(:,F) = Level{F}(S(:,F));
end

temp = [];
Result = zeros(Nall,1);
for C=1:Ncell
    F1 = []; F2 = [];
    F0 = find(sum(Data(:,1:NF)==(ones(Nall,1)*S(C,:)),2)==NF);
    if isempty(F0)
        temp(C,:) = [0 0 0 0 0];
        disp(['  [findoutlier]:  No data in the cell ' num2str(S(C,:)) '.']);
    else
        NowData = Data(F0,NF+1);
        M = mean(NowData);
        SD = std(NowData);
        if (1-isempty(C1))
            if isempty(C1type)
                disp('  [findoutlier]:  Please specify C1type.');
                Result = []; S = [];
                return;
            elseif C1type==1
                NowC1 = C1;
            elseif C1type==2
                NowC1 = M + C1;
            elseif C1type==3
                NowC1 = M + C1*SD;
            else
                disp('  [findoutlier]:  Please specify 1, 2 or 3 as C1type.');
                Result = []; S = [];
                return;
            end
            F1 = find(NowData > NowC1);
            temp(C,2) = length(F1); % 上側外れ値の数
            temp(C,4) = NowC1; % 上側しきい値
            Result(F0(F1)) = ones(length(F1),1);
        else
            temp(C,2) = 0; % 上側外れ値の数
            temp(C,4) = 0; % 上側しきい値
        end
        if (1-isempty(C2))
            if isempty(C2type)
                disp('  [findoutlier]:  Please specify C2type.');
                Result = []; S = [];
                return;
            elseif C2type==1
                NowC2 = C2;
            elseif C2type==2
                NowC2 = M - C2;
            elseif C2type==3
                NowC2 = M - C2*SD;
            else
                disp('  [findoutlier]:  Please specify 1, 2 or 3 as C2type.');
                Result = []; S = [];
                return;
            end
            F1 = find(NowData < NowC2);
            temp(C,3) = length(F1); % 下側外れ値の数
            temp(C,5) = NowC2; % 下側しきい値
            Result(F0(F1)) = ones(length(F1),1);
        else
            temp(C,3) = 0; % 下側外れ値の数
            temp(C,5) = 0; % 下側しきい値
        end
        temp(C,1) = length(F0) - sum(temp(C,2:3)); % 非外れ値の数
    end
end
S = [S temp];
```
