## Extended Data 1 for "Scale space calibrates present and subsequent spatial learning in Barnes maze in mice": fitroc.m

```
function [d,sdratio_est] = fitroc(HR,FA,varargin)
% 
% fitroc: ROC曲線のあてはめとd'の推定
% 
% [d,sdratio_est] = fitroc(HR,FA,sdratio_hyp,wannaplot)
%    d             d'値（感度の指標）
%    sdratio_est   N分布の標準偏差に対するSN分布の標準偏差の比（データに基づく推定値）
%    HR            ヒット率
%    FA            誤警報率
%    sdratio_hyp   N分布の標準偏差に対するSN分布の標準偏差の比（仮定値）
%    wannaplot     ROC曲線をプロットするには1を指定して下さい
% 
% 与えられたヒット率と誤警報率のデータにROC曲線をあてはめ、d'を推
% 定します。FAとHRにはスカラーまたは要素数の等しいベクトルを指定し
% て下さい。N分布とSN分布には正規分布が用いられます。両分布の標準
% 偏差の比がわかっている場合には、sdratio_hypを指定して下さい。例
% えば、両分布が等分散であるという仮定の下でデータにROC曲線をあて
% はめるには、sdratio_hypに1を指定します。
% 
% sdratio_hypが省略された場合、FAに対応するz得点とHRに対応するz得点
% の関係に直線をあてはめ、その傾きによってN分布とSN分布の標準偏差の
% 比を推定します。推定された比の値はsdratio_estとして出力されます。
% （sdratio_hypが指定された場合は、その値がそのままsdratio_estに出
% 力されます。）
% 
% ただし、FAとHRがスカラーの場合（つまり、データが1点しかない場合）、
% N分布とSN分布の分散比は推定できません。この場合、（sdratio_hypが
% 指定されていない限り）自動的に標準偏差の比が1であると仮定して（つ
% まり、sdratio_hyp=1）計算が行われます。
% 
% wannaplotに1が指定されると、ROC曲線とデータをグラフにプロットしま
% す。省略されるとプロットしません。
% 
% ひとつひとつのヒット率と誤警報率の組み合わせに対してちょうどその
% 値を生じるようなd'を計算するには、dprimeを使って下さい。
% 
% この関数は、関数dprime.mを利用します。
% 
% see also:   dprime   roc   rocinv
% 
% (2009/04/14, by R. NIIMI)

d = []; sdratio_est = [];

if 1-isvector(HR)
    disp('  [fitroc]:  HR should be a scalar or vector.');
    return;
end
if 1-isvector(FA)
    disp('  [fitroc]:  FA should be a scalar or vector.');
    return;
end
L = length(HR);
if L~=length(FA)
    disp('  [fitroc]:  Numbers of elements of FA and d should be equal.');
    return;
end
Check = find((HR<0) | (HR>1));
if 1-isempty(Check)
    disp('  [fitroc]:  HR should be 0~1.');
    return;
end
Check = find((FA<0) | (FA>1));
if 1-isempty(Check)
    disp('  [fitroc]:  FA should be 0~1.');
    return;
end

sdratio_hyp = [];
wannaplot = 0;
if length(varargin)>0
    sdratio_hyp = varargin{1};
    if length(sdratio_hyp)>1
        disp('  [fitroc]:  sdratio_hyp should be a scalar.');
        return;
%    elseif length(sdratio_hyp)==1
%        sdratio_hyp = 0;
%    end
%    if sdratio_hyp<=0
%        disp('  [fitroc]:  sdratio_hyp should be larger than zero.');
%        return;
    end
    if length(varargin)>1
        wannaplot = varargin{2}(1);
    end
    if isempty(wannaplot)
        wannaplot = 0;
    end
end

if L==1
    if isempty(sdratio_hyp)
        sdratio_est = 1;
    else
        sdratio_est = sdratio_hyp;
    end
    d = dprime(HR,FA,sdratio_est,0);
else
    if isempty(sdratio_hyp)
        Zn  = erfinv((1-FA)*2-1) * sqrt(2); % z-value of the criterion in terms of N distribution
        Zsn = erfinv((1-HR)*2-1) * sqrt(2); % z-value of the criterion in terms of S+N distribution
        FitParam = polyfit(Zn,Zsn,1);
        sdratio_est = 1/FitParam(1);
        d = FitParam(2) * (-1) / FitParam(1);
    elseif sdratio_hyp<=0
        disp('  [fitroc]:  sdratio_hyp should be larger than zero.');
        return;
    else
        sdratio_est = sdratio_hyp;
        Zn  = erfinv((1-FA)*2-1) * sqrt(2); % z-value of the criterion in terms of N distribution
        Zsn = erfinv((1-HR)*2-1) * sqrt(2); % z-value of the criterion in terms of S+N distribution
        FitParam = [1/sdratio_hyp polyfit(Zn,(Zsn-Zn/sdratio_hyp),0)];
        d = FitParam(2) * (-1) * sdratio_hyp;
    end
end

if wannaplot==1
    plotFA  = [0.001 0.005 ([1:99]/100) 0.995 0.999];
    plotZn  = erfinv((1-plotFA)*2-1) * sqrt(2);
    plotZsn = d - plotZn;
    plotZsn = plotZsn / sdratio_est;
    plotHR  = (erf(plotZsn/sqrt(2))+1)/2;
    plotFA = [0 plotFA 1];
    plotHR = [0 plotHR 1];
    plot(plotFA,plotHR,'r-'), hold on;
    plot(FA,HR,'b.'), hold on;
    plot([0 1],[0 1],'k-');
    axis square;
    xlim([0 1]); ylim([0 1]);
    xlabel('False Alarm'); ylabel('Hit');
end
```
