## Extended Data 1 for "Scale space calibrates present and subsequent spatial learning in Barnes maze in mice": Ftest.m

```
function [Report,F,p,df,S2] = Ftest(A,B)
% 
% Ftest: F検定（2つの分散の差の検定）
% 
% [Report,F,p,df,S2] = Ftest(A,B)
%    A        標本1（ベクトル）
%    B        標本2（ベクトル）
%    Report   検定結果
%    F        観測されたF値（不偏分散の比）
%    p        観測されたp値（両側）
%    df       自由度
%    S2       観測された分散
% 
% 分子側を標本1、分母側を標本2としたF値が、両標本の母分散が等しいと
% いう帰無仮説のもとで観測されるかを検定します。検定は両側です。
% 標本1と標本2を入れ替えると（e.g., Ftest(A,B) -> Ftest(B,A) ）、F値
% は逆数になりますが、p値は変わりません。
% 
% この関数は、関数 p4F.m を利用します。
% 
% see also:   vartest2 (Statistics Toolbox)
% 
% (2009/02/10, by R. NIIMI)

Report = ['']; F = []; p = []; df = []; S2 = [];

if ~(isvector(A) & isvector(B))
    disp('  [Ftest]:  Please input vectors as data.');
    return;
end

N = [length(A) length(B)];
M = [mean(A) mean(B)];
S2 = [sum((A-M(1)).^2) sum((B-M(2)).^2)]./(N-1);
F = S2(1)/S2(2);
df = N-1;

if F < 1
    p = p4F(1/F,df(2),df(1));
else
    p = p4F(F,df(1),df(2));
end
p = p*2;
Report = ['  [Ftest]:  F(' num2str(df(1)) ',' num2str(df(2)) ') = ' num2str(F) ',  p = ' num2str(p) ' (two-tailed)'];
```
