## Extended Data 1 for "Scale space calibrates present and subsequent spatial learning in Barnes maze in mice": Htest.m

```
function [Report,Level,M,N,H,p,df] = Htest(Data)
% 
% Htest: クラスカル・ウォリスのH検定
% 
% [Report,Level,M,N,H,p,df] = Htest(Data)
%    Data     対象となるデータ（2列の行列）
%    Report   検定結果
%    Level    水準番号の一覧
%    M        各水準の中央値
%    N        各水準のサンプル数
%    H        観測されたH値
%    p        観測されたp値
%    df       カイ2乗分布の自由度
% 
% H統計量を用いて、対応がない複数の条件の中央値に差があるか
% を検定します。検定はH統計量の分布がカイ2乗分布で近似でき
% ることを利用して行われます。標本数が少ないと、この近似は
% 正確ではありません。標本数が5以下の条件がひとつでもあると
% 警告メッセージを表示します（計算は行われます）。この場合
% は、統計法の専門書などに掲載されているH検定のための数表を
% 利用して検定を行うことをお勧めします。
% 
% 入力データ（Data）はリスト形式です。すなわち、1列目に水準
% 番号、2列目に測定値が書かれた行列です。（anova_1bなどと同
% 形式です）。
% 
% この関数は、関数 p4chisq.m および num2rank.m を利用します。
% 
% see also:   kruskalwallis (Statistics Toolbox)
% 
% (2009/02/10, by R. NIIMI)

Report = ['']; H = []; p = []; df = [];

if ~(size(Data,1)>1 & size(Data,2)==2)
    disp('  [Htest]:  Data must be a N by 2 matrix.');
    return;
end

[RankData, EqData] = num2rank(Data(:,2),1,0);
Data = [Data RankData];
Nall = size(Data,1);
Level = min(Data(:,1));
for k=min(Data(:,1))+1:max(Data(:,1))
   if sum(Data(:,1)==k) > 0
       Level = [Level k];
   end
end
NL = length(Level);
Rs = [];
for k=1:NL
    I = find(Data(:,1)==Level(k));
    N(k) = length(I);
    M(k) = median(Data(I,2));
    Rs(k) = sum(Data(I,3)); % rank-sum
end
H = 12/(Nall*(Nall+1)) * sum((Rs.^2)./N) - 3*(Nall+1);

if sum(N<=5)~=0
    disp('  [Htest]:  Very small data (n<=5). Result may be imprecise.');
    disp('            Please consult statistical tables for H-test.');
end

if ~isempty(find(EqData))
    temp = RankData.*EqData;
    temp = [0 sort(temp)' 0];
    temp = find((temp(2:Nall+2)-temp(1:Nall+1))~=0);
    temp = temp(2:end)-temp(1:end-1);
    temp = sum(temp.^3 - temp);
    temp = 1 - (temp/(Nall*(Nall^2-1)));
    H = H / temp;
end

df = NL-1;
p = p4chisq(H,df);
Report = ['  [Htest]:  H = ' num2str(H) ',  p = ' num2str(p) ' (two-tailed)'];
Report = strvcat(Report,['            (Tested with approximation by chi-square distribution of df = ' num2str(df) ').']);
```
