## Extended Data 1 for "Scale space calibrates present and subsequent spatial learning in Barnes maze in mice": interpNev.m

```
function [Yc] = interpNev(X,Y,Xc)
% 
% interpNev: 関数のNeville補間
% 
% [Yc] = interpNev(X,Y,Xc)
%    X     xの値（ベクトル）
%    Y     Xに対する既知のyの値（Xと同じ長さのベクトル）
%    Xc    Xに含まれていないxの値（スカラー）
%    Yc    Xcに対するyの推定値
% 
% interpNevは、関数y=f(x)において、xとyの値が離散的にわかっている場合、それら
% の値を元にあるxに対する未知のyの値を推定する。例えば関数y=log(x)について、
%    X = [1 2 3 4 5 6]
%    Y = [0    0.6931    1.0986    1.3863    1.6094    1.7918]
% がわかっているとする。これらの値を元に、x=3.5のときのyの値を推定すると、
%    Yc = interpNev(X,Y,3.5)
% により、log(3.5)の推定値1.2521を得る（実際の値は1.2528）。X, Yのデータ数
% が多いほど、また補間する関数f(x)が直線に近いほど、正確な推定が期待できる。
% 
% (2008/12/19, by R. NIIMI)

Yc = [];
if ~(isvector(X) & isvector(Y))
    disp('  [interpNev]:  Please input vectors as X and Y.');
    return;
end
if length(X)~=length(Y)
    disp('  [interpNev]:  X and Y must have identical numbers of elements.');
    return;
end
if ~isscalar(Xc)
    disp('  [interpNev]:  Please input a scalar as Xc.');
    return;
end

Dat = sortrows([(abs(X-Xc))' X' Y']);
X = Dat(:,2);
Y = Dat(:,3);
N = length(X);

j = 1;
I = Y;

while j < N
	for k=j:(N-1)
		I(k+1,j+1) = ( (X(k+1)-Xc)*I(k,j) - (X(k-j+1)-Xc)*I(k+1,j) ) / (X(k+1)-X(k-j+1));
	end
	j = j + 1;
end

Yc = I(N,N);
```
