## Extended Data 1 for "Scale space calibrates present and subsequent spatial learning in Barnes maze in mice": list2cell.m

```
function [DataCell,Level,N] = list2cell(Data)
% 
% list2cell:  リスト形式のデータをセル配列に変換
% 
% [DataCell,Level,N] = list2cell(Data)
%    Data      対象となるデータ（リスト形式）
%    DataCell  集計表形式に変換されたデータ
%    Level     水準番号の一覧のセルアレイ
%    N         各セルのデータ数
% 
% 2次元行列のリスト形式データを多次元セル配列の変数に格納し
% ます。
% 
% 例えば、2水準の要因A×3水準の要因B（対応なし）の計画で、
% 各セルの参加者数が0〜3名の実験結果 Data が下のようなリスト
% 形式だったとします。第1列は要因Aの水準番号、第2列は要因Bの
% 水準番号、第3列がデータです。
%   Data = [
%       0   -5   87
%       0   -5   61
%       0   -5   77
%       0   10   49
%       0   10   50
%       0   25   66
%       0   25   74
%       1   -5   39
%       1   -5   41
%       1   -5   55
%       1   25   34
%       1   25   44
%   ];
% 上例ではわかりやすくするため行の並び順を整理していますが、
% 実際には行の順序はランダムでもかまいません。この Data を
% list2cell によってセル配列に変換すると下のようになります。
% DataCell = 
%     [3x1 double]    [3x1 double]
%     [2x1 double]    [0x1  double]
%     [2x1 double]    [2x1 double]
% DataCell の第1次元は要因Bに、第2次元は要因Aに対応しています。
% 例えば要因Aが第1水準（水準番号は0）で、要因Bが第3水準（水準
% 番号は25）のデータは、 DataCell{3,1} で与えられます。各水準
% は、水準番号の値が小さい順に第1水準、第2水準…とみなされます。
% 
% Level には、水準番号の一覧を返します。Level はセル配列です。
% 例えば第2の要因（要因B）の水準番号の一覧は、Level{2} で与え
% られます（上例では [-5 10 25] となります）。
% 
% N は各セルのデータ数です。DataCellと同じ次元数のdouble変数
% です。例えば要因Aが第1水準、要因Bが第3水準のセルのデータ数
% は、 N(3,1)で与えられます（上例では2です）。
% 
% 各セルのデータ数が等しい場合（完全に対応のある実験計画など）
% では、セル配列ではなく行列変数に変換する list2matrix も利用
% できます。
% 
% see also:   list2matrix   sumoflist
% 
% (2009/02/13, by R. NIIMI)

DataCell = []; Level = {[]}; N = [];
Nall = size(Data,1);
NF = size(Data,2)-1;
if NF < 1
    disp('  [list2cell]:  Data must contain 2 or more columns.');
    return;
end

% 水準番号の抽出
Level = []; NL = [];
for F = 1:NF
    Level{F} = min(Data(:,F));
    for k=min(Data(:,F))+1:max(Data(:,F))
        if sum(Data(:,F)==k) > 0
           Level{F} = [Level{F} k];
        end
    end
    NL(F) = length(Level{F}); % 水準数
end

Ncell = prod(NL); % セル数

S = [];
for F = 1:NF-1
    temp = mod([0:Ncell-1],prod(NL(F:NF)))+1;
    temp = ceil(temp/prod(NL((F+1):NF)));
    S = [S temp'];
end
temp = mod([0:Ncell-1],NL(NF))+1;
S = [S temp'];
for F = 1:NF
    S(:,F) = Level{F}(S(:,F));
end

if NF<=1
    DataCell = cell(NL,1);
    N = zeros(NL,1);
else
    DataCell = cell(fliplr(NL));
    N = zeros(fliplr(NL));
end
for C=1:Ncell
    F = find(sum(Data(:,1:NF)==(ones(Nall,1)*S(C,:)),2)==NF);
    DataCell{C} = Data(F,NF+1);
    N(C) = length(F);
end
```
