## Extended Data 1 for "Scale space calibrates present and subsequent spatial learning in Barnes maze in mice": list2matrix.m

```
function [Matrix,Level] = list2matrix(Data)
% 
% list2matrix:  リスト形式のデータを集計表形式に変換
% 
% [Matrix,Level] = list2matrix(Data)
%    Data    対象となるデータ（リスト形式）
%    Matrix  集計表形式に変換されたデータ
%    Level   水準番号の一覧のセルアレイ
% 
% 2次元行列のリスト形式データを多次元行列の集計表形式に変換し
% ます。各セルのデータ数は等しくなければなりません。
% 
% 例えば、2水準の要因A×3水準の要因Bという計画で、2名の参加者
% （もしくは、2水準の要因C）による実験結果 Data が下のような
% リスト形式だったとします。第1列は要因Aの水準番号、第2列は要
% 因Bの水準番号、第3列が参加者番号もしくは要因Cの水準番号、そ
% して第4列がデータです。
%   Data = [
%       0   -5   1   87
%       0   -5   2   61
%       0   10   1   49
%       0   10   2   50
%       0   25   1   66
%       0   25   2   74
%       1   -5   1   39
%       1   -5   2   55
%       1   10   1   62
%       1   10   2   29
%       1   25   1   34
%       1   25   2   44
%   ];
% 上例ではわかりやすくするため行の並び順を整理していますが、
% 実際には行の順序はランダムでも同じ結果になります。この Data
% を list2matrix によって変換すると、下のようになります。
%    Matrix(:,:,1) = [
%       87    49    66
%       61    50    74
%    ];
%    Matrix(:,:,2) = [
%       39    62    34
%       55    29    44
%    ];
% Matrix の第1次元は参加者番号もしくは要因Cに、第2次元は要因B
% に、第3次元は要因Aに対応しています。例えば要因Aが第1水準（水
% 準番号は0）で、要因Bが第3水準（水準番号は25）のデータは、
%  Matrix(:,3,1) で与えられます。各水準は、水準番号の値が小さ
% い順に第1水準、第2水準…とみなされます。
% 
% 上例では各セルのデータ数が1でしたが、例えば Data の第3列目が
% ない場合（つまり、Data(:,[1 2 4]) ）のように、各セルのデータ
% 数が2以上でも動作可能です。例えば対応のない実験計画で、参加者
% 番号が存在しない場合にはそのようなデータになるでしょう。ただ
% しこの場合、元の Data の行の順序が変わると変換結果 Matrix の
% 第1次元でのデータの並び順も変わります。
% 
% Level には、水準番号の一覧を返します。Level はセル配列です。
% 例えば第2の要因（要因B）の水準番号の一覧は、Level{2} で与えら
% れます（上例では [-5 10 25] となります）。
% 
% see also:   matrix2list   list2cell   sumoflist
% 
% (2009/02/16, by R. NIIMI)

Matrix = []; Level = {[]};
Nall = size(Data,1);
NF = size(Data,2)-1;
if NF < 1
    disp('  [list2matrix]:  Data must contain 2 or more columns.');
    return;
end

NL = [];
temp = [];
Alert = 0;
for f=1:NF
    Level{f} = [];
    Nlevel{f} = [];
    for k=min(Data(:,f)):max(Data(:,f))
        I = find(Data(:,f)==k);
        if 1-isempty(I)
            Level{f} = [Level{f} k];
            Nlevel{f} = [Nlevel{f} length(I)];
            temp(I,f) = length(Level{f})*ones(length(I),1);
        end
        if sum(mean(Nlevel{f})==Nlevel{f}) ~= length(Level{f})
            Alert = 1;
        end
    end
    NL = [NL length(Level{f})];
end
if Alert
    disp('  [list2matrix]:  Cells are not balanced. Please equate the N of all cell.');
    return;
end
temp = fliplr(temp);
Data = [temp Data(:,NF+1)];
Data = sortrows(Data,[NF:-1:1]);

Ncell = prod(NL);
if Nall < Ncell
    disp('  [list2matrix]:  Empty cell is found. Please equate the N of all cell.');
    return;
else
    Rep = Nall / Ncell;
    if Rep~=round(Rep)
        disp('  [list2matrix]:  Cells are not balanced. Please equate the N of all cell.');
        return;
    elseif Rep==1
        Matrix = zeros(fliplr(NL));
        Matrix(1:Nall) = Data(:,NF+1);
    else
        temp = mod([0:(Nall-1)],Rep)+1;
        Data = [Data(:,1:NF) temp' Data(:,NF+1)];
        Matrix = zeros(fliplr([NL Rep]));
        Matrix(1:Nall) = Data(:,NF+2);
    end
end
```
