## Extended Data 1 for "Scale space calibrates present and subsequent spatial learning in Barnes maze in mice": matrix2list.m

```
function [List] = matrix2list(Data)
% 
% matrix2list:  集計表形式のデータをリスト形式に変換
% 
% [List] = matrix2list(Data)
%    Data    対象となるデータ（集計表形式）
%    List    リスト形式に変換されたデータ
% 
% 例えば、2水準の要因A×3水準の要因Bという計画で、4名の参加者
% （もしくは、4水準の要因C）による実験結果 Data が下のような
% 集計表形式だったとします（要因Aが第3次元、要因Bが第2次元、
% 参加者もしくは要因Cが第1次元に相当しています）。
% 
%   Data(:,:,1) = [
%       10     8    13
%        9     6    15
%       12    10    14
%       15    16    20
%   ];
%   Data(:,:,2) = [
%       13    11    16
%       12     9    18
%       15    13    17
%       18    19    23
%   ];
% 
% MATLAB上で扱うにはこのように行列になっている方が集計に便利
% ですが、一般的な統計ソフトでは水準番号を用いたリスト形式が
% 使われます。そこで、この関数によって上のデータを変換すると、
% 
%   List = [
%     1     1     1    10
%     1     1     2     9
%     1     1     3    12
%     1     1     4    15
%     1     2     1     8
%     1     2     2     6
%       ---（中略）---
%     2     3     3    17
%     2     3     4    23
%  ];
% 
% のようになります。変数Listの第1列が要因Aの水準番号、第2列が
% 要因Bの水準番号、第3列が参加者番号もしくは要因Cの水準番号、
% そして第4列がデータです。
% 
% anova_1w や anova_2ww などの関数でリスト形式のデータに対して
% 対応のある分析をする際には、参加者番号は不要です。上例であれ
% ば3列目を削除して (e.g., List = List(:,[1:2 4]); ）から与え
% てください。
% 
% see also:   list2matrix
% 
% (2008/10/21, by R. NIIMI)

List = [];

DataSize = size(Data);
N = numel(Data);
DataDim  = length(DataSize);

for D = DataDim:-1:1
    temp = mod([0:N-1],prod(DataSize(1:D)))+1;
    temp = ceil(temp/prod(DataSize(1:D-1)));
    List = [List temp'];
end
List = [List Data(1:N)'];
```
