## Extended Data 1 for "Scale space calibrates present and subsequent spatial learning in Barnes maze in mice": num2rank.m

```
function [rankdata,eqdata] = num2rank(data,varargin)
% 
% num2rank: 数値に順位をつける
% 
% [RankData,EqData] = num2rank(Data,HowAboutEq,Ord)
%    Data        入力データ（行列可）
%    HowAboutEq  結びの処理方法
%                 0 -> 結びの処理をしません
%                 1 -> 結びの順位は平均順位（デフォルト）
%                 2 -> 結びの順位は最小同順位
%    Ord         昇順の場合は0、降順の場合は1を指定して下さい
%    RankData    順位
%    EqData      結びの有無
% 
% Dataの各要素の大きさ順に順位をつけます。Dataは多次元行列可
% です。RankDataは順位で、Dataと同じサイズの行列です。例えば
% num2rank([5 9 2]) は、RankDataに行列 [2 3 1] を出力します。
% 
% 引数Ordに0を指定するとRankDataは昇順の順位に、Ordに1を指定
% するとRankDataは降順の順位（上例なら[2 1 3]）となります。
% Ordは省略可です。省略されるとRankDataは昇順になります。
% 
% Dataに複数の同値の要素（結び）がある場合（例：[5 9 2 5]）、
% 引数HowAboutEqで結び要素に対する順位のつけかたを決めます。
% 
% HowAboutEqが0の場合、結びの処理は行わず、同値の要素にも連
% 続した異なる順位を自動的に与えます（例：[2 4 1 3]）。
% 
% HowAboutEqに1が指定されていると結びの要素の順位は平均値に
% なります（例：[2.5 4.0 1.0 2.5]）。従って、RankDataは小数
% になったり、「飛び」（存在しない順位）が生じたりします。
% 
% HowAboutEqに2が指定されると、結びの要素には最小の同順位が
% 与えられ、他の要素の後続順位はこの順位に続くように（「飛
% び」がないように）調整されます。上例であれば [2 3 1 2] と
% なります。この場合、RankDataの最大値はDataの要素数より小
% さくなります。
% 
% HowAboutEqは省略可です。省略されると1が指定されたものとみ
% なします。
% 
% EqDataはDataと同じサイズの行列で、Dataで結びがある位置に対応
% する要素に1を、それ以外の要素に0を与えます（例：[1 0 0 1]）。
% 
% (2009/02/19, by R. NIIMI)

rankdata = []; eqdata = [];

howabouteq = 1; ord = 0;
L = length(varargin);
if L>0
    howabouteq = varargin{1};
    if L>1
        ord = varargin{2};
    end
end
if isempty(howabouteq)
    howabouteq = 1;
else
    howabouteq = howabouteq(1);    
end
if isempty(ord)
    ord = 0;
else
    howabouteq = howabouteq(1);    
end
if sum(howabouteq==[0 1 2])==0
    disp('  [num2rank]:  Please input 0, 1, or 2 as HowAboutEq.');
    return;
end
if sum(ord==[0 1])==0
    disp('  [num2rank]:  Please input 0 or 1 as Ord.');
    return;
end

rankdata = data*0;
eqdata = data*0;
N = numel(data);
I = [1:N];

data2 = data(I);
if size(data2,1)>1
    data2 = data2';
end
data2 = [data2; I]';
data2 = sortrows(data2,1);
data2 = [data2 I'];

Ieq = (data2(1:N-1,1)==data2(2:N,1));
eqdata(data2(:,2)) = ([Ieq' 0] | [0 Ieq']);
if howabouteq>0
    Ieq = [0 Ieq' 0];
    Ieq = Ieq(2:N+1) - Ieq(1:N);
    IeqTop = find(Ieq==1);
    IeqEnd = find(Ieq==-1);
    if howabouteq==1
        for J = 1:length(IeqTop)
            data2(IeqTop(J):IeqEnd(J),3) = mean(data2(IeqTop(J):IeqEnd(J),3));
        end
    elseif howabouteq==2
        for J = 1:length(IeqTop)
            Neq = IeqEnd(J)-IeqTop(J)+1;
            NowRank = min(data2(IeqTop(J):IeqEnd(J),3));
            data2(IeqTop(J):IeqEnd(J),3) = ones(Neq,1) * NowRank;
            temp = find(data2(:,3)>NowRank);
            data2(temp,3) = data2(temp,3)-(Neq-1);
        end
    end
end
rankdata(data2(:,2)) = data2(:,3);
if ord==1
    rankdata = max(data2(:,3))+1-rankdata;
end
```
