## Extended Data 1 for "Scale space calibrates present and subsequent spatial learning in Barnes maze in mice": p4chisq.m

```
function [p] = p4chisq(ChiSqData,dfdata,varargin)
% 
% p4chisq: 与えられたカイ2乗値が臨界値となるようなp値（上側確率）
% 
% [p] = p4chisq(chisq,df,TOL)
%    p       カイ2乗分布における、与えられたカイ2乗値から無限大までの累積確率
%    chisq   カイ2乗値
%    df      自由度
%    TOL     求積の絶対誤差許容範囲
% 
% chisq, dfともにスカラーを与えて下さい。
% 
% p値はMATLABの数値積分関数（quadl）を利用して計算していま
% す。引数TOLはその際の誤差許容範囲です。詳しくはquadlのヘ
% ルプを参照して下さい。TOLは省略されるとデフォルト値とし
% て1e-06が用いられます。
% 
% see also:   chi2cdf (Statistics Toolbox)
% 
% (2008/12/19, by R. NIIMI)

p = [];

TOL = 1e-06; % default value
if 1-isempty(varargin)
    TOL = varargin{1};
end
if isempty(TOL)
    TOL = 1e-06;
else
    TOL = TOL(1);
end

if 1-isscalar(dfdata)
    disp('  [p4chisq]:  df must be a scalar.');
    return;
end
if 1-isscalar(ChiSqData)
    disp('  [p4chisq]:  chisq must be a scalar.');
    return;
end
if ChiSqData<0
    disp('  [p4chisq]:  Please specify 0 or larger value as chisq.');
    return;
end
if ChiSqData==0
    p = 1;
    return;
end

A = (2*gamma(dfdata/2))*(2^(dfdata/2-1));
if dfdata>=2
    ChiSqDist = @(ChiSq) ChiSq.^(dfdata/2-1) .* exp(-ChiSq/2) ./A;
else % To avoid y=+Inf within the region of quadrature, the function is transformed.
    ChiSqDist = @(u) exp( (u.*dfdata/2).^(2/dfdata) ./(-2) ) ./A;
    ChiSqData = ChiSqData^(dfdata/2) *2/dfdata;
end
p = quadl(ChiSqDist,0,ChiSqData,TOL);
p = 1 - p;
if p<0
    p = 0;
end
```
