## Extended Data 1 for "Scale space calibrates present and subsequent spatial learning in Barnes maze in mice": p4F.m

```
function [p] = p4F(Fdata,dfdata1,dfdata2,varargin)
% 
% p4F: 与えられたF値が臨界値となるようなp値（上側確率）
% 
% [p] = p4F(F,df1,df2,TOL)
%    p     F分布における、与えられたF値から無限大までの累積確率
%    F     F値
%    df1   分子の自由度
%    df2   分母の自由度
%    TOL   求積のための絶対誤差許容範囲
% 
% F, df1, df2 いずれもスカラーを与えて下さい。
% 
% p値はMATLABの数値積分関数（quadl）を利用して計算していま
% す。引数TOLはその際の誤差許容範囲です。詳しくはquadlのヘ
% ルプを参照して下さい。TOLは省略されるとデフォルト値とし
% て1e-06が用いられます。
% 
% see also:   fcdf (Statistics Toolbox)
% 
% (2008/02/10, by R. NIIMI)

p = [];

TOL = 1e-06; % default value
if 1-isempty(varargin)
    TOL = varargin{1};
end
if isempty(TOL)
    TOL = 1e-06;
else
    TOL = TOL(1);
end

if Fdata<0
    disp('  [p4F]:  F must be larger than zero.');
    return;
end
if 1 - isscalar(Fdata)
    disp('  [p4F]:  F must be a scalar.');
    return;
end
if 1 - (isscalar(dfdata1) & isscalar(dfdata2))
    disp('  [p4F]:  df1 and df2 must be scalars.');
    return;
end
if Fdata==0
    p = 1;
    return;
end

A = dfdata1^(dfdata1/2) * dfdata2^(dfdata2/2) / beta(dfdata1/2,dfdata2/2);
if dfdata1>=2
    Fdist = @(F) A .* (F.^(dfdata1/2-1) .* (dfdata2+dfdata1.*F).^((dfdata1+dfdata2)/(-2)));
else % To avoid y=+Inf within the region of quadrature, the function is transformed.
    Fdist = @(u) A .* (dfdata2 + dfdata1 .* (u.*dfdata1/2).^(2/dfdata1)).^((dfdata1+dfdata2)/(-2));
    Fdata = Fdata.^(dfdata1/2) *2/dfdata1;
end
p = quadl(Fdist,0,Fdata,TOL);
p = 1 - p;
if p<0
    p = 0;
end
```
