## Extended Data 1 for "Scale space calibrates present and subsequent spatial learning in Barnes maze in mice": p4ztest.m

```
function [p] = p4ztest(zdata,varargin)
% 
% p4ztest: z検定において、与えられたz値が臨界値となるようなp値（両側確率）
% 
% [p] = p4ztest(z,d)
%    p     標準正規分布における、与えられたzの絶対値から+∞までの累積確率の2倍
%    z     z値（行列可）
%    d     求積のための分割幅
% 
% 片側検定の場合は、pを2で割った値を使って下さい。
% 
% p値は合成シンプソン公式による数値積分で計算しています。
% 引数 d を小さくするとp値はより正確になりますが、計算時間や
% メモリ使用量が増えます。
% 
% dは省略できます。省略すると、デフォルト値として1e-04が用い
% られます。なお、dの値に比して積分区間が非常に小さく、積分
% 区間の分割数が100未満になるデータに対しては、分割数が100に
% なるような分割幅がdのかわりに用いられます。
% 
% p値の1.e-06以下の桁の値は、正確でない可能性があります。
% 
% メモリ不足が発生する場合は、zに行列データを与えるのをやめて
% スカラーを与える、dを大きくする、などして下さい。
% 
% see also:   normcdf (Statistics Toolbox)   erf
% 
% (2009/02/10, by R. NIIMI)

p = [];

d = 1e-04;
if 1-isempty(varargin)
    d = varargin{1};
end
if isempty(d)
    d = 1e-04;
else
    d = d(1);
end

data = abs(zdata);
data = data(1:numel(data));
Ndist = @(z) exp((z.^2)/(-2))/sqrt(2*pi);

x = [0:d:max(data)];
n1 = length(x);
Mask = ones(length(data),1) * x;
Mask2 = data' * ones(1,n1) + (d/2);
Mask = (Mask < Mask2);
clear Mask2;
y =  Ndist(x);
y = ones(length(data),1) * y;
y = y .* Mask;
n2 = sum(Mask,2);
clear Mask;
y = y';
p = y(1,:) + y(n1*[0:(length(data)-1)] + n2');
y(n1*[0:(length(data)-1)] + n2') = data*0;
p = p + 4*sum(y(2:2:(n1-1),:)) + 2*sum(y(3:2:(n1-1),:));
p = p * d / 3;

% If division number for the quadrature is less than 100...
SmallDiv = find(n2<100);
if 1-isempty(SmallDiv)
    div = 100;
    data2 = data(SmallDiv);
    n = length(data2);
    d2 = data2 / div;
    x = (ones(div,1) * data2) .* ([1:div]' * ones(1,n)) ./ div;
    x = [zeros(1,n); x];
    y = Ndist(x);
    p2 = y(1,:) + y(div+1,:) + 4*sum(y([2:2:div],:)) + 2* sum(y([3:2:div],:));
    p2 = p2 .* d2 / 3;
    p(SmallDiv) = p2;
end

p = 1 - (p*2); % p for two-tailed tests
I = find(p<0);
p(I) = zeros(size(I));
zdata = zeros(size(zdata));
zdata(1:numel(zdata)) = p;
p = zdata;
```
