## Extended Data 1 for "Scale space calibrates present and subsequent spatial learning in Barnes maze in mice": pr.m

```
function [Report,R,N,A,B,t,p,df] = pr(Data,varargin)

% pr: ピアソンの積率相関係数Rの算出と無相関検定，直線回帰
% 
% [Report,R,N,A,B,t,p,df] = pr(Data,wannaplot)
%    Data       入力データ。2列の行列。1列目をX, 2列目をYとする。
%    wannaplot  散布図を表示する場合は1,しない場合は0を与えて下さい。
%    Report     結果の一覧
%    R          相関係数
%    N          サンプル数
%    A          回帰直線の傾き
%    B          回帰直線の切片
%    t          無相関検定の結果，観測されたt値
%    p          上記t値が観測される両側確率
%    df         上記tの自由度
% 
% 入力データ（Data）の相関係数を算出し、無相関検定を行い、直線回帰を行い
% ます。引数wannaplotが1のときは、散布図が別ウィンドウで表示されます。
% wannaplotは省略可です。
% 
% この関数は無相関検定のために関数p4ttest.mを必要とします。
%
% see also:   prtest_1   prtest_2
% 
% (2009/01/27, by R. NIIMI)

Report = ['']; R = []; N = []; A = []; B = []; t = []; p = []; df = [];

if size(Data,2)~=2
    disp('  [pr]:  Data must be a N by 2 matrix.');
    return;
end

wannaplot = 0;
if 1-isempty(varargin)
    wannaplot = varargin{1}(1);
end

% Rの算出
X = Data(:,1);
Y = Data(:,2);
N = size(Data,1);
S = sum(Data);
R = ( N*(sum(X.*Y)) - S(1)*S(2) ) / sqrt(N*sum(X.^2)-S(1)^2) /sqrt(N*sum(Y.^2)-S(2)^2);
% R2 = R^2;

% 無相関検定
t = R*sqrt(N-2)/sqrt(1-R^2);
df = N-2;
p = p4ttest(t,df);

% 直線回帰
A = [X ones(N,1)] \ Y; % A = polyfit(X,Y,1);
B = A(2); A = A(1);

% 散布図の表示
if wannaplot ==1
    MarginX = std(X)/2;
    MarginY = std(Y)/2;
    Range = [min(X) max(X) min(Y) max(Y)] + [-MarginX MarginX -MarginY MarginY];
    plot(X,Y,'r.'), hold on;
    plot(Range(1:2),[A*Range(1:2)+B],'k-'), axis(Range);
    title(['R = ' num2str(R) ',  p = ' num2str(p)]);
end

% 結果の一覧
Report = ['  [pr]:  r = ' num2str(R) ',  t(' num2str(df) ') = ' num2str(t) ',  p = ' num2str(p) ' (two-tailed)'];
Report = strvcat(Report,['         Y = ' num2str(A) 'X + ' num2str(B)]);
```
