## Extended Data 1 for "Scale space calibrates present and subsequent spatial learning in Barnes maze in mice": prtest_1.m

```
function [Report,z,p] = prtest_1(rdata,N,rhyp)

% prtest_1: 1つの相関係数の仮説値との差の検定
% 
% [Report,z,p] = prtest_1(rdata,N,rhyp)
%    rdata   観測された相関係数（スカラー）
%    N       標本数（スカラー）
%    rhyp    仮説相関係数（スカラー）
%    Report  検定結果
%    z       観測されたz値
%    p       観測されたp値
% 
% 1つの相関係数rdataについて、これが仮説相関係数
% rhypを母相関係数とする母集団から得られたもので
% あるという帰無仮説について検定します。rdata, 
% N, rhypはいずれもスカラーです。
% 
% この検定は、相関係数をフィッシャーのz変換によって
% 変換した値が正規分布に近似的に従うことを利用して
% z検定により行われます。検定は両側です。
% 
% この関数は、関数 r2z.m と p4ztest.m を利用します。
% 
% see also:   pr   prtest_2
% 
% (2009/01/27, by R. NIIMI)

Report = ['']; z = []; p = [];

if ~isscalar(rdata)
    disp('  [prtest_1]:  rdata must be a scalar.');
    return;
end
if ~isscalar(rhyp)
    disp('  [prtest_1]:  rhyp must be a scalar.');
    return;
end
if ~isscalar(N)
    disp('  [prtest_1]:  N must be a scalar.');
    return;
end

if ((rdata<-1) | (rdata>1))
    disp('  [prtest_1]:  rdata must be -1 ~ 1.');
    return;
end
if ((rhyp<-1) | (rhyp>1))
    disp('  [prtest_1]:  rhyp must be -1 ~ 1.');
    return;
end
if round(N)~=N
    disp('  [prtest_1]:  N must be a integer.');
    return;
end
if N<1
    disp('  [prtest_1]:  N must be 1 or larger number.');
    return;
end

rdata = r2z(rdata);
rhyp  = r2z(rhyp);
z = (rdata-rhyp)*sqrt(N-3);
p = p4ztest(z);
Report = ['  [prtest_1]:  z = ' num2str(z) ',  p = ' num2str(p)];
```
