## Extended Data 1 for "Scale space calibrates present and subsequent spatial learning in Barnes maze in mice": prtest_2.m

```
function [Report,Xsq,p,df] = prtest_2(r,N)

% prtest_2: 複数の相関係数の差の検定
% 
% [Report,Xsq,p,df] = prtest_2(r,N)
%    r       相関係数（ベクトル）
%    N       標本数（ベクトル）
%    Report  検定結果
%    Xsq     観測されたカイ2乗値
%    p       観測されたp値
%    df      自由度
% 
% 互いに独立な標本から計算された複数の相関係数 r の
% 間に差があるかを調べるため、すべての標本の母相関
% 係数が等しいという帰無仮説について検定を行います。
% 検定は、計算されたXsqの値が近似的にカイ2乗分布に
% 従うことを利用して行われます。検定は両側です。
% 
% r は検定した複数の相関係数、N は r の元となった
% データの標本数です。両者は等しい要素数のベクトルで
% す。例えば50個の標本から観測された相関係数0.68と、
% これとは独立な60個の標本から観測された相関係数0.40
% の間に差があるかを検定するには下のようにします。
% prtest_2([0.68 0.40],[50 60])
% 
% この関数は、関数 r2z.m と p4chisq.m を利用します。
% 
% see also:   pr   prtest_1
% 
% (2009/01/27, by R. NIIMI)

Report = ['']; Xsq = []; p = []; df = [];

if ~isvector(r)
    disp('  [prtest_2]:  r must be a vector.');
    return;
end
if ~isvector(N)
    disp('  [prtest_2]:  N must be a vector.');
    return;
end

k = length(r);
if k~=length(N)
    disp('  [prtest_2]:  r and N must have identical number of element.');
    return;
end
if k<2
    disp('  [prtest_2]:  r and N must have 2 or more elements.');
    return;
end

if ~isempty(find((r<-1) | (r>1)))
    disp('  [prtest_2]:  r must be -1 ~ 1.');
    return;
end
if sum(round(N)==N)~=k
    disp('  [prtest_2]:  N must have integers.');
    return;
end
if sum(N>0)~=k
    disp('  [prtest_2]:  N must have positive numbers.');
    return;
end
if size(r,1)>1
    r = r';
end
if size(N,1)>1
    N = N';
end

z = r2z(r);
Xsq = sum((z.^2).*(N-3)) - (sum(z.*(N-3)).^2)/sum(N-3);
df = k-1;
p = p4chisq(Xsq,df);
Report = ['  [prtest_2]:  chi-square(' num2str(df) ') = ' num2str(Xsq) ',  p = ' num2str(p)];
```
