## Extended Data 1 for "Scale space calibrates present and subsequent spatial learning in Barnes maze in mice": r2z.m

```
function [z] = r2z(r)

% r2z: フィッシャーのz変換
% 
% [z] = r2z(r)
%    r   相関係数（-1 〜 +1）
%    z   変換されたz値（-Inf 〜 +Inf）
% 
% 入力された相関係数 r に対して、log((1+r)/(1-r))/2
% を計算します。r は行列可です。
% 
% rの元となったデータの標本数をNとすると、Nが十分に
% 大きいとき、zの分布は平均が0、分散が1/(N-3)の正規
% 分布に近似的に従うとされています。
% 
% see also:   z2r
% 
% (2015/09/03, by R. NIIMI)

z = [];

if ~isempty(find((r<-1) | (r>1)))
    disp('  [r2z]:  r must be -1 ~ 1.');
    return;
end

if r==1
	z = Inf;
else
	z = log((1+r)./(1-r))/2;
end
```
