## Extended Data 1 for "Scale space calibrates present and subsequent spatial learning in Barnes maze in mice": readme.txt

-----------------------------------------------------------------------------
私家版・心理実験データ統計解析のためのOctave関数集
しんりとうけい for Octave 略してsto (ver. 0.30)

■0. 目次
  1. 関数一覧
  2. 使い方
  3. 解析手法について
  4. 注意事項
  5. 作成および動作確認環境
  6. 参考文献
  7. 更新履歴メモ
  8. 連絡先

■1. 関数一覧

[t検定関連]
ttest_1      1標本のt検定
ttest_2      対応のない2標本のt検定、分散が等質な場合
ttest_2w     対応のない2標本のt検定、分散が等質でない場合（ウェルチの検定）
ttest_2p     対応のある2標本のt検定
bonf_ttest_2    ボンフェローニ法による多重比較、対比較はttest_2
bonf_ttest_2w   ボンフェローニ法による多重比較、対比較はttest_2w
bonf_ttest_2p   ボンフェローニ法による多重比較、対比較はttest_2p

[1要因分散分析関連]
anova_1b     対応のない1要因分散分析
anova_1w     対応のある1要因分散分析（データはリスト形式）
anova_1wm    対応のある1要因分散分析（データは集計表形式）
tukey_1b     対応のない1要因分散分析における、TukeyのHSD法による多重比較
tukey_1w     対応のある1要因分散分析における、TukeyのHSD法による多重比較
            （データはリスト形式）
tukey_1wm    対応のある1要因分散分析における、TukeyのHSD法による多重比較
            （データは集計表形式）

[2要因分散分析関連]
anova_2bb    2要因分散分析、2要因とも対応のない場合
anova_2bw    2要因分散分析、混合計画の場合
anova_2ww    2要因分散分析、2要因とも対応のある場合（データはリスト形式）
anova_2wwm   2要因分散分析、2要因とも対応のある場合（データは集計表形式）
tukey_2bb    2要因分散分析（2要因とも対応のない場合）での、TukeyのHSD法に
             よる多重比較
tukey_2bw    2要因分散分析（混合計画の場合）での、TukeyのHSD法による多重
             比較
tukey_2ww    2要因分散分析（2要因とも対応のある場合）での、TukeyのHSD法に
             よる多重比較（データはリスト形式）
tukey_2wm    2要因分散分析（2要因とも対応のある場合）での、TukeyのHSD法に
             よる多重比較（データは集計表形式）
sme_2bb      2要因分散分析（2要因とも対応のない場合）での単純主効果の検定
sme_2bw      2要因分散分析（混合計画の場合）での単純主効果の検定)
sme_2ww      2要因分散分析（2要因とも対応のある場合）での単純主効果の検定
            （データはリスト形式）
sme_2wwm     2要因分散分析（2要因とも対応のある場合）での単純主効果の検定
            （データは集計表形式）
tukey_sme_2bb   2要因分散分析（2要因とも対応のない場合）での、単純主効果に
                ついてのTukeyのHSD法による多重比較
tukey_sme_2bw   2要因分散分析（混合計画の場合）での、単純主効果についての
                TukeyのHSD法による多重比較
tukey_sme_2ww   2要因分散分析（2要因とも対応のある場合）での、単純主効果に
                ついてのTukeyのHSD法による多重比較（データはリスト形式）
tukey_sme_2wwm  2要因分散分析（2要因とも対応のある場合）での、単純主効果に
                ついてのTukeyのHSD法による多重比較（データは集計表形式）

[相関係数関連]
pr              ピアソンの積率相関係数Rの算出と無相関検定、直線回帰
prtest_1        1つの相関係数の仮説値との差の検定
prtest_2        複数の相関係数の差の検定
r2z             フィッシャーのz変換
z2r             フィッシャーのz変換の逆

[カイ2乗検定]
chisqtest_fit   適合度のカイ2乗検定
chisqtest_ind   独立性のカイ2乗検定
bonf_chisqtest_ind   ボンフェローニ法による多重比較、対比較はchisqtest_ind

[その他の検定等]
Utest           マン・ホイットニーのU検定
Htest           クラスカル・ウォリスのH検定
Ftest           F検定（2つの分散の差の検定）
binom_c         二項分布の累積確率

[信号検出理論]
dprime          d'の計算
fitroc          ROC曲線のあてはめとd'の推定
roc             ROC曲線上で誤警報率に対応するヒット率の計算
rocinv          ROC曲線上でヒット率に対応する誤警報率の計算
aprime          A'の計算

[ユーティリティ]
asintr       変数の逆正弦変換
matrix2list  集計表形式のデータをリスト形式に変換
list2matrix  リスト形式のデータを集計表形式に変換
list2cell    リスト形式のデータをセル配列に変換
sumoflist    リスト形式のデータの集計
findoutlier  リスト形式のデータから外れ値を検出する

[その他]
　主に他の関数が内部で呼び出して利用するために作成したものですが、単体で
使うこともできます。詳しくはヘルプを参照して下さい。
p4ztest      z検定において、与えられたz値が臨界値となるようなp値（両側確率）
p4ttest      t検定において、与えられたt値が臨界値となるようなp値（両側確率）
p4F          与えられたF値が臨界値となるようなp値（上側確率）
p4chisq      与えられたカイ2乗値が臨界値となるようなp値（上側確率）
studrange_5  スチューデント化された範囲の5%点（TukeyのHSD法におけるq値）
studrange_1  スチューデント化された範囲の1%点（TukeyのHSD法におけるq値）
bonf         ボンフェローニ法による多重比較
num2rank     数値に順位をつける
interpNev    関数のNeville補間

[ヘルプ]
helpsto      このreadme.txtファイルの中身を表示します

■2. 使い方
　普通の関数と同じです。適当な名前のディレクトリに置いて、そこにカレント
ディレクトリを移動するかパスを通すかして使って下さい。なお、いくつかの関数
は内部で他の関数を呼び出して使用しますので（例：tukey_1b は内部で anova_1b 
を走らせます）、みんな同じディレクトリに置いておかないと動作しない可能性が
あります。
　各関数の使い方については、それぞれのヘルプを参照して下さい（例：Octaveの
コンソールで help ttest_2 と入力）。また、help [ディレクトリ名] と打つと、
関数一覧を見ることができます（例： help sto ）。この readme.txt の内容は、
helpsto コマンドによってコンソールから見ることができます。

■3. 解析手法について
　t検定や分散分析の計算手続きは文献[1]に、信号検出理論に関しては文献[4]に、
ボンフェローニ法に関しては文献[5]に主に依拠しています。検定結果については、
文献[1]の例題などを与えて検算しています。統計量やp値の計算結果は、各種文献
に掲載の例題の答えや数表の値、MATLAB の Statistics Toolbox、R、ANOVA4 on 
the web, Excel などの出力とつき合わせてチェックしています。
　なお、分散分析で各セルのデータ数が等しくない場合には、SSの算出方法にいく
つかの方法があり、どれを用いるかによって結果が異なります。そのため、他の統
計ソフトとやや異なる結果が出力される場合があります。

■4. 注意事項
　統計学者ではない個人が自分で使う用に作成したものですので、結果の正確さに
保証はありません。ご自身の責任においてご利用ください。市販の統計パッケージ
が利用可能な方は、その利用も検討して下さい。
　各種検定のp値は近似的に算出しています。1.e-06以下の桁の非常に小さなp値は
正確でないと思われます。p = 5.6247e-009 のような値が出てきても、その数値自
体は気にせず、p値が心理実験で一般的な有意水準（5%とか1%）に比べて「非常に
小さかった」ということが結果であると考えてください。

■5. 作成および動作確認環境
・GNU Octave 3.2.4
・OS: Ubuntu 12.04LTS

■6. 参考文献
[1] 森　敏昭・吉田寿夫（編著） 1990　心理学のためのデータ解析テクニカル
        ブック　北大路書房
[2] 東京大学教養学部統計学教室（編） 1991　統計学入門　東京大学出版会
[3] 山内二郎（編） 1972　統計数値表　日本規格協会
[4] Gescheider, G. A. 1997. Psychophysics: the fundamentals. 3rd ed. 
        New York: Lawrence Arlbaum Associates.
        （ゲシャイダー，G. A.　宮岡　徹（監訳） 2002-03　心理物理学 ―方
        法・理論・応用―　北大路書房　2巻）
[5] 永田　靖・吉田道弘　1997　統計的多重比較法の基礎　サイエンティスト社
[6] 大久保街亜・岡田謙介　2012　伝えるための心理統計　勁草書房
[7] Cohen, J. 1988. Statistical power analysis for the behavioral 
        sciences. 2nd ed. Hillsdale, NJ: Lawrence Arlbaum Associates.
[8] Pashler, H. & Wixted, J. (eds.) 2002. Stevens's handbook of experi-
        mental psychology, 3rd ed., Vol.4 Methodology in experimental
        psychology.

■7. 更新履歴メモ
2015/9/22 ver. 0.30
　・stm から sto に改名しました。
　・文字コードをUTF-8に変更しました。
　・分散分析で効果量（偏イータ2乗）を算出するようにしました。
　・t検定で効果量（Cohen's d）を算出するようにしました（ttest_2wにおいて
　　はGlass' Deltaも）。
　・ttest_2pは、内部でttest_1を利用して動作するよう変更しました。
　・r2zでrが1のとき0除算でエラーが出るのを修正しました。
　・z2rでzが大きな値のとき結果がNaNとなるのを修正しました。
　・helpsto（旧 helpstm）を修正しました。
　・いくつかの関数でヘルプの内容を修正しました。
2009/04/14 ver. 0.21
  ・aprimeの計算式が間違っていたので修正しました。
  ・fitrocのヘルプの文を修正しました。
2009/03/01 ver. 0.2
  ・webで公開しました。
  ・GNU Octaveに対応しました（Octave 3.0.3, Windows XPにて動作確認）。
  ・信号検出理論に関する関数群（dprime, fitroc, roc, rocinv, aprime）が新
    しく追加されました。
  ・ボンフェローニ法による多重比較を行う関数群（bonf_ttest_2, bonf_ttest_2w,
    bonf_ttest_2p, bonf_chisqtest_ind, bonf）が新しく追加されました。また、
    この検定内で利用するために、関数list2cellが新しく追加されました。
  ・相関係数に関する検定（関数prtest_1, prtest_2）および、この検定内で
    行うフィッシャーのz変換（関数r2z）とその逆変換（関数z2r）が新しく追加
    されました。
  ・マン・ホイットニーのU検定（関数Utest）が新しく追加されました。
  ・クラスカル・ウォリスのH検定（関数Htest）が新しく追加されました。
  ・上記Htest内で用いるため、数値に順位をつける関数（num2rank）が新しく
    追加されました。
  ・sumoflistで、平均値のかわりに中央値も出力できるようにしました。
  ・p4ztest, p4ttestについて、p値の計算をMATLABのquadl関数を利用する方法か
    ら合成シンプソン公式による計算に変更しました。また、行列変数を入力でき
    るようにしました。
  ・p4F, p4chisqについて、自由度が2未満のときにOctaveで動作させると積分に
    失敗するバグを修正しました。また、絶対誤差許容範囲を引数で指定できるよ
    うにしました。
  ・binom_cを修正し，計算精度と速度を改善しました。
  ・Ftestで，F値の計算過程に誤りがあったのを修正しました。
  ・ほとんどの関数で、入力データが正しくない形式のときにエラーメッセージを
    返すようにしました。
  ・その他、いくつかのバグを修正しました。
2008/10/23 ver. 0.1

■8. 連絡先等
詳しくは http://rnpsychology.org/sto/ もご覧下さい。
お問い合わせ等は info //at// rnpsychology.org までお願いします。

                                     (Last modified: 2015/09/22, by R. NIIMI)
-----------------------------------------------------------------------------
