## Extended Data 1 for "Scale space calibrates present and subsequent spatial learning in Barnes maze in mice": roc.m

```
function [HR] = roc(FA,d,varargin)
% 
% roc: ROC曲線上で誤警報率に対応するヒット率の計算
% 
% [HR] = roc(FA,d,sdratio,wannaplot)
%    HR         ヒット率
%    FA         誤警報率
%    d          d'値（N分布の標準偏差を単位とする）
%    sdratio    N分布の標準偏差に対するSN分布の標準偏差の比
%    wannaplot  ROC曲線をプロットするには1を指定して下さい
% 
% 与えられたdとsdratioに基づいてROC曲線を決定し、その曲線において
% 与えられた誤警報率FAに対応するヒット率HRを計算します。逆にヒット
% 率から誤警報率を計算するにはrocinvを用いて下さい。
% 
% FAおよびdにはスカラーか等しい要素数のベクトルを指定します。ただ
% し、FAがベクトルでdがスカラーの場合、FAのすべての要素について
% 同じdの値に基づいて（つまり、単一のROC曲線に基づいて）対応する
% ヒット率を計算します。FAとdがともにベクトルの場合、FAの第n要素に
% 対応するヒット率は、dの第n要素に基づいて計算されます。
% 
% sdratioはスカラーです。sdratioは省略された場合、1が指定されたも
% のとみなされます（つまり、両分布は等分散であると仮定します）。
% 
% wannaplotに1が指定されると、ROC曲線をグラフにプロットします。省
% 略されるとプロットしません。
% 
% see also:   rocinv   dprime   fitroc
% 
% (2009/02/23, by R. NIIMI)

HR = [];

if 1-isvector(d)
    disp('  [roc]:  d should be a scalar or vector.');
    return;
end
if 1-isvector(FA)
    disp('  [roc]:  FA should be a scalar or vector.');
    return;
end
Check = find((FA<0) | (FA>1));
if 1-isempty(Check)
    disp('  [roc]:  FA should be 0~1.');
    return;
end

sdratio = 1;
wannaplot = 0;
if length(varargin)>0
    sdratio = varargin{1};
    if length(sdratio)>1
        disp('  [roc]:  sdratio should be a scalar.');
        return;
    end
    if sdratio<=0
        disp('  [roc]:  sdratio should be larger than zero.');
        return;
    end
    if isempty(sdratio)
        sdratio = 1;
    end
    if length(varargin)>1
        wannaplot = varargin{2}(1);
    end
    if isempty(wannaplot)
        wannaplot = 0;
    end
end

L = length(FA);
if length(d)==1 & L>1
    d = ones(1,L)*d;
elseif length(d)~=L
    disp('  [roc]:  Numbers of elements of FA and d are not equal.');
    return;
end

Zn  = erfinv((1-FA)*2-1) * sqrt(2);
Zsn = (d - Zn) / sdratio;
HR  = (erf(Zsn/sqrt(2))+1)/2;

if wannaplot==1
    plotFA  = [0.001 0.005 ([1:99]/100) 0.995 0.999];
    plotZn  = erfinv((1-plotFA)*2-1) * sqrt(2);
    plotZsn = d'*ones(1,103) - ones(length(d),1)*plotZn;
    plotZsn = plotZsn / sdratio;
    plotHR  = (erf(plotZsn/sqrt(2))+1)/2;
    plotFA = [0 plotFA 1];
    plotHR = [zeros(length(d),1) plotHR ones(length(d),1)];
    plot([0 1],[0 1],'k-'), hold on;
    plot(plotFA,plotHR,'r-'), hold on;
    plot(FA,HR,'b.'), hold on;
    axis square;
    xlim([0 1]); ylim([0 1]);
    xlabel('False Alarm'); ylabel('Hit');
end
```
