## Extended Data 1 for "Scale space calibrates present and subsequent spatial learning in Barnes maze in mice": sme_2bb.m

```
function [Report,Level,M,N,F,p,MS,df] = sme_2bb(Data,Fct)
% 
% sme_2bb:  2要因分散分析（2要因とも対応のない場合）での単純主効果の検定
% 
% [Report,Level,M,N,F,p,MS,df] = sme_2bb(Data,Fct)
%    Data    対象となるデータ（3列の行列）
%    Fct     検定する要因はどちらかを、下のように指定します。
%             1 -> 要因Bの各水準における要因Aの単純主効果の検定
%             2 -> 要因Aの各水準における要因Bの単純主効果の検定
%    Report  分析結果の一覧表
%    Level   水準番号の一覧（セルアレイ）
%    M       各条件の平均値
%    N       サンプル数
%    F       F値
%    p       p値
%    MS      平均平方
%    df      自由度
% 
% 入力データ（Data）は、1列目に要因Aの水準番号、2列目に要因Bの
% 水準番号、そして3列目に測定値が書かれた行列（anova_2bb.mに準
% じる）。
% 
% see also:   anova_2bb   sme_2ww   sme_2bw   tukey_2bb
% 
% (2008/12/25, by R. NIIMI)

Report = ['']; Level = []; M = []; N = []; F = []; p = []; MS = []; df = [];

if ~(size(Data,1)>1 & size(Data,2)==3)
    disp('  [sme_2bb]:  Data must be a N by 3 matrix.');
    return;
end

%%%%%% ここからanova_2bb.mと共通コード %%%%%%
Nall = size(Data,1);
Level{1} = min(Data(:,1));
for k=min(Data(:,1))+1:max(Data(:,1))
   if sum(Data(:,1)==k) > 0
       Level{1} = [Level{1} k];
   end
end
Level{2} = min(Data(:,2));
for k=min(Data(:,2))+1:max(Data(:,2))
   if sum(Data(:,2)==k) > 0
       Level{2} = [Level{2} k];
   end
end
Dat = [];
NL = [length(Level{1}) length(Level{2})]; % 水準数
for k=1:NL(1)
    for l=1:NL(2)
        I = find(Data(:,1)==Level{1}(k) & Data(:,2)==Level{2}(l));
        N(k,l) = length(I);
        M(k,l) = sum(Data(I,3));
%        Dat(:,l,k) = Data(I,3); % データを3次元（S*B*A）の集計表形式にする
    end
end

ABS = sum(Data(:,3).^2);
AB = sum(sum((M.^2)./N));
M = M./N; % 各セルの平均値
G = sum(sum(M));
X = G*G / (NL(1)*NL(2));
A = NL(2) * sum(mean(M').^2);
B = NL(1) * sum(mean(M).^2);
% AB = sum(sum(M.^2));
Ntil = NL(1)*NL(2) / sum(sum(1./N));

%%%%%% ここまでanova_2bb.mと共通コード %%%%%%

dfa = NL(1) - 1;
dfb = NL(2) - 1;
dfwc = Nall - NL(1)*NL(2);
df = [dfa dfb dfwc]';
SSwc = ABS - AB;
MSwc = SSwc / dfwc;

if Fct==2
    M = M';
elseif Fct~=1
    disp('  [sme_2bb]:  Fct must be 1 or 2.');
    return;
end
SS = Ntil * (sum(M.^2) - (sum(M).^2)/NL(Fct));
if Fct==2
    M = M';
end
MS = SS / df(Fct);
F = MS / MSwc;
for k=1:NL(3-Fct)
    p(k) = p4F(F(k),df(Fct),df(3));
end
SS = [SS SSwc];
MS = [MS MSwc];
df = [ones(1,NL(3-Fct))*df(Fct) dfwc];

sp = char(ones(NL(3-Fct)+2,3)*32);
if Fct==1
    F1 = 'A';
    F2 = 'b';
    temp = Level{2};
elseif Fct==2
    F1 = 'B';
    F2 = 'a';
    temp = Level{1};
end
Report = 'Source';
for k=1:NL(3-Fct)
    Report = strvcat(Report,[F1 '(' F2 num2str(temp(k)) ')']);
end
Report = strvcat(Report,'Error');
Report = [Report sp strvcat('SS',num2str(SS')) sp strvcat('df',num2str(df'))];
Report = [Report sp strvcat('MS',num2str(MS')) sp strvcat('F',num2str(F'), ' ') sp strvcat('p',num2str(p'), ' ')];
temp = char(ones(1,size(Report,2))*45);
Report = [temp; Report(1,:); temp; Report(2:NL(3-Fct)+1,:); temp; Report(NL(3-Fct)+2,:); temp];
Report = [char(ones(size(Report,1),3)*32) Report];
```
