## Extended Data 1 for "Scale space calibrates present and subsequent spatial learning in Barnes maze in mice": sme_2ww.m

```
function [Report,Level,M,N,F,p,MS,df] = sme_2ww(Data,Fct)
% 
% sme_2ww:  2要因分散分析（2要因とも対応のある場合）での単純主効果の検定（データはリスト形式）
% 
% [Report,Level,M,N,F,p,MS,df] = sme_2ww(Data,Fct)
%    Data    対象となるデータ（3列の行列）
%    Fct     検定する要因はどちらかを、下のように指定します。
%             1 -> 要因Bの各水準における要因Aの単純主効果の検定
%             2 -> 要因Aの各水準における要因Bの単純主効果の検定
%    Report  分析結果の一覧表
%    Level   水準番号の一覧（セルアレイ）
%    M       各水準の平均値
%    N       サンプル数（e.g.,参加者数）
%    F       F値
%    p       p値
%    MS      平均平方
%    df      自由度
% 
% 入力データ（Data）は、1列目に要因Aの水準番号、2列目に要因Bの
% 水準番号、そして3列目に測定値が書かれた行列（anova_2ww.mに準
% じる）。
% 
% see also:   anova_2ww   anova_2wwm   sme_2bb   sme_2bw
% 
% (2008/12/25, by R. NIIMI)

Report = ['']; Level = []; M = []; N = []; F = []; p = []; MS = []; df = [];

if ~(size(Data,1)>1 & size(Data,2)==3)
    disp('  [sme_2ww]:  Data must be a N by 3 matrix.');
    return;
end

%%%%%% ここからanova_2ww.mと共通コード %%%%%%
Nall = size(Data,1);
Level{1} = min(Data(:,1));
for k=min(Data(:,1))+1:max(Data(:,1))
   if sum(Data(:,1)==k) > 0
       Level{1} = [Level{1} k];
   end
end
Level{2} = min(Data(:,2));
for k=min(Data(:,2))+1:max(Data(:,2))
   if sum(Data(:,2)==k) > 0
       Level{2} = [Level{2} k];
   end
end
Dat = [];
NL = [length(Level{1}) length(Level{2})]; % 水準数
N = Nall / (NL(1) * NL(2));
for k=1:NL(1)
    for l=1:NL(2)
        I = find(Data(:,1)==Level{1}(k) & Data(:,2)==Level{2}(l));
        M(k,l) = sum(Data(I,3));
        Dat(:,l,k) = Data(I,3); % データを3次元（S*B*A）の集計表形式にする
    end
end

ABS = sum(sum(sum(Dat.^2)));
X = sum(sum(sum(Dat))).^2 / (N*NL(1)*NL(2));
A = sum(sum(M').^2) / (N*NL(2));
B = sum(sum(M).^2) / (N*NL(1));
AB = sum(sum(M.^2)) / N;
S = sum(sum(sum(Dat,2),3).^2) / (NL(1)*NL(2));
AS = sum(sum(sum(Dat,2).^2)) / NL(2);
BS = sum(sum(sum(Dat,3).^2)) / NL(1);

SS = [
    S - X; % SSs
    A - X; % SSa
    AS - A - S + X; % SSas
    B - X; % SSb
    BS - B - S + X; % SSbs
    AB - A - B + X; % SSab
    ABS - AB - AS - BS + A + B + S - X; % SSabs
    ABS - X; % SSt
];

dfs = N - 1;
dfa = NL(1) - 1;
dfas = dfa * dfs;
dfb = NL(2) - 1;
dfbs = dfb * dfs;
dfab = dfa * dfb;
dfabs = dfas * dfb;
dft = Nall - 1;
df = [dfs dfa dfas dfb dfbs dfab dfabs dft]';

%%%%%% ここまでanova_2ww.mと共通コード %%%%%%

df = [df(Fct*2) df(Fct*2+1)+df(7)];
MSpool = (SS(Fct*2+1)+SS(7)) / df(2);

if Fct==1
    M = M / N;
elseif Fct==2
    M = M' / N;
else
    disp('  [sme_2ww]:  Fct must be 1 or 2.');
    return;
end
SS = (sum(M.^2) - (sum(M).^2)./NL(Fct)) * N;
if Fct==2
    M = M';
end
MS = SS / df(1);
F = MS / MSpool;
for k=1:NL(3-Fct)
    p(k) = p4F(F(k),df(1),df(2));
end
MS = [MS MSpool];
df = [ones(1,NL(3-Fct))*df(1) df(2)];

sp = char(ones(NL(3-Fct)+2,3)*32);
if Fct==1
    F1 = 'A';
    F2 = 'b';
    temp = Level{2};
elseif Fct==2
    F1 = 'B';
    F2 = 'a';
    temp = Level{1};
end
Report = 'Source';
for k=1:NL(3-Fct)
    Report = strvcat(Report,[F1 '(' F2 num2str(temp(k)) ')']);
end
Report = strvcat(Report,'Error');
Report = [Report sp strvcat('SS',num2str(SS'),' ') sp strvcat('df',num2str(df'))];
Report = [Report sp strvcat('MS',num2str(MS')) sp strvcat('F',num2str(F'), ' ') sp strvcat('p',num2str(p'), ' ')];
temp = char(ones(1,size(Report,2))*45);
Report = [temp; Report(1,:); temp; Report(2:NL(3-Fct)+1,:); temp; Report(NL(3-Fct)+2,:); temp];
Report = [char(ones(size(Report,1),3)*32) Report];
```
