## Extended Data 1 for "Scale space calibrates present and subsequent spatial learning in Barnes maze in mice": studrange_1.m

```
function [q] = studrange_1(r,df)
% 
% studrange_1: ステューデント化された範囲の1%点（TukeyのHSD法におけるq値）
% 
% q = studrange_1(r,df)
%     q   与えられたr, dfにおけるqの臨界値（有意水準=1%）
%     r   多重比較する平均値の数（要因の水準数）, 2〜30の整数
%     df  誤差項の自由度, 1〜infの整数
% 
% 山内編（1972）『統計数値表』の表D1による。表に含まれていないr, dfに
% ついては、Neville補間による補間計算結果（interpNev.m使用）を返す。
% 
% (2007/12/26, by R. NIIMI)

qTable = [
% r= 2       3       4      5       6       8      10      15      20      30      
90.0242	135.0407	164.2577	185.5753	202.2097	227.1663	245.5416	277.0034	297.9972	325.9682 %df=1
14.0358	19.0189	22.2937	24.7172	26.6290	29.5301	31.6894	35.4261	37.9435	41.3221 %   2
8.2603	10.6185	12.1695	13.3243	14.2407	15.6410	16.6908	18.5219	19.7648	21.4429 %   3
6.5112	8.1198	9.1729	9.9583	10.5832	11.5418	12.2637	13.5298	14.3939	15.5662 %   4
5.7023	6.9757	7.8042	8.4215	8.9131	9.6687	10.2393	11.2436	11.9318	12.8688 %   5
5.2431	6.3305	7.0333	7.5560	7.9723	8.6125	9.0966	9.9508	10.5378	11.3393 %   6
4.9490	5.9193	6.5424	7.0050	7.3730	7.9390	8.3674	9.1242	9.6454	10.3586 %   7
4.7452	5.6354	6.2038	6.6248	6.9594	7.4738	7.8632	8.5517	9.0265	9.6773  %   8
4.5960	5.4280	5.9567	6.3473	6.6574	7.1339	7.4945	8.1323	8.5726	9.1767  %   9
4.4820	5.2702	5.7686	6.1361	6.4275	6.8749	7.2133	7.8121	8.2256	8.7936  %  10
4.3198	5.0459	5.5016	5.8363	6.1011	6.5069	6.8136	7.3558	7.7305	8.2456  %  12
4.2099	4.8945	5.3215	5.6340	5.8808	6.2583	6.5432	7.0466	7.3943	7.8726  %  14
4.1306	4.7855	5.1919	5.4885	5.7223	6.0793	6.3483	6.8233	7.1512	7.6023  %  16
4.0707	4.7034	5.0942	5.3788	5.6028	5.9443	6.2013	6.6546	6.9673	7.3973  %  18
4.0239	4.6392	5.0180	5.2933	5.5095	5.8389	6.0865	6.5226	6.8232	7.2366  %  20
3.9555	4.5456	4.9068	5.1684	5.3735	5.6850	5.9187	6.3296	6.6123	7.0008  %  24
3.8891	4.4549	4.7992	5.0476	5.2418	5.5361	5.7563	6.1423	6.4074	6.7710  %  30
3.8247	4.3672	4.6951	4.9308	5.1145	5.3920	5.5989	5.9606	6.2083	6.5471  %  40
3.7622	4.2822	4.5944	4.8178	4.9913	5.2525	5.4466	5.7845	6.0149	6.3290  %  60
3.7016	4.1999	4.4970	4.7085	4.8722	5.1176	5.2992	5.6138	5.8272	6.1168  % 120
3.6428	4.1203	4.4028	4.6028	4.7570	4.9872	5.1566	5.4485	5.6452	5.9106  % inf
];

df1 = [1 2 3 4 5 6 7 8 9 10 12 14 16 18 20 24 30 40 60 120 inf];
r1 = [2 3 4 5 6 8 10 15 20 30];
df2 = 120./df1;
r2 = 1./r1;

q = [];
if (r<2) | (r>30)
    disp('  [studrange_1]:  q = studrange_1(r,df); r must be 2~30.');
    return;
elseif (df<1)
    disp('  [studrange_1]:  q = studrange_1(r,df); df must be 1 or larger.');
    return;
elseif (floor(r)~=r) | (floor(df)~=df) % unacceptable cases
    disp('  [studrange_1]:  q = studrange_1(r,df); r and df must be integer.');
    return;
elseif sum(r==r1) & sum(df==df1)
    q = qTable(find(df==df1),find(r==r1));
elseif sum(r==r1) & sum(df~=df1)
    I = max(find(df1 < df));
    I = [I-1:I+2];
    if max(I) > 21
        I = [18:21];
    end
    J = find(r==r1);
    q = interpNev(df2(I),qTable(I,J)',120/df);
elseif sum(r~=r1) & sum(df==df1)
    I = find(df==df1);
    J = max(find(r1 < r));
    J = [J-1:J+2];
    if max(J) > 10
        J = [7:10];
    end
    q = interpNev(r2(J),qTable(I,J),1/r);
else
    I = max(find(df1 < df));
    I = [I-1:I+2];
    if max(I) > 21
        I = [18:21];
    end
    J = max(find(r1 < r));
    J = [J-1:J+2];
    if max(J) > 10
        J = [7:10];
    end
    for k=1:4
        q2(k) = interpNev(df2(I),qTable(I,J(k))',120/df);
    end
    q = interpNev(r2(J),q2,1/r);
end
```
