## Extended Data 1 for "Scale space calibrates present and subsequent spatial learning in Barnes maze in mice": studrange_5.m

```
function [q] = studrange_5(r,df)
% 
% studrange_5: ステューデント化された範囲の5%点（TukeyのHSD法におけるq値）
% 
% q = studrange_5(r,df)
%     q   与えられたr, dfにおけるqの臨界値（有意水準=5%）
%     r   多重比較する平均値の数（要因の水準数）, 2〜30の整数
%     df  誤差項の自由度, 1〜infの整数
% 
% 山内編（1972）『統計数値表』の表D1による。表に含まれていないr, dfに
% ついては、Neville補間による補間計算結果（interpNev.m使用）を返す。
% 
% (2007/12/26, by R. NIIMI)

qTable = [
% r= 2       3       4      5       6       8      10      15      20      30      
17.9693	26.9755	32.8187	37.0815	40.4076	45.3973	49.0710	55.3607	59.5576	65.1490 %df=1
6.0849	8.3308	9.7980	10.8811	11.7343	13.0273	13.9885	15.6503	16.7688	18.2690 %   2
4.5007	5.9096	6.8245	7.5017	8.0371	8.8525	9.4620	10.5222	11.2400	12.2073 %   3
3.9265	5.0402	5.7571	6.2870	6.7064	7.3465	7.8263	8.6640	9.2334	10.0034 %   4
3.6354	4.6017	5.2183	5.6731	6.0329	6.5823	6.9947	7.7163	8.2080	8.8747  %   5
3.4605	4.3392	4.8956	5.3049	5.6284	6.1222	6.4931	7.1428	7.5864	8.1889  %   6
3.3441	4.1649	4.6813	5.0601	5.3591	5.8153	6.1579	6.7586	7.1691	7.7275  %   7
3.2612	4.0410	4.5288	4.8858	5.1672	5.5962	5.9183	6.4831	6.8694	7.3953  %   8
3.1992	3.9485	4.4149	4.7554	5.0235	5.4319	5.7384	6.2758	6.6435	7.1444  %   9
3.1511	3.8768	4.3266	4.6543	4.9120	5.3042	5.5984	6.1141	6.4670	6.9480  %  10
3.0813	3.7729	4.1987	4.5077	4.7502	5.1187	5.3946	5.8780	6.2089	6.6600  %  12
3.0332	3.7014	4.1105	4.4066	4.6385	4.9903	5.2534	5.7139	6.0290	6.4586  %  14
2.9980	3.6491	4.0461	4.3327	4.5568	4.8962	5.1498	5.5932	5.8963	6.3097  %  16
2.9712	3.6093	3.9970	4.2763	4.4944	4.8243	5.0705	5.5006	5.7944	6.1950  %  18
2.9500	3.5779	3.9583	4.2319	4.4452	4.7676	5.0079	5.4273	5.7136	6.1039  %  20
2.9188	3.5317	3.9013	4.1663	4.3727	4.6838	4.9152	5.3186	5.5936	5.9682  %  24
2.8882	3.4864	3.8454	4.1021	4.3015	4.6014	4.8241	5.2114	5.4750	5.8335  %  30
2.8582	3.4421	3.7907	4.0391	4.2316	4.5205	4.7345	5.1056	5.3575	5.6996  %  40
2.8288	3.3987	3.7371	3.9774	4.1632	4.4411	4.6463	5.0011	5.2412	5.5663  %  60
2.8000	3.3561	3.6846	3.9169	4.0960	4.3630	4.5595	4.8979	5.1259	5.4336  % 120
2.7718	3.3145	3.6332	3.8577	4.0301	4.2863	4.4741	4.7959	5.0117	5.3013  % inf
];

df1 = [1 2 3 4 5 6 7 8 9 10 12 14 16 18 20 24 30 40 60 120 inf];
r1 = [2 3 4 5 6 8 10 15 20 30];
df2 = 120./df1;
r2 = 1./r1;

q = [];
if (r<2) | (r>30)
    disp('  [studrange_5]:  q = studrange_5(r,df); r must be 2~30.');
    return;
elseif (df<1)
    disp('  [studrange_5]:  q = studrange_5(r,df); df must be 1 or larger.');
    return;
elseif (floor(r)~=r) | (floor(df)~=df) % unacceptable cases
    disp('  [studrange_5]:  q = studrange_5(r,df); r and df must be integer.');
    return;
elseif sum(r==r1) & sum(df==df1)
    q = qTable(find(df==df1),find(r==r1));
elseif sum(r==r1) & sum(df~=df1)
    I = max(find(df1 < df));
    I = [I-1:I+2];
    if max(I) > 21
        I = [18:21];
    end
    J = find(r==r1);
    q = interpNev(df2(I),qTable(I,J)',120/df);
elseif sum(r~=r1) & sum(df==df1)
    I = find(df==df1);
    J = max(find(r1 < r));
    J = [J-1:J+2];
    if max(J) > 10
        J = [7:10];
    end
    q = interpNev(r2(J),qTable(I,J),1/r);
else
    I = max(find(df1 < df));
    I = [I-1:I+2];
    if max(I) > 21
        I = [18:21];
    end
    J = max(find(r1 < r));
    J = [J-1:J+2];
    if max(J) > 10
        J = [7:10];
    end
    for k=1:4
        q2(k) = interpNev(df2(I),qTable(I,J(k))',120/df);
    end
    q = interpNev(r2(J),q2,1/r);
end
```
