## Extended Data 1 for "Scale space calibrates present and subsequent spatial learning in Barnes maze in mice": sumoflist.m

```
function [S] = sumoflist(Data,varargin)
% 
% sumoflist:  リスト形式のデータの集計
% 
% [S] = sumoflist(Data,Mtype)
%    Data    対象となるデータ（リスト形式）
%    Mtype   代表値の種類を指定します
%             1 -> 平均値（デフォルト）
%             2 -> 中央値
%    S       集計結果
% 
% リスト形式のデータを集計し、各セルのデータ数および代表値、標
% 準偏差の一覧表を出力します。Data は少なくとも2列以上の行列で
% ある必要があります。要因数は1以上で制限なし。Data の行の並び
% 順はランダムでかまいません。
% 
% 代表値の種類は引数Mtypeで指定します（平均値もしくは中央値）。
% Mtypeは省略可です。省略されると平均値を出力します。
% 
% 例えば下のようなデータ（1列目が要因Aの水準番号、2列目が要因Bの
% 水準番号、3列目が測定値）があったとします（水準番号は整数にし
% て下さい）。
% 
%   Data = [
%       1   1   24
%       1   1   18
%       1   2   30
%       1   2   26
%       1   1   25
%       1   2   30
%       0   1   15
%       0   2   13
%       0   1   18
%       0   1   14
%       0   2   11
%   ];
% 
% このデータを入力すると、下のような結果を返します。
% 
%   S = [
%       0    1    3   15.6667    2.0817
%       0    2    2   12.0000    1.4142
%       1    1    3   22.3333    3.7859
%       1    2    3   28.6667    2.3094
%   ];
% 
% 左列から順に要因Aの水準番号、要因Bの水準番号、そのセルのデータ
% 数、平均値、標準偏差です。各行は、水準番号の数値の小さい順に並
% びます。なお、ひとつもデータのないセルがあった場合には、データ
% 数・平均値・標準偏差としてすべて0を返します。
% 
% see also:   findoutlier   list2matrix   list2cell
% 
% (2009/02/16, by R. NIIMI)

S = [];

Nall = size(Data,1);
NF = size(Data,2)-1; % 要因数
if NF < 1
    disp('  [sumoflist]:  Data must contain 2 or more columns.');
    return;
end

Mtype = 1;
if ~isempty(varargin)
    if ~isempty(varargin{1})
        Mtype = varargin{1}(1);
    end
end
if sum(Mtype==[1 2])==0
    Mtype = 1;
end

% 水準番号の抽出
Level = [];
for F = 1:NF
    Level{F} = min(Data(:,F));
    for k=min(Data(:,F))+1:max(Data(:,F))
        if sum(Data(:,F)==k) > 0
           Level{F} = [Level{F} k];
        end
    end
    NL(F) = length(Level{F}); % 水準数
end

Ncell = prod(NL); % セル数
S = [];
for F = 1:NF-1
    temp = mod([0:Ncell-1],prod(NL(F:NF)))+1;
    temp = ceil(temp/prod(NL((F+1):NF)));
    S = [S temp'];
end
temp = mod([0:Ncell-1],NL(NF))+1;
S = [S temp'];
for F = 1:NF
    S(:,F) = Level{F}(S(:,F));
end
temp = [];
for C=1:Ncell
    F = find(sum(Data(:,1:NF)==(ones(Nall,1)*S(C,:)),2)==NF);
    if isempty(F)
        temp(C,1) = 0; % n
        temp(C,2) = 0; % mean/median
        temp(C,3) = 0; % std
    else
        temp(C,1) = length(F); % n
        if Mtype==1
            temp(C,2) = mean(Data(F,NF+1)); % mean
        elseif Mtype==2
            temp(C,2) = median(Data(F,NF+1)); % median
        end
        temp(C,3) = std(Data(F,NF+1)); % std
    end
end
S = [S temp];
```
