## Extended Data 1 for "Scale space calibrates present and subsequent spatial learning in Barnes maze in mice": ttest_1.m

```
function [Report,t,p,df,N,ES] = ttest_1(Data,M)
% 
% ttest_1:  1標本のt検定
% 
% [Report,t,p,df,N] = ttest_1(Data,M)
%    Data     対象となる標本（ベクトル）
%    M        仮説平均
%    Report   検定結果
%    t        観測されたt値
%    p        観測されたp値（両側）
%    df       自由度
%    N        標本サイズ
%    ES       効果量（Cohen's d）
% 
% 母分散がわからない場合の1つの平均値の検定です。標本が
% 仮説平均を母平均とする母集団から得られたものであるとい
% う帰無仮説のもとで、両側t検定を行います。
% 
% この関数は、関数 p4ttest を利用します。
% 
% (2015/09/11, by R. NIIMI)

Report = []; t = []; p = []; df = []; N = []; ES = [];

if 1-isvector(Data)
    disp('  [ttest_1]:  Data must be a vector.');
    return;
end

N = length(Data);
df = N - 1;
% S2 = sumsq(Data-mean(Data))/df;
t = (mean(Data)-M) / sqrt(sumsq(Data-mean(Data))/df/N); %t = (mean(Data)-M) / sqrt(var(Data)/N);
p = p4ttest(t,df);
ES = abs(mean(Data)-M) / sqrt(sumsq(Data-mean(Data))/N);
Report = ['  [ttest_1]: t(' num2str(df) ') = ' num2str(t) ', p = ' num2str(p) ' (two-tailed), Cohen''s d = ' num2str(ES)];
```
