## Extended Data 1 for "Scale space calibrates present and subsequent spatial learning in Barnes maze in mice": ttest_2.m

```
function [Report,t,p,df,N,ES] = ttest_2(A,B)
% 
% ttest_2:  対応のない2標本のt検定、分散が等質な場合
% 
% [Report,t,p,df,N,ES] = ttest_2(A,B)
%    A        標本1（ベクトル）
%    B        標本2（ベクトル）
%    Report   検定結果
%    t        観測されたt値
%    p        観測されたp値（両側）
%    df       自由度
%    N        標本サイズ
%    ES       効果量（Cohen's d）
% 
% この関数は、関数 p4ttest を利用します。
% 
% see also:   ttest_2w   ttest_2p   bonf_ttest_2   Ftest
% 
% (2015/09/11, by R. NIIMI)

Report = []; t = []; p = []; df = []; N = []; ES = [];

if (~isvector(A)) | (~isvector(B))
    disp('  [ttest_2]:  Please input vectors as data.');
    return;
end

N = [length(A) length(B)];
df = sum(N)-2;
M = [mean(A) mean(B)];
S = sum((A-M(1)).^2) + sum((B-M(2)).^2);
t = (M(1)-M(2)) / sqrt(S/df*((1/N(1))+(1/N(2))));
p = p4ttest(t,df);
ES = abs(M(1)-M(2)) / sqrt((sum((A-M(1)).^2) + sum((B-M(2)).^2)) / sum(N));
Report = ['  [ttest_2]: t(' num2str(df) ') = ' num2str(t) ', p = ' num2str(p) ' (two-tailed), Cohen''s d = ' num2str(ES)];
```
