## Extended Data 1 for "Scale space calibrates present and subsequent spatial learning in Barnes maze in mice": ttest_2p.m

```
function [Report,t,p,df,N,ES] = ttest_2p(Data)
% 
% ttest_2p:  対応のある2標本のt検定 (Paired t-test)
% 
% [Report,t,p,df,N,ES] = ttest_2p(Data)
%    Data     データ（2列の行列）
%    Report   検定結果
%    t        観測されたt値
%    p        観測されたp値（両側）
%    df       自由度
%    N        標本サイズ
%    ES       効果量（Cohen's d）
% 
% 入力データ（Data）は、N行2列からなる集計表形式の行列。
% すなわち、同一行の2つの数値が対応したデータです。
% 
% 効果量 ES は、対応のある2群の差 [Data(:,1) - Data(:,2)]
% の分散に基づいた分母を用いて算出されます。対応のないt検定
% と同様に2群それぞれの分散に基づいた分母を用いた効果量は、
% 同じデータを ttest_2 に与えることで算出できます。
% 
% この関数は、関数 ttest_1, p4ttest を利用します。
% 
% see also:   ttest_2   ttest_2w   bonf_ttest_sp
% 
% (2015/09/11, by R. NIIMI)

Report = []; t = []; p = []; df = []; N = []; ES = [];

if ~(size(Data,1)>1 & size(Data,2)==2)
    disp(' [ttest_2p]:  Data must be a N by 2 matrix.');
    return;
end

Data = Data(:,1) - Data(:,2);
[Report,t,p,df,N,ES] = ttest_1(Data,0);
%%%%% otherwise, following sentences do the paired t-test without using ttest_1 function.
% N = size(Data,1);
% df = N-1;
% D = Data(:,1)-Data(:,2);
% M = [mean(Data(:,1)) mean(Data(:,2))];
% t = sum(D) / sqrt( (N*sum(D.^2)-(sum(D))^2) / df );
% p = p4ttest(t,df);
% ES = abs(M(1)-M(2)) / sqrt(sumsq(D-mean(D))/N);
%%%%%

Report = ['  [ttest_2p]: t(' num2str(df) ') = ' num2str(t) ', p = ' num2str(p) ' (two-tailed), Cohen''s d = ' num2str(ES)];
```
