## Extended Data 1 for "Scale space calibrates present and subsequent spatial learning in Barnes maze in mice": ttest_2w.m

```
function [Report,t,p,df,N,ES] = ttest_2w(A,B,varargin)
% 
% ttest_2w:  対応のない2標本のt検定、分散が等質でない場合（ウェルチの検定）
% 
% [Report,t,p,df,N,ES] = ttest_2w(A,B,EStype)
%    A        標本1（ベクトル）
%    B        標本2（ベクトル）
%    EStype   算出する効果量の種類を指定します
%             ('cohen':Cohen's d, 'glass': Glass' Delta)
%    Report   検定結果
%    t        観測されたt値
%    p        観測されたp値（両側）
%    df       自由度
%    N        標本サイズ
% 
% 検定手法の性質上、自由度 df は小数になることがあります。
% 
% 引数 EStype には文字列で 'cohen' または 'glass' を指定して下さい。
% 省略可です。省略時にはデフォルト値として 'cohen' が用いられます。
% 
% EStype に 'cohen' を指定すると、ES として Cohen's d を報告します。
% 分母として、標本1と標本2の標本分散をNで重み付けた平均の平方根を使
% 用します。
% 
% EStype に 'glass' を指定すると、ES として Glass' Delta を報告しま
% す。このとき、ES は長さ2のベクトルとなり、ES(1) は分母に標本1の不
% 偏分散の平方根を用いた値、ES(2) は分母に標本2の不偏分散の平方根を
% 用いた値です。
% 
% この関数は、関数 p4ttest を利用します。
% 
% see also:   ttest_2   ttest_2p   bonf_ttest_2w   Ftest
% 
% (2015/09/11, by R. NIIMI)

Report = []; t = []; p = []; df = []; N = []; ES = [];

if (~isvector(A)) | (~isvector(B))
    disp('  [ttest_2w]:  Please input vectors as data.');
    return;
end
if isempty(varargin)
    EStype = 1;
elseif ~ischar(varargin{1})
    disp('  [ttest_2w]:  Invalid EStype');
    return;
else
    if strcmp(varargin{1},'cohen')
        EStype = 1;
    elseif strcmp(varargin{1},'glass')
        EStype = 2;
    else
        disp('  [ttest_2w]:  EStype must be ''cohen'' or ''glass''.');
        return;
    end
end

N = [length(A) length(B)];
M = [mean(A) mean(B)];
S = [sum((A-M(1)).^2) sum((B-M(2)).^2)]./N;
t = (M(1)-M(2)) / sqrt(sum(S./(N-1)));
df = sum(S./(N-1))^2 / sum(S.^2 ./ (N-1).^3);
p = p4ttest(t,df);

ES = abs(M(1)-M(2));
if EStype==1
    ES = ES / sqrt((sum((A-M(1)).^2) + sum((B-M(2)).^2)) / sum(N));
    Report = ['  [ttest_2w]:  t(' num2str(df) ') = ' num2str(t) ',  p = ' num2str(p) ' (two-tailed), Cohen''s d = ' num2str(ES)];
elseif EStype==2
    ES = [ES ES] ./ [sqrt(sum((A-M(1)).^2)/(N(1)-1)) sqrt(sum((B-M(2)).^2)/(N(2)-1))]; 
    Report = ['  [ttest_2w]:  t(' num2str(df) ') = ' num2str(t) ',  p = ' num2str(p) ' (two-tailed), Glass'' Delta = [' num2str(ES) ']'];
end
```
