## Extended Data 1 for "Scale space calibrates present and subsequent spatial learning in Barnes maze in mice": tukey_1b.m

```
function [Report,HSD5,HSD1,Result5,Result1,Md] = tukey_1b(Data)
% 
% tukey_1b: 対応のない1要因の分散分析における、TukeyのHSD法による多重比較
% 
% [Report,HSD5,HSD1,Result5,Result1,Md] = tukey_1b(Data)
%    Data    対象となるデータ（2列の行列）
%    Report  検定結果の表（* p < .05, ** p < .01）
%    HSD5    各比較における、有意水準5%でのHSD値
%    HSD1    各比較における、有意水準1%でのHSD値
%    Result5 各比較における、有意水準5%での検定結果（1:有意, 0:n.s.）
%    Result1 各比較における、有意水準1%での検定結果（1:有意, 0:n.s.）
%    Md      平均値の差の表
% 
% 入力データ（Data）は、anova_1b.mと同様、1列目に水準番号、2列目に測定値が
% 書かれた行列。すなわち、1行が1サンプル。行の順序はランダムでかまわない。
% 水準番号は整数（負も可）。
% 3水準のデータの例：
%       Data = [
%          1    12.3
%          3    15.1
%          2     9.9
%          3    18.7
%          2    14.0
%          ...  ...
%       ];
% 
% HSD5, HSD1, Result5, Result1, Md は、水準数×水準数の行列となります。これ
% らの変数の行および列の順序は、水準番号の値の小さい順序となっています。
% 
% この関数は、関数 anova_1b, studragen_5, studrange_1 を利用します。
%  
% see also:   tukey_1w   tukey_1wm   anova_1b
% 
% (2015/09/03, by R. NIIMI)

Report = ['']; HSD5 = []; HSD1 = []; Result5 = []; Result1 = []; Md = [];

if ~(size(Data,1)>1 & size(Data,2)==2)
    disp('  [tukey_1b]:  Data must be a N by 2 matrix.');
    return;
end

[Report,Level,M,N,F,p,MS,df] = anova_1b(Data);
if isempty(p) % failure on anova_1b
    return;
end

NL = length(M); % 水準数
Md = (ones(NL,1) * M) - (M' * ones(1,NL));
ND = (1./(ones(NL,1) * N) +  1./(N' * ones(1,NL)))/2;
q5 = studrange_5(NL,df(2));
q1 = studrange_1(NL,df(2));
HSD5 = sqrt(MS(2)*ND) * q5;
HSD1 = sqrt(MS(2)*ND) * q1;

Result5 = abs(Md)>HSD5;
Result1 = abs(Md)>HSD1;

Sp = char(ones(NL,3)*32);
Report = [Sp char(ones(NL,1)*91) num2str(Level') char(ones(NL,1)*93)];
Report = [blanks(size(Report,2)); Report];
Sp = [blanks(3); Sp];
for j=1:NL
    NowCol = num2str(Md(:,j));
    temp = size(NowCol,2) - (length(num2str(Level(j)))+2);
    if temp>=0
        NowCol = [blanks(temp) '[' num2str(Level(j)) ']'; NowCol];
    else
        NowCol = ['[' num2str(Level(j)) ']'; [char(ones(NL,-temp)*32) NowCol]]; 
    end
    temp = [0; Result5(:,j) + Result1(:,j)];
    for i=1:NL+1
        Ast(i,:) = [char(ones(1,temp(i))*42)  blanks(2-temp(i))];
    end
    Report = [Report Sp NowCol Ast];
end
```
