## Extended Data 1 for "Scale space calibrates present and subsequent spatial learning in Barnes maze in mice": stats_ntw_v0.m

```
function [] = stats_ntw_v0(tbl,ntwObj,parameters)


phase = unique(tbl.Phase);


%% load data
names = fieldnames(ntwObj);
for i = 1:length(names)
    iT = getfield(ntwObj(i),names{i});
    eval([names{i}, '=iT;']);
end


%% 

if ismember(phase, {'training','retraining','reversal_training'})
        Data = tbl_trn_daily;
        ntws = Data.Properties.VariableNames(4:end);
        Days = unique(Data.Day);
elseif ismember(phase, {'probe_test'})
        Data = tbl_prb_daily;
        ntws = Data.Properties.VariableNames(3:end);
        Days = 1;
end


[groups,~,ic] = unique(Data.Group);

ntwsnames = {'Order','Shortest path length','Degree',...
        'Clustering coefficient','Density', 'Betweenness centrality', 'Closeness centrality',...
        'No. of stops','Order','Shortest path length','Degree',...
        'Clustering coefficient','Density','Betweenness centrality', 'Closeness centrality'};


y = cell(length(ntws)*length(Days),7);
yy = [];
n = 1;
for i = 1:length(ntws)
    for j = 1:length(Days)
        
        if ismember(phase, {'training','retraining','reversal_training'})
            I = Days(j)==tbl_trn_daily.Day;
            x = Data{I, ntws{i}};
        elseif ismember(phase, {'probe_test'})
            I = 1:length(ic);
            x = Data{:, ntws{i}};
        end
        
        switch ntws{i}(end)
            case '2'
                ngm = 'dynamic';
            case '1'
                ngm = 'static';
        end
        
        if length(groups)==2
            [p,~,stats] = ranksum(x(ic(I)==1),x(ic(I)==2),'method','approximate');
            r = stats.zval/sqrt(length(ic(I)));
            y(n,5) = {sprintf('z = %0.2f',stats.zval)};
            y(n,7) = {sprintf('r = %0.2f',r)};
            y(n,3) = {'Wilcoxon rank-sum test'};
        elseif length(groups)>2
            [p,statstable,~] = kruskalwallis(x, ic(I), 'off');
            y(n,5) = {sprintf('X^2 (%d) = %0.2f',length(groups)-1,statstable{2,5})};
            y(n,3) = {'Kruskall-Wallis test'};
            
            C = nchoosek(1:length(groups),2);
            yy_k = cell(size(C,1),8);
            for k = 1:size(C,1)
                [pp,~,stats] = ranksum(x(ic(I)==C(k,1)),x(ic(I)==C(k,2)));
                r = stats.zval/sqrt(sum(ismember(ic(I),C(k,:))));
                yy_k(k,1) = ntwsnames(i);
                yy_k(k,2) = {ngm};
                yy_k(k,3) = {'Wilcoxon rank-sum test'};
                yy_k(k,4) = {sprintf('%s vs. %s', groups{C(k,1)},groups{C(k,2)})};
                if ismember(phase, {'training','retraining','reversal_training'})
                    yy_k(k,5) = {Days(j)};
                elseif ismember(phase, {'probe_test'})
                    yy_k(k,5) = {''};
                end
                yy_k(k,6) = {sprintf('z = %0.2f',stats.zval)};
                yy_k(k,7) = {sprintf('%0.3f',pp)};
                yy_k(k,8) = {sprintf('r = %0.2f',r)};
            end
            yy = [yy; yy_k];
        else
            n = n + 1;
            continue
        end
        
        y(n,6) = {sprintf('%0.3f',p)};
        
        y(n,1) = ntwsnames(i);
        y(n,2) = {ngm};
        
        if ismember(phase, {'training','retraining','reversal_training'})
            y(n,4) = {Days(j)};
        elseif ismember(phase, {'probe_test'})
            y(n,4) = {''};
        end
        
        n = n + 1;
    end
end

y = cell2table(y);
y.Properties.VariableNames = {'Feature', 'Network', 'Test' 'Day', 'stats', 'p', 'ES'};
y = sortrows(y,{'Network','Feature','Day'});


%%
writetable(y, [parameters.PrjDir,'\ntw_stats.csv'], 'WriteVariableNames',1, 'delimiter',',')

if length(groups)>2
    yy = cell2table(yy);
    yy.Properties.VariableNames = {'Feature', 'Network', 'Test'...
        'Comparison' 'Day', 'stats', 'p', 'ES'};
    yy = sortrows(yy,{'Network','Feature','Day'});
    writetable(yy, [parameters.PrjDir,'\ntw_stats_mltcmp.csv'], 'WriteVariableNames',1, 'delimiter',',')
end
```
