## Extended Data 1 for "Scale space calibrates present and subsequent spatial learning in Barnes maze in mice": stats_str_v0.m

```
function [] = stats_str_v0(strObj,parameters)


%% load data
names = fieldnames(strObj);
for i = 1:length(names)
    iT = getfield(strObj(i),names{i});   
    eval([names{i}, '=iT;']);
end

Data = tbl_trn_daily;
strs = Data.Properties.VariableNames(4:end);
Days = unique(Data.Day);
[groups,~,ic] = unique(Data.Group);


y = cell(length(strs)*length(Days),6);
yy = [];
n = 1;
for i = 1:length(strs)
    for j = 1:length(Days)
        I = Days(j)==tbl_trn_daily.Day;
        x = Data{I, strs{i}};
        
        if length(groups)==2
            [p,~,stats] = ranksum(x(ic(I)==1),x(ic(I)==2),'method','approximate');
            r = stats.zval/sqrt(length(ic(I)));
            y(n,4) = {sprintf('z = %0.2f',stats.zval)};
            y(n,6) = {sprintf('r = %0.2f',r)};
            y(n,2) = {'Wilcoxon rank-sum test'};
        elseif length(groups)>2
            [p,statstable,~] = kruskalwallis(x, ic(I), 'off');
            y(n,4) = {sprintf('X^2 (%d) = %0.2f',length(groups)-1,statstable{2,5})};
            y(n,2) = {'Kruskall-Wallis test'};
            
            C = nchoosek(1:length(groups),2);
            yy_k = cell(size(C,1),7);
            for k = 1:size(C,1)
                [pp,~,stats] = ranksum(x(ic(I)==C(k,1)),x(ic(I)==C(k,2)));
                r = stats.zval/sqrt(sum(ismember(ic(I),C(k,:))));
                yy_k(k,1) = strs(i);
                yy_k(k,2) = {'Wilcoxon rank-sum test'};
                yy_k(k,3) = {sprintf('%s vs. %s', groups{C(k,1)},groups{C(k,2)})};
                yy_k(k,4) = {Days(j)};
                yy_k(k,5) = {sprintf('z = %0.2f',stats.zval)};
                yy_k(k,6) = {sprintf('%0.3f',pp)};
                yy_k(k,7) = {sprintf('r = %0.2f',r)};
            end
            yy = [yy; yy_k];
        else
            n = n + 1;
            continue
        end
        y(n,1) = strs(i);
        y(n,3) = {Days(j)};
        y(n,5) = {sprintf('%0.3f',p)};
        
        n = n + 1;
    end
end

y = cell2table(y);
y.Properties.VariableNames = {'Feature','Test','Day','stats','p','ES'};
y = sortrows(y,{'Feature','Day'});
writetable(y, [parameters.PrjDir,'\str_stats.csv'], 'WriteVariableNames',1, 'delimiter',',')

if length(groups)>2
    yy = cell2table(yy);
    yy.Properties.VariableNames = {'Feature', 'Test'...
        'Comparison' 'Day', 'stats', 'p', 'ES'};
    yy = sortrows(yy,{'Feature','Day'});
    writetable(yy, [parameters.PrjDir,'\str_stats_mltcmp.csv'], 'WriteVariableNames',1, 'delimiter',',')
end
```
