## Extended Data 1 for "Scale space calibrates present and subsequent spatial learning in Barnes maze in mice": summary_ntw_v0.m

```
function [] = summary_ntw_v0(parameters,options,tbl)

%%
fprintf('\nAnalyze network indices......DONE\n')
ntwObj = analyze_ntw_med_170215(parameters, tbl);

if options.order{'Network','Stats'}
    stats_ntw_v0(tbl,ntwObj,parameters)
end

if options.order{'Network','Output'}
    
    if ismember(unique(tbl.Phase),{'training','retraining','reversal_training'})
        sheetnames = {'training(daily_mean)', 'training(summarize)'};
    else
        sheetnames = {'probetest(daily_mean)', 'probetest(summarize)'};
    end
    
    filename = fullfile(parameters.PrjDir, 'NetworkScore.xls');
    names = fieldnames(ntwObj);
    empt = [];
    a = 0;
    
    for i = 1:length(names)
        
        Ti = getfield(ntwObj(i), names{i});
        
        if isstruct(Ti)
            names_Tj = fieldnames(Ti);
            for j = 1:length(names_Tj)
                Tj = getfield(Ti, {j}, names_Tj{j});
                a = a + 1;
                if isempty(Tj)
                    empt = [empt, a];
                else
                    dat = [Tj.Properties.VariableNames; table2cell(Tj)];
                    xlswrite(filename, dat, a)
                end
            end
            
        else
            a = a + 1;
            if isempty(Ti)
                empt = [empt, a];
            else
                dat = [Ti.Properties.VariableNames; table2cell(Ti)];
                xlswrite(filename, dat, a)
            end
        end
    end
    sheetnames(empt) = [];
    
    
    % excel ÌeV[gÌ¼OðÏX·é
    % http://goo.gl/8S2QXW
    e = actxserver('Excel.Application'); % # open Activex server
    ewb = e.Workbooks.Open(filename); % # open file (enter full path!)
    for i = 1:length(sheetnames)
        ewb.Worksheets.Item(i).Name = sheetnames{i}; % # rename 1st sheet
        ewb.Save % # save to the same file
    end
    ewb.Close(false)
    e.Quit
    
    fprintf('\nOutput......DONE\n')
else
    fprintf('\nOutput......SKIP\n')
end


if options.order{'Network','View'}
    FigMaker_ntw_v0(tbl, ntwObj, parameters, options)
    fprintf('\nVisualize......DONE\n')
else
    fprintf('\nVisualize......SKIP\n')
end
```
