## Extended Data 1 for "Scale space calibrates present and subsequent spatial learning in Barnes maze in mice": summary_str_v0.m

```
function [] = summary_str_v0(parameters,options,tbl)

fprintf('\nAnalyze strategy indices......DONE\n')
strObj = analyze_str_v0(tbl);

if options.order{'Strategy','Stats'}
    stats_str_v0(strObj,parameters)
end

if options.order{'Strategy','Output'}
    % output data of strategy indices
    filename = fullfile(parameters.PrjDir, 'StrategyScore.xls');
    names = fieldnames(strObj);
    ind = find(~cellfun(@nnz,strfind(names,'outlier')));
    names = names(ind);
    for i = 1:length(names)
        T = getfield(strObj(ind(i)), names{i});
        dat = [T.Properties.VariableNames; table2cell(T)];
        xlswrite(filename, dat, i)
    end
    % excel ÌeV[gÌ¼OðÏX·é
    % http://goo.gl/8S2QXW
    e = actxserver('Excel.Application'); % # open Activex server
    sheetnames = {'training(daily_sum)',...
        'training(summarize)'};
    ewb = e.Workbooks.Open(filename); % # open file (enter full path!)
    for i = 1:length(sheetnames)
        ewb.Worksheets.Item(i).Name = sheetnames{i}; % # rename 1st sheet
        ewb.Save % # save to the same file
    end
    ewb.Close(false)
    e.Quit
    fprintf('\nOutput......DONE\n')
else
    fprintf('\nOutput......SKIP\n')
end


if options.order{'Strategy','View'}
    FigMaker_str_v0(strObj,tbl,options,parameters)
    fprintf('\nVisualize......DONE\n')
else
    fprintf('\nVisualize......SKIP\n')
end
```
